# Functional Vagotopy in the Cervical Vagus Nerve of the Domestic Pig: Implications for the Study of Vagus Nerve Stimulation

**DOI:** 10.1101/856989

**Authors:** Megan L. Settell, Bruce E. Knudsen, Aaron M. Dingle, Andrea L. McConico, Evan N. Nicolai, James K. Trevathan, Erika K. Ross, Nicole A. Pelot, Warren M. Grill, Kenneth J. Gustafson, Andrew J. Shoffstall, Justin C. Williams, Weifeng Zeng, Samuel O. Poore, Luis C. Populin, Aaron J. Suminski, Kip A. Ludwig

## Abstract

Given current clinical interest in vagus nerve stimulation, there are surprisingly few studies characterizing the anatomy of the vagus nerve in large animal models as it pertains to on-and off-target engagement of local fibers. We sought to address this gap by evaluating vagal anatomy in the domestic pig, whose vagus nerve organization and size approximates the human cervical vagus nerve. We provide data on key features across the cervical vagus nerve including diameter, number and diameter of fascicles, and distance of fascicles from the epineural surface where stimulating electrodes are placed. We also characterized the relative locations of the superior and recurrent laryngeal branches of the vagus nerve that have been implicated in therapy limiting side effects with common electrode placement. We identified key variants across the cohort that may be important for vagus nerve stimulation with respect to changing sympathetic/parasympathetic tone, such as cross-connections to the sympathetic trunk. We discovered that cell bodies of pseudo-unipolar cells aggregate together to form a very distinct grouping within the nodose ganglion. This distinct grouping gives rise to a larger number of smaller fascicles as one moves caudally down the cervical vagus nerve. This often leads to a distinct bimodal organization, or ‘vagotopy’ that may be advantageous to exploit in design of electrodes/stimulation paradigms. Finally, we placed our data in context of historic and recent histology spanning mouse, rat, canine, pig, non-human primate and human models, thus providing a comprehensive resource to understand similarities and differences across species.

## 1. INTRODUCTION

The Food and Drug Administration (FDA) approved cervical vagus nerve stimulation (VNS) in 1997 as an adjunctive therapy in adults with partial onset epilepsy refractory to medications (Morris et al. 2013)(FDA, 1997). Subsequently, VNS was FDA-approved for the treatment of depression (Wheless, Gienapp, and Ryvlin 2018), and is in clinical trials for diverse conditions such as hypertension (Ng et al. 2016), heart failure (De Ferrari et al. 2017), rheumatoid arthritis (Koopman et al. 2016), tinnitus (Tyler et al. 2017) and stroke rehabilitation (Kimberley et al. 2018). Despite the growing clinical interest and some remarkable success in individual patients, VNS therapeutic effects are variable from patient to patient and are often limited by side effects including cough, throat pain, voice alteration and dyspnea (De Ferrari et al. 2017).

The most common clinical VNS electrode has two open helical cuffs, with each metal contact wrapping 270**°** around the exterior surface of the cervical vagus. Off-target activation of the neck muscles occurs at stimulation levels below or near average therapeutic parameters (Tosato et al. 2007; Yoo et al. 2013), possibly precluding activation of higher threshold parasympathetic efferents and/or baroreceptor afferents to and from the cardiopulmonary system (De Ferrari et al. 2017). These off-target effects have been attributed to activation of somatic fibers within the cervical vagus that eventually become the recurrent laryngeal branch, and potentially by current escaping the open helix electrodes to activate the somatic fibers of the nearby superior laryngeal branch (Tosato et al. 2007; Yoo et al. 2013). Given the off-target effects of VNS, there is renewed interest in developing novel stimulation strategies and multi-contact electrodes to stimulate selectively specific fibers within the vagus – typically preganglionic efferents or sensory afferents to and from the visceral organs – while avoiding motor efferent fibers coursing within or near the cervical vagus trunk (Plachta et al. 2014; Rozman and Bunc 2004; Thompson, Mastitskaya, and Holder 2019; Aristovich et al. 2019; Pečlin et al. 2009).

It is necessary to understand how well anatomical features in animal models approximate humans to optimize new electrode designs and stimulation strategies for eventual clinical deployment. The domestic pig may be an appropriate model for VNS given the similarity to humans with respect to autonomic control of cardiovascular function and diameter of the cervical vagus (Ding, Tufano, and German 2012; Hayes et al. 2013; Tosato et al. 2006; Trevathan et al. 2019). However, there are no studies characterizing the anatomical features of the pig cervical vagus that would impact on- and off-target engagement by VNS.

To address this gap, we used microdissection and post-mortem histology to quantify anatomical features of the cervical vagus nerve of the domestic pig including nerve diameter, number of fascicles, diameter of fascicles and distance of fascicles from the epineural surface where VNS electrodes are placed. Additionally, we characterized the locations of the superior and recurrent laryngeal branches of the vagus in relation to common electrode placement, which are pertinent to therapy limiting side effects. These data are placed in the context of both historic and recent histological data from a variety of animal models commonly used in the evaluation of VNS.

## 2. MATERIALS AND METHODS

### 2.1. Subjects and Surgical Methods

All study procedures were conducted under the guidelines of the American Association for Laboratory Animal Science (AALAS) in accordance with the National Institutes of Health Guidelines for Animal Research (Guide for the Care and Use of Laboratory Animals) and approved by Mayo Clinic Institutional Animal Care and Use Committee. The subject group included eleven (5 male, 6 female) healthy domestic swine (38.10 kg ± 2.67, mean ± SD). Animals were housed individually (21°C and 45% humidity) with ad libitum access to water and were fed twice a day. Each subject was given an injectable induction anesthesia; telazol (6 mg/kg), xylazine (2 mg/kg), and glycopyrrolate (0.006 mg/kg). An intramuscular injection of buprenorphine was given as an analgesic (0.03 mg/kg). An intravenous catheter was placed in the peripheral ear vein, followed by endotracheal intubation. The femoral artery was catheterized and instrumented with a pressure catheter (Millar, Inc., Houston, TX, Model # SPR-350S) for hemodynamic monitoring. Subjects were maintained with a mechanical ventilator using 1.5-3% isofluorane, and vital signs, including heart rate and temperature, were monitored every 15 minutes.

In a dorsal recumbence position, a ventral incision (15-20 cm) was made 3 cm lateral and parallel to midline starting at the level of the mandible. Tissue was divided to expose the carotid sheath and incised to expose the carotid artery (CA), internal jugular vein (IJV) and vagus nerve (VN). The area of interest on the cervical vagus nerve was cleared of surrounding tissues, taking care not to disturb small nerve branches. In a separate functional experiment not described here, animals were instrumented with a clinical LivaNova stimulating cuff electrode (LivaNova, Houston, TX). To best approximate the human implant procedure, placement of the LivaNova electrode was guided by a neurosurgeon who performs clinical VNS surgeries. Placement was caudal to the nodose ganglion, approximately 0.5-0.7 cm from the bifurcation of superior laryngeal nerve (SLN) for the cranial lead and 1.0-1.2 cm for the caudal lead. After installation, the electrode was functionally tested to ensure proper placement by measuring stimulation-induced decreases in heart rate, Hering-Breuer response, or neck muscle contraction. The incision site was kept moist with 0.9% sterile saline until the completion of experiment.

### 2.2. Microdissection Methods

Following the stimulation experiment, animals were euthanized using sodium pentobarbital (390 mg/ml) and underwent microdissection to expose further the vagus nerve from the cervical level (nodose ganglion) and the SLN to just caudal to the recurrent laryngeal (RLN) bifurcation (Figure 2). Care was taken to minimize disruption of any branching between the vagus and the sympathetic trunk. The SLN was identified as the nerve emerging medial and slightly ventral from the nodose and extending to the thyroid cartilage, splitting into the external superior laryngeal (ESL) and internal superior laryngeal (ISL). The ISL was identified as extending medially and ventrally, following the upper level of the thyroid cartilage, running parallel to the laryngeal artery. The ESL was identified as running parallel to the esophagus and terminating in the cricothyroid muscles (Bacchi, Miani, and Piemonte 1990; Cernea et al. 1992; Cheruiyot et al. 2018; Kierner, Aigner, and Burian 1998; Knight, McDonald, and Birchall 2005; Kochilas et al. 2008; Orestes and Chhetri 2014; Ozlugedik et al. 2007). The trunk of the vagus nerve was then exposed caudally from the nodose to the RLN by following the carotid artery and identifying the RLN as passing under the aortic arch (left side) or subclavian artery (right side). The RLN was identified as extending up from either the aortic arch or subclavian artery to the cricoarytenoid muscles. More cranially it forms an anastomosis with the ISL (Bowden 1955; Knight, McDonald, and Birchall 2005; Monfared et al. 2001).

**Figure 1:**
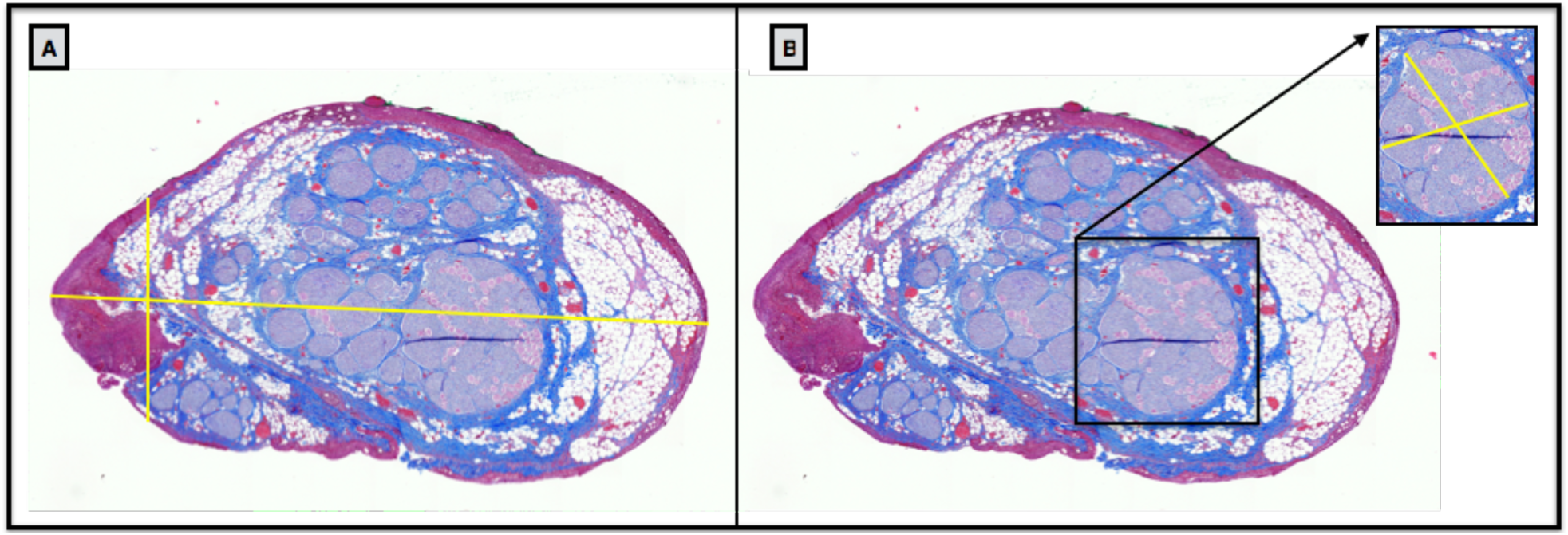
Example measurements from histological cross section of the cervical vagus nerve of a pig. A) Two measurements were taken of the nerve diameter, due to the irregular shape of nerve sections: one at the widest point and one at the narrowest, illustrated by the yellow cross hairs. B) The diameters of the largest fascicle were measured in a similar manner, with the largest and narrowest diameters measured.

**Figure 2:**
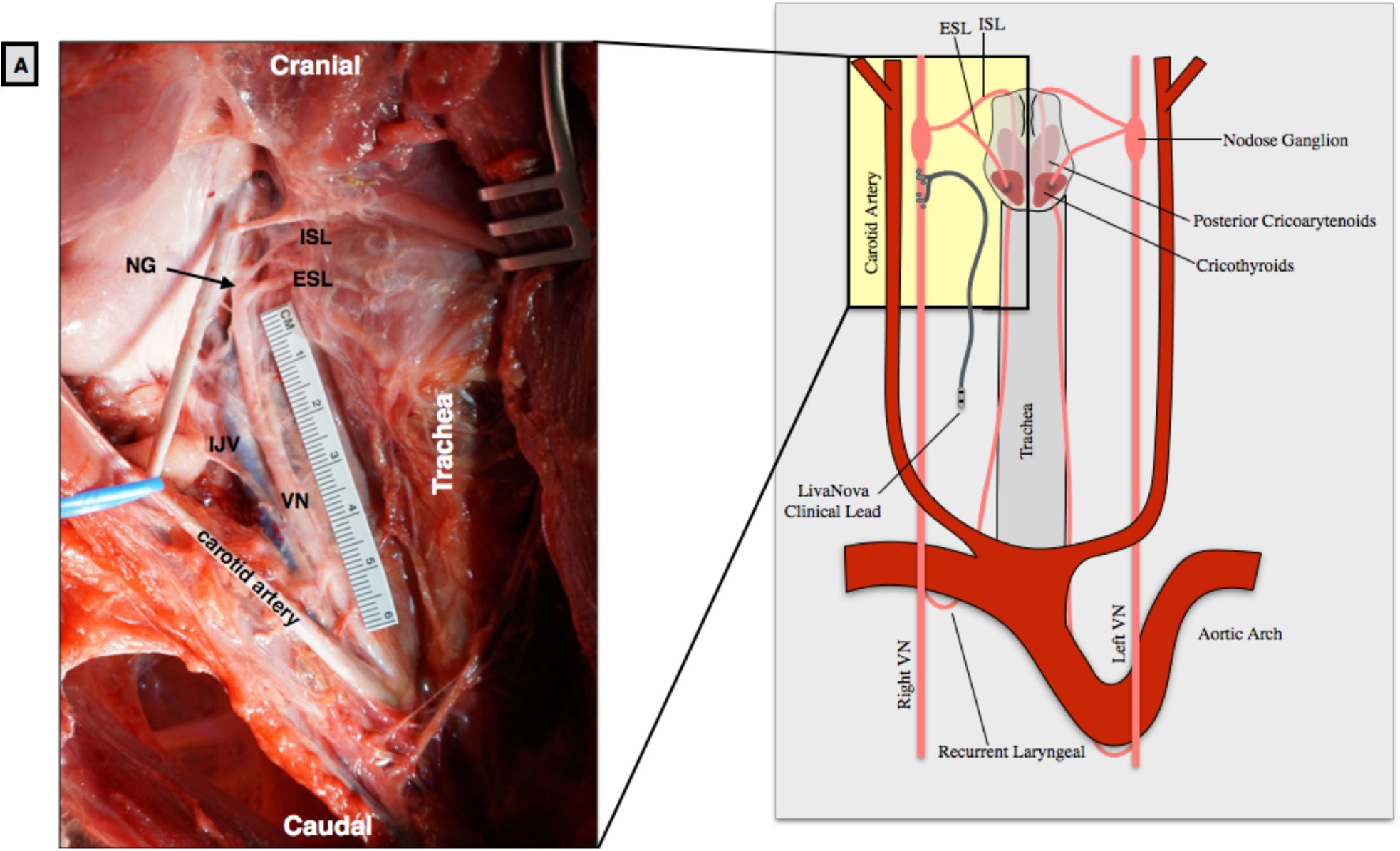
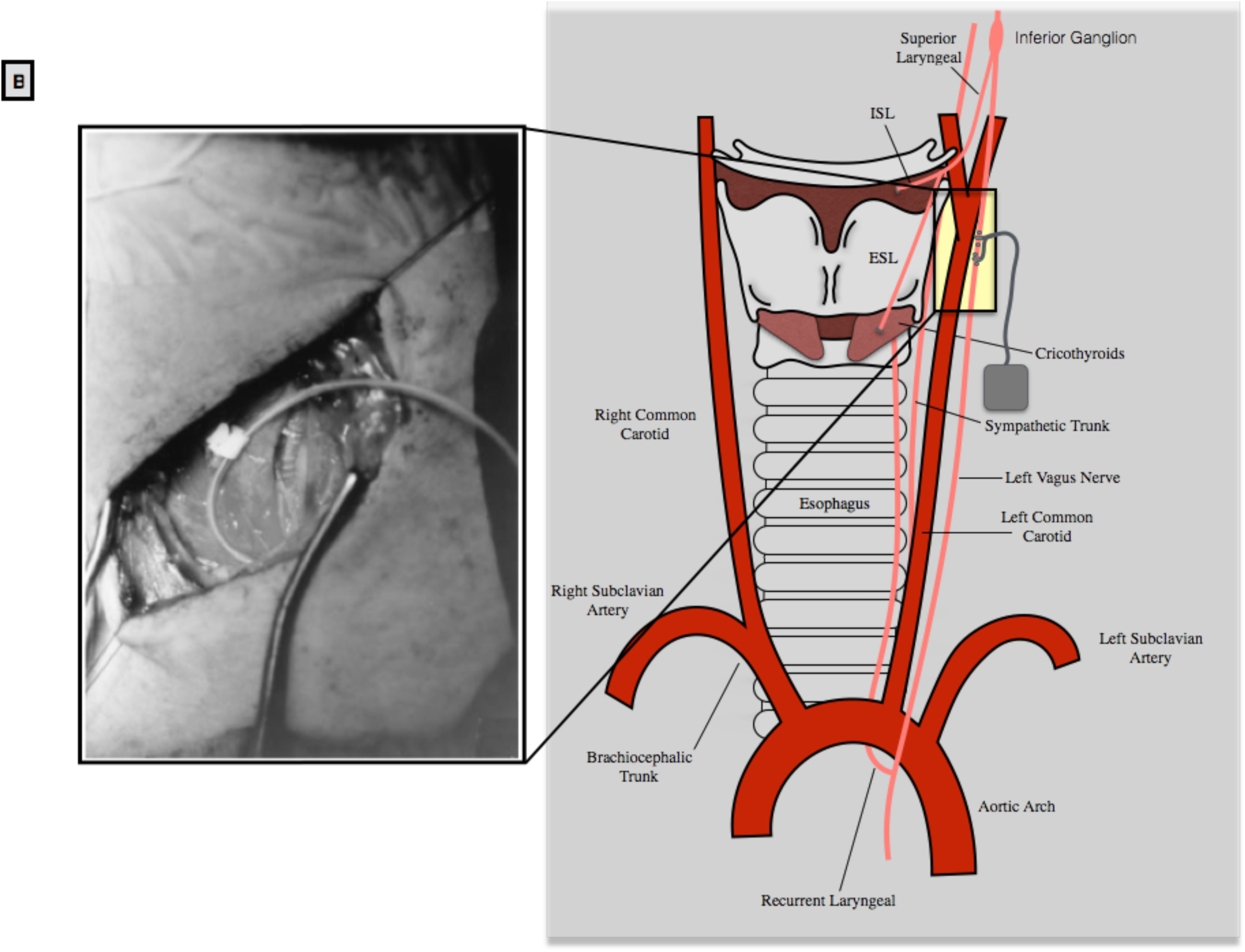
(A) The swine surgical window in the pig includes the inferior (nodose) ganglion, and by extension the superior laryngeal bifurcating nearby into the internal superior laryngeal (ISL) and external superior laryngeal (ESL). The clinical LivaNova electrode is placed approximately 0.5-0.7 cm from the bifurcation of the superior laryngeal nerve for the cranial lead and 1.0-1.2 cm for the caudal lead within the surgical window. Note the presence of the recurrent laryngeal running parallel to the trachea and esophagus, near the electrode. (B) In the human surgical window, the inferior (nodose) ganglion is more cranial, and not included in the surgical window, though the superior laryngeal branch extends down into the window. The carotid bifurcation in both the human and pig is located near the stimulating electrode, though the vagus nerve (VN) runs medial to the carotid artery in the pig, and lateral in the human. Human surgical window (B) reprinted from (Patil, Chand, and Andrews 2001) with permission from Elsevier, **©** 2013.

### 2.3. Histological Analysis of Fascicle Organization and Diameter

#### Porcine Histology

Histology dye (Bradley Products, Inc., Davidson Marking System, Bloomington, MN) was placed along the ventral and lateral edge of the nerve of interest (VN, ESL, ISL, RL) to maintain orientation and nerve sections of each were placed in 10% neutral buffered formalin for ∼24 hours at 4°C. The VN section was taken from just cranial to the nodose ganglion, including a small section of SLN to be embedded with the nodose ganglion, to just caudal of the RLN bifurcation. Samples were then placed in a Research and Manufacturing Paraffin Tissue Processor (RMC Ventana Renaissance PTP 1530, Ventana Medical Systems, Oro Valley, AZ) and underwent a series of standard processing steps: 1) dehydration (ethanol, 50-100%), 2) clearing (xylene), and 3) infiltration (paraffin) (Feldman and Wolfe 2014). Sections were then embedded in paraffin wax and allowed to set. Each block was placed in an ice water bath for approximately one hour to rehydrate the tissue and allow 5 µm sections to be cut using a Leica Biosystems Rotary Microtome (Buffalo Grove, Illinois) and stained using Gomori’s trichrome. Slides were imaged using a Motic Slide Scanner (Motic North America, Richmond, British Columbia) at 20x. Image analysis was performed using ImageJ software. Using the “straight line tool”, each slice of interest was measured at both the widest and narrowest portions for diameter (Figure 1 and supplemental materials). The largest fascicle was measured in a similar manner, and the number of fascicles was manually counted.

#### Mouse, Rat and Non Human Primate Histology

The cervical vagus nerves of rats, mice and non-human primates were removed following anatomical dissections and prepared as previously described (Dingle et al. 2019). Briefly, nerves were fixed in 10% neutral buffered formalin at 4 °C overnight. Each nerve was cut at the mid-point along the length and both halves were embedded in paraffin. Sections in both the caudal and cranial directions were obtained simultaneously and mounted. Whole slides were scanned at 40x magnification using a PathScan Enabler IV (Meyer Instruments, Houston, TX). Image analysis was performed using ImageJ software.

#### Human Specimen Histology

Fresh, frozen, and de-identified human cephalic specimens (to T2-T3) were obtained through Science Care (Phoenix, AZ). A lateral midline incision was made in the neck from the front of the ear to the distal edge of the sample (approximately at C2). The platysma muscle was cut to expose the sternohyoid muscle and the sternocleidomastoid muscle. Blunt dissection was used to locate the anterior edge and separate sternocleidomastoid muscle from the sternohyoid muscle, and to expose and identify the internal jugular vein, vagus nerve, and common carotid artery. Blunt dissection was used to expose the vagus nerve from the vein and artery, with care taken to avoid disturbing any surrounding branches. The pharyngeal branch of the vagus nerve and superior laryngeal nerve were also identified.

The pharyngeal branch of the vagus nerve ran across the surgical window, and cranial to the common carotid artery and terminated in the superior pharyngeal constrictor muscle. The superior laryngeal nerve ran across the incision, under the common carotid artery and terminated in the cricothyroid muscles. The vagus nerve was dissected up to the base of the skull and the styloid process was removed. The vagus nerve sample for histology was taken just after exiting the foramen, at the base of the skull to the C2 level. The sample did not include the recurrent laryngeal bifurcation, or the cervical cardiac branch of vagus nerve. The sample was stained and processed as described in the sections above, with this sample being cut into seven (2 cm) sections, and fixed in 10% neutral buffered formalin, before being processed. Samples were then cut into 6 mm segments, embedded, and mounted.

## 3. RESULTS

At the level of the vagus nerve stimulating electrode, the average diameter of the cervical vagus of the domestic pig was 2.45 mm at the most narrow point and 3.53 mm at the widest point (SD 0.52 and 1.01, respectively), making the porcine vagus nerve an appropriate human analog. Through microdissection and histological analysis, we determined that there was variability in several key anatomical features: 1) branching patterns of nerves exiting the VN, 2) non-vagal nerves integrated with the trunk of the vagus nerve, i.e. ‘hitch-hiking’ nerves, and 3) a predominantly bimodal anatomical organization at the level of the cervical vagus between fascicles putatively arising from sensory afferents from the organs and the remaining fibers within the vagus.

### 3.1. Variations in the Surgical Window

Within the surgical window in the pig, the sternocleidomastoid was retracted laterally, and the carotid sheath opened to expose the vagus nerve, carotid artery (CA) and bifurcation, internal jugular vein (IJV) nodose ganglion, and SLN (Figure 2). Running parallel to the vagus nerve the sympathetic trunk was exposed and any cross connections to the vagus nerve were identified. Unlike the human where the nodose (inferior) ganglion is outside the surgical window near the jugular foramen, the nodose ganglion was notably more caudal in the pig and identifiable within the surgical window (Figure 2).

### 3.2. Variations in Branching

The cervical vagus extended from the nodose ganglion caudally, and the ESL and ISL projected from the nodose ganglion medially to the esophagus. The ISL projected medial and slightly ventral and followed the cranial margin of the thyroid cartilage, parallel to the laryngeal artery, to the insertion point, just above the thyroid cartilage. The ESL projected medially and then once at the level of the esophagus, projected caudally along the trachea to the cricothyroid muscle.

The first inter-subject variation in anatomy was in how the ESL and ISL emerged from the nodose ganglion (Figure 3). In some subjects we observed that the ESL and ISL first formed a single nerve bundle and then bifurcated into the ISL and ESL. In other subjects, the ESL and ISL exited the nodose ganglion as two separate nerve bundles, and both of these arrangements confirmed the findings of Hayes et al. (Hayes et al. 2013) (Figure 3).

**Figure 3:**
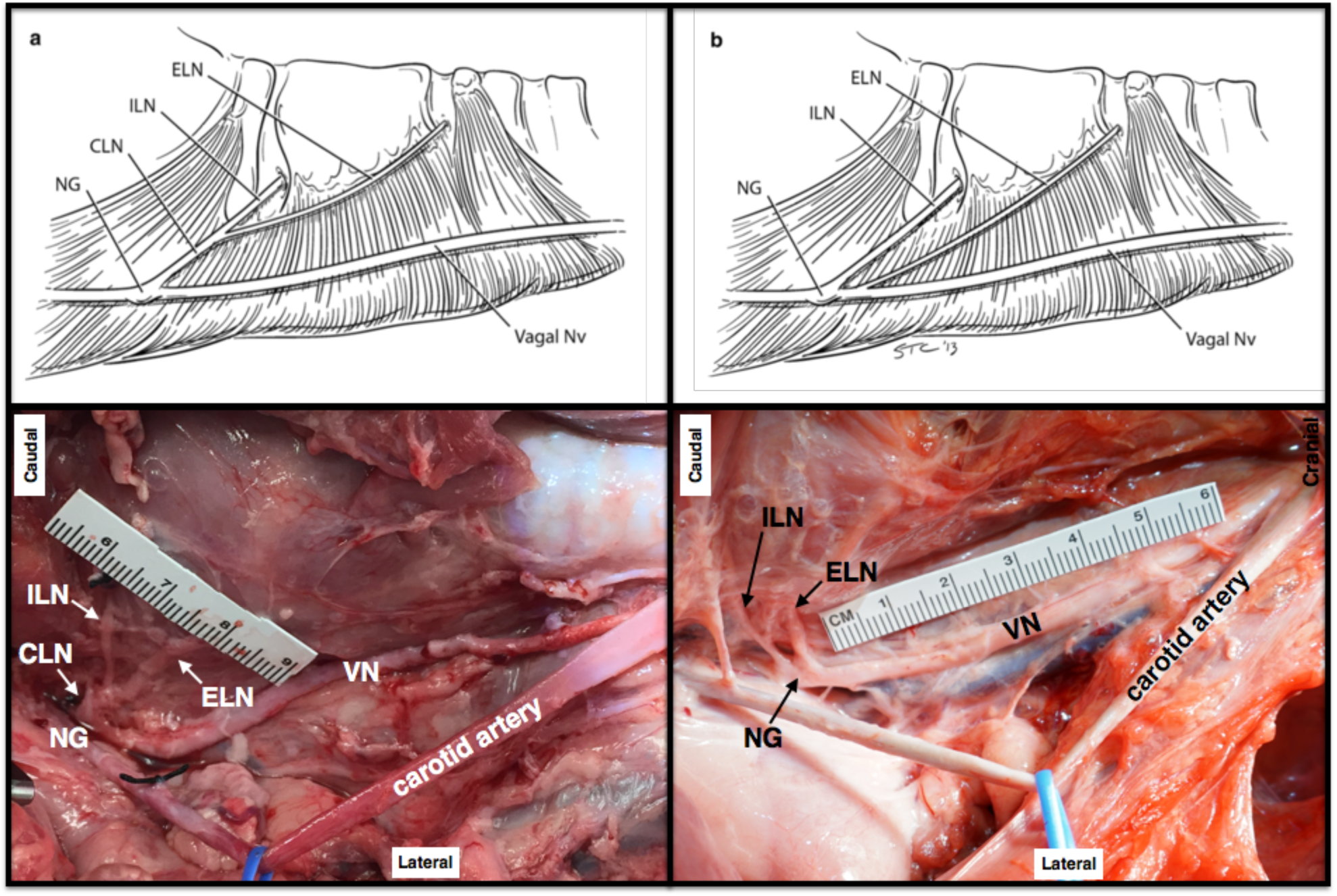
Depiction of two branching patterns of the superior laryngeal branch into the external and internal superior laryngeal branches. a) The superior (cranial) laryngeal nerve (CLN) extending from the nodose ganglion (NG) as a single process, and bifurcating into the external superior laryngeal nerve (ELN) and internal superior laryngeal nerve (ILN); b) The ELN and ILN protruding from the NG as separate processes. Top row reprinted from (Hayes et al. 2013) with permission from Springer Nature, **©** 2013.

The second location with variation in anatomy was the length of the recurrent laryngeal from its bifurcation from the vagal trunk to its insertion point into the muscle (Table 2, Figure 4). The average length of the RLN to the insertion in the muscle was longer on the left side (15.54 cm ± 3.11 SD) than the right (9.58 cm ± 1.31 SD) as expected given that the path of the RLN on the left side courses under the aortic arch. The length of the SLN (internal) from branching to muscle insertion was longer on the left side (3.88 cm ± 1.25 SD) than the right (2.81 cm ± 0.49 SD). Finally, the length of vagal trunk from the SLN to RLN on the left side (20.54 cm ± 1.32 SD) was longer than right (14.55 cm ± 0.74 SD).

**Table 2:** Nerve length measurements across the cohort; vagal trunk from superior laryngeal bifurcation to recurrent laryngeal bifurcation (SLN to RLN), recurrent laryngeal bifurcation from the vagal trunk to insertion into the muscle (RLN to Insertion), and superior laryngeal bifurcation from the nodose ganglion to insertion (SLN to Insertion). Figure 3 illustrates the location of measurement for each of these nerve lengths. The first three subjects (1-3) underwent bilateral microdissection to obtain measurements, and each subject thereafter (4-11) underwent unilateral microdissection for measurements. *The SLN to insertion for subject three may be an outlier, potentially due to significant stretching of the superior laryngeal during microdissection. Nerve length measurements were not taken for subject 11.

|  |  | Left Vagus Nerve |  |  | Right Vagus Nerve |  |  |
| --- | --- | --- | --- | --- | --- | --- | --- |
| Subject | Sex | SLN to RLN | RLN to Insertion | SLN to Insertion | SLN to RLN | RLN to Insertion | SLN to Insertion |
| 1 | F | 19.8 | 16.5 | 3.4 | 15.1 | 10.4 | 3.0 |
| 2 | F | 19.0 | 10.2 | 3.0 | 14.0 | 10.0 | 3.3 |
| 3 | M | 22.5 | 16.7 | 6.0* | 15.4 | 10.2 | 3.4 |
| 4 | M | - | - | - | 13.2 | 9.1 | 3.0 |
| 5 | M | - | - | - | 15.2 | 11.4 | 2.0 |
| 6 | F | 21.0 | 16.0 | 3.0 | - | - | - |
| 7 | M | 20.4 | 18.3 | 4.0 | - | - | - |
| 8 | M | - | - | - | 14.6 | 8.8 | 2.5 |
| 9 | F | - | - | - | 14.8 | 7.0 | 3.0 |
| 10 | F | - | - | - | 14.1 | 9.7 | 2.3 |
| **Average** |  | 20.54 | 15.54 | 3.88 | 14.55 | 9.58 | 2.81 |
| **Standard Deviation** |  | 1.32 | 3.11 | 1.25 | 0.74 | 1.31 | 0.49 |

All measurements in centimeters

**Figure 4:**
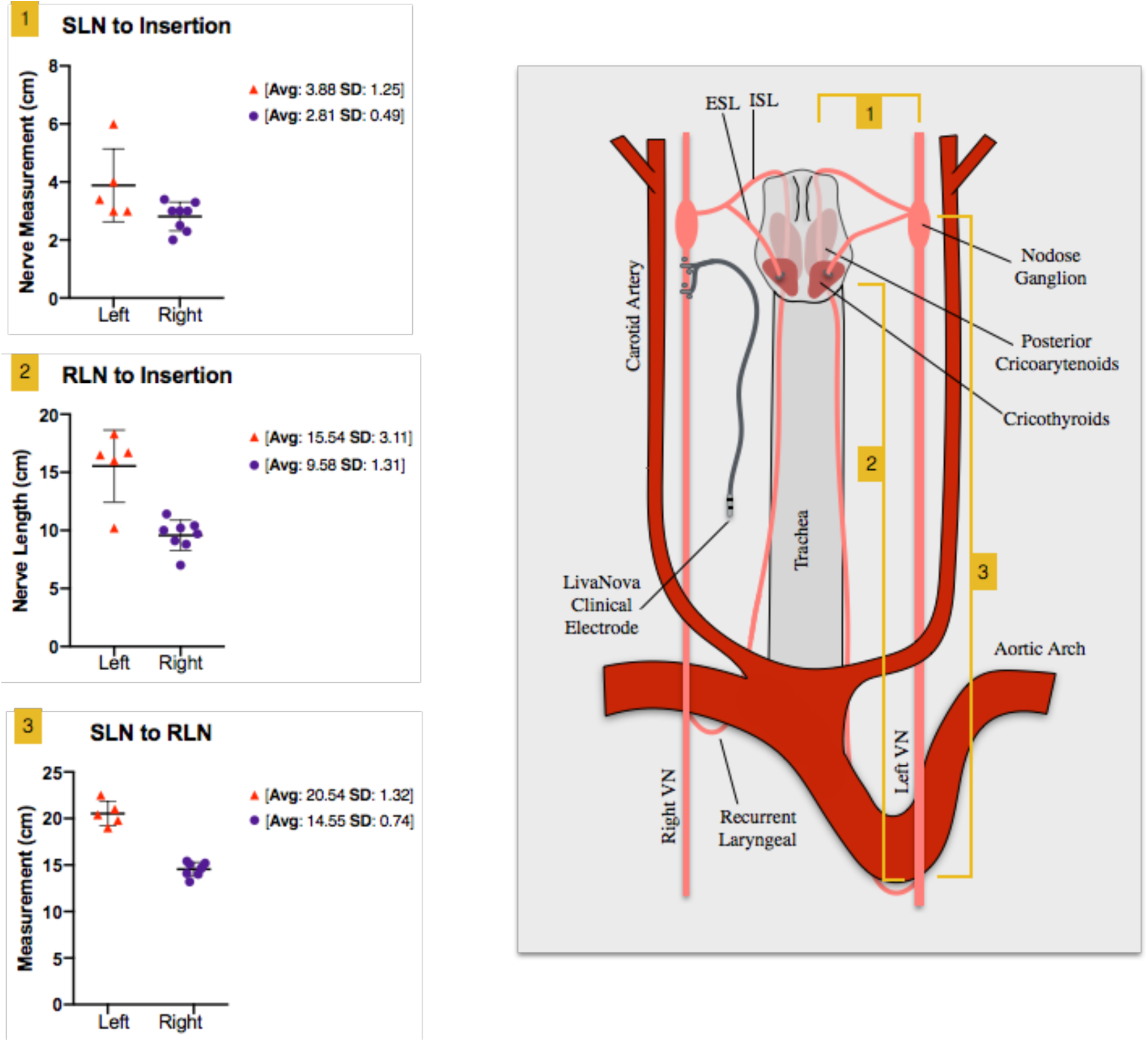
Individual nerve length measurements (Table 2) with corresponding measurement locations. Nerve lengths were measured along the nerve by laying a vessel loop along the nerve and measuring the length of the vessel loop on a ruler. The left and right group means are given with standard deviations for each of the three measurements 1) superior laryngeal (SLN) to insertion, 2) recurrent laryngeal (RLN) to muscle, and 3) SLN to RLN.

In contrast to the locations of the RLN and SLN branches with respect to the cervical vagus trunk, which may impact unwanted neck muscle activation, the location of the sympathetic trunk and any cross-connections from the trunk to the vagus nerve introduce variability in stimulation-induced changes in sympathetic/parasympathetic tone mediated by the baroreceptors/chemoreceptors. There are three key variables that may impact how well the stimulating electrode isolates the cervical vagus contributions to sympathetic/parasympathetic tone: 1) location of the sympathetic trunk, 2) cross-connections between the cervical vagus and the sympathetic trunk or its cardiac branches, and 3) the location of the aortic depressor nerve. These additional pathways are important because in addition to a ‘depressor’ response causing a reduction of heart rate/blood pressure, these pathways generate a ‘pressor’ response at certain stimulation amplitudes and frequencies (Peterson and Brown 1971; Randall and Rohse 1956; Schmidt 1968). The human cervical sympathetic trunk is often located within the carotid sheath (Kiray et al. 2005). We noted that in some cases, the sympathetic trunk was intimately joined to the vagus in certain areas, where they could not be separated during microdissection, both along the length, and at the nodose ganglion (Figure 5).

**Figure 5:**
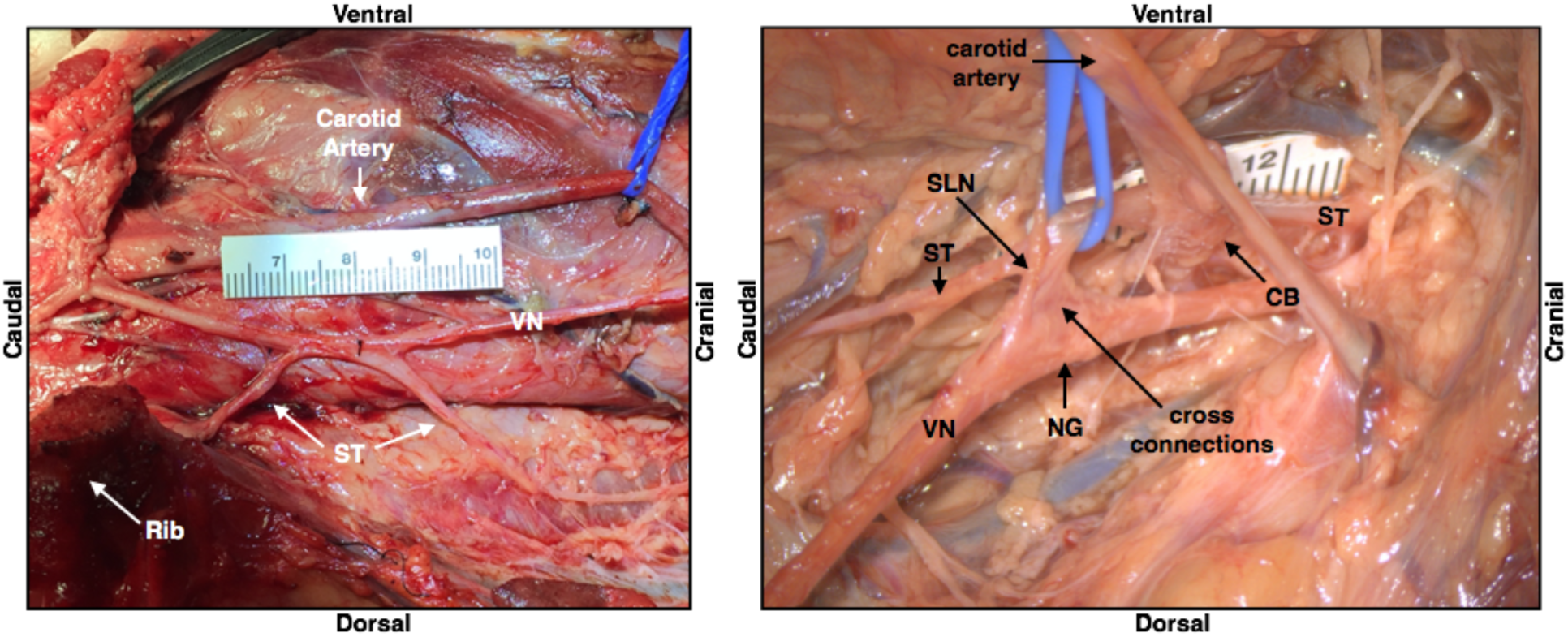
Depiction of the sympathetic trunk (ST) running parallel to the vagus nerve (VN) in one subject, and also “hitch-hiking” along sections of the nerve including cross-connections to the nodose ganglion (NG); superior laryngeal (SLN), carotid bifurcation (CB), vagus nerve (VN).

In some cases, we observed a separate small nerve adjacent to the vagus during microdissection, or apparent in cross-sectional histology integrated into the cervical vagus trunk, consistent with previous descriptions of the location of the aortic depressor nerve in pig (Schmidt 1968). However, we also observed this small nerve trunk on the right side in a portion of the histological cross sections (Figure 7).

**Figure 6:**
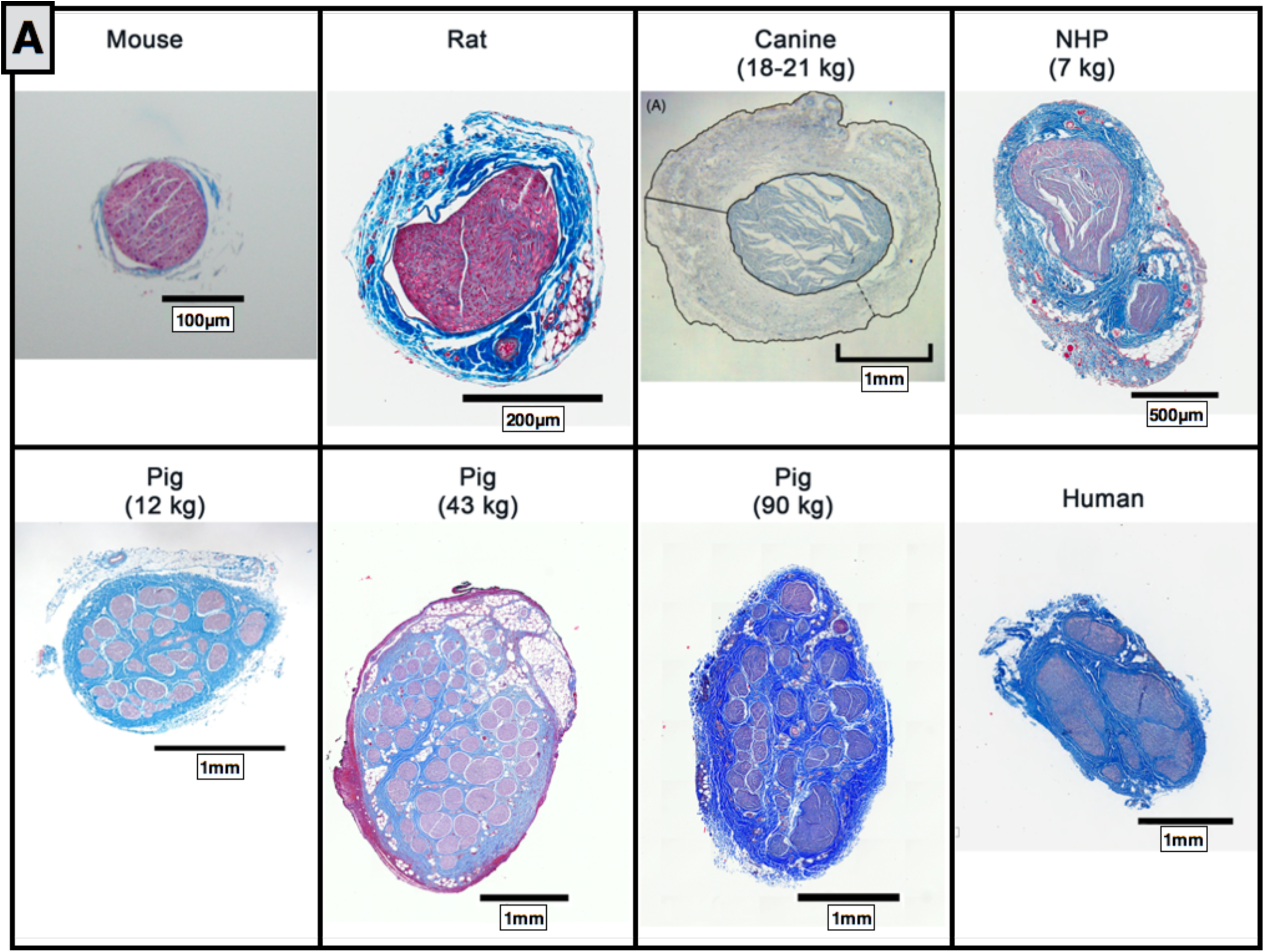
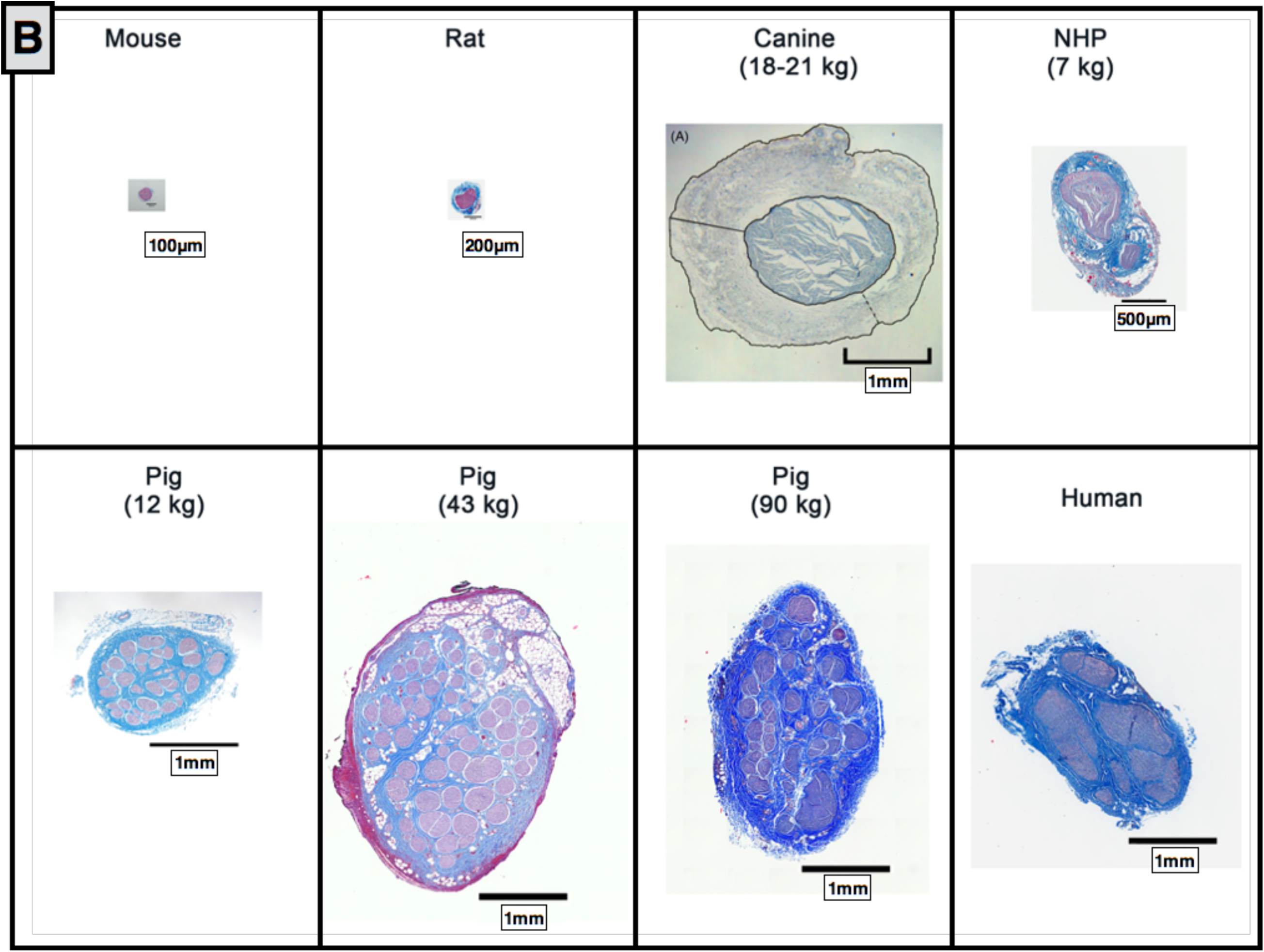
A) Comparative anatomy of the cervical vagus nerve between mouse, rat, canine, non human primate (NHP), pig (12 kg, 43 kg, and 90 kg) and human. B) Comparative anatomy at relative sizing. Canine histology reprinted from <u>(Yoo et al. 2013)</u> with permission from IOP Publishing, **©** 2013.

**Figure 7:**
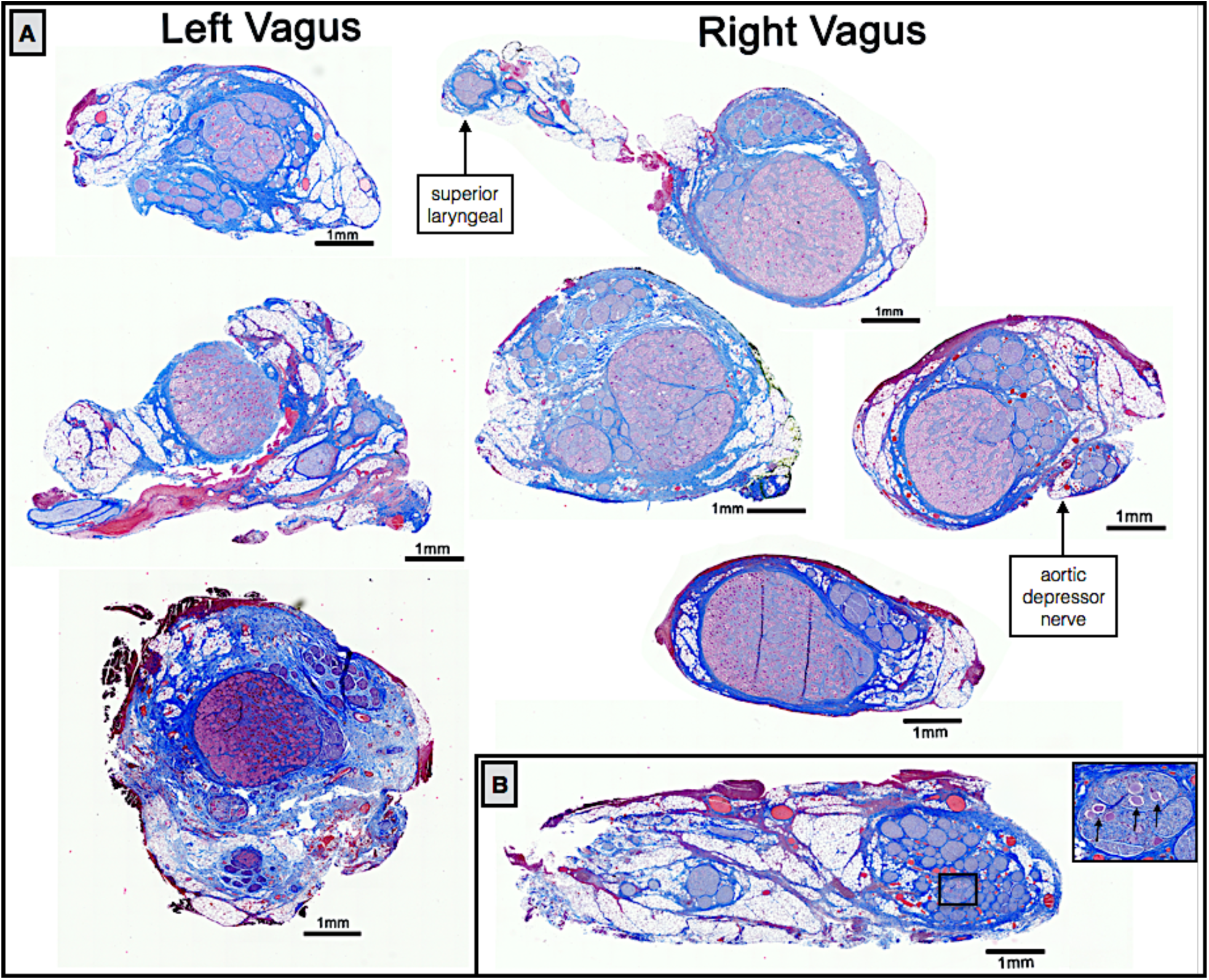
Examples of left and right side vagal nerve cross-sections showing the singular large fascicles of aggregated pseudo-unipolar cell bodies in a single plane when the nodose was sectioned extensively (A), whereas the aggregated pseudo-unipolar cells were not visible in early subjects, when nerves were sampled more sparsely, black arrows indicate a few sparse pseudo-unipolar cells (B).

### 3.3. Histological Analysis of the Vagus Nerve

The diameter, number of fascicles, average fascicle diameter, and closest distance from the epineural surface to the nearest fascicle varied greatly across species (Figure 6). The mouse, rodent, canine and NHP all had notably less complex fascicular organization than human, typically consisting of 1-2 fascicles. The canine had the thickest epineurium across all models, resulting in a greater distance from an electrode placed on the epineural surface to the nerve fibers, and presumably increasing activation thresholds (Yoo et al. 2013).

In addition to the mouse, rodent, canine, and NHP, samples were also processed from several domestic pigs weighing less than 20kg and weighing as much as 100kg (Figure 6). The cervical vagus nerve from pigs under 20kg had a notably smaller diameter than large pigs, but still had more fascicles than humans. No differences were observed in the diameter of the vagus nerve or number of fascicles between pigs ranging from 35-45 kg and their larger counterparts.

When scaled to relative size (Figure 6b), the pig nerves were the most representative of the human samples in terms of diameter of the vagus and distance from the epineural surface to the closest fascicle compared to the other animal models assessed. However, the pig had a greater number of fascicles than the human, and this may have significant implications in assessing electrode strategies to isolate specific physiological effects.

### 3.4. Vagotopy of the Cervical Vagus Nerve Swine

Vagal motor fibers pass through the nodose ganglion, while visceral afferent sensory fibers arise from pseudo-unipolar cells with their cell bodies located in the nodose ganglion (Câmara and Griessenauer 2015; Ellis 1964; Rea 2014). There was a clear organizational structure with respect to pseudo-unipolar cell bodies within the nodose ganglion. Fascicles that arose from a large consistent grouping of pseudo-unipolar cell bodies continued caudally along the cervical vagus to course within the VNS bipolar electrodes.

We initially sectioned the nodose samples sparsely (∼3-5 sections spanning the rostral/caudal nodose, approximately 5 mm between sections, 5 µm slices), and in some animals we identified a very large grouping containing an aggregation of classic pseudo-unipolar cell bodies, which we also refer to as a “fascicle” for simplicity (Figure 7 and supplemental material for additional animals). Cell bodies of pseudo-unipolar cells were identified by surrounding supporting satellite glia cells and by the nucleus of the soma (Haberberger et al. 2019; Ling and Wong 1988; Ohara et al. 2009). In other animals, we observed a large number of fascicles roughly organized into a grouping, in which the occasional pseudo-unipolar cell body was visible (Figure 7B and supplemental materials for additional animals). In subsequent animals (n=10), we increased the density of our cross sections along the nodose to evaluate whether the smaller fascicles merged to form the single large “fascicle” containing a large number of pseudo-unipolar cell bodies in all animals. By increasing the density of our sampling, we identified this single large “fascicle” containing a large aggregation of pseudo-unipolar cell bodies in either the right or left vagus in 6 of 10 of pigs (Figure 7). Two additional pigs (for a total of 8) were identified as having incomplete planes, most likely due to simply missing the exact “plane of distribution” or the plane with the large “fascicle” during sectioning. Additionally, we compared the vagus nerve of the domestic pigs to those of several (n=5) mini-pigs to determine if this plane was present across strains (supplementary materials).

In a subset of animals (n=10), sectioning was extended caudally covering the length of nerve within the VNS electrode and beyond the branching point of the RL (Figure 8). At sections just caudal to the nodose ganglion, there was often a distinct bimodal organization to the cross section of the nerve (n=7), consisting of a large grouping of fascicles emerging from the singular large “fascicle” containing pseudo-unipolar cell bodies, and one or more groupings of fascicles not arising from this pseudo-unipolar fascicle. Fascicles mostly maintained their grouped organization with respect to each other along the cranial/caudal axis (Figure 8). The secondary grouping of fascicles not arising from the pseudo-unipolar cell “fascicle” rotated medially towards the recurrent laryngeal branching point as sections progressed caudally. This distinct bimodal grouping then disappeared beyond the RL branching point.

**Figure 8:**
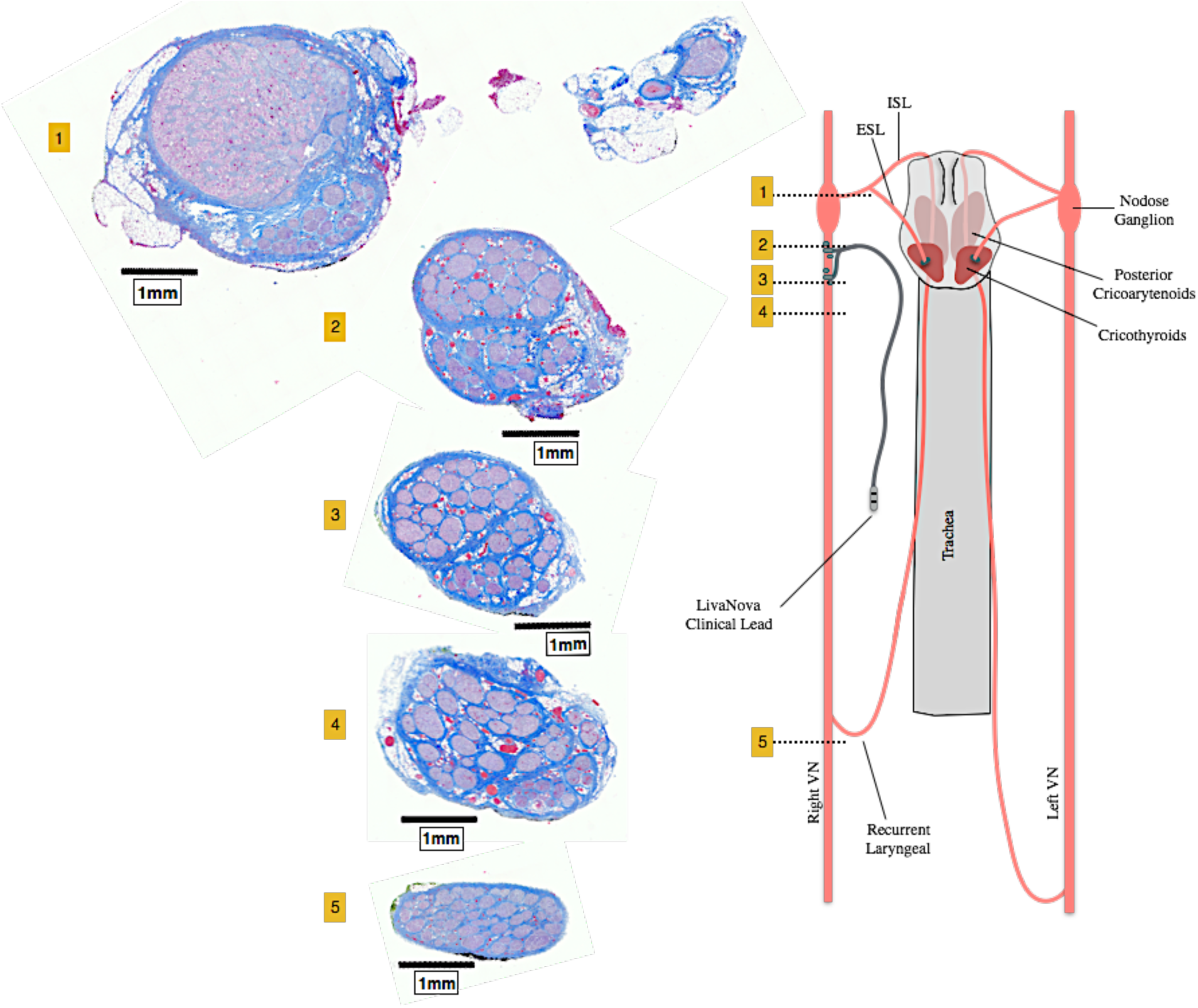
Histological sections taken at several locations along the length of the vagus nerve demonstrating bimodal organization or vagotopy. Section 1 was taken through the nodose ganglion and superior laryngeal branch, and contains the pseudo-unipolar cells aggregated in a large “fascicle” that gives rise to a distinct smaller grouping of fascicles in sections 2 through 4. After the recurrent laryngeal bifurcates from the vagus nerve trunk, the bimodal organization is no longer evident (section 5). Orientation of fascicle groupings was maintained and analyzed using a histological dye applied *in vivo.*

The diameter of the vagus, the number of fascicles, and the diameter of the largest fascicle at cross-sections obtained cranial to the nodose, at the nodose, and caudal to the nodose at the level of the bipolar electrodes, and at the RLN bifurcation are summarized in Table 3. Note that the widest average diameter of the largest fascicle at the level of the nodose was 2.19 mm (± 0.8 SD). This was much larger than both the average diameter of the largest fascicle in a more cranial vagal nerve section (0.65 mm ± 0.35 SD), and a more caudal vagal nerve section (0.38 mm ± 0.21 SD). Most likely due to the presence of the singular large “fascicle” containing the pseudo-unipolar cells as described above. In addition to the aforementioned bimodal grouping of fascicles, two other organized groupings of fascicles were frequently observed cranial to or at the level of the nodose. One additional fascicle grouping was identified as belonging to the SLN branch of the vagus, which was included in the cross section. The second grouping of fascicles, not attributed to the SLN branch, appeared to be part of its own small trunk. This small trunk was identifiable as a separate merged epineurium and/or separated by fatty tissue from the main trunk (Figure 7). The source of this latter grouping could be from 1) the aortic depressor nerve embedded within the larger cervical vagus trunk, cross connections from the vagus to 2) the sympathetic trunk or 3) the carotid sinus nerve. These cross-connections may have been unintentionally severed during microdissection and therefore missed in our gross observations.

**Table 3:**
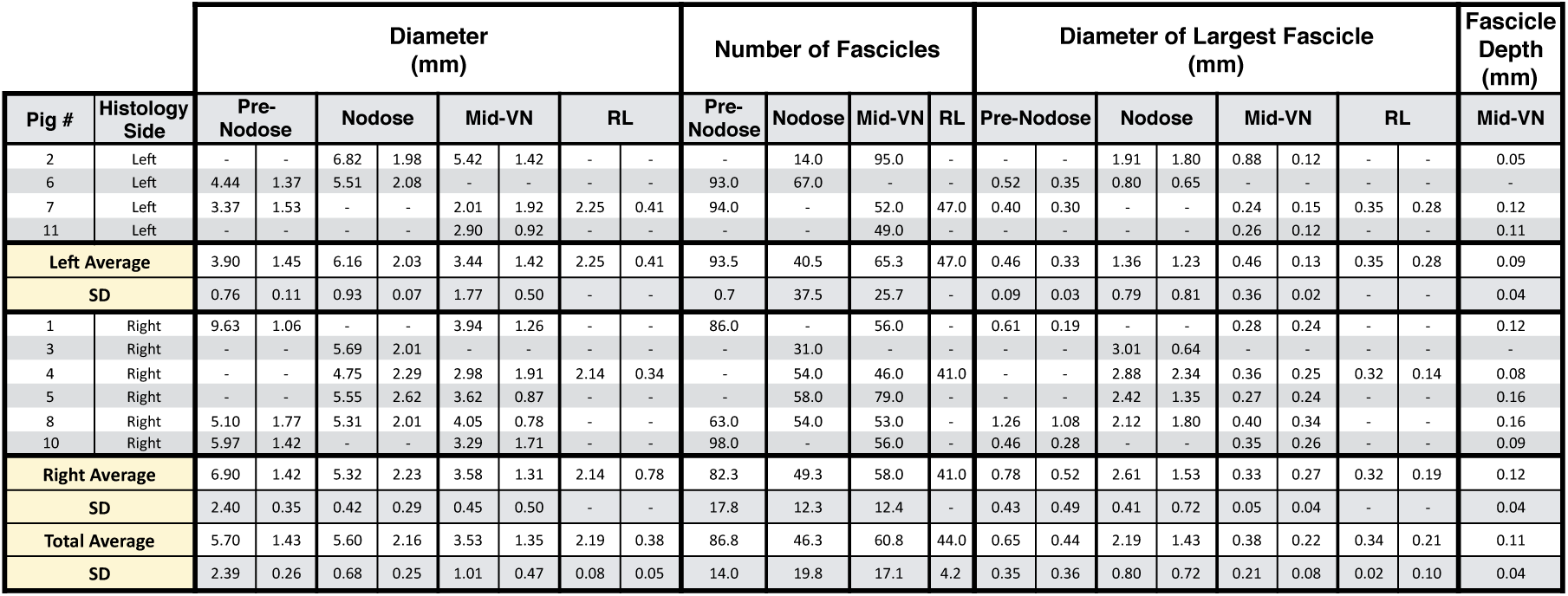
The diameter of the nerve, number of fascicles, and diameter of the largest fascicle were taken at multiple locations along the vagus nerve, at the region just cranial to the nodose ganglion (Pre-Nodose), at the nodose ganglion (Nodose), near the region where the LivaNova electrode was placed (Mid-VN) and near the recurrent laryngeal bifurcation (RL). Additionally, the distance from epineural surface to the edge of the closest fascicle (Fascicle Depth), in the region that runs within the electrode, was measured. Two measurements are given for diameter, the widest and narrowest diameter (see methods section), given that the nerve and fascicles were not perfectly round (column one and two, under each subheading). Histology side indicates from which vagus nerve (left or right) samples were sectioned. Dashes indicate suboptimal samples (torn during processing, folded during mounting, etc.), which could not be accurately measured.

To understand the relevance of the pig anatomy at the level of the nodose ganglion to the human condition, we conducted pilot studies using trichrome staining of the cervical vagus from human cadavers. Blocks were taken from just below the jugular foramen and sectioned and stained as described above. As in the pig, there was a singular large fascicle containing pseudo-unipolar cell bodies, whereas other fascicles did not contain pseudo-unipolar cell bodies (Figure 9) (Seki et al. 2014). There was a distinct inhomogeneity across the fascicle in distribution of fiber sizes. This includes a grouping of large diameter fibers approximately 10 µm in diameter consistent with Aα fibers commonly associated with motor fibers, and fibers approximately 1/10^th^ this diameter more consistent with Aδ, B and C fibers typically associated with mechanoreceptor afferents, parasympathetic efferents to the viscera, and unmyelinated afferent fibers.

**Figure 9:**
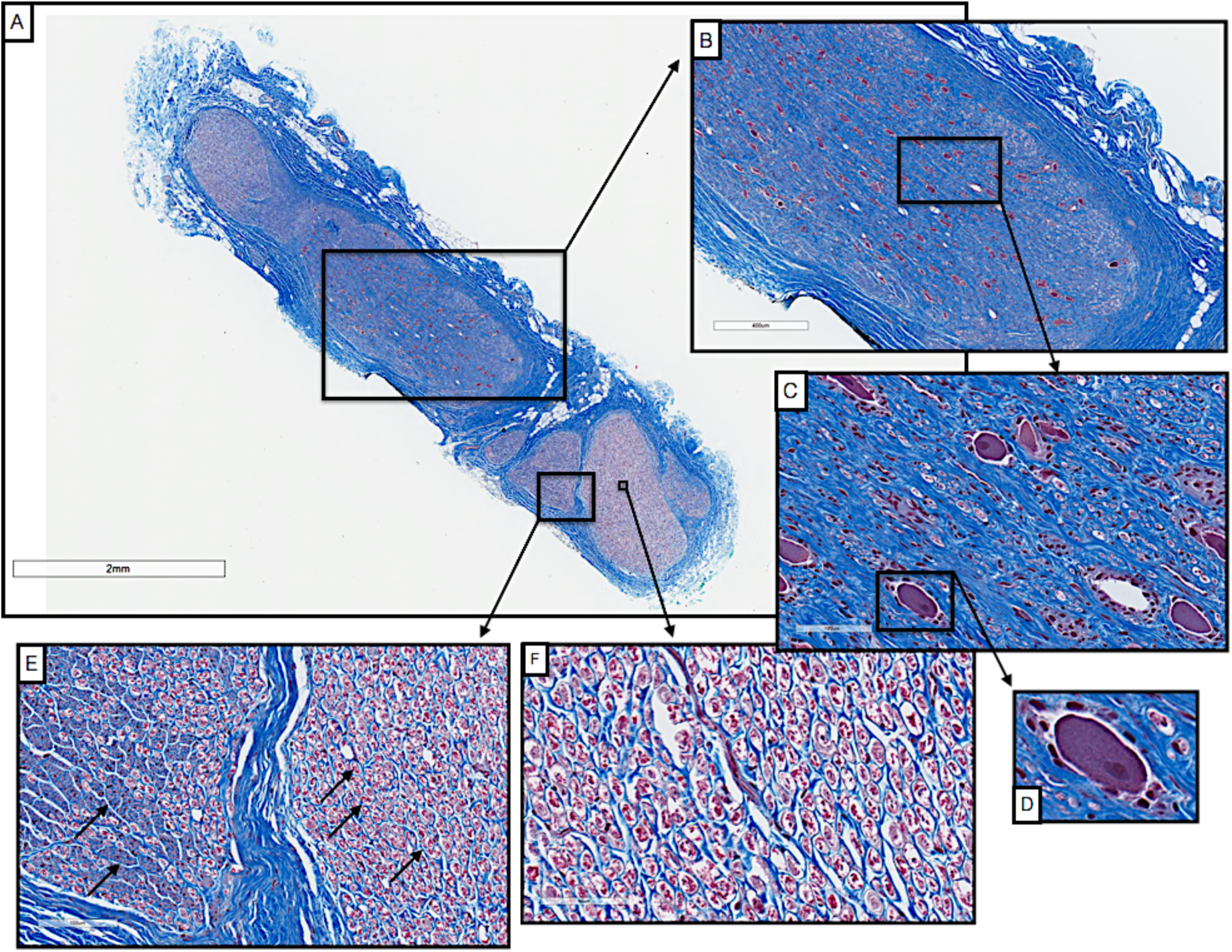
Cross-section of a human vagus nerve (n=1) at the level of the jugular foramen (A) with pseudo-unipolar cells evident in one large fascicle (B, C). Pseudo-unipolar cells were identifiable by the accompanying surrounding satellite cells (D). In the remaining fascicles there were no obvious pseudo-unipolar cells, but instead large diameter fibers (10 µm), and fibers approximately 1/10^th^ this diameter (E & F, black arrows).

## 4. Discussion

### 4.1. Vagotopy

When comparing the surgical window in humans to that of the pig, the presence of the nodose (inferior) ganglion within the surgical window in pig but not in humans has two important implications. In terms of intended and unintended neural pathways of activation, the SLN of the vagus branches at or near the nodose ganglion. The LivaNova electrode is placed near the SL bifurcation in our pig functional studies. In canine studies of VNS, electromyographic recordings (EMG) of the laryngeal muscles have demonstrated a short latency response that was eliminated with neuromuscular block (Yoo et al. 2013). The short latency (3 to 5 ms) was not consistent with activation of the recurrent laryngeal branch via motor fibers traveling within the electrode. The response could be explained, however, by ‘stimulation spillover’ due to current spread outside of the electrode activating motor nerves passing nearby, such as the SL. Although the SLN of the vagus branches more cranially in humans, the external superior laryngeal (ESL) branch of the superior laryngeal that innervates the cricothyroid muscles can pass near the carotid bifurcation, which in turn can be located quite close to the traditional clinical VNS lead placement (Monfared et al. 2001). Consequently, activation of the ESL branch may be a contributor to unwanted neck muscle activation in clinical VNS.

The second implication of a more caudal nodose ganglion is the neuronal composition of the ganglion itself. In rodents and guinea pigs, neurons with particular attributes were found to have discrete localizations within the nodose ganglion (Hayes et al. 2013; Travagli et al. 2003). These studies suggest that motor efferent fibers linked to unwanted side effects may have a discrete location, or vagotopy, to separate them from VNS target fibers at the level of the nodose. How far this vagotopy extends caudally from the nodose remains uncertain, but could be an important variable in differences in VNS physiological responses across locations at the cervical level.

A primary goal of this study was to characterize the nodose ganglion as a useful frame of reference to identify a functional vagotopy for potential preferential stimulation of visceral afferent sensory fibers, and avoidance of the motor nerves of the laryngeal neck muscles associated with therapy-limiting side effects. The recurrent laryngeal branch contains motor efferent fibers that innervate the cricoarytenoid, a laryngeal neck muscle. To help identify these pathways, ultrasound video (Supplementary Video 1) with fascicular resolution obtained by placing the transducer in the surgical pocket was taken from the nodose to the point of placement of the stimulating electrodes on the cervical vagus (Supplementary Video 1). This video was then matched to histology from a separate pig to visualize the location of the singular large pseudo-unipolar cell body “fascicle” and secondary fascicles grouped at the level of the nodose. The ultrasound video then follows those groupings to the level of the electrode placement. Using this ultrasound method, in conjunction with additional histological analysis, our data suggest one could minimize off-target activation of the laryngeal neck muscles by placing a small electrode along the cervical vagus trunk away from groupings identified as arising from muscle groups known to elicit side effects.

### 4.2. Variations in Branching

The superior cardiac branches of the vagus, the aortic depressor nerve, the sympathetic trunk, and the carotid sinus nerve are difficult to distinguish in microdissections. As noted by Duncan, differentiating the superior cardiac branches of the vagus - which are often identified as forming a ramus with the cardiac branches of the sympathetic trunk - from the aortic depressor nerve, has been a point of contention in literature (Duncan 1929). The superior cardiac branches are highly variable in their point of origin and can vary in the number and size of fibers they contain. The sympathetic trunk can also be difficult to distinguish from the aortic depressor nerve; it has been reported that in pigs the aortic depressor nerve is found only on the left side (Prakash and Safanie 1967; Schmidt 1968). Schmidt et al. reported that the aortic nerve was either 1) a separate nerve adjacent to the vagus, or 2) projected from the nodose ganglion adjacent to the superior laryngeal nerve, traveling alongside the vagus for a short distance, and then looping back into the vagus (Schmidt 1968).

Additionally, cross-connections between the vagus nerve and the carotid sinus nerve at the level of the carotid bifurcation have been reported in human studies (Toorop et al. 2009); cross-connection to the carotid sinus nerve presumably could easily be conflated with cross-connections to the sympathetic trunk without careful dissection. Unfortunately, detailed microdissection to identify and isolate the sympathetic trunk, the carotid sinus nerve, cross-connections between the vagus and these two structures, and the aortic depressor nerve may not be safe or practical to perform during human VNS surgery. Therefore, post-mortem microdissection in both animals and humans should be performed to understand the neural pathways that might be engaged by VNS.

### 4.3. Histology

As compared to other animal models, the domestic pig best approximated the diameter of the human vagus (Hammer et al. 2018), which is an important point of emphasis for preclinical testing of electrodes and stimulation approaches to minimize off-target effects at human scale. Given that multiple human VNS clinical studies failed to meet their primary efficacy end-point to treat hypertension or heart failure, despite successful studies in animal models (De Ferrari et al. 2017; Ludwig et al. 2017; Musselman, Pelot, and Grill 2019), it is important to understand how the cervical vagus may differ between animal models and human patients with respect to target engagement and dosing. For example, studies by Grill and by Woodbury suggest that the current amplitudes necessary to activate specific fiber types within the vagus between the canine and rat models can differ by up to 100x (Woodbury and Woodbury 1990; Yoo et al. 2013). Given that bipolar stimulating electrodes are typically placed around the vagus nerve, key variables determining threshold for activation are diameter of the cervical vagus nerve under the stimulating electrodes, thickness of the epineurium, electrode geometry, and distance from the electrode to each fascicle. The distance of the electrode to each fiber is particularly important for determining stimulation thresholds, as falloff of an electric field from a bipolar electrode is ∼1/r^2^ (depending on cathode/anode separation) with ‘r’ being the distance from the electrode (Plonsey and Barr 1995).

Estimates of the diameter and number of fascicles in the human vagus vary and depend on the degree of dissection, and on the level and side of the cross section (Hammer et al. 2015; Verlinden et al. 2016). With the exception of the canine data in Figure 6, which was taken from a previous study, the cross-sectional histology spanning mouse, rat, several sizes of pig, non-human primates (NHP), and humans was executed by our group to ensure consistency. In the human sample presented in Figure 9, the diameter of the longest axis is ∼3mm and there are 8 fascicles. Hammer et al reported the average largest diameter of the human vagus is ∼4.6 ± 1.2mm (SD) (Hammer et al. 2015). Verlinden et al. reported the average number of fascicles as 8 ± 2 (SEM) for the right vagus compared to 5 ± 1 (SEM) for the left vagus (Verlinden et al. 2016).

It is interesting to note that although NHPs are often considered the standard for comparison to human in many therapeutic indications, the morphology of the vagus nerve is quite different. NHPs have smaller vagus nerves than humans and notably less complex fascicular organization, and thus smaller distances from the epineural surface to the nerve fibers (Figure 6). This suggests that NHPs may be suboptimal in terms of replicating the clinical environment in VNS studies, and other models may be more appropriate.

The pilot human histology data presented in this paper suggest there may be functional inhomogeneity (vagotopy) in organization in humans that may be exploitable to isolate specific physiological responses. How consistent and distinct this functional organization is between individuals, or whether this distinct functional organization extends caudally along the cervical vagus trunk where VNS electrodes are placed, should be explored in future studies.

### 4.4. Limitations

The present study is intended as a guide for other labs to understand better how the pig cervical vagus may or may not represent the human anatomy with respect to VNS. Microdissection and Gomori’s trichrome staining were used as they can be implemented in any lab and can be extended to make comparisons between other animal models and post-mortem human cadaver work. However, these methods do not allow identification of specific cell types or visceral targets. There may be functional organization at the cervical vagus level that is not only exploitable in terms of preferentially stimulating sensory afferents from the visceral organs, but separable based on organ of origin and specific sensory sub-types (Thompson, Mastitskaya, and Holder 2019; Travagli et al. 2003). Identifying discrete functional subtypes at the level of the cervical vagus with this resolution should be explored in large animal models in future studies.

## 5. CONCLUSION

There is a significant gap in characterization of the vagus nerve both in terms of morphology, on- and off-target effects, and differences across animal models. We quantified vagal morphology in the pig, and the vagus nerve organization and size approximates the clinical environment not only in diameter, but fascicle size and complexity of organization. Our findings revealed potentially exploitable functional vagotopy throughout the length of the vagus nerve, from the level of the nodose to the recurrent laryngeal bifurcation, with respect to the location of sensory afferent fibers arising from pseudo-unipolar cells with their cell bodies in the nodose ganglion. This organization could be leveraged when designing electrodes to stimulate the vagus or determining optimal placement to minimize unwanted side effects such as neck muscle activation. Our data in pig, in conjunction with histology spanning mouse, rat, canine, pig, non-human primate and human models, provides a comprehensive view of the vagus, and is the first necessary step towards selective VNS^1^.

## Supporting information

Additional Pseudo-unipolar Cells, Supplemental Figure 1

Supplemental Figure 2, Fascicle and Nerve Diameter Measurement Record

Supplemental Figure 3, Ultrasound Video of Vagus Nerve

## 6. Acknowledgements

The authors would like to acknowledge funding from The Defense Advanced Research Projects Agency (DARPA) Biological Technologies Office (BTO) Targeted Neuroplasticity Training Program under the auspices of Doug Weber and Tristan McClure-Begley through the Space and Naval Warfare Systems Command (SPAWAR) Systems Center with (SSC) Pacific grant no. N66001-17-2-4010, NIH SPARC OT2 OD025340, and CTSA Grant Number TL1 TR002380 from the National Center for Advancing Translational Science (NCATS). Its contents are solely the responsibility of the authors and do not necessarily represent the official views of the NIH.

We would also like to acknowledge Dr. Jamie Van Gompel (Mayo Clinic) for his instruction on electrode placement procedures, Dr.’s Shigao Chen, Chenyun Zhou, and Chengwu Huang (Mayo Clinic) for their assistance in ultrasound data collection, Aaron Skubal (University of Wisconsin-Madison) for his assistance in creating annotated supplementary ultrasound video, Kim Butters and Karen Lien (Mayo Clinic) for their assistance in the histological processes, and Dr. Joseph Larson (Mayo Clinic) for his statistical direction in interpretation of our data.

## 7. Conflict of Interest Statement

JW and KAL are scientific board members and have stock interests in NeuroOne Medical Inc., a company developing next generation epilepsy monitoring devices. JW also has an equity interest in NeuroNexus technology Inc., a company that supplies electrophysiology equipment and multichannel probes to the neuroscience research community. KAL is also paid member of the scientific advisory board of Cala Health, Blackfynn, Abbott and Battelle. KAL also is a paid consultant for Galvani and Boston Scientific. KAL is a consultant to and co-founder of Neuronoff Inc.

None of these associations are directly relevant to the work presented in this manuscript.

## Footnotes

1 Additional supplemental information can be found online, DOI: 10.26275/fbzm-3eii

