## Additional Pseudo-unipolar Cells, Supplemental Figure 1 for "Functional Vagotopy in the Cervical Vagus Nerve of the Domestic Pig: Implications for the Study of Vagus Nerve Stimulation"

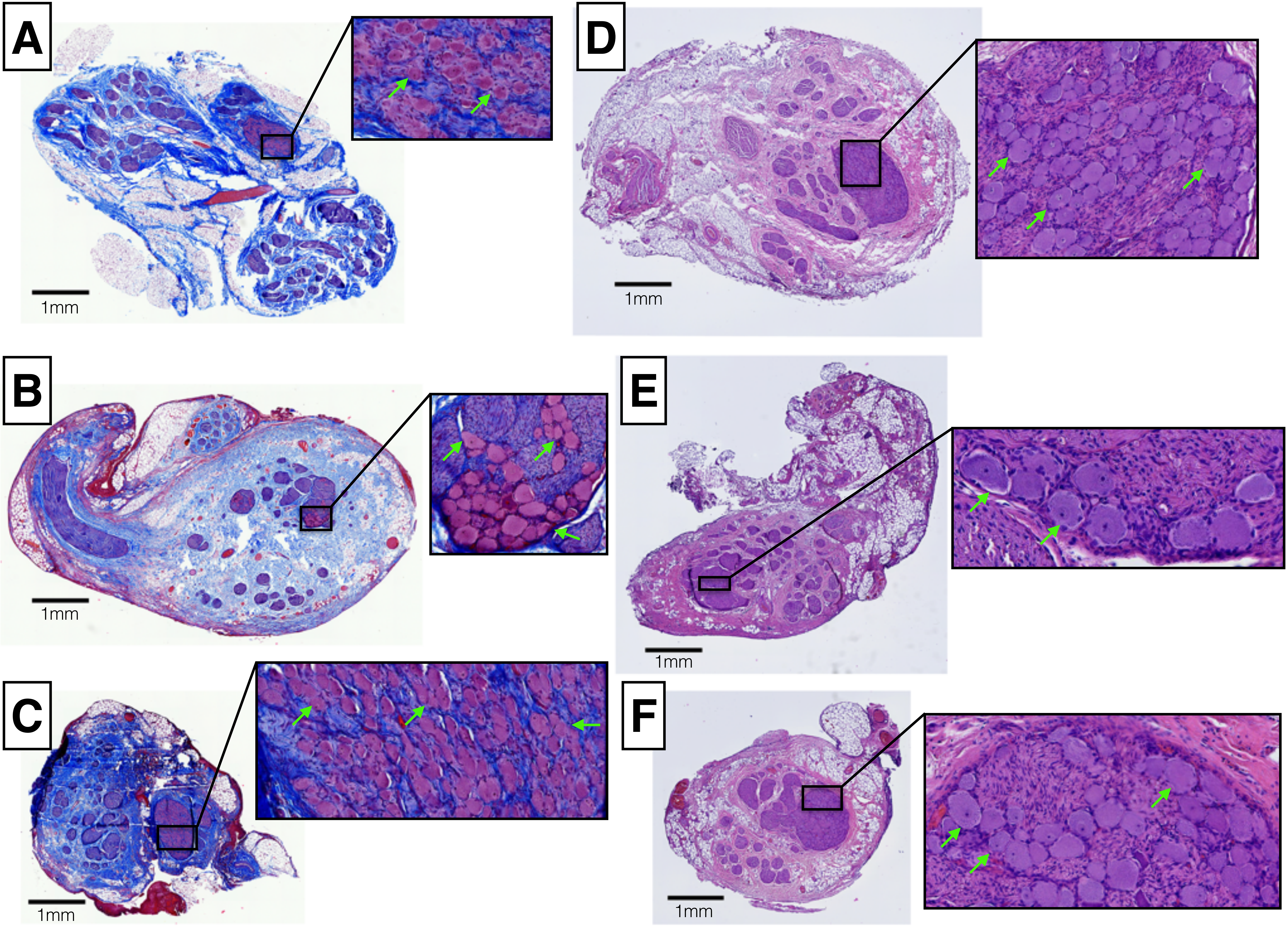

A
1mm
B
1mm
C
1mm
D
1mm
E
1mm
F
1mm

Supplemental Figure 1: Six subjects had vagal nerve sections removed as discussed in methods. These subjects were more sparsely sampled and therefore the pseudo-unipolar cells are visible, but the aggregated plane with the vast majority were not identified. Subjects A, B and C were stained with Gomori's trichrome. Subjects D, E, and F were completed during pilot studies, and stained with H & E. Green arrows indicate pseudo-unipolar cells with surrounding satellite cells.

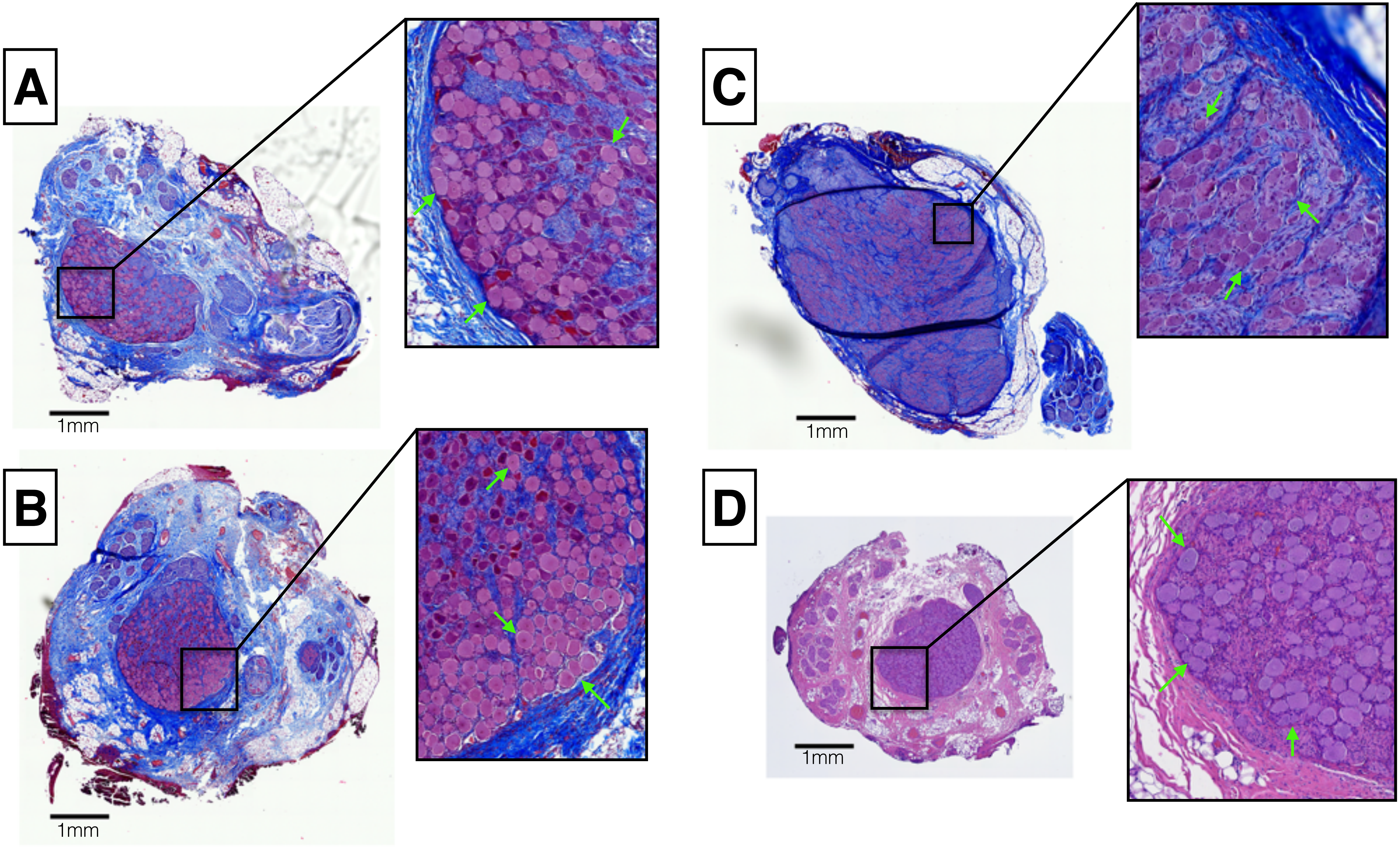

A
1mm
B
1mm
C
1mm
D
1mm

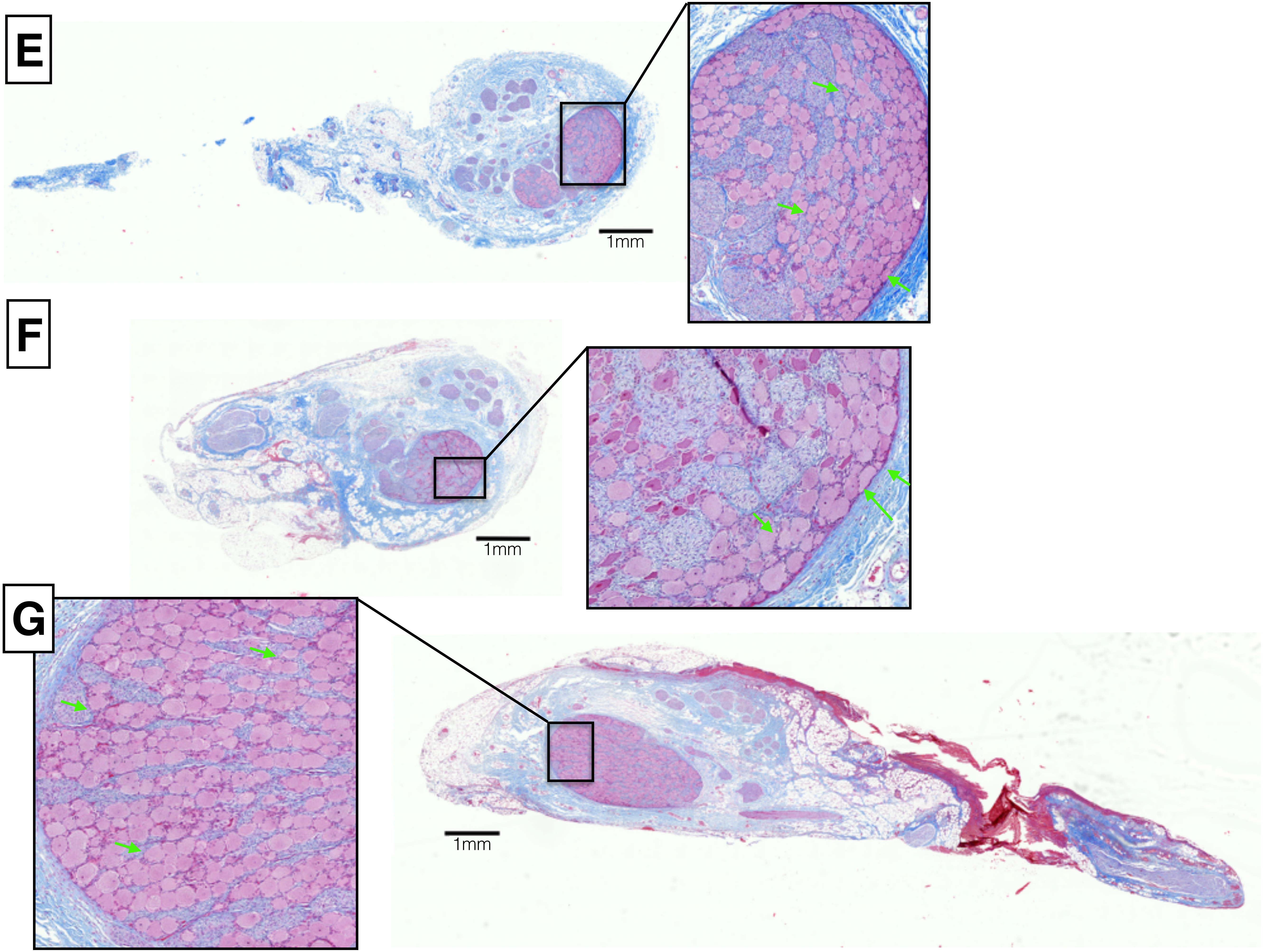
E
1mm
F
1mm
G
1mm

Supplemental Figure 2: Six subjects (in addition to those presented in the manuscript) had vagal nerve sections removed as discussed in methods. In these subjects we were able to locate the pseudo-unipolar cells in an aggregated plane. Subjects A, B, C, E and F were stained with Gomori's trichrome. Subject D was completed during pilot studies, and stained with Hematoxylin & Eosin. Green arrows indicate pseudo-unipolar cells with surrounding satellite cells.

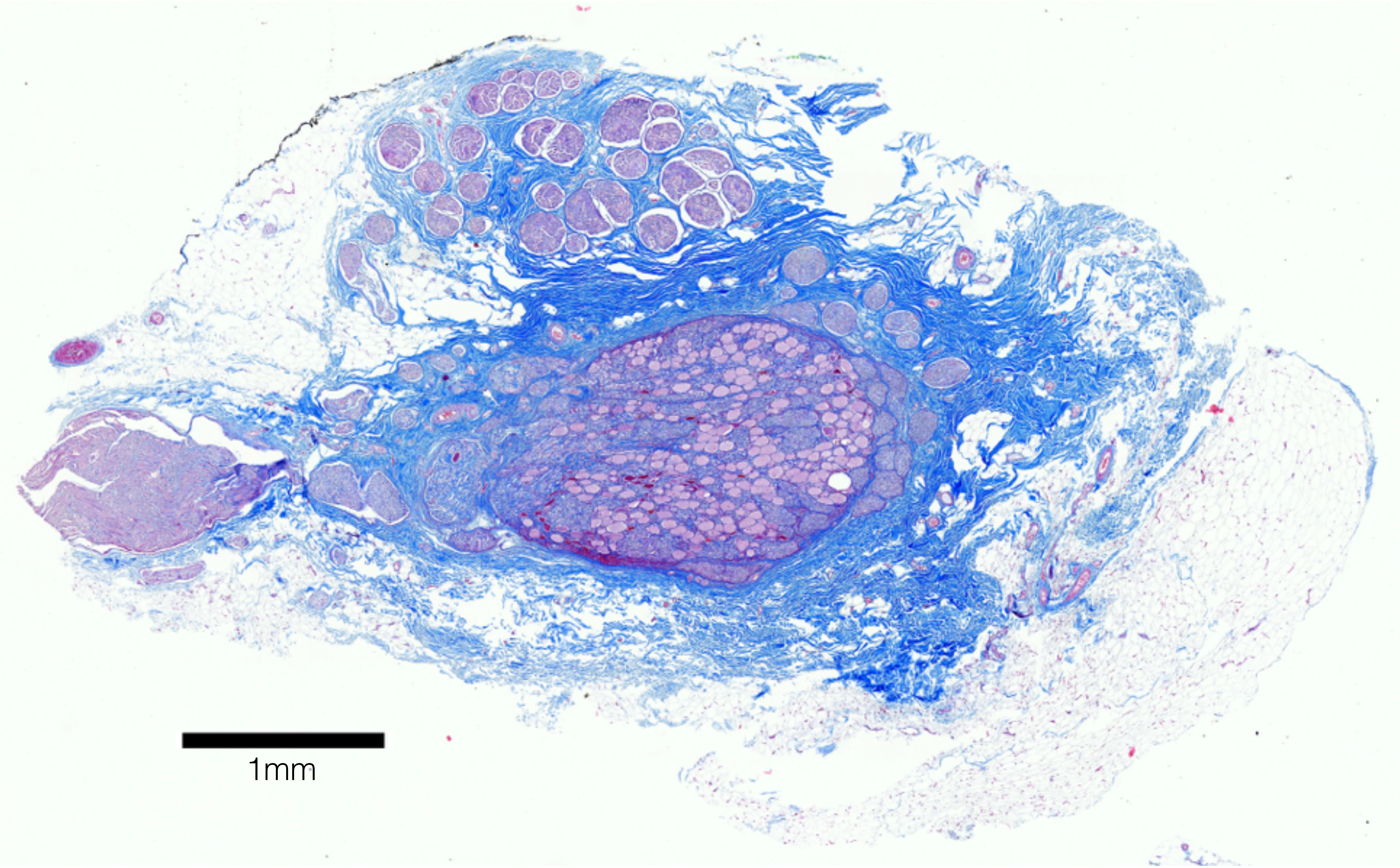
1mm

Supplemental Figure 3: As shown in this representative animal, current vagal nerve studies extending to miniature pigs have shown similar organization in the aggregation of pseudo-unipolar cells at the level of the nodose ganglia. Data has been collected in an additional five miniature pigs, and is currently being analyzed.
