## Supplemental Figure 2, Fascicle and Nerve Diameter Measurement Record for "Functional Vagotopy in the Cervical Vagus Nerve of the Domestic Pig: Implications for the Study of Vagus Nerve Stimulation"

Scale: 1894 pixels/mm

Subject 2 Record_P818

### Nodose

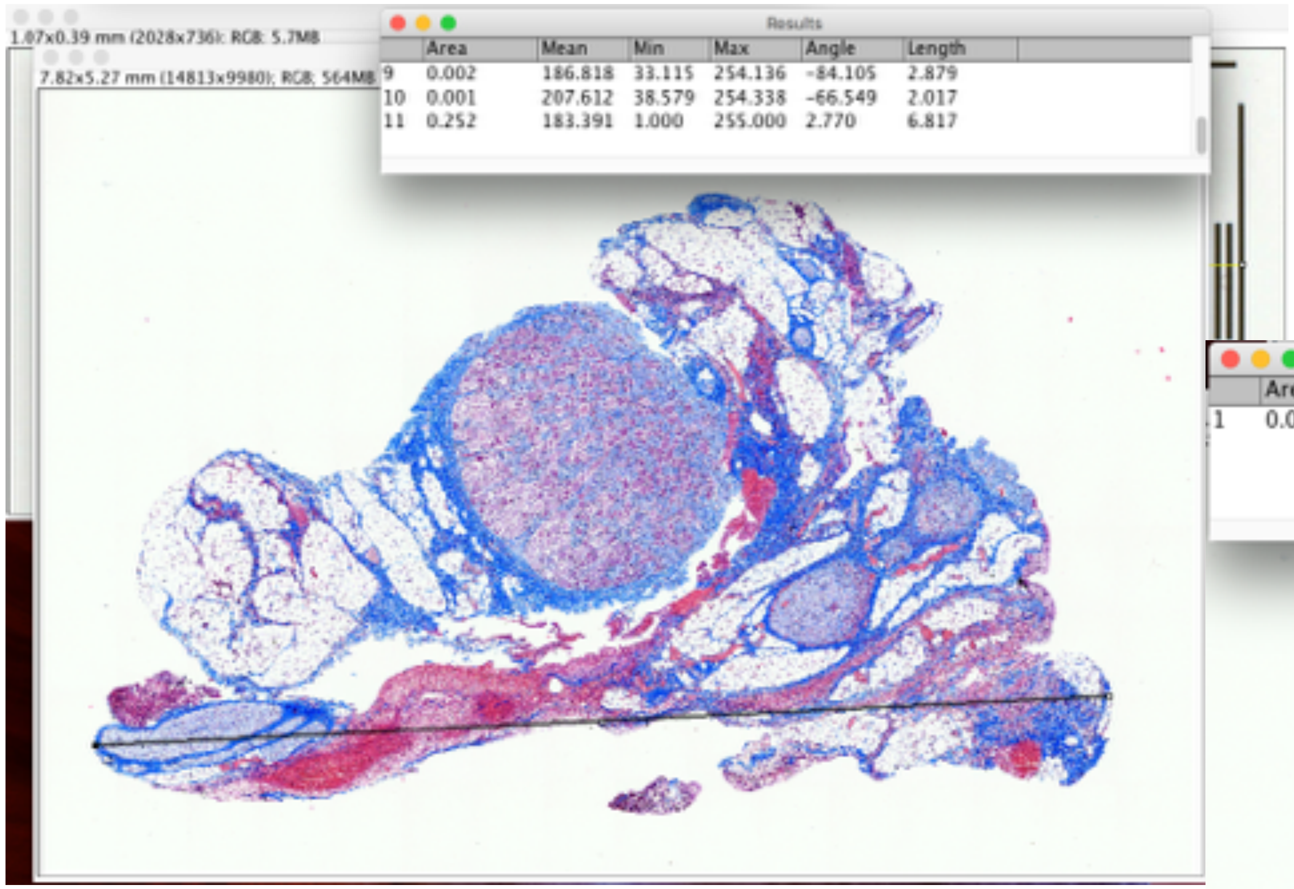


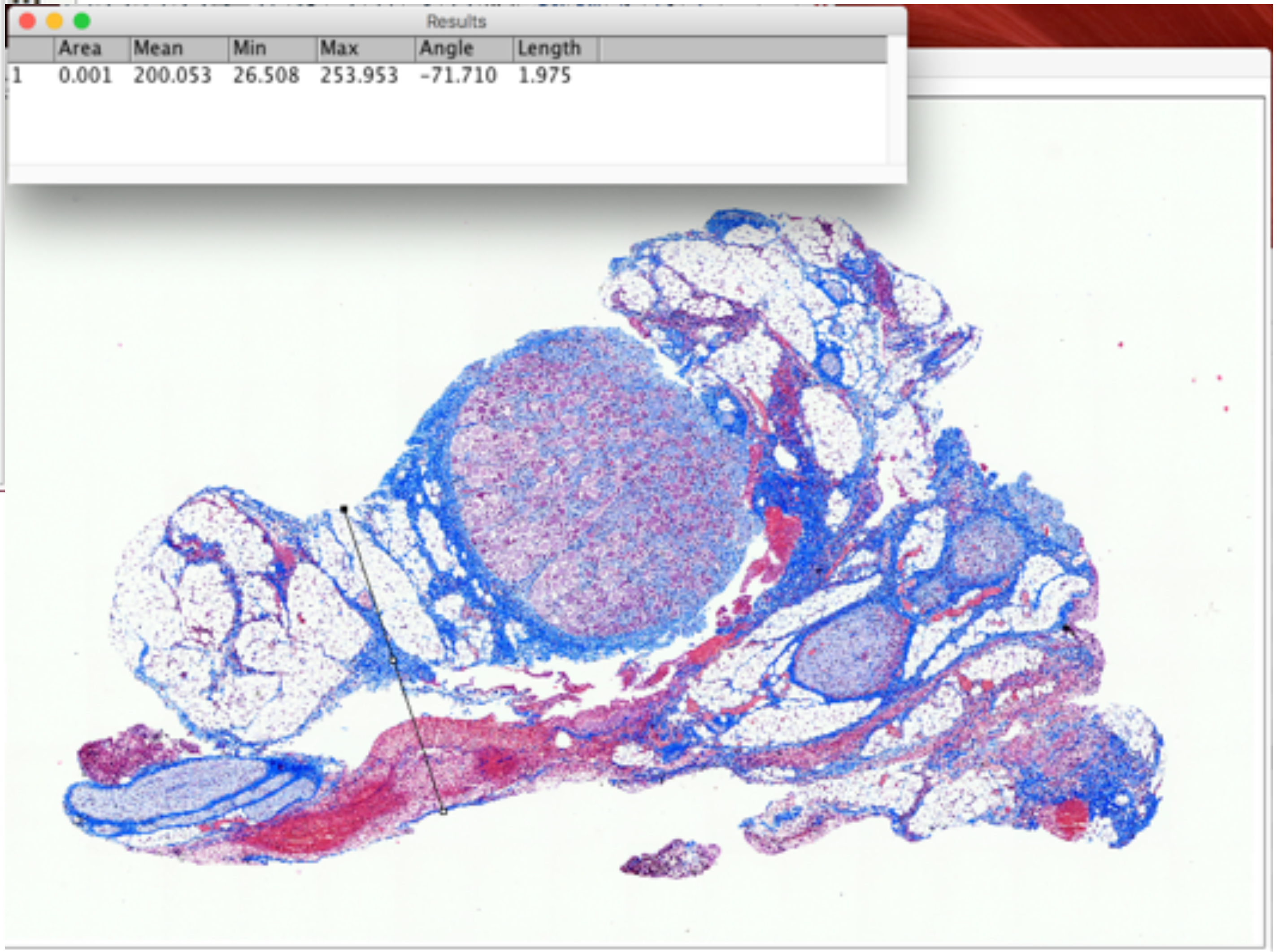


P818_LV4

widest and narrowest diameter

Nodose

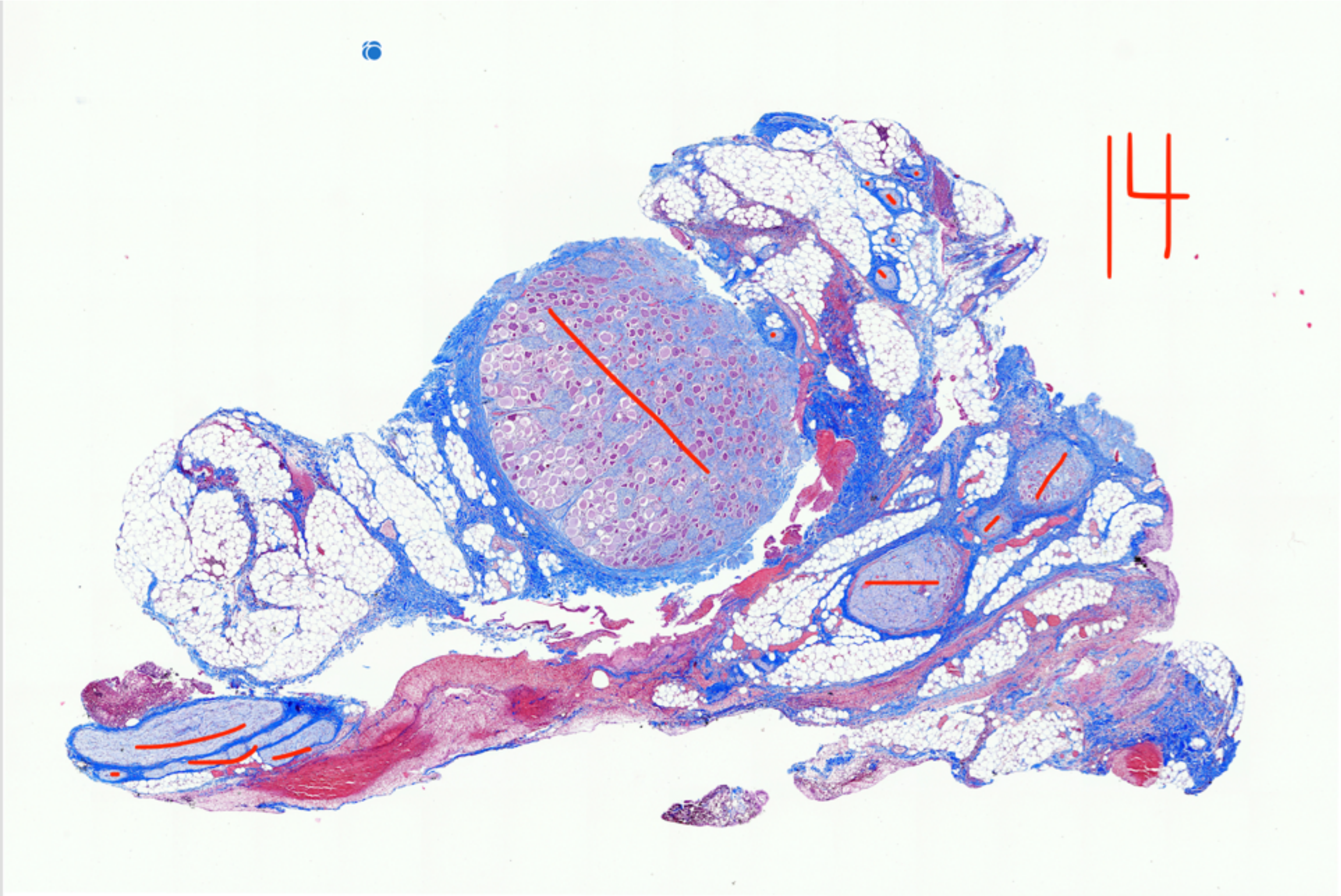


P818_LV4

Fascicle count, 14

### Nodose

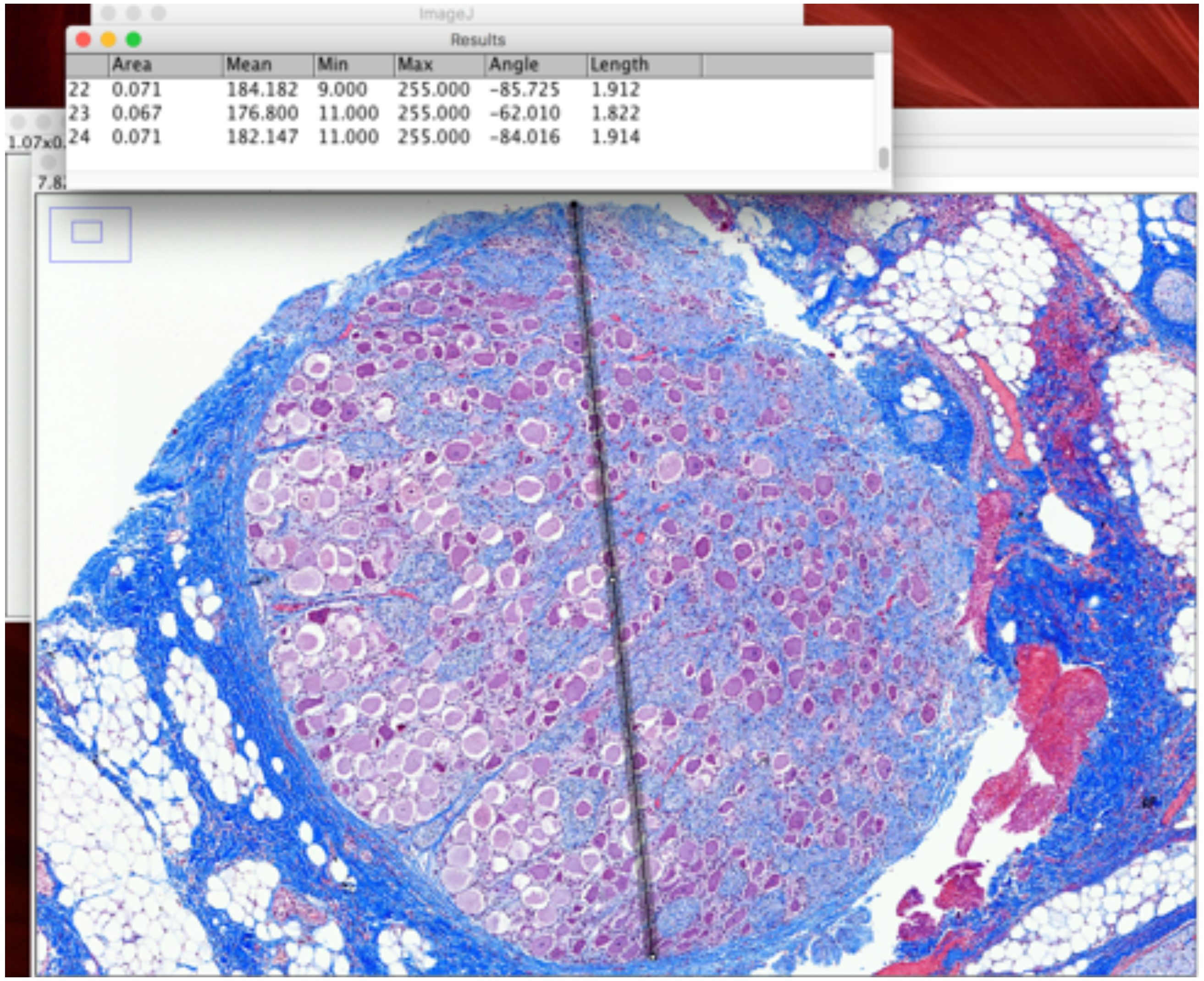


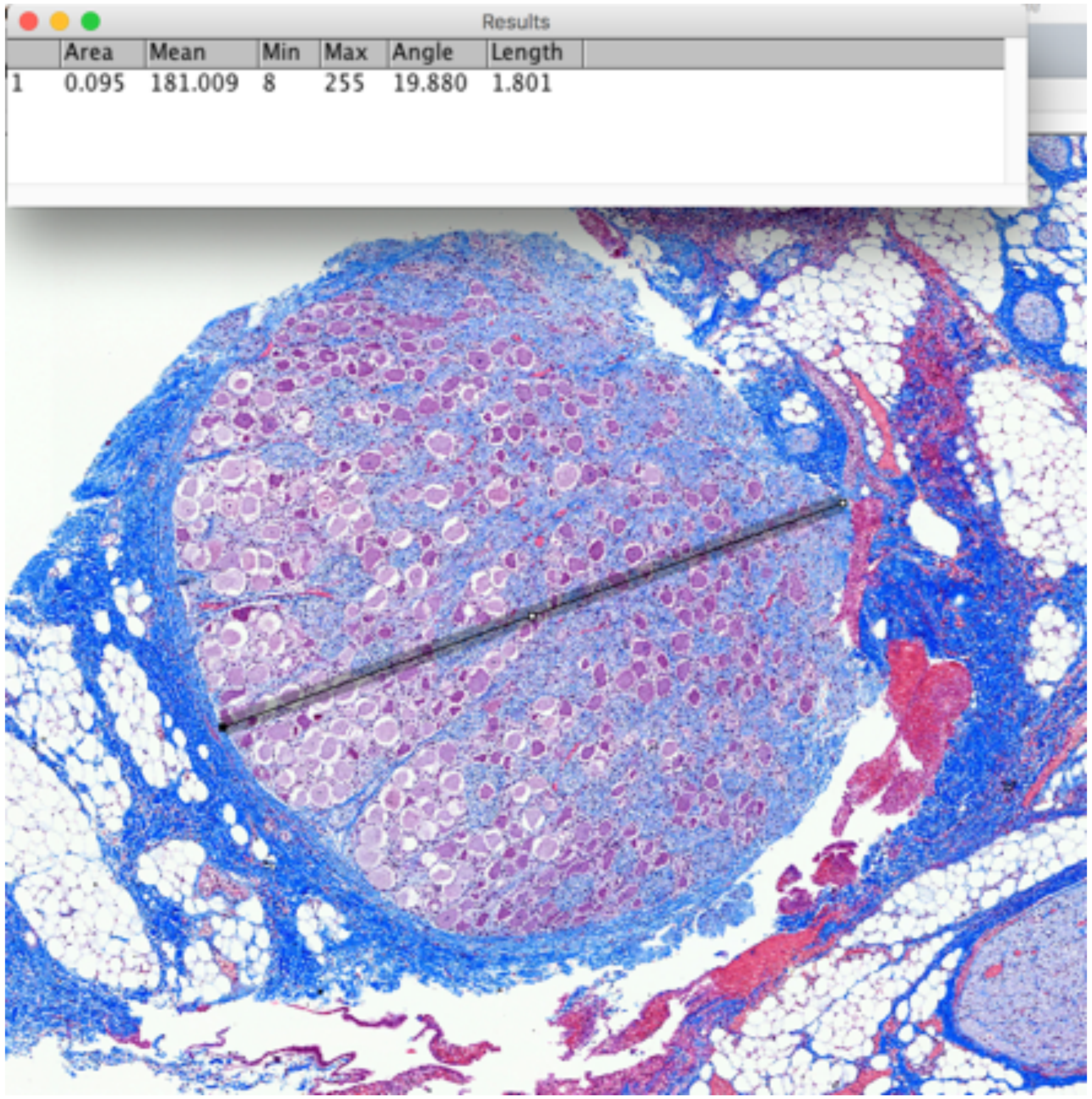


P818_LV4 Largest Fascicle_widest and narrowest diameter

### Mid-VN

Results

| | Area | Mean | Min | Max | Angle | Length |
|---|---|---|---|---|---|---|
| 24 | 0.071 | 182.147 | 11.000 | 255.000 | -84.016 | 1.914 |
| 25 | 0.071 | 182.147 | 11.000 | 255.000 | -84.016 | 1.914 |
| 26 | 0.201 | 190.344 | 0.000 | 255.000 | -110.086 | 5.424 |

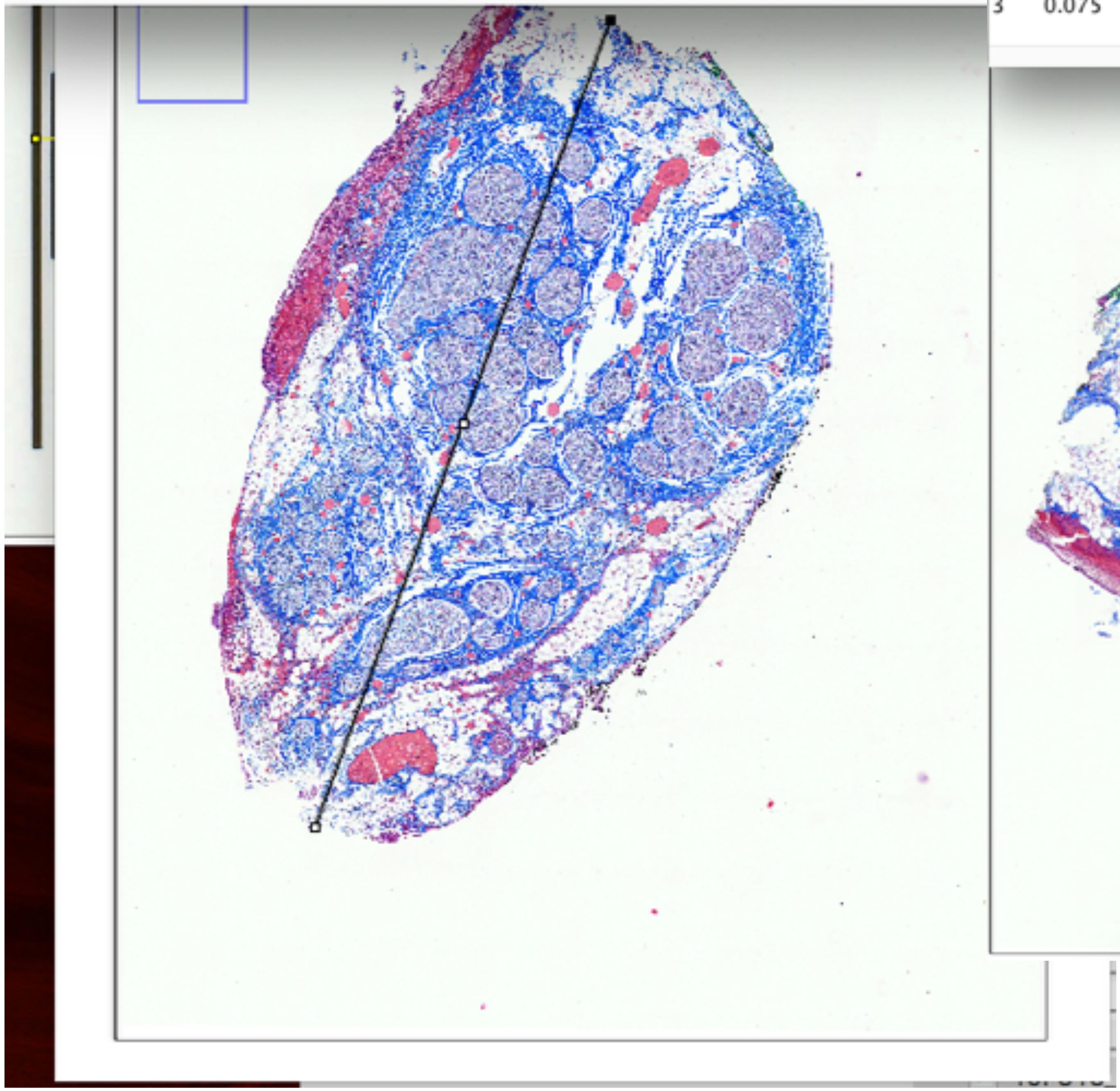

Results

| | Area | Mean | Min | Max | Angle | Length |
|---|---|---|---|---|---|---|
| 1 | 8.157E-4 | 183.671 | 40.595 | 253.834 | 72.451 | 1.544 |
| 2 | 0.074 | 216.195 | 3.000 | 255.000 | -111.413 | 1.388 |
| 3 | 0.075 | 204.255 | 2.000 | 255.000 | -112.306 | 1.424 |

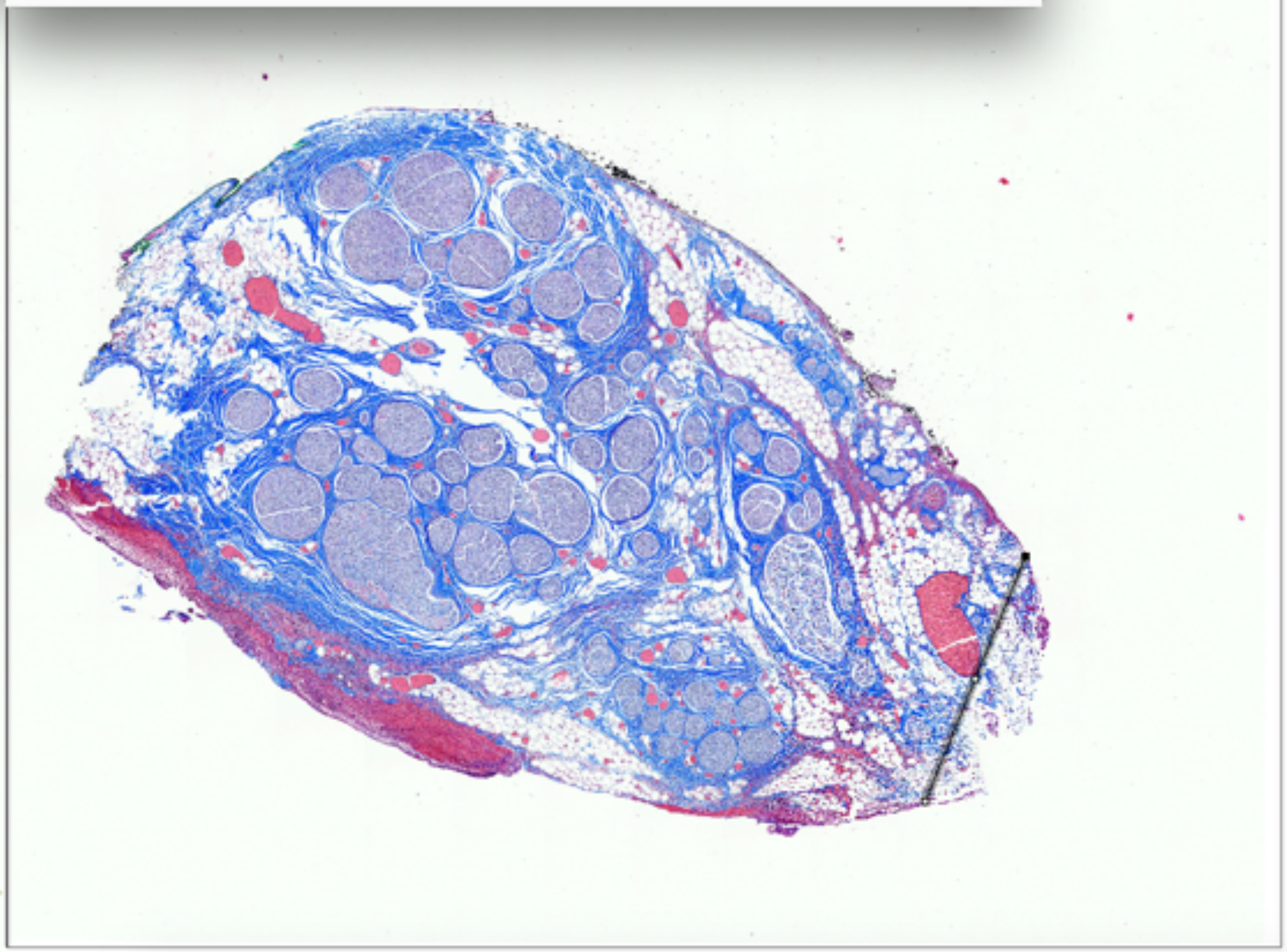

P818_LV5b

widest and narrowest diameter

Mid-VN

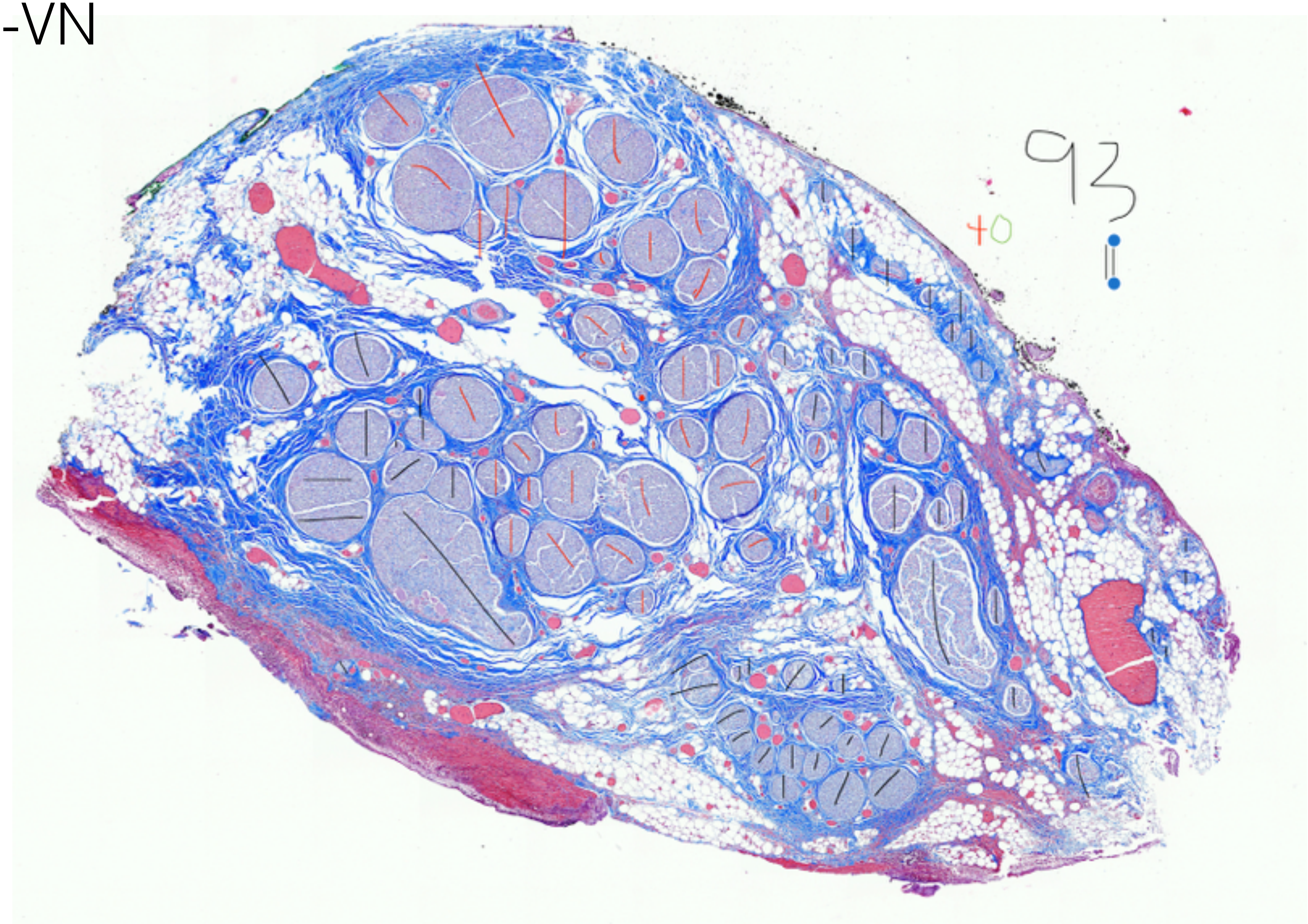


P818_LV5b

Fascicle count, 95

Mid-VN

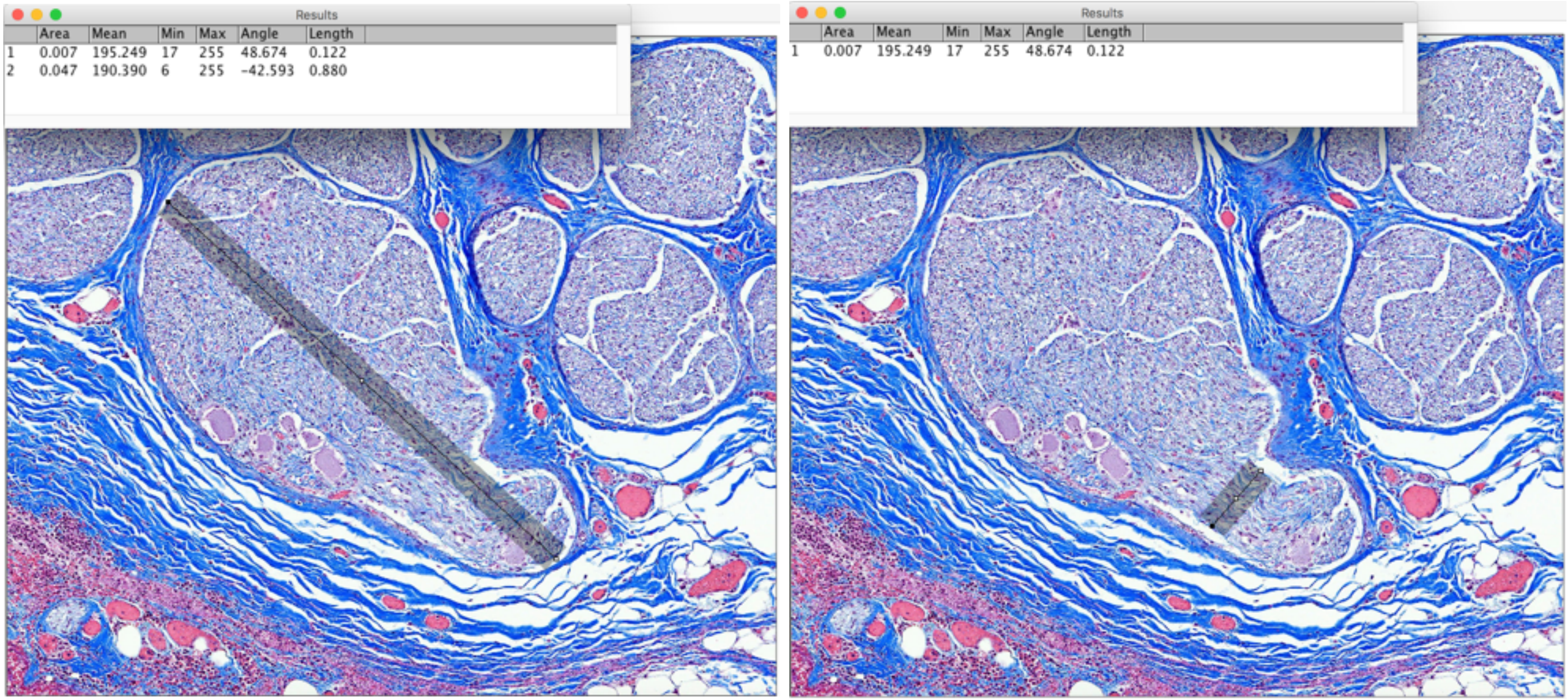


P818_LV5b     Largest Fascicle_widest and narrowest diameter

Mid-VN

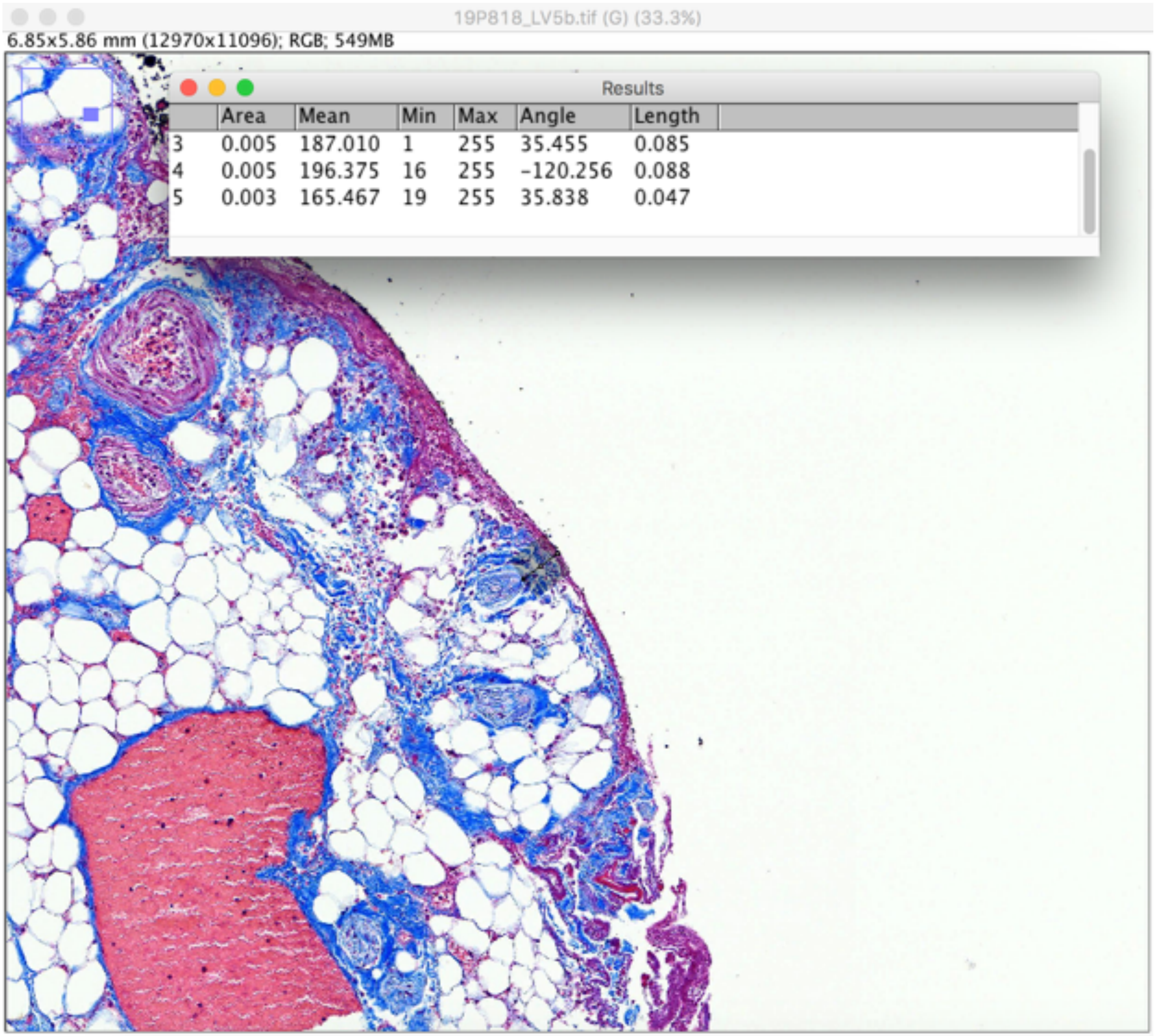


P818_LV5b

fascicle depth

Subject 6_P839

### Pre-Nodose

Results

| | Area | Mean | Min | Max | Angle | Length |
|---|---|---|---|---|---|---|
| 31 | 0.206 | 203.180 | 30.000 | 255.000 | -105.933 | 5.508 |
| 32 | 0.110 | 180.568 | 28.000 | 255.000 | -9.408 | 2.946 |
| 33 | 0.166 | 163.476 | 0.000 | 255.000 | -98.655 | 4.435 |

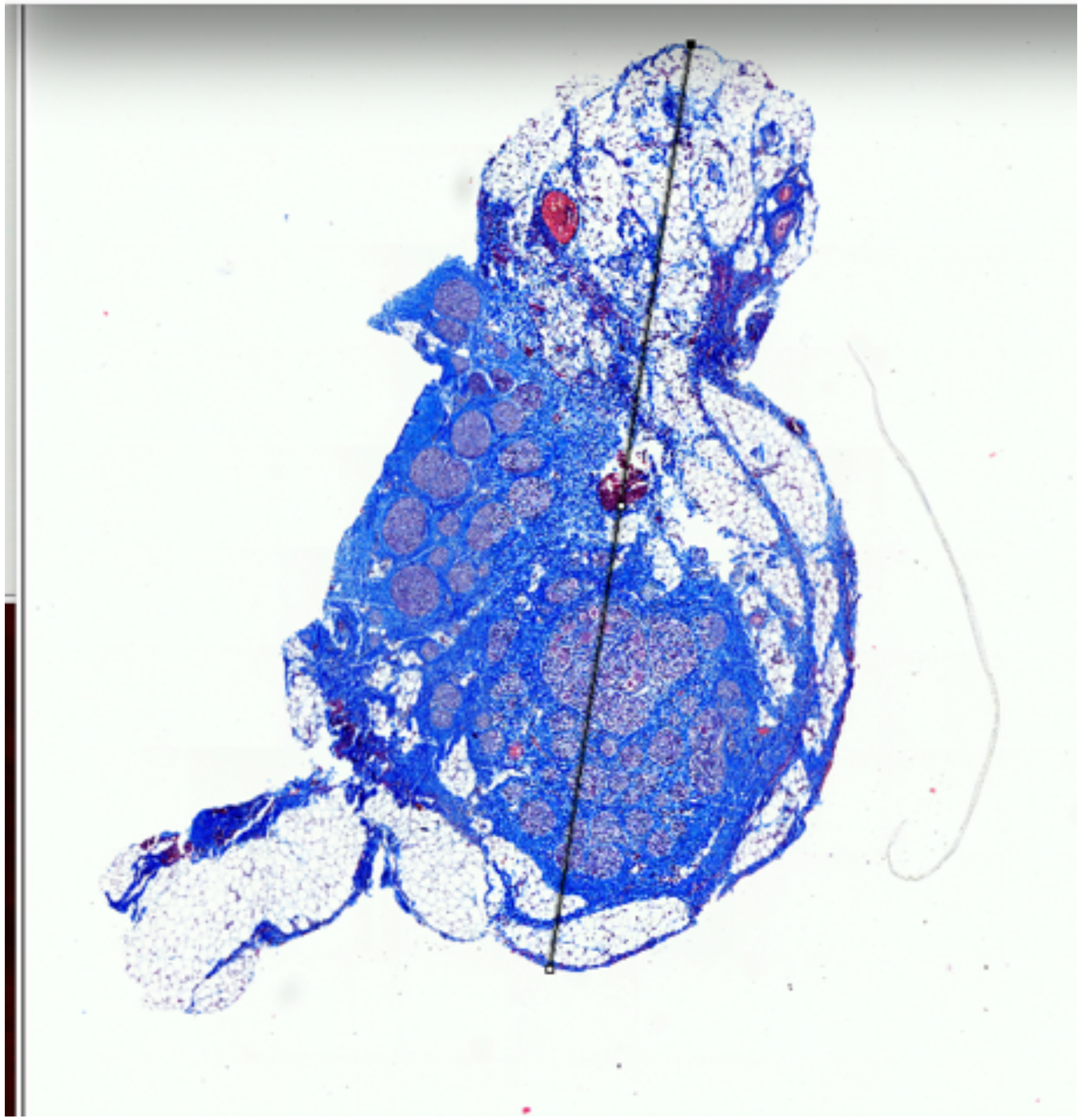

Results

| | Area | Mean | Min | Max | Angle | Length |
|---|---|---|---|---|---|---|
| 4 | 0.005 | 196.375 | 16 | 255 | -120.256 | 0.088 |
| 5 | 0.003 | 165.467 | 19 | 255 | 35.838 | 0.047 |
| 6 | 0.072 | 148.915 | 3 | 255 | -0.796 | 1.369 |

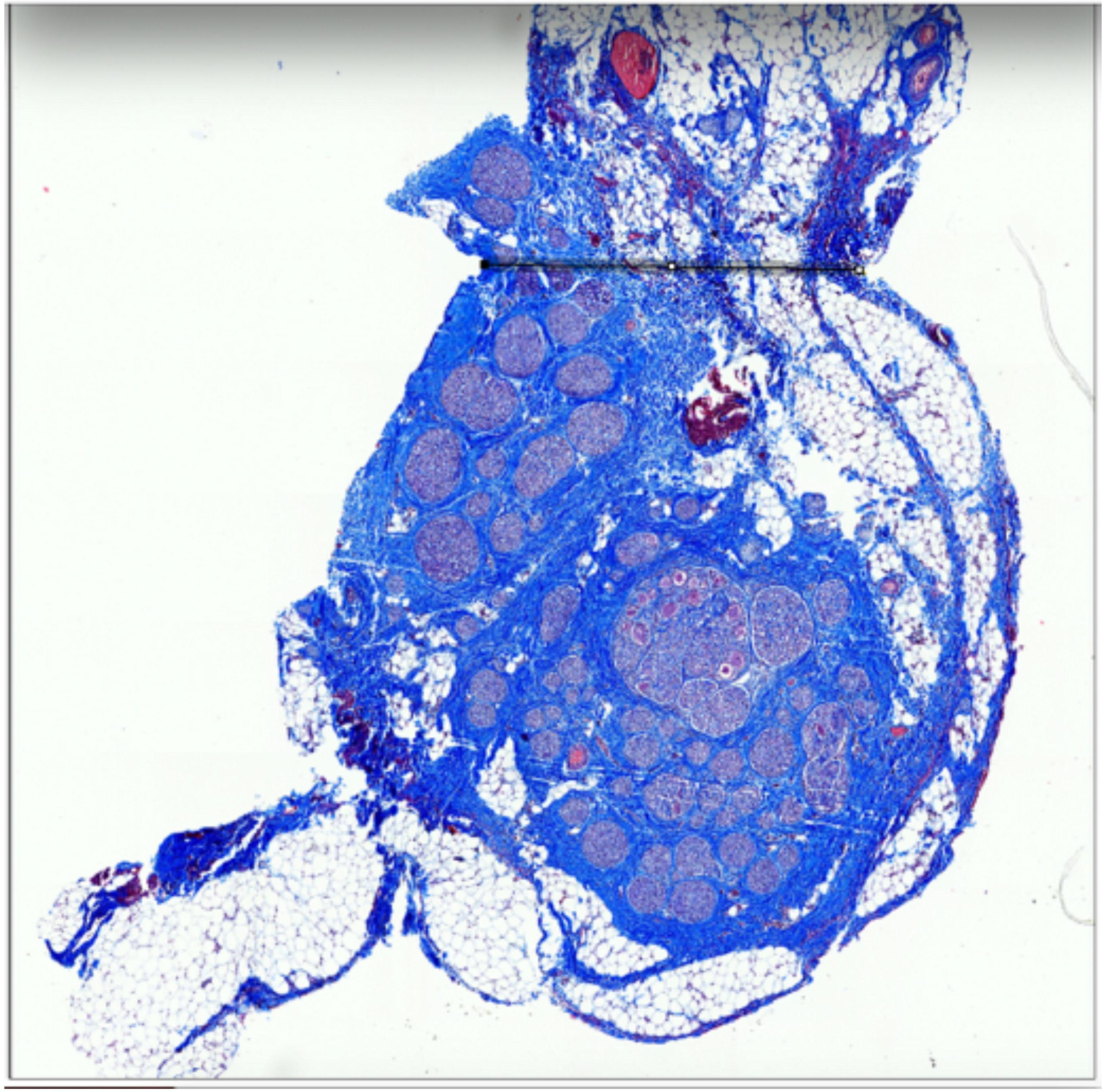

P839_Nodose_4

Widest and narrowest diameters

### Pre-Nodose

Results

| | Area | Mean | Min | Max | Angle | Length |
|---|---|---|---|---|---|---|
| 33 | 0.166 | 163.476 | 0.000 | 255.000 | -98.655 | 4.435 |
| 34 | 0.095 | 140.053 | 7.000 | 255.000 | -27.154 | 2.573 |
| 35 | 0.019 | 151.138 | 7.000 | 255.000 | 47.222 | 0.520 |

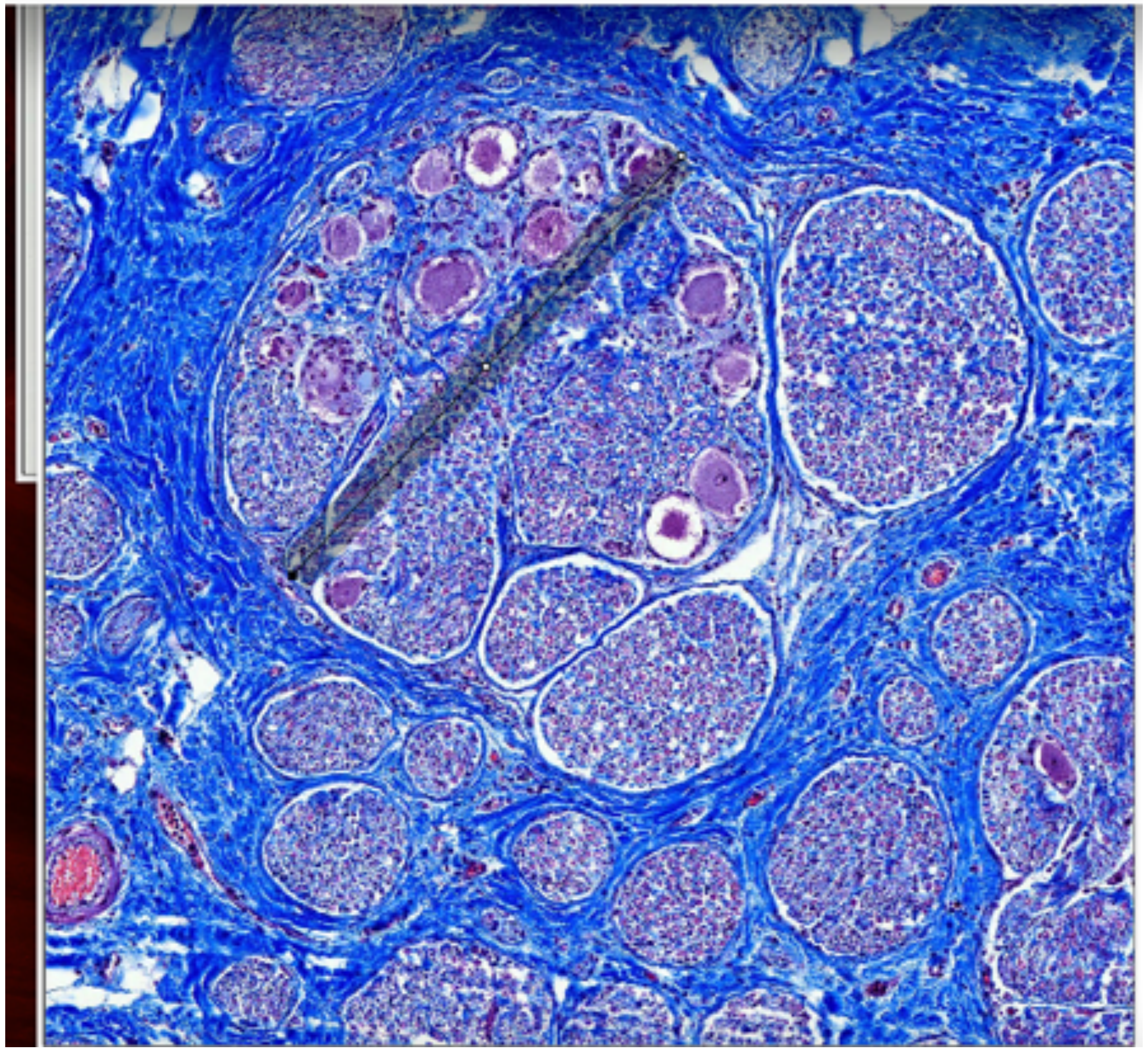

Results

| | Area | Mean | Min | Max | Angle | Length |
|---|---|---|---|---|---|---|
| 2 | 0.011 | 154.088 | 3 | 255 | -48.180 | 0.215 |
| 3 | 0.011 | 154.088 | 3 | 255 | -48.180 | 0.215 |
| 4 | 0.018 | 146.596 | 15 | 255 | -71.346 | 0.350 |

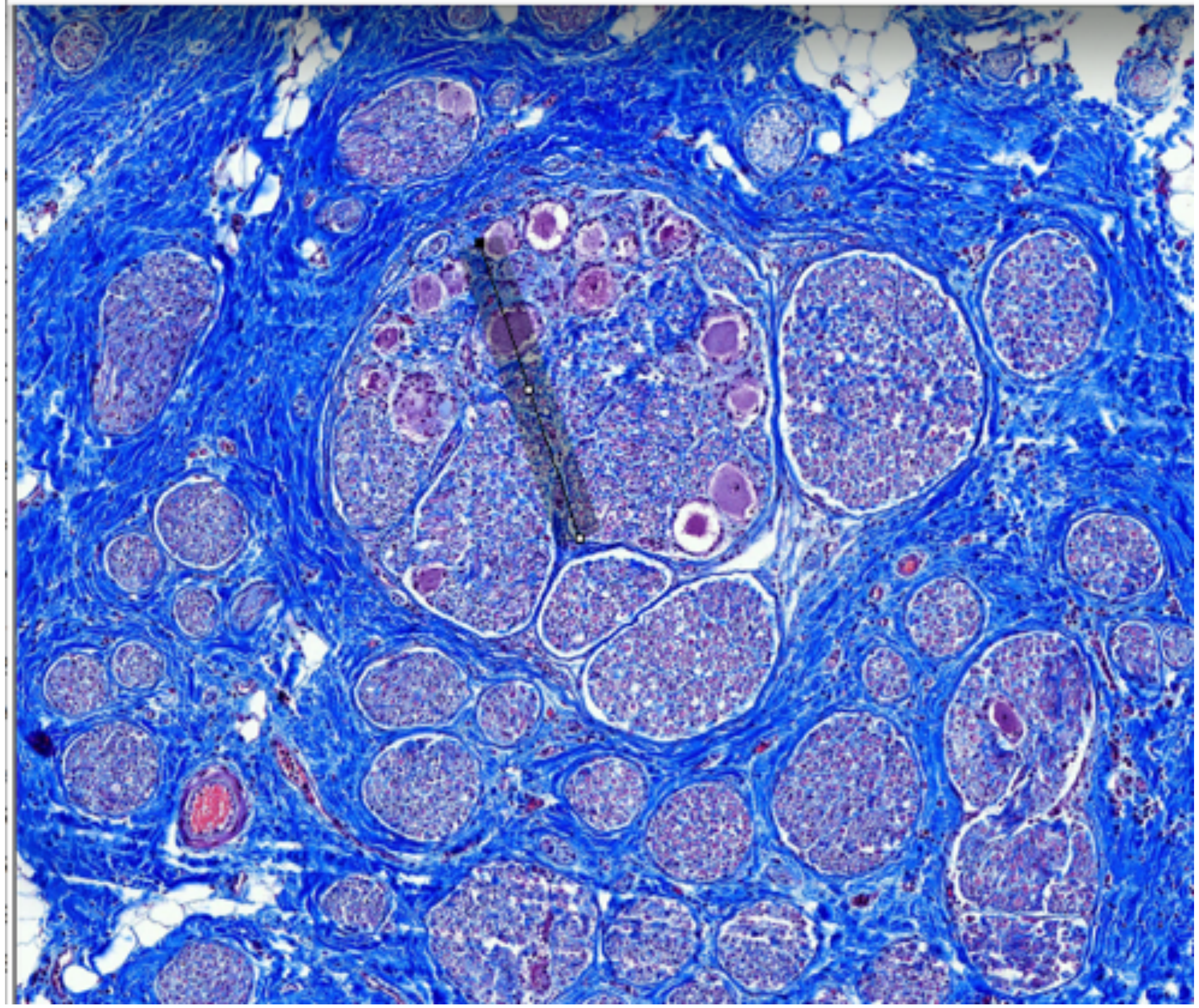

P839_Nodose_4 Largest fascicle _widest and narrowest diameters

Pre-Nodose

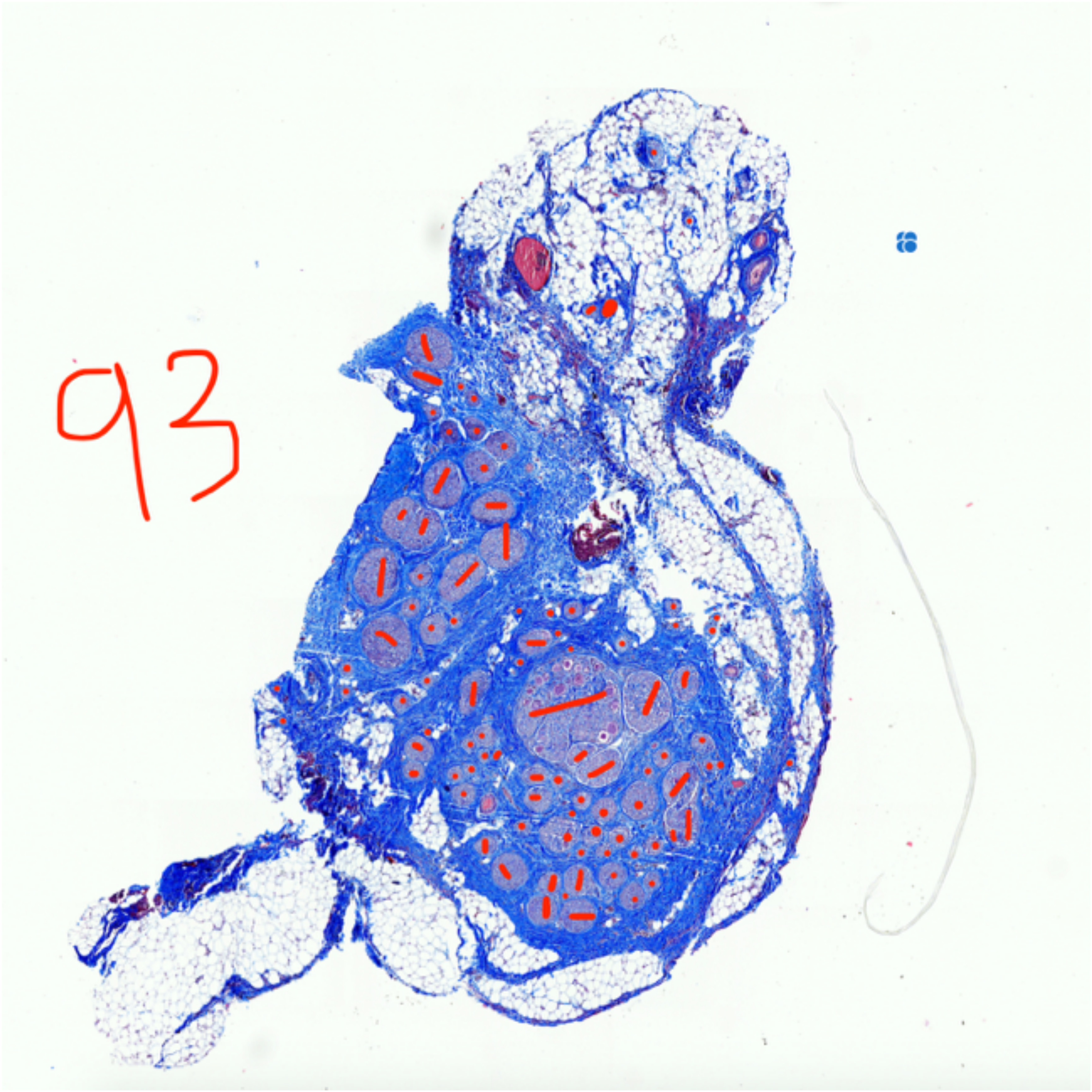


P839_Nodose_4

Fascicle count, 93

### Nodose

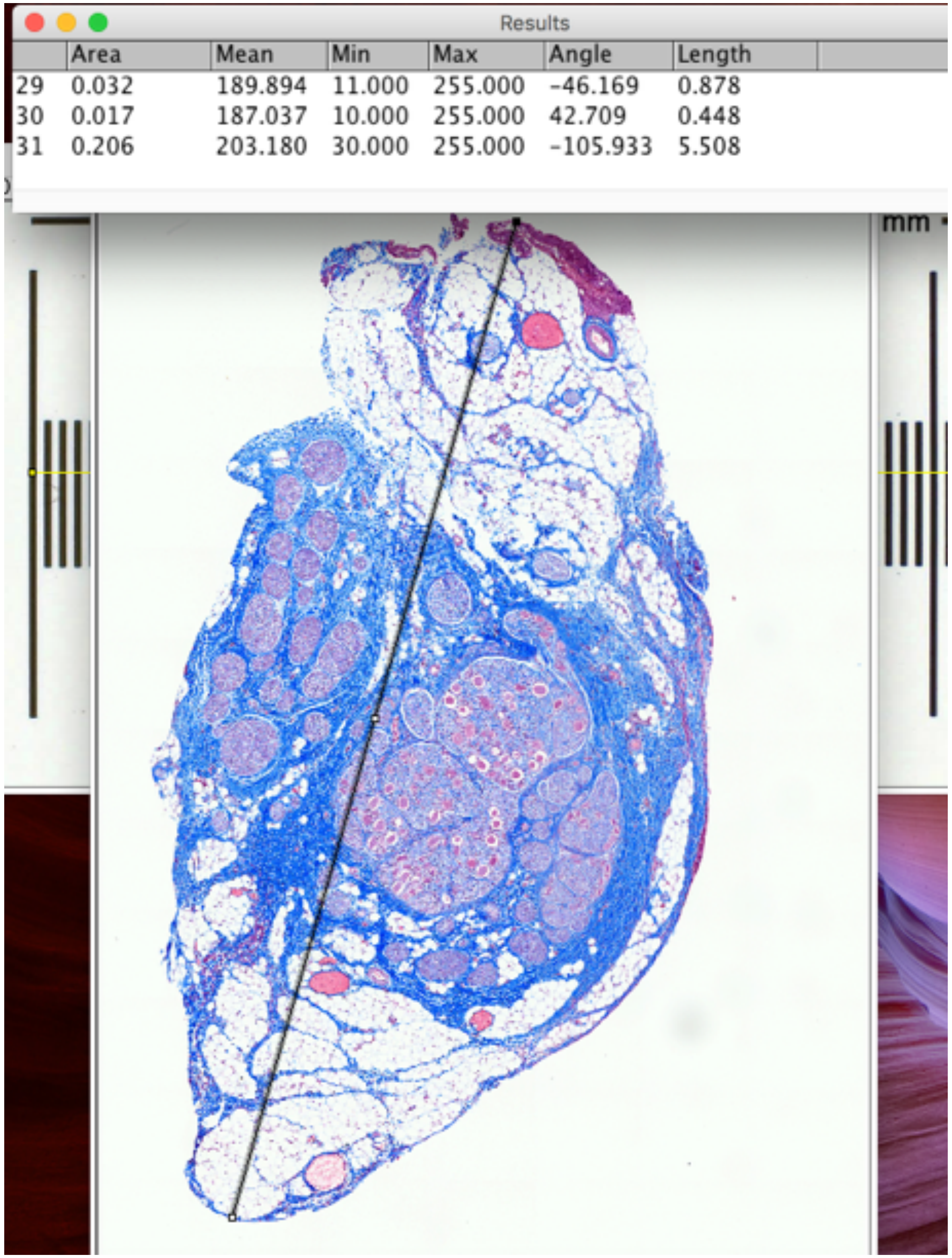


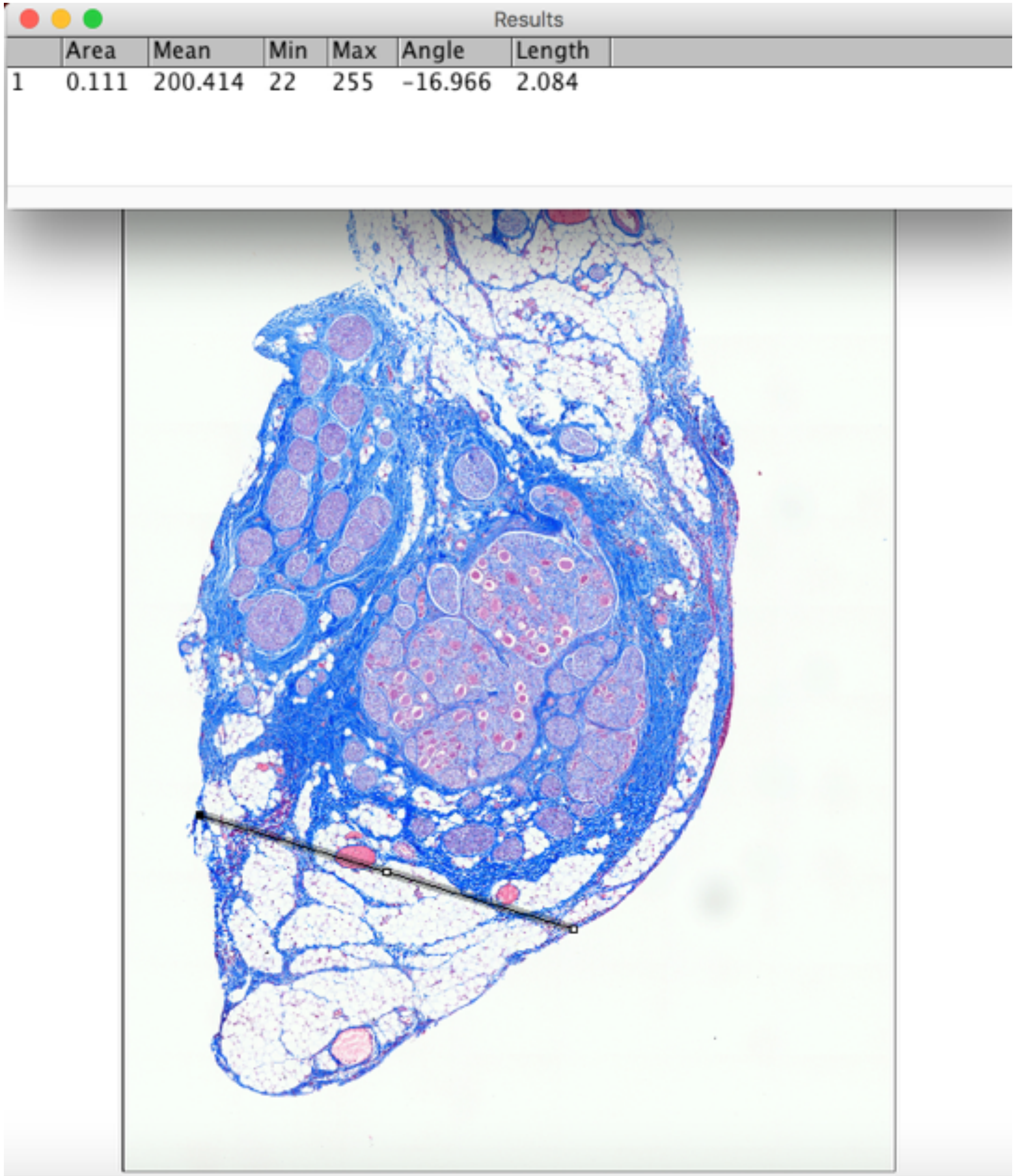


P839_Nodose_43

Widest and narrowest diameters

### Nodose

Results

| | Area | Mean | Min | Max | Angle | Length |
|---|---|---|---|---|---|---|
| 36 | 0.026 | 190.767 | 33.000 | 255.000 | -79.695 | 0.708 |
| 37 | 0.029 | 188.655 | 50.000 | 255.000 | 5.208 | 0.791 |
| 38 | 0.030 | 186.739 | 32.000 | 255.000 | 8.688 | 0.797 |

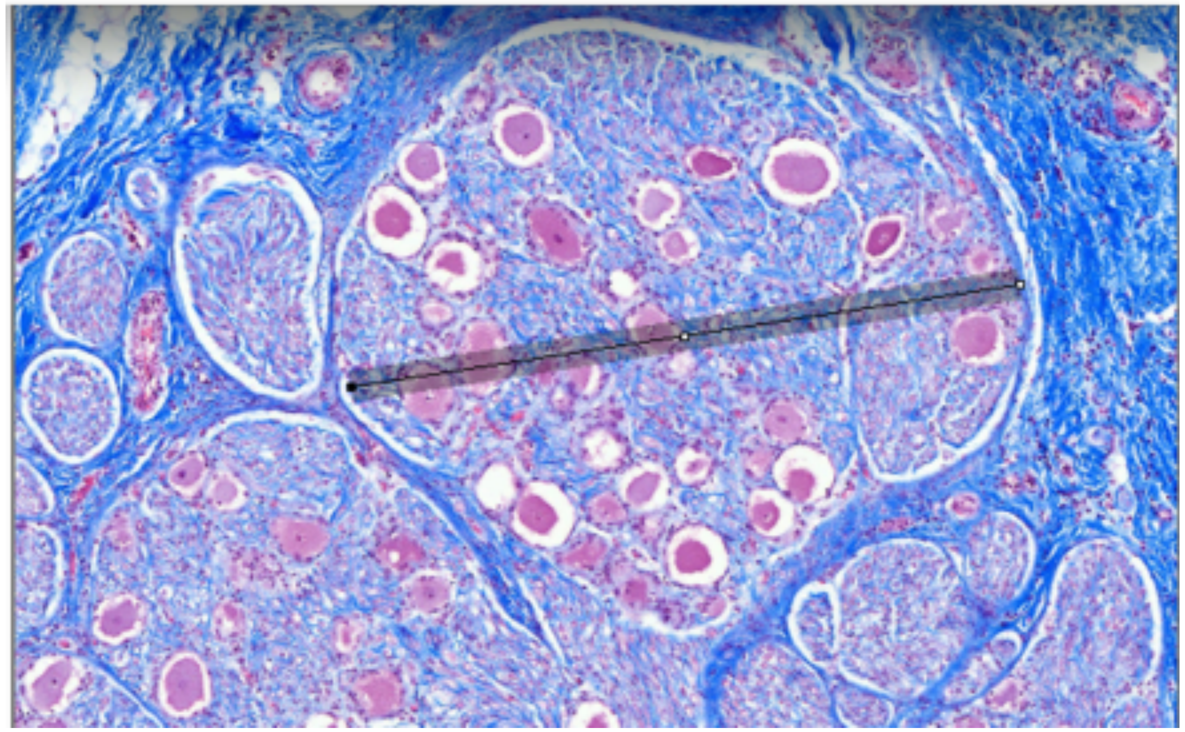

Results

| | Area | Mean | Min | Max | Angle | Length |
|---|---|---|---|---|---|---|
| 1 | 0.034 | 190.507 | 38 | 255 | -47.971 | 0.648 |

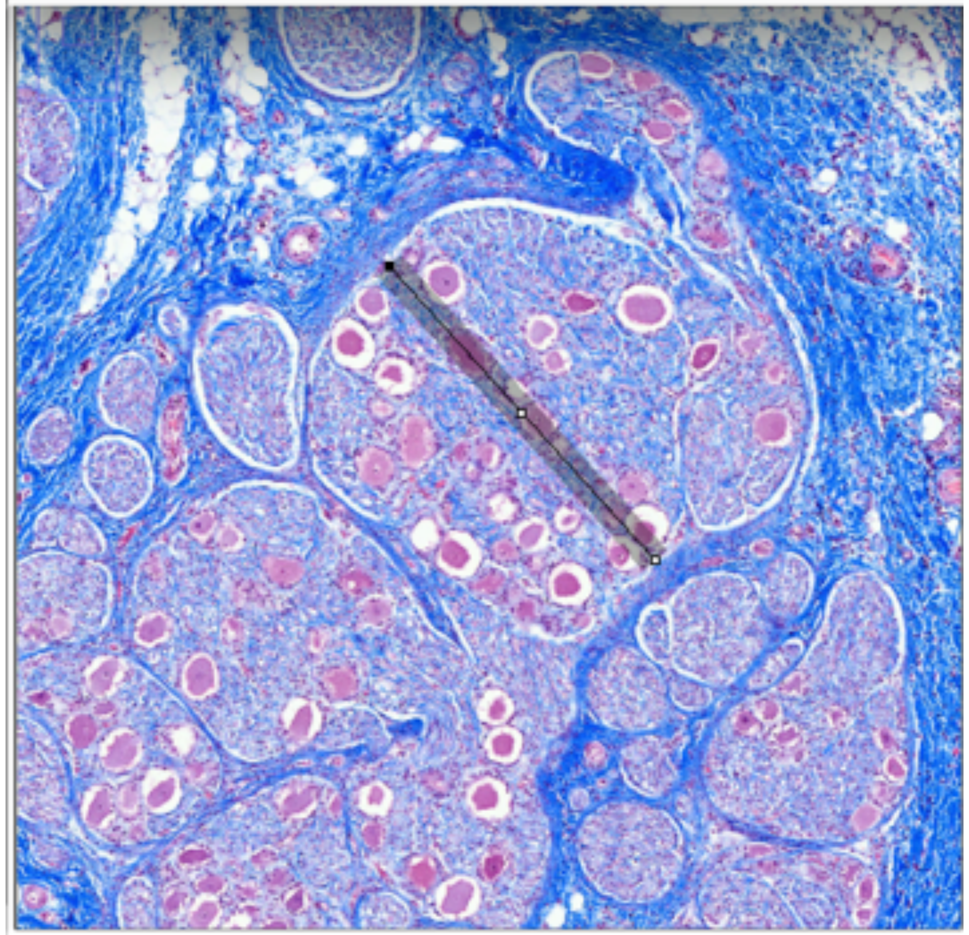

P839_Nodose_43 Largest Fascicle_Widest and narrowest diameters

### Nodose

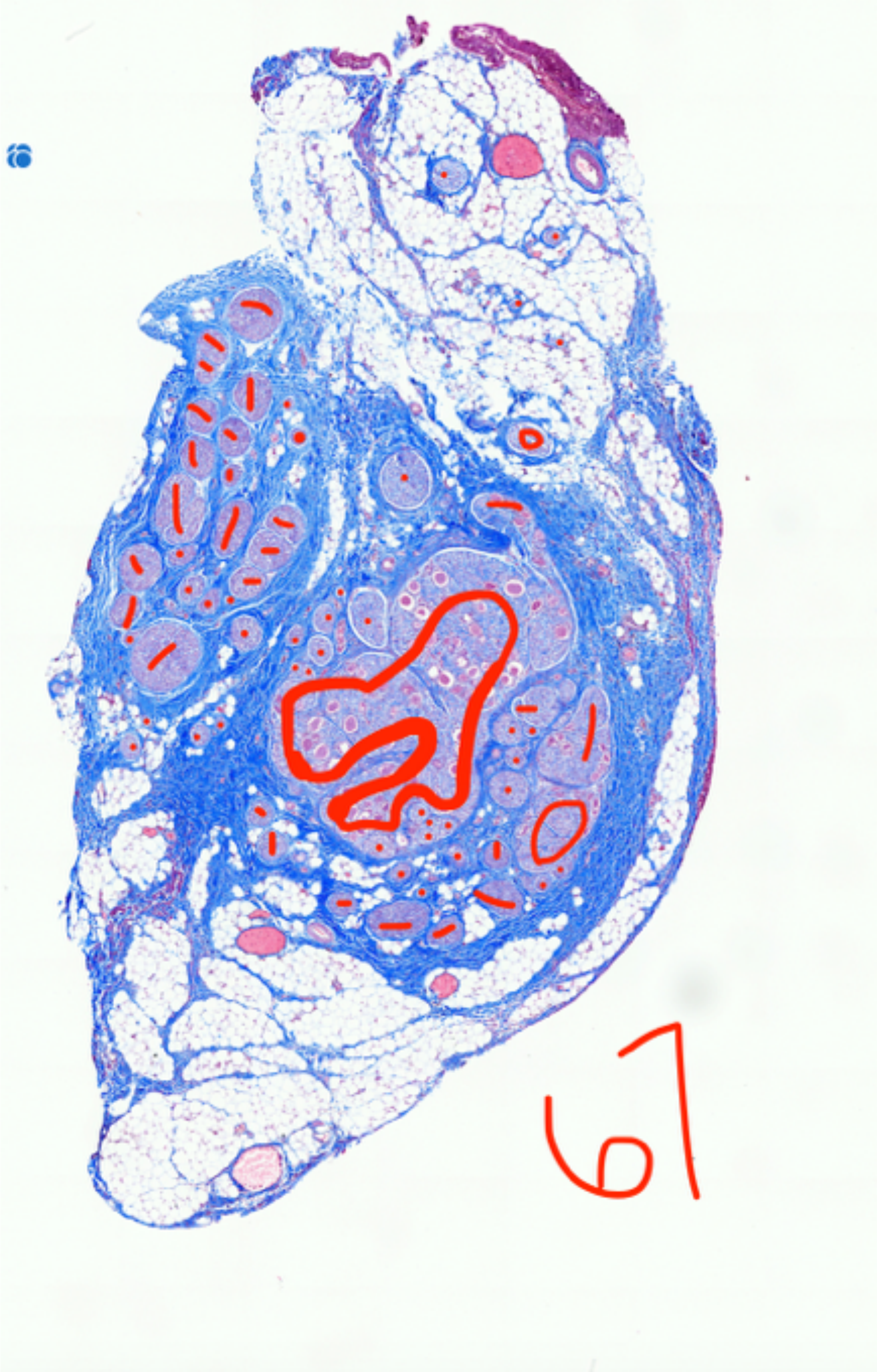


P839_Nodose_4

Fascicle count, 67

Subject 7_P845

### Pre-nodose

Results

| | Area | Mean | Min | Max | Angle | Length |
|---|---|---|---|---|---|---|
| 39 | 0.011 | 179.848 | 8.000 | 255.000 | -83.949 | 0.301 |
| 40 | 0.015 | 174.962 | 8.000 | 255.000 | 19.592 | 0.397 |
| 41 | 0.125 | 167.558 | 8.000 | 255.000 | -95.077 | 3.365 |

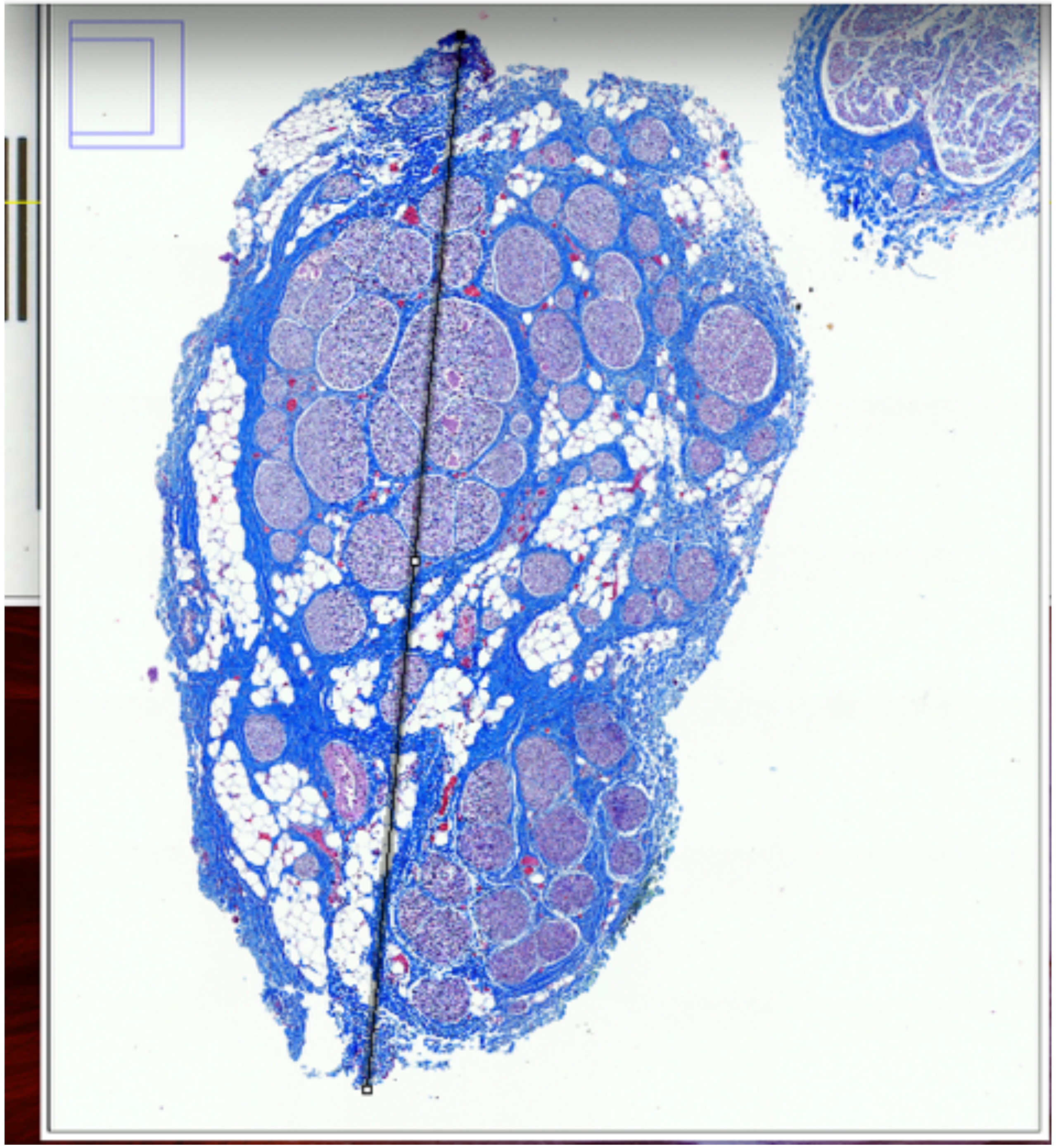

Results

| | Area | Mean | Min | Max | Angle | Length |
|---|---|---|---|---|---|---|
| 1 | 0.081 | 168.558 | 6 | 255 | -2.369 | 1.533 |

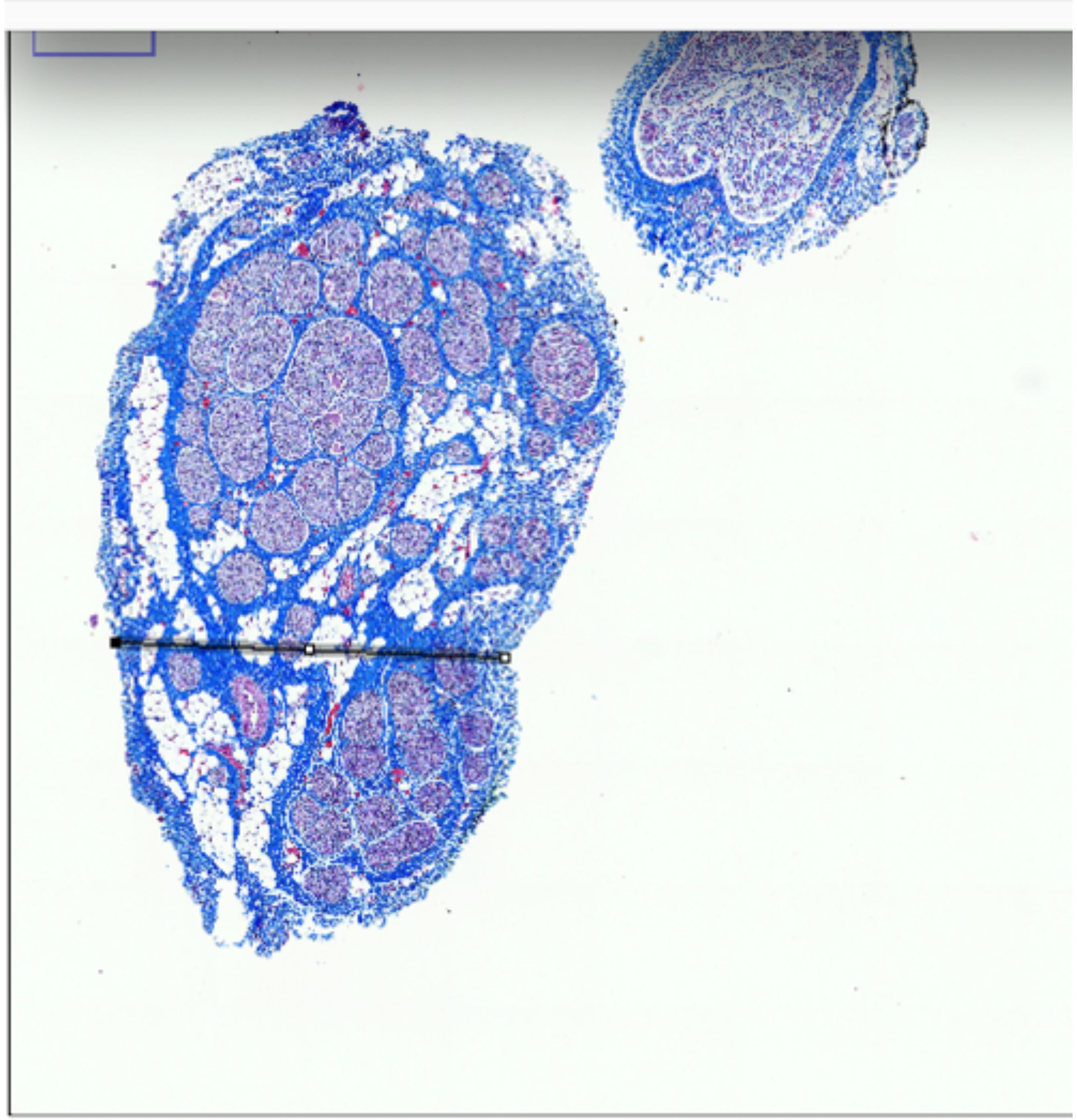

P845_Nodose_SL_16

widest and narrowest diameters

Pre-nodose

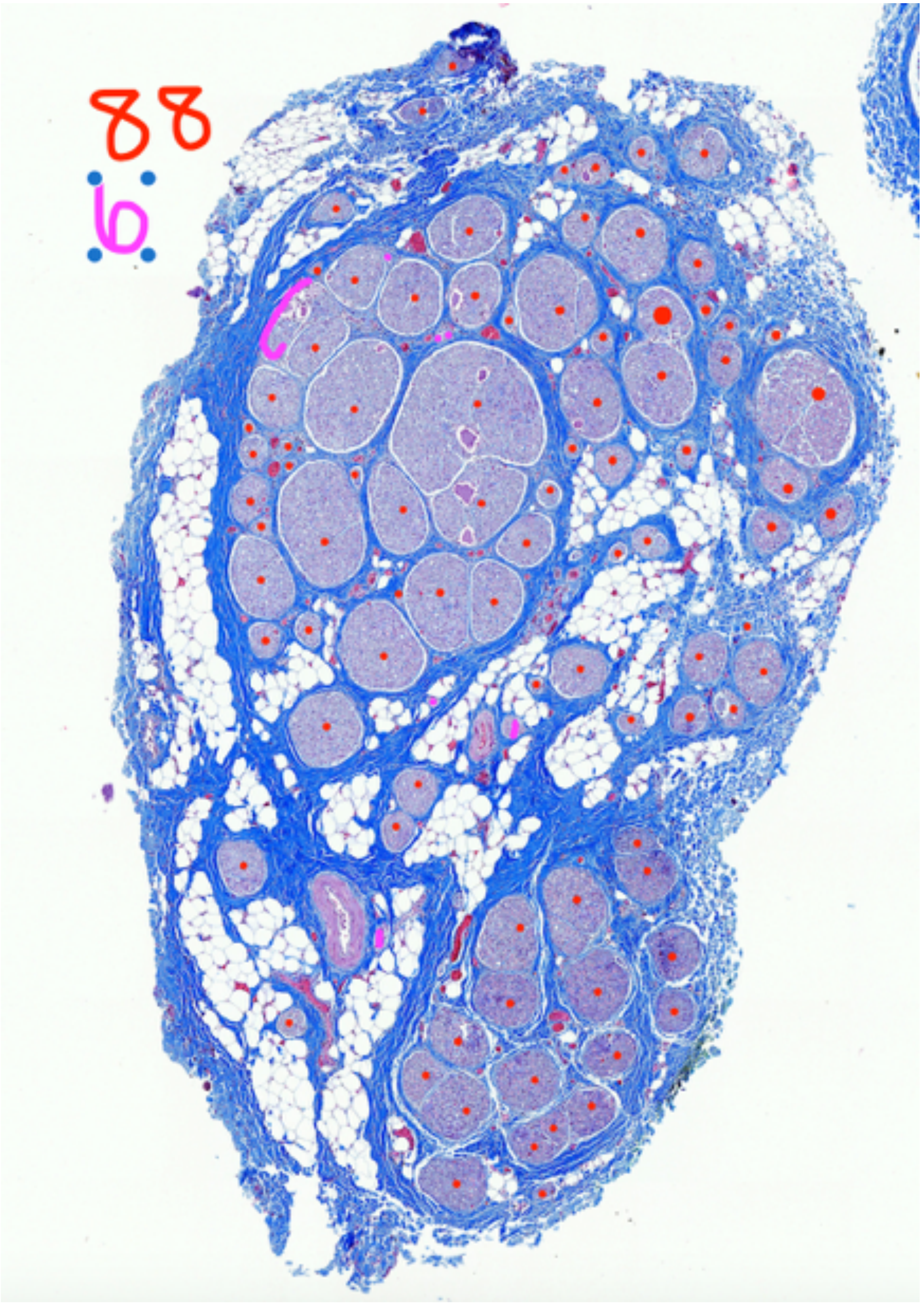


P845_Nodose_SL_16

Fascicle count, 94

### Pre-nodose

Results

| | Area | Mean | Min | Max | Angle | Length |
|---|---|---|---|---|---|---|
| 37 | 0.029 | 188.655 | 50.000 | 255.000 | 5.208 | 0.791 |
| 38 | 0.030 | 186.739 | 32.000 | 255.000 | 8.688 | 0.797 |
| 39 | 0.011 | 179.848 | 8.000 | 255.000 | -83.949 | 0.301 |

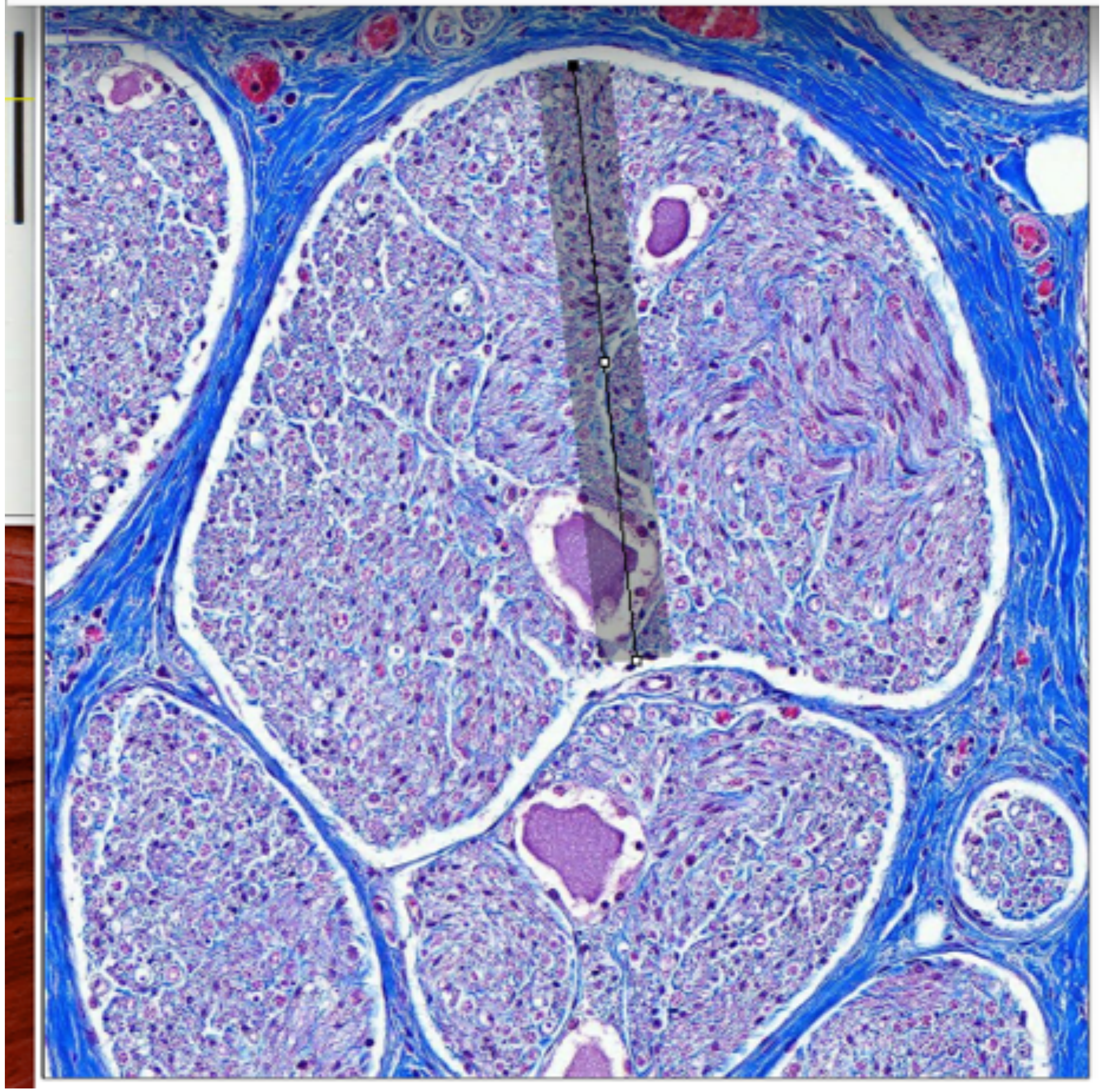

Results

| | Area | Mean | Min | Max | Angle | Length |
|---|---|---|---|---|---|---|
| 38 | 0.030 | 186.739 | 32.000 | 255.000 | 8.688 | 0.797 |
| 39 | 0.011 | 179.848 | 8.000 | 255.000 | -83.949 | 0.301 |
| 40 | 0.015 | 174.962 | 8.000 | 255.000 | 19.592 | 0.397 |

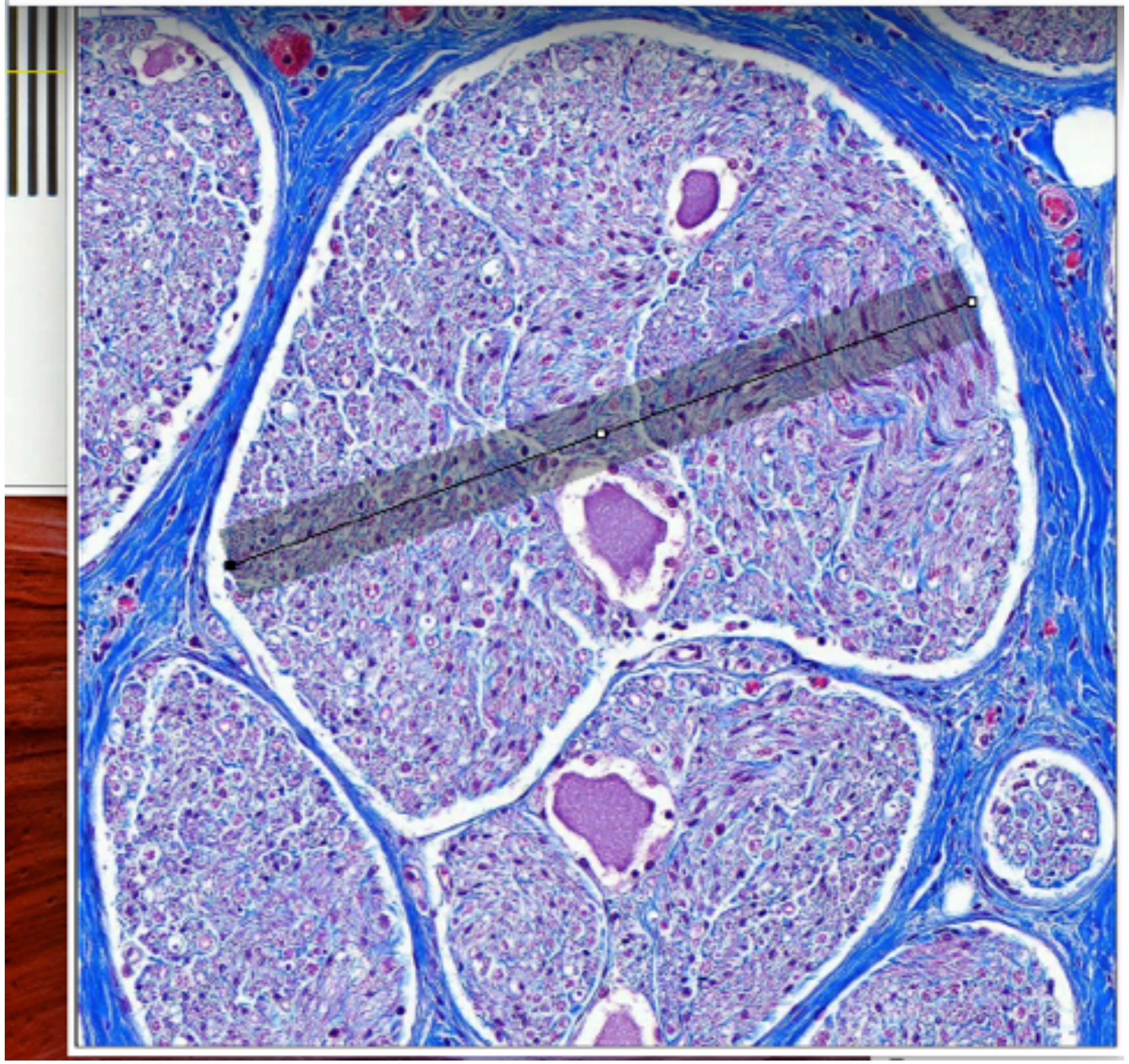

P845_Nodose_SL_16

Largest fascicle_widest and narrowest diameters

### Mid-VN

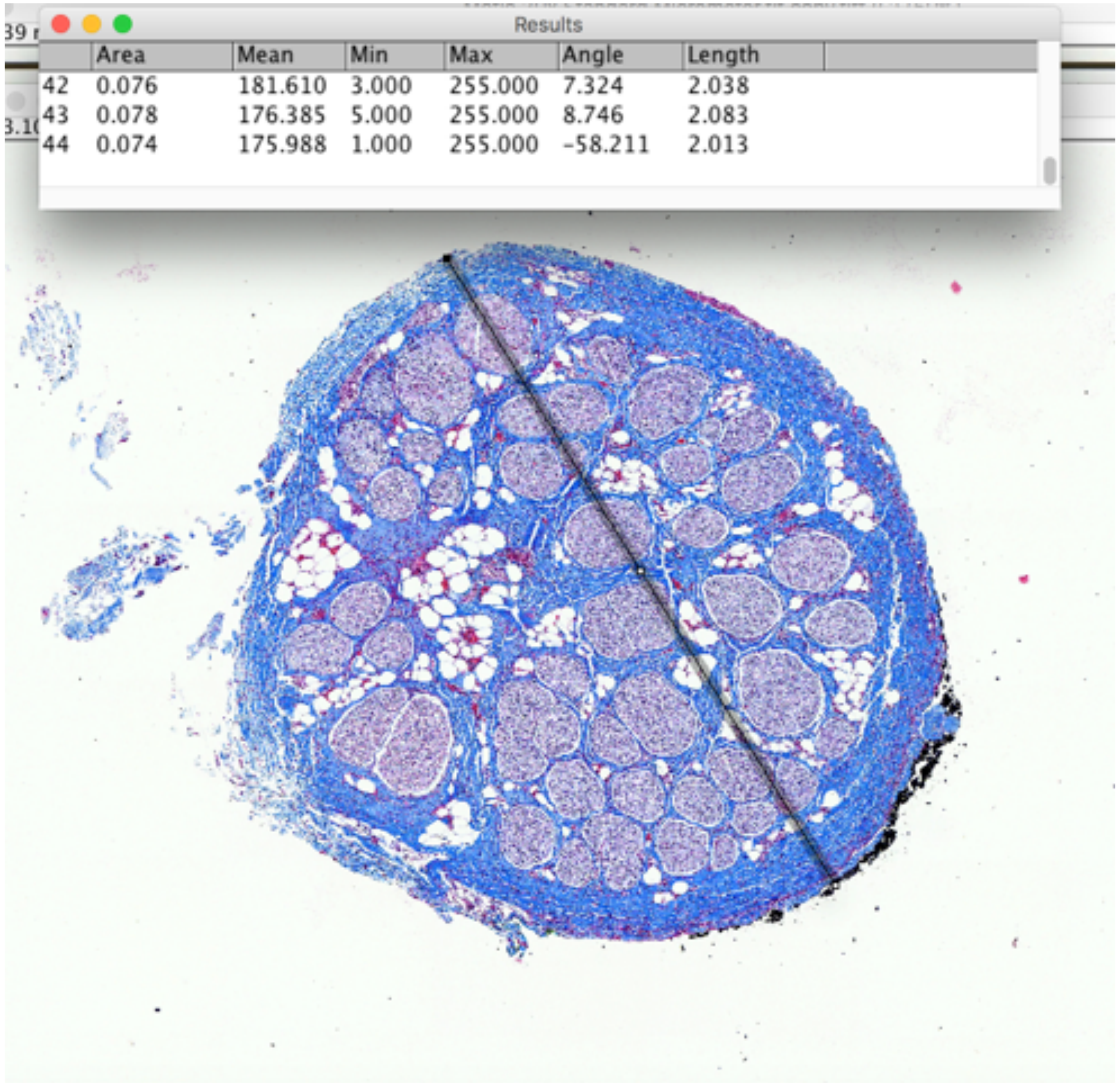


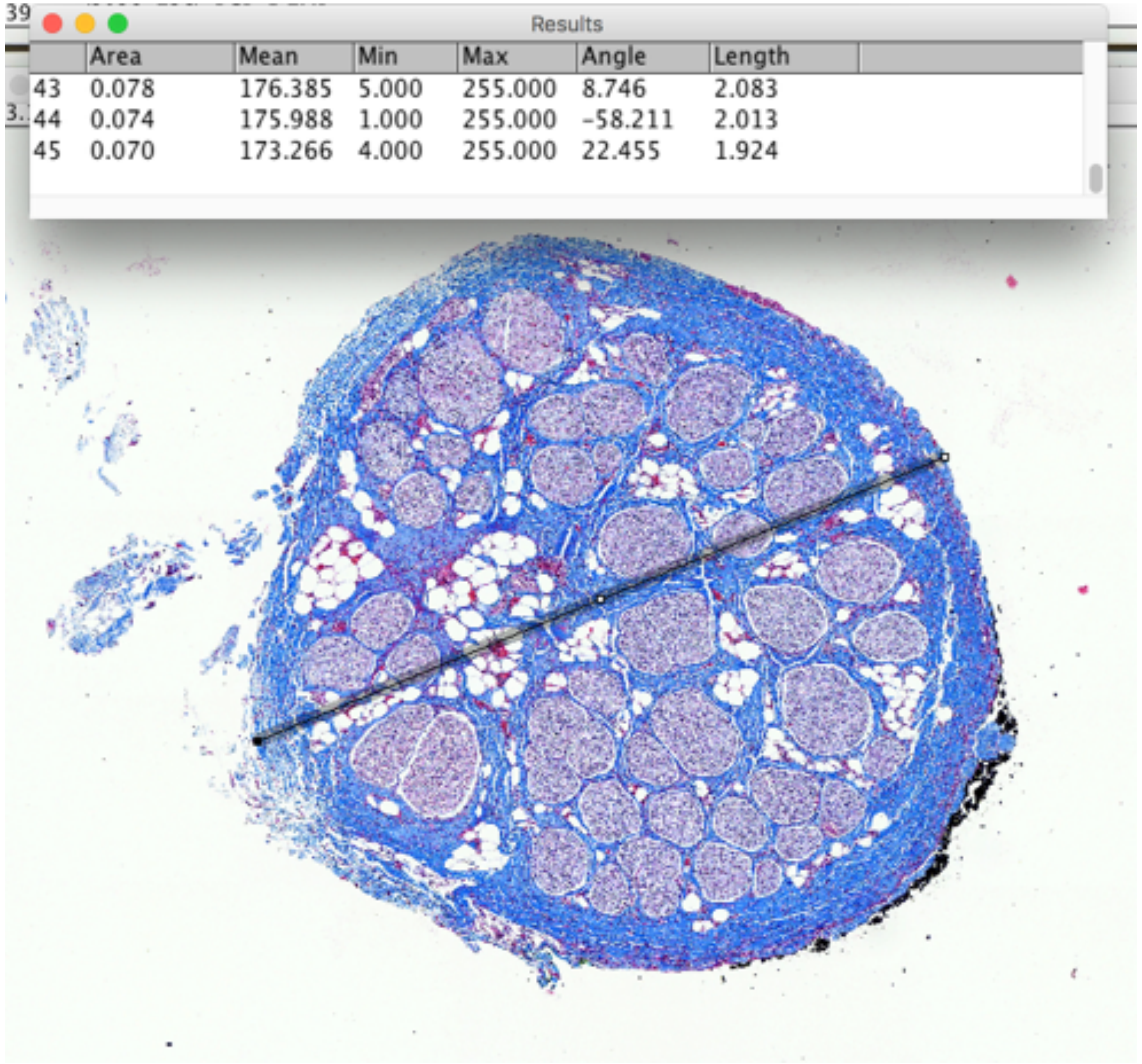


P845_VagusA_3_Cranial contact

widest and narrowest diameters

### Mid-VN

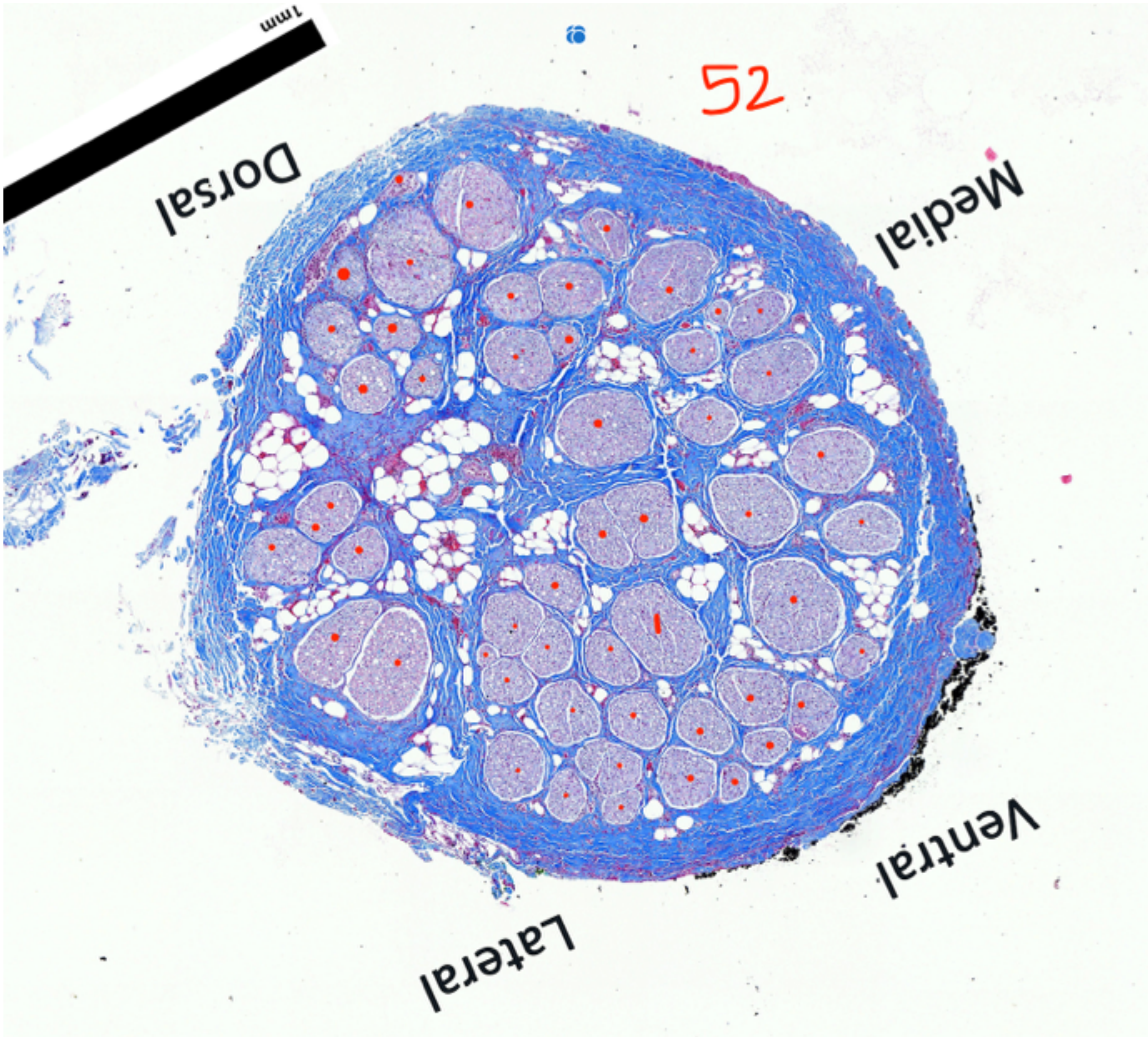


P845_VagusA_3_Cranial contact

Fascicle Count, 52

### Mid-VN

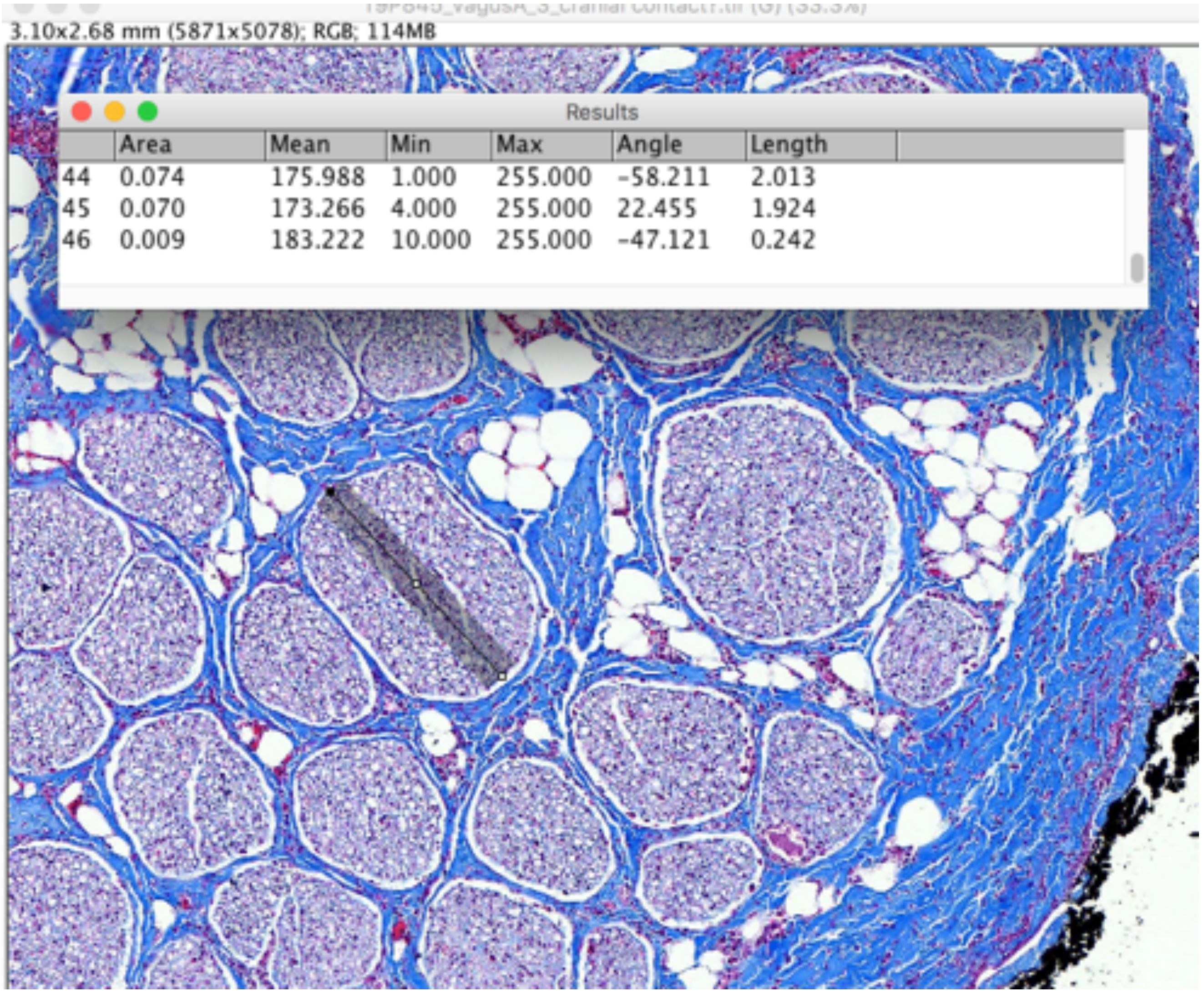


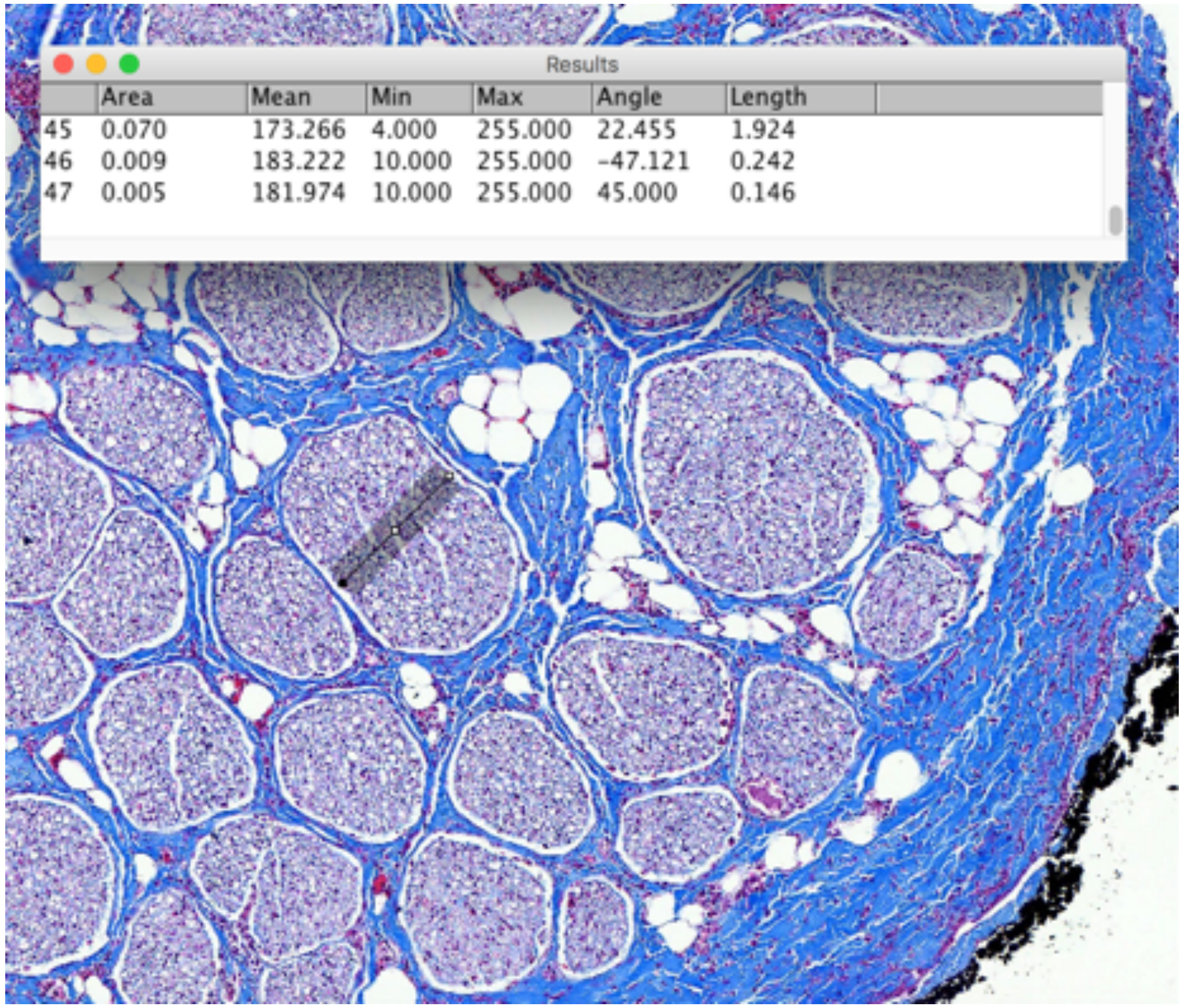


P845_VagusA_3_Cranial contact     largest fascicle_widest and narrowest diameters

### Mid-VN

Results

| | Area | Mean | Min | Max | Angle | Length |
|---|---|---|---|---|---|---|
| 3 | 0.011 | 154.088 | 3 | 255 | -48.180 | 0.215 |
| 4 | 0.018 | 146.596 | 15 | 255 | -71.346 | 0.350 |
| 5 | 0.006 | 166.922 | 26 | 255 | 0.000 | 0.118 |

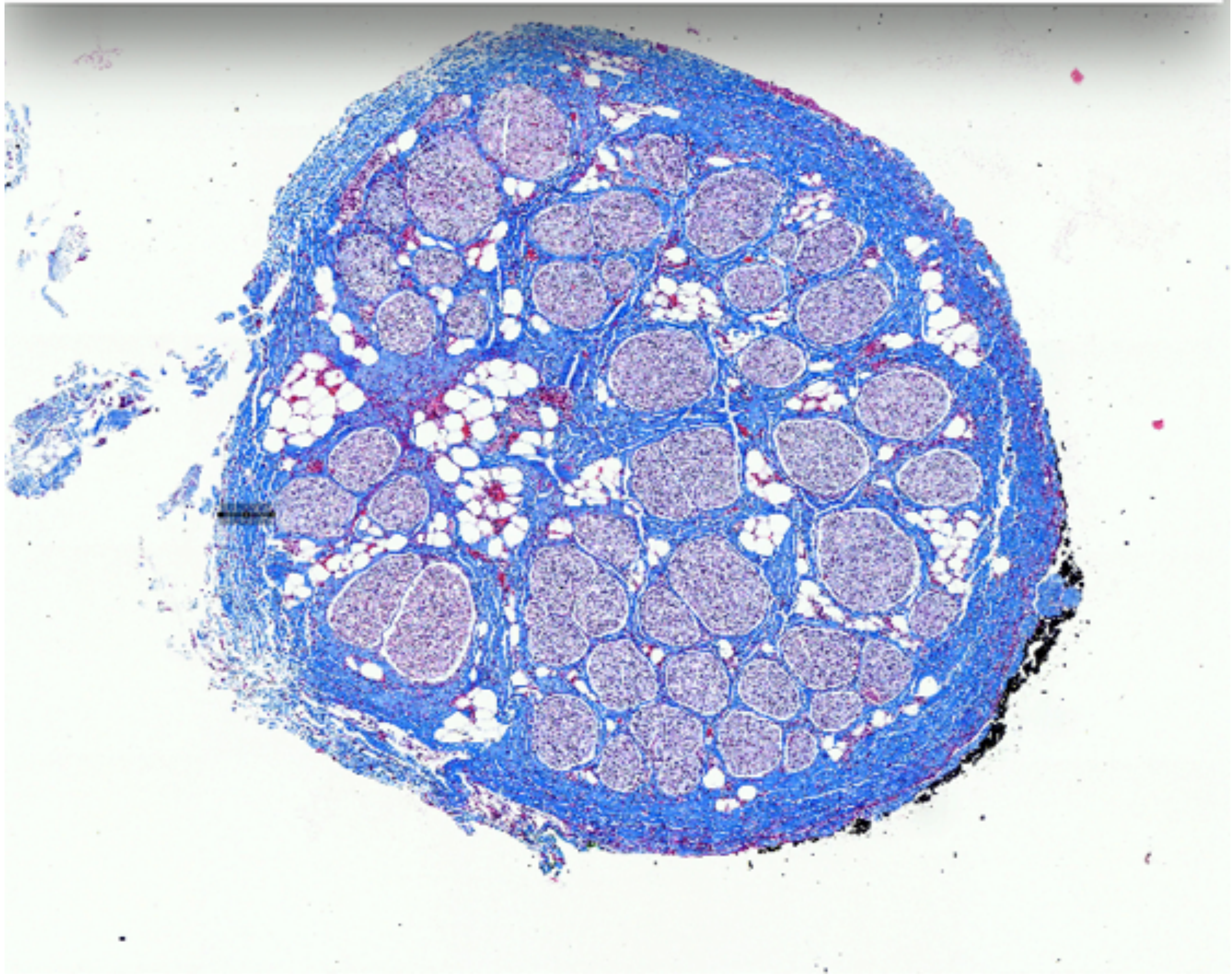

P845_VagusA_3

Fascicle depth

RL

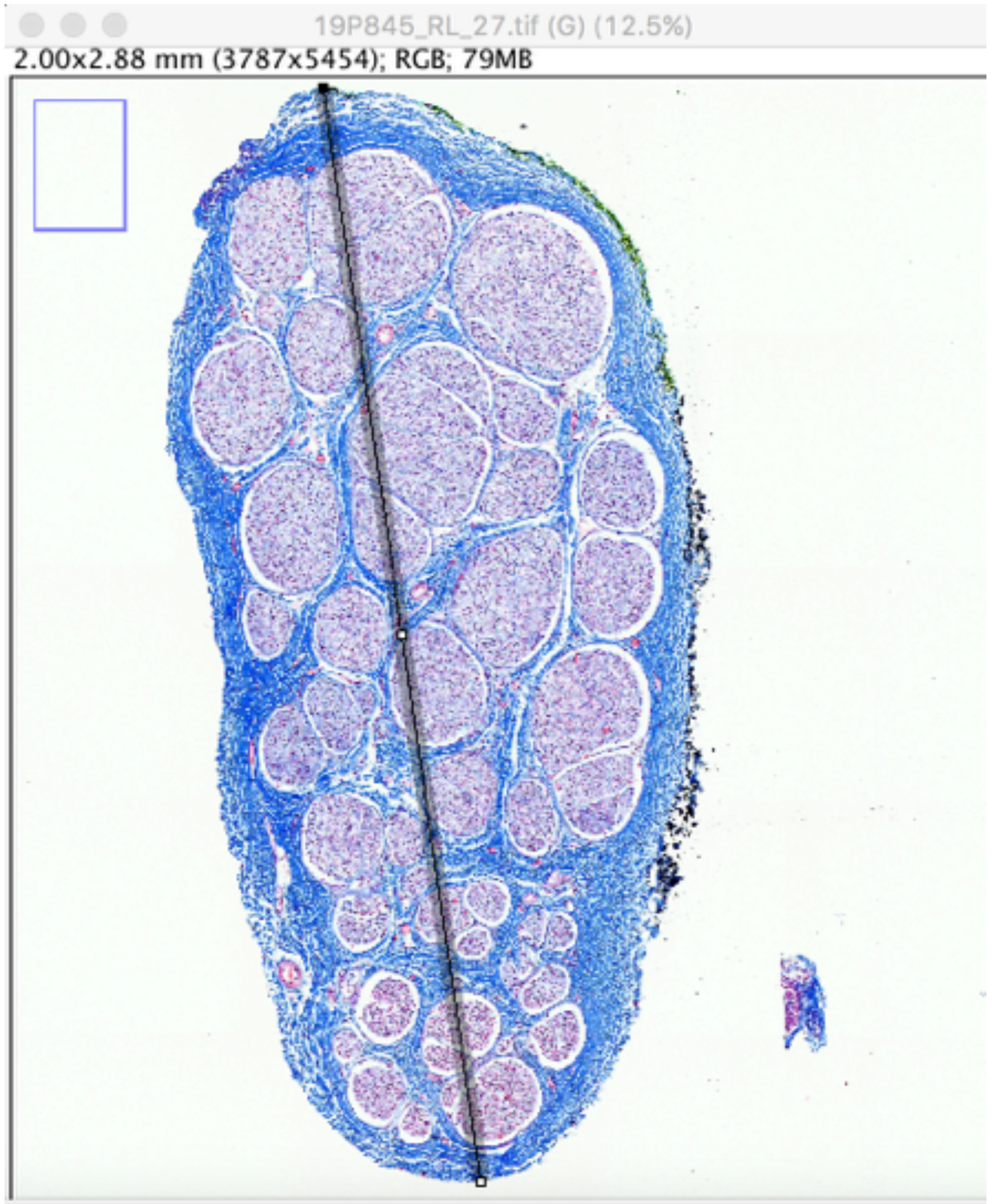


Results

| [illegible]a | Mean | Min | Max | Angle | Length |
|---|---|---|---|---|---|
| [illegible]9 | 183.222 | 10.000 | 255.000 | -47.121 | 0.242 |
| [illegible]5 | 181.974 | 10.000 | 255.000 | 45.000 | 0.146 |
| [illegible]4 | 193.017 | 4.000 | 255.000 | -81.809 | 2.253 |

P845_RL_27

Results

| | Area | Mean | Min | Max | Angle | Length |
|---|---|---|---|---|---|---|
| 1 | 0.081 | 168.558 | 6 | 255 | -2.369 | 1.533 |
| 2 | 0.022 | 177.834 | 10 | 255 | -0.591 | 0.410 |

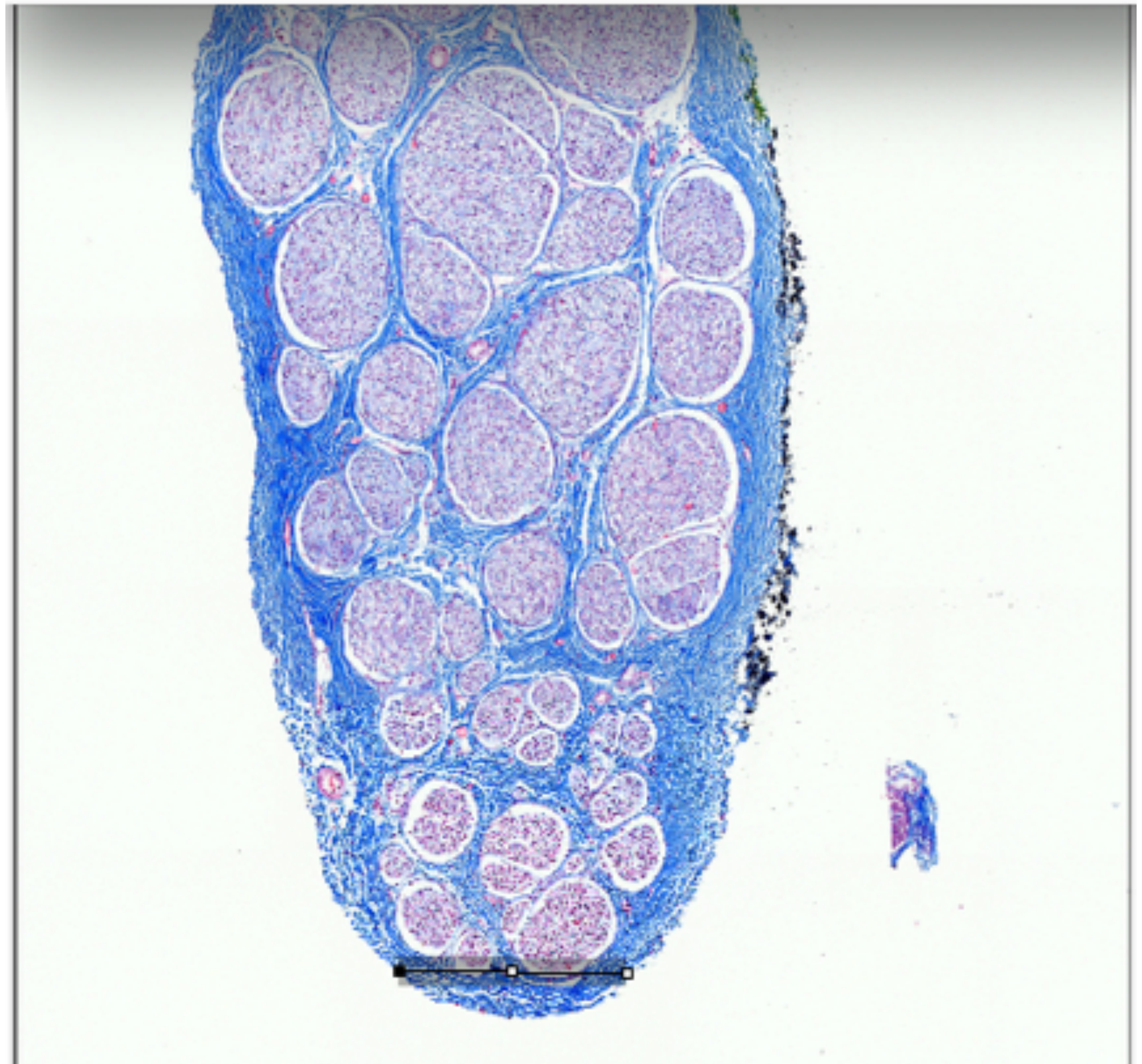

diameter taken at narrowest region near a fascicle

widest and narrowest diameter

RL

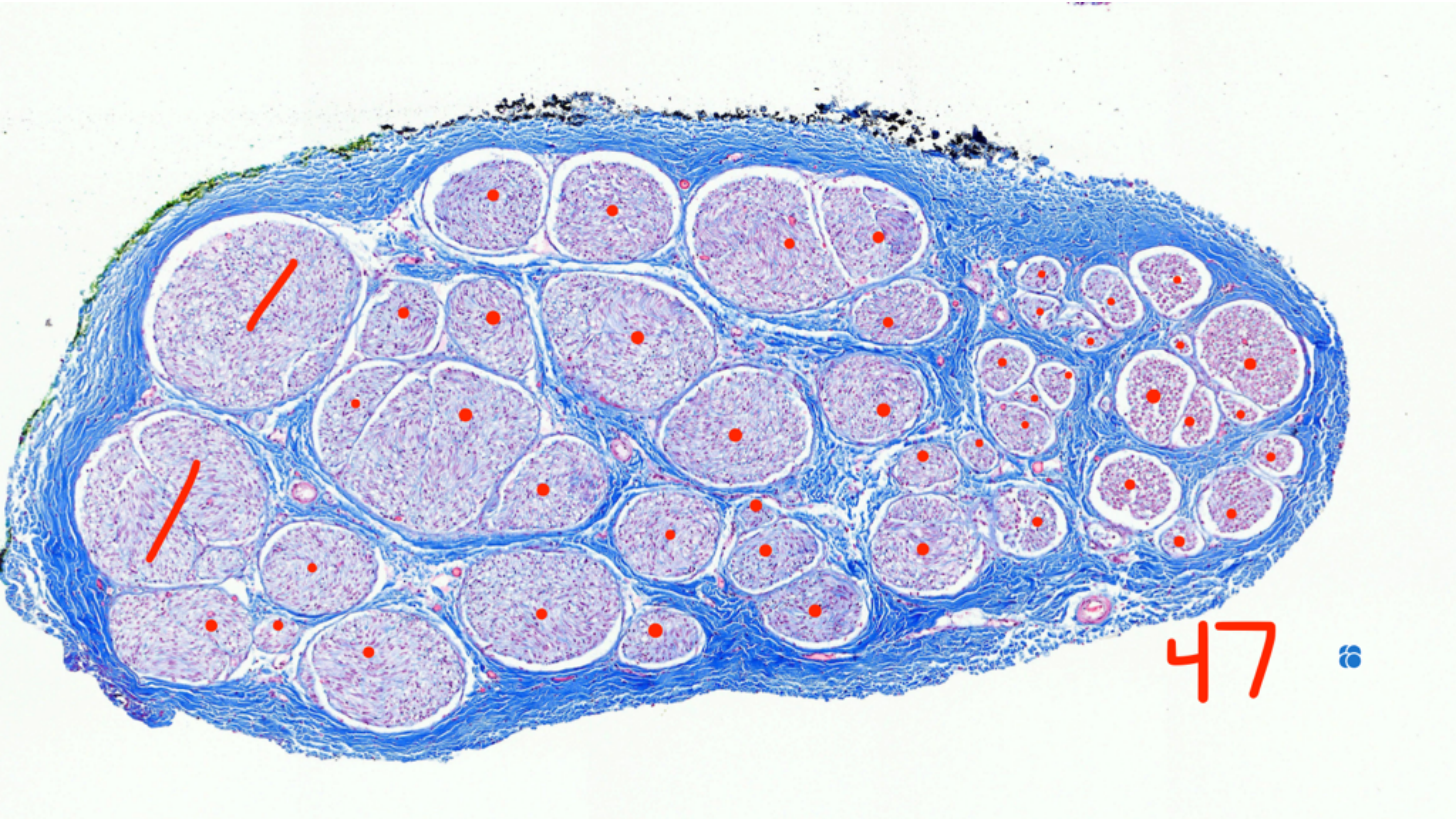


P845_RL_27

Fascicle count, 47

# RL

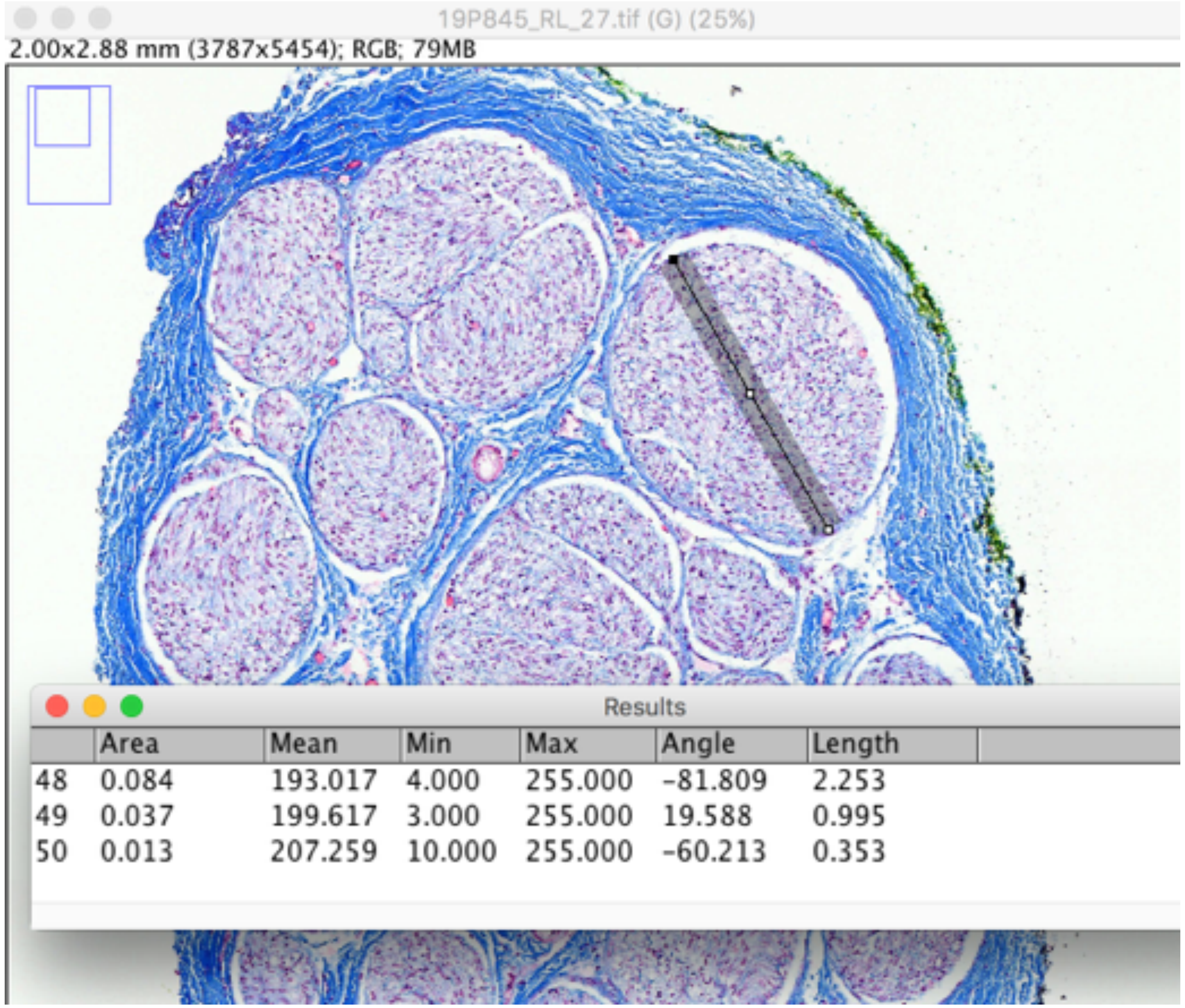


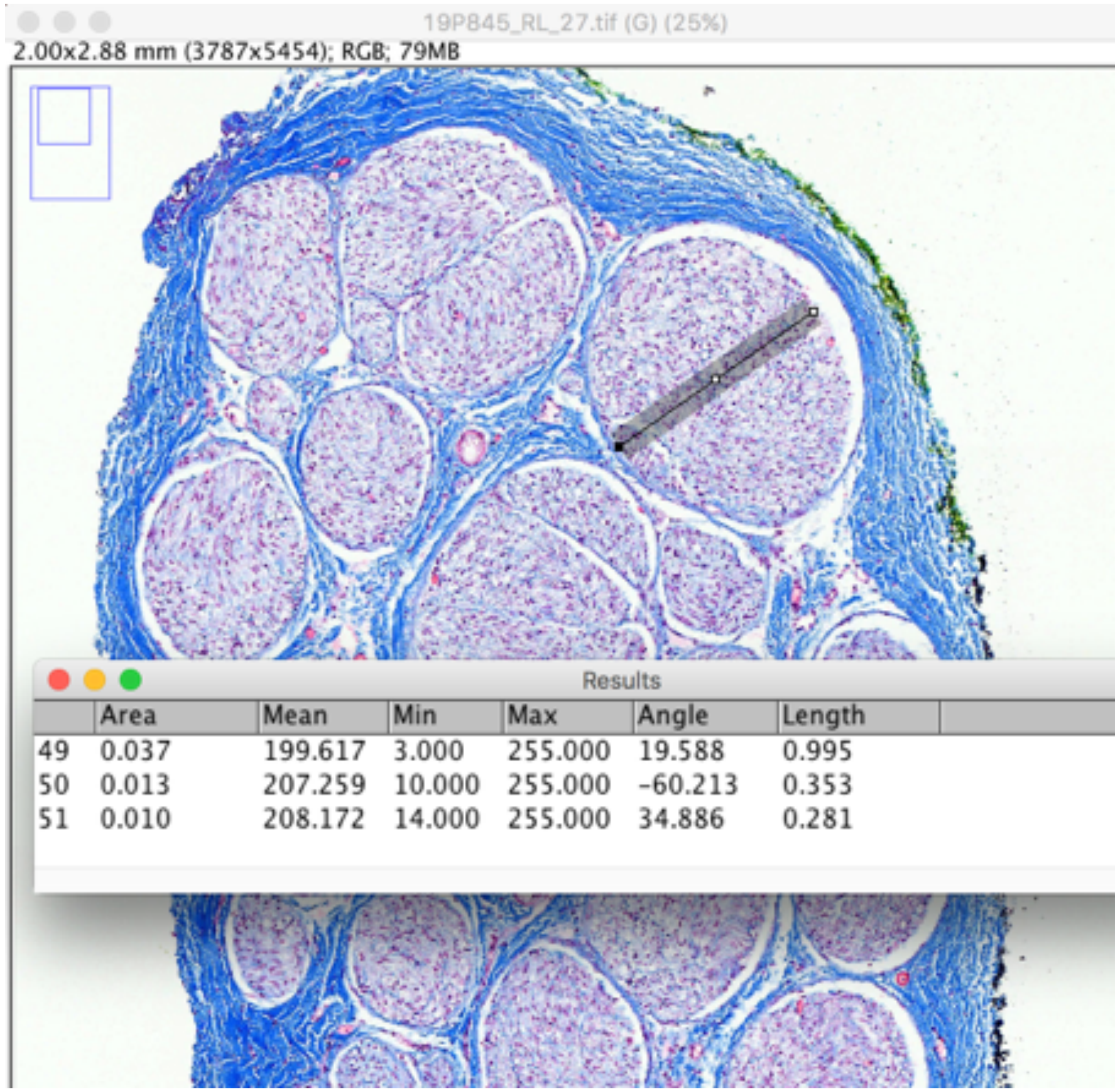


P845_RL_27

Largest fascicle_widest and narrowest diameter

Subject 11_P853

Mid-VN

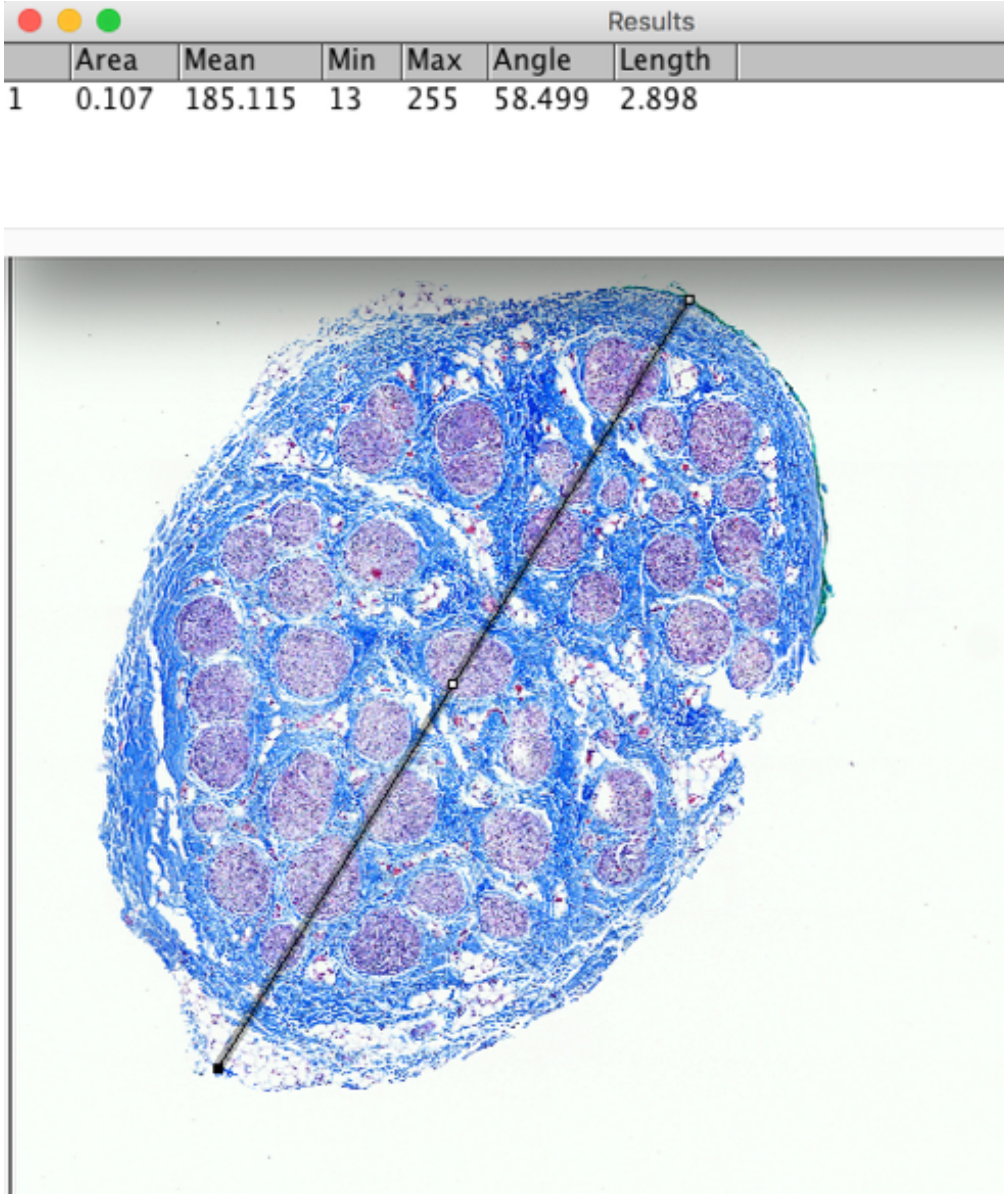


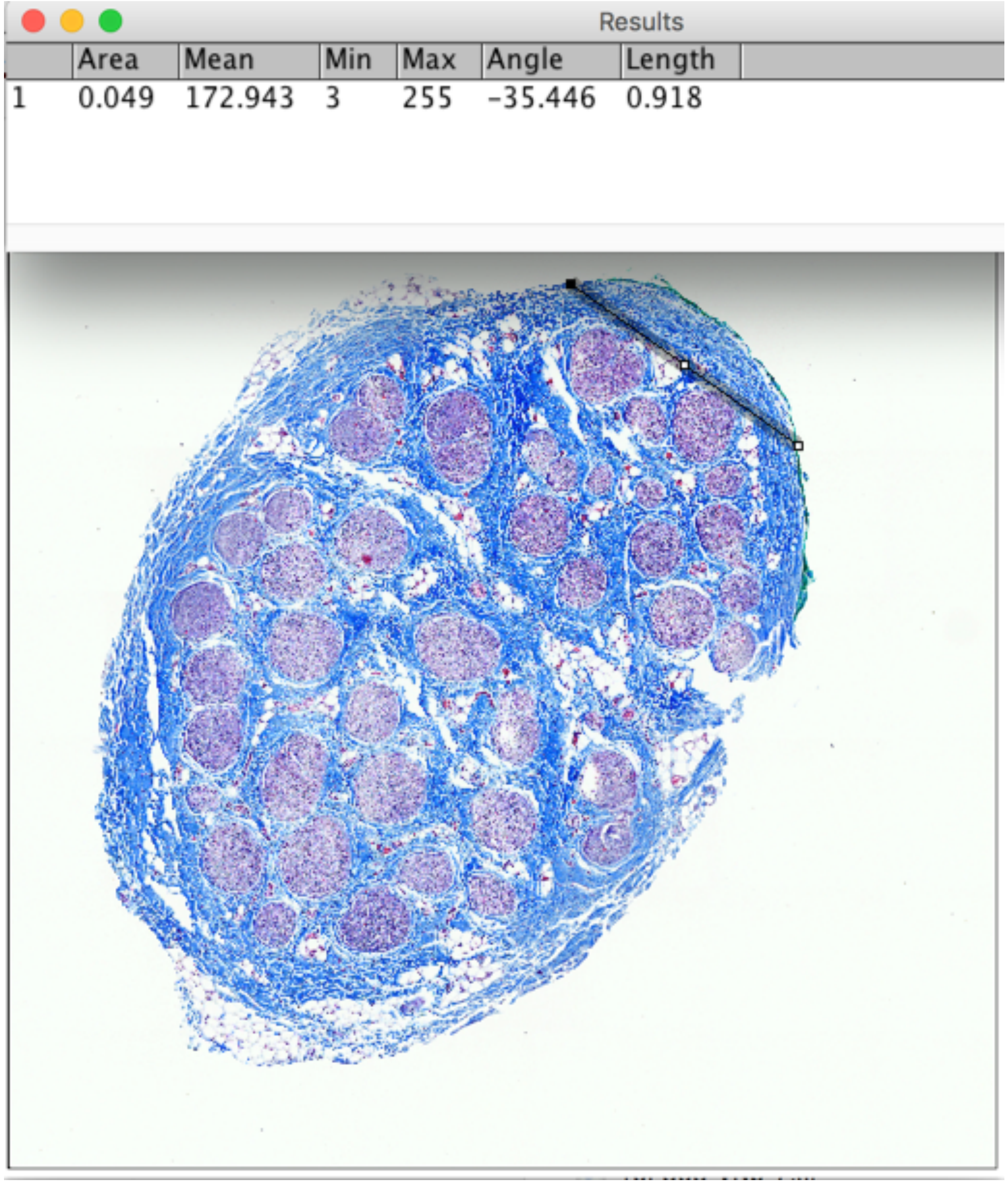


P853_VNb_4

Widest and narrowest diameter

Mid-VN

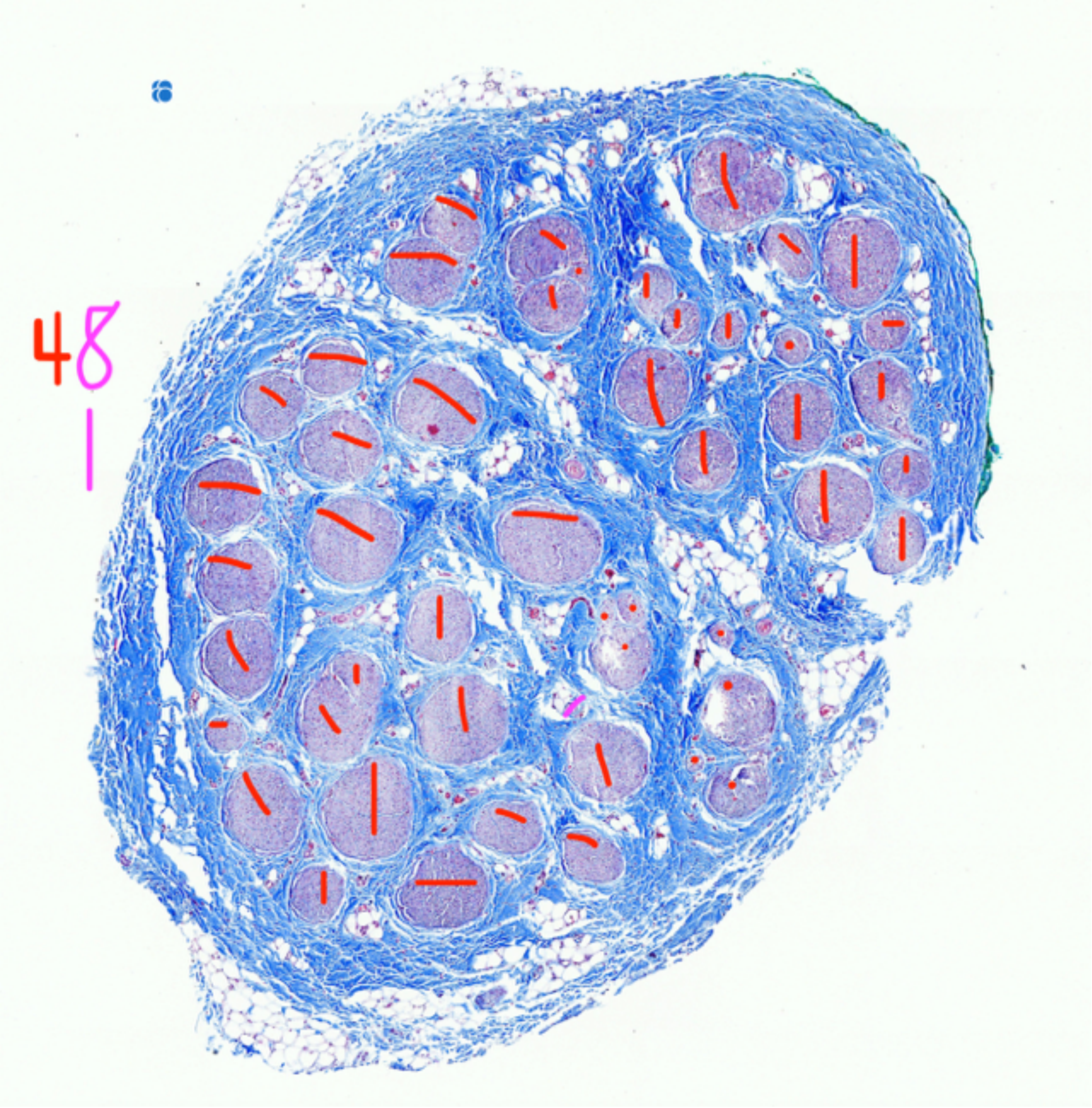


P853_VNb_4

Fascicle count, 49

### Mid-VN

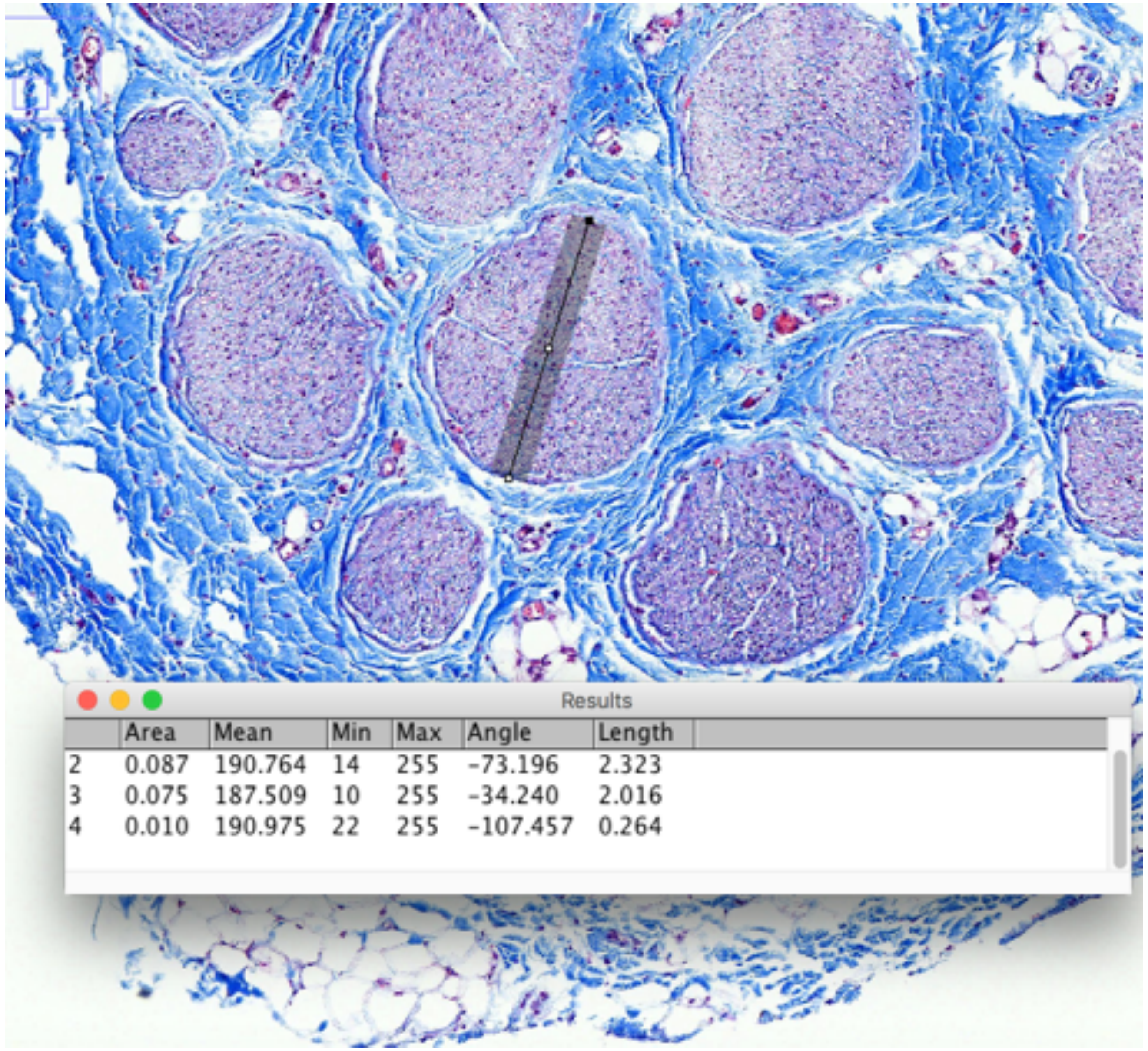

| | Area | Mean | Min | Max | Angle | Length |
|---|---|---|---|---|---|---|
| 2 | 0.087 | 190.764 | 14 | 255 | -73.196 | 2.323 |
| 3 | 0.075 | 187.509 | 10 | 255 | -34.240 | 2.016 |
| 4 | 0.010 | 190.975 | 22 | 255 | -107.457 | 0.264 |

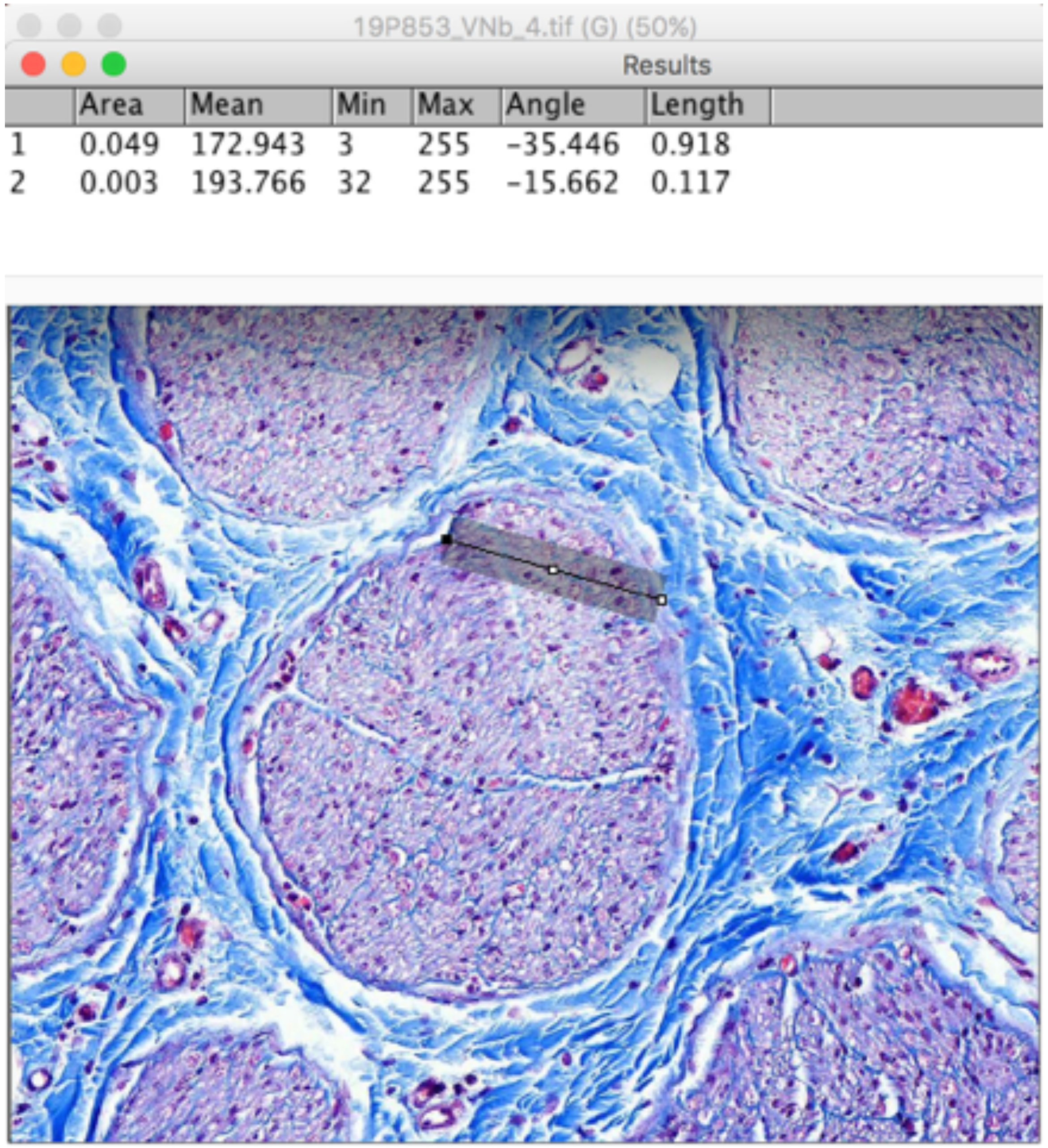



| | Area | Mean | Min | Max | Angle | Length |
|---|---|---|---|---|---|---|
| 1 | 0.049 | 172.943 | 3 | 255 | -35.446 | 0.918 |
| 2 | 0.003 | 193.766 | 32 | 255 | -15.662 | 0.117 |

P853_VNb_4

Largest fascicle_Widest and narrowest diameter

### Mid-VN

Results

| | Area | Mean | Min | Max | Angle | Length |
|---|---|---|---|---|---|---|
| 4 | 0.018 | 146.596 | 15 | 255 | -71.346 | 0.350 |
| 5 | 0.006 | 166.922 | 26 | 255 | 0.000 | 0.118 |
| 6 | 0.006 | 185.318 | 16 | 255 | -36.048 | 0.110 |

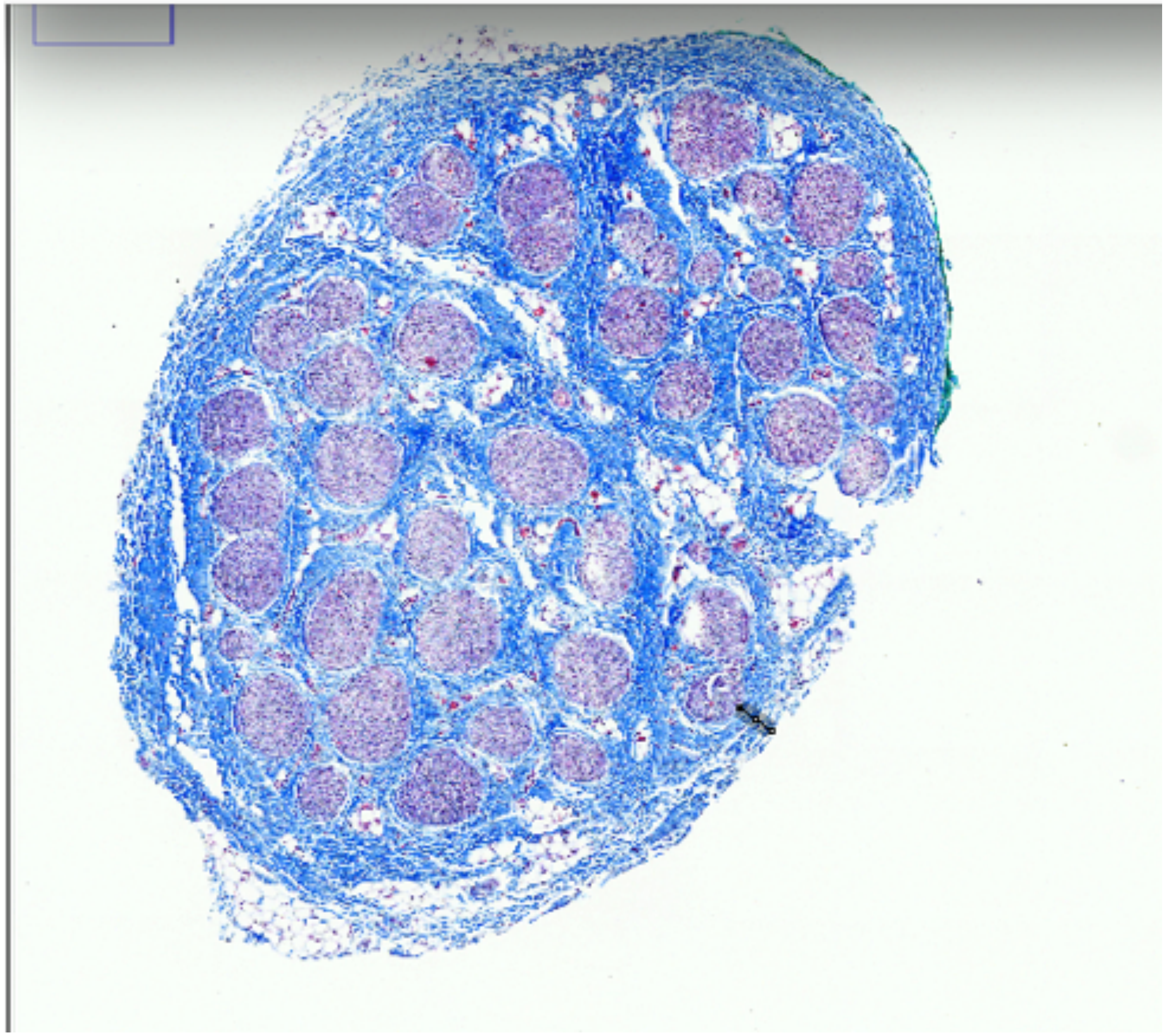

P853_VNb_4

Fascicle depth

Subject 1_P786

### Pre-nodose

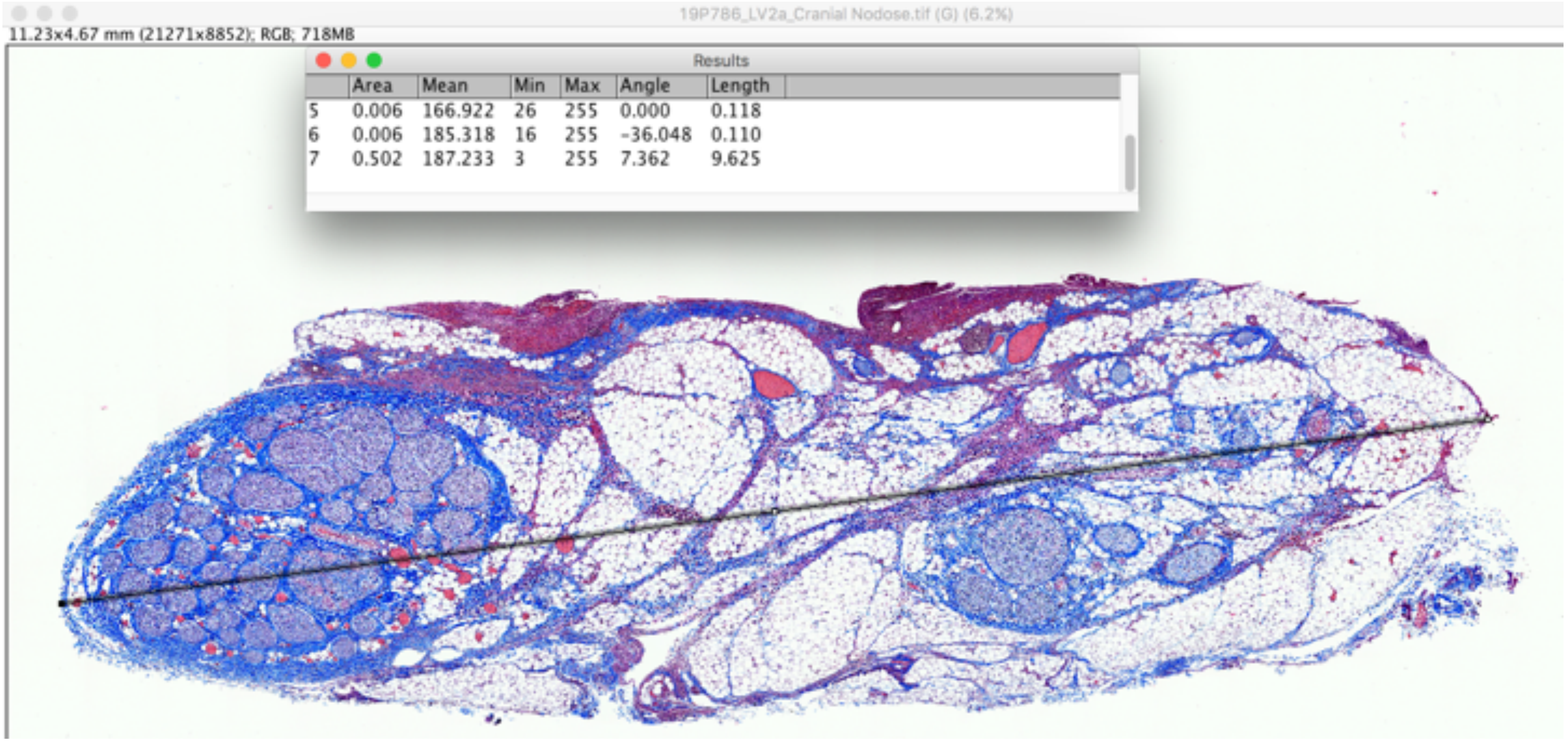


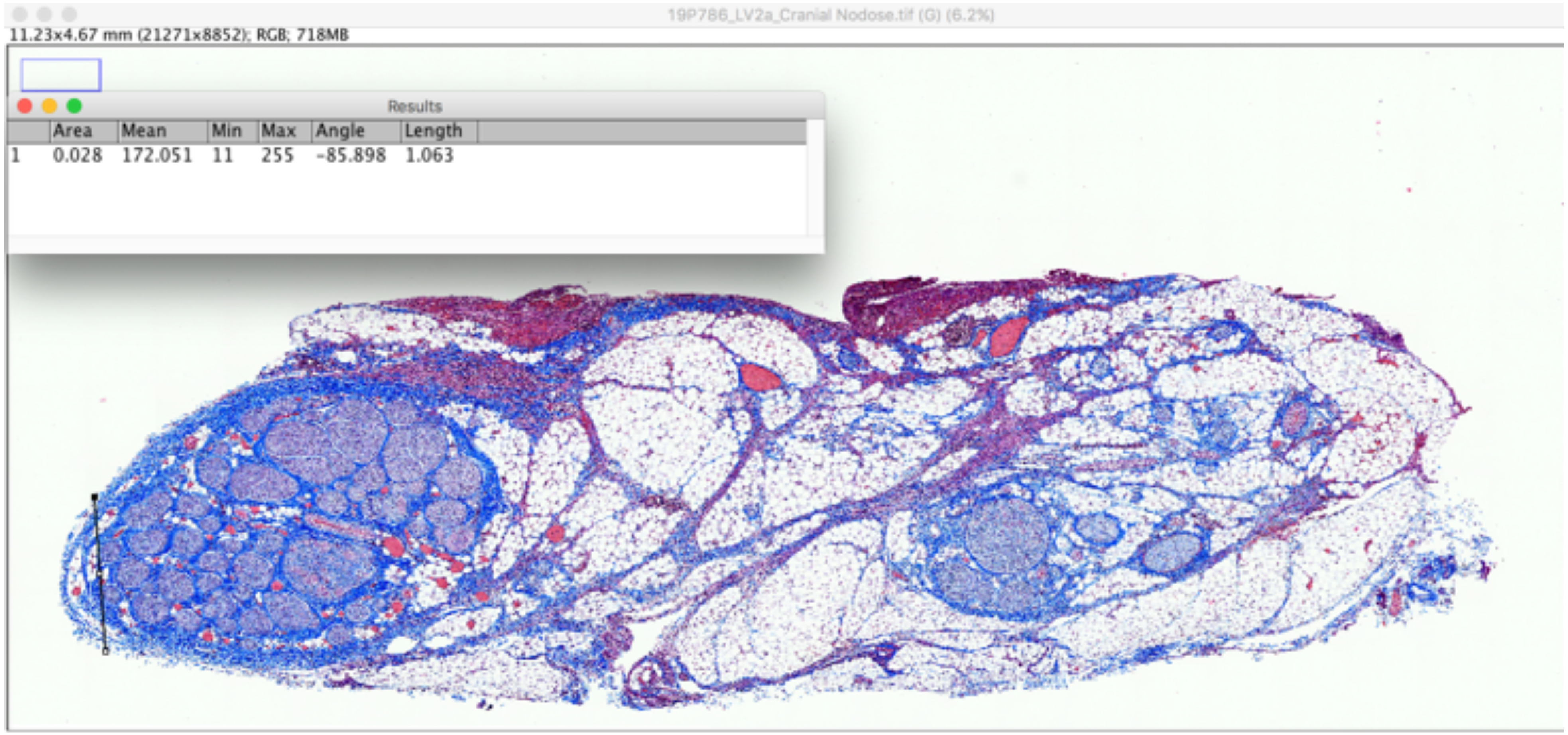


19P786_LV2b_cranial nodose

Widest and narrowest diameter

Pre-nodose

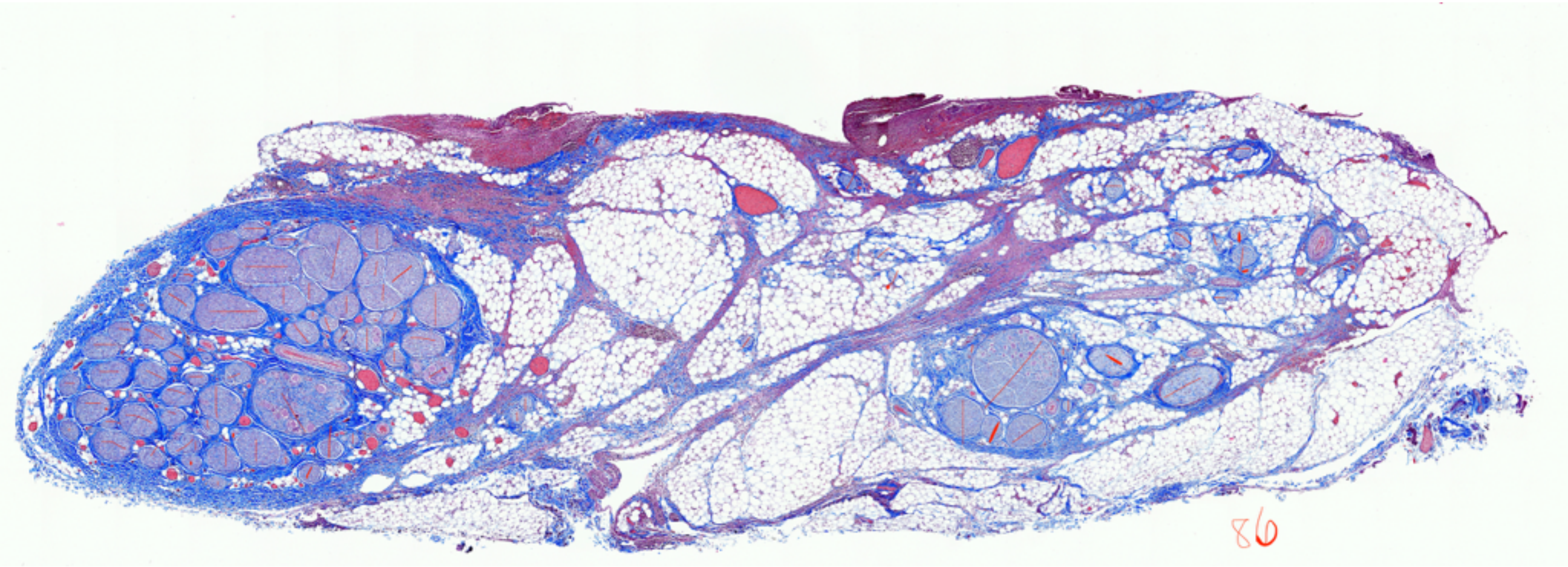


19P786_LV2b_cranial nodose

Fascicle count, 86

### Pre-nodose

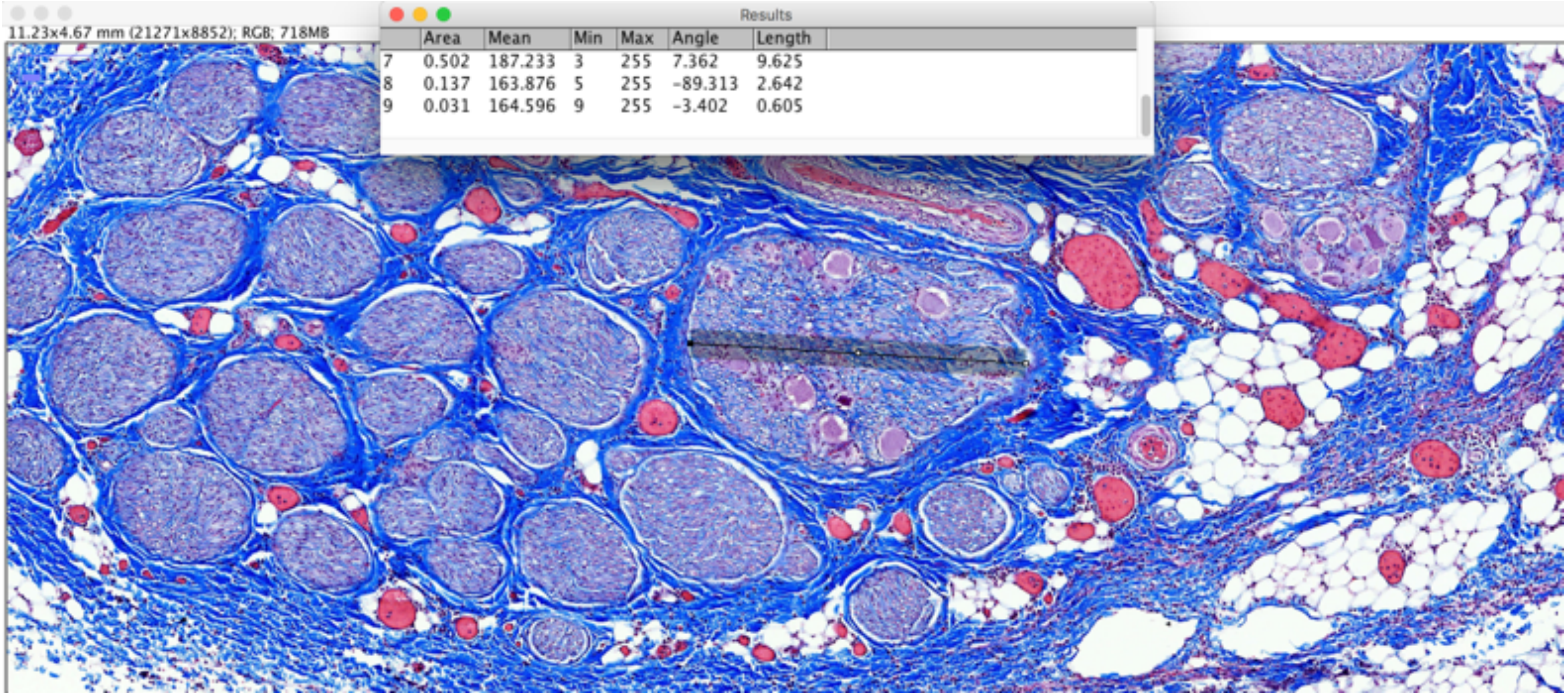


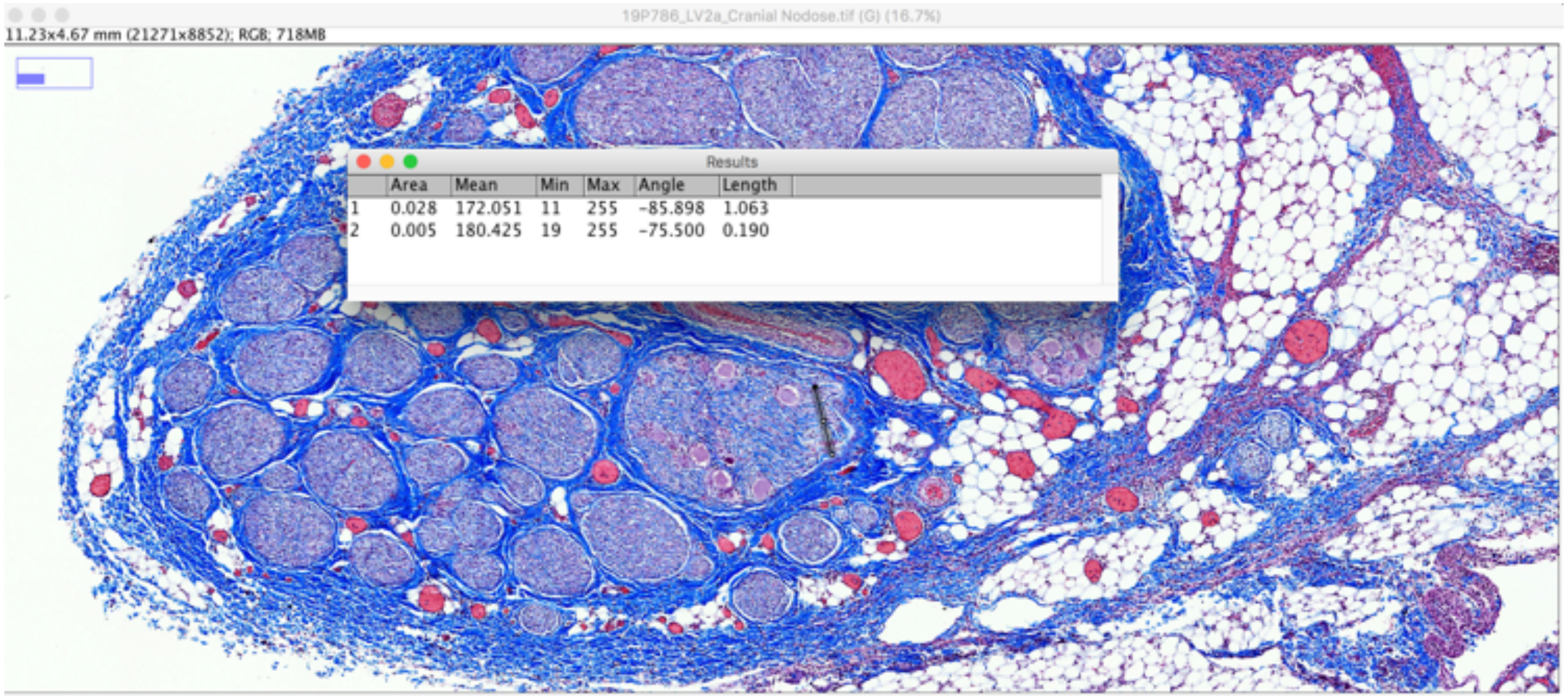


19P786_LV2b_cranial nodose    Largest fascicle_widest and narrowest diameter

### Mid-VN

Results

| | Area | Mean | Min | Max | Angle | Length |
|---|---|---|---|---|---|---|
| 4 | 0.010 | 190.975 | 22 | 255 | -107.457 | 0.264 |
| 5 | 0.008 | 186.969 | 11 | 255 | -24.341 | 0.219 |
| 6 | 0.145 | 190.955 | 3 | 255 | -103.529 | 3.936 |

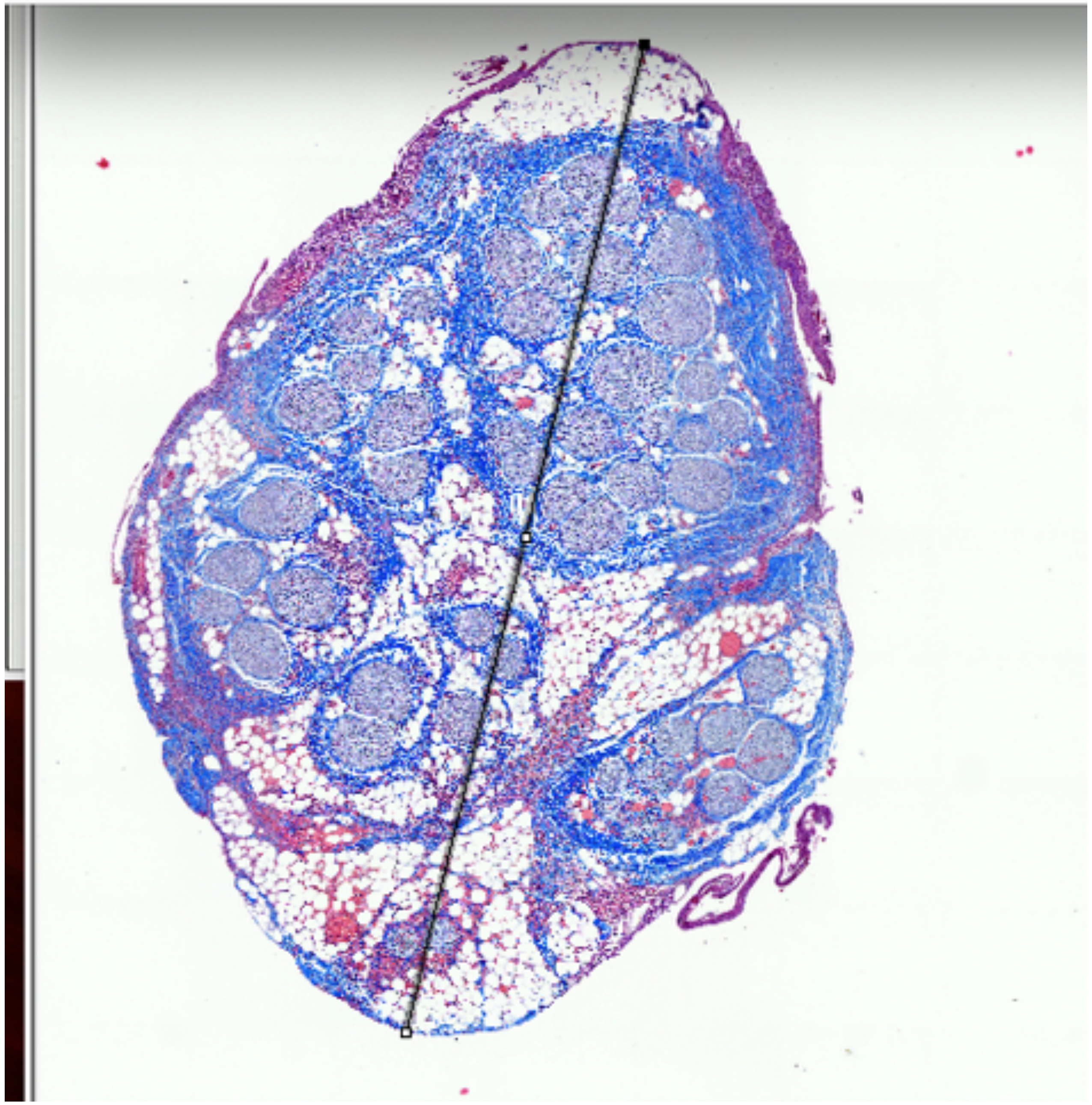

Results

| | Area | Mean | Min | Max | Angle | Length |
|---|---|---|---|---|---|---|
| 1 | 0.028 | 172.051 | 11 | 255 | -85.898 | 1.063 |
| 2 | 0.005 | 180.425 | 19 | 255 | -75.500 | 0.190 |
| 3 | 0.034 | 165.655 | 9 | 255 | -9.273 | 1.258 |

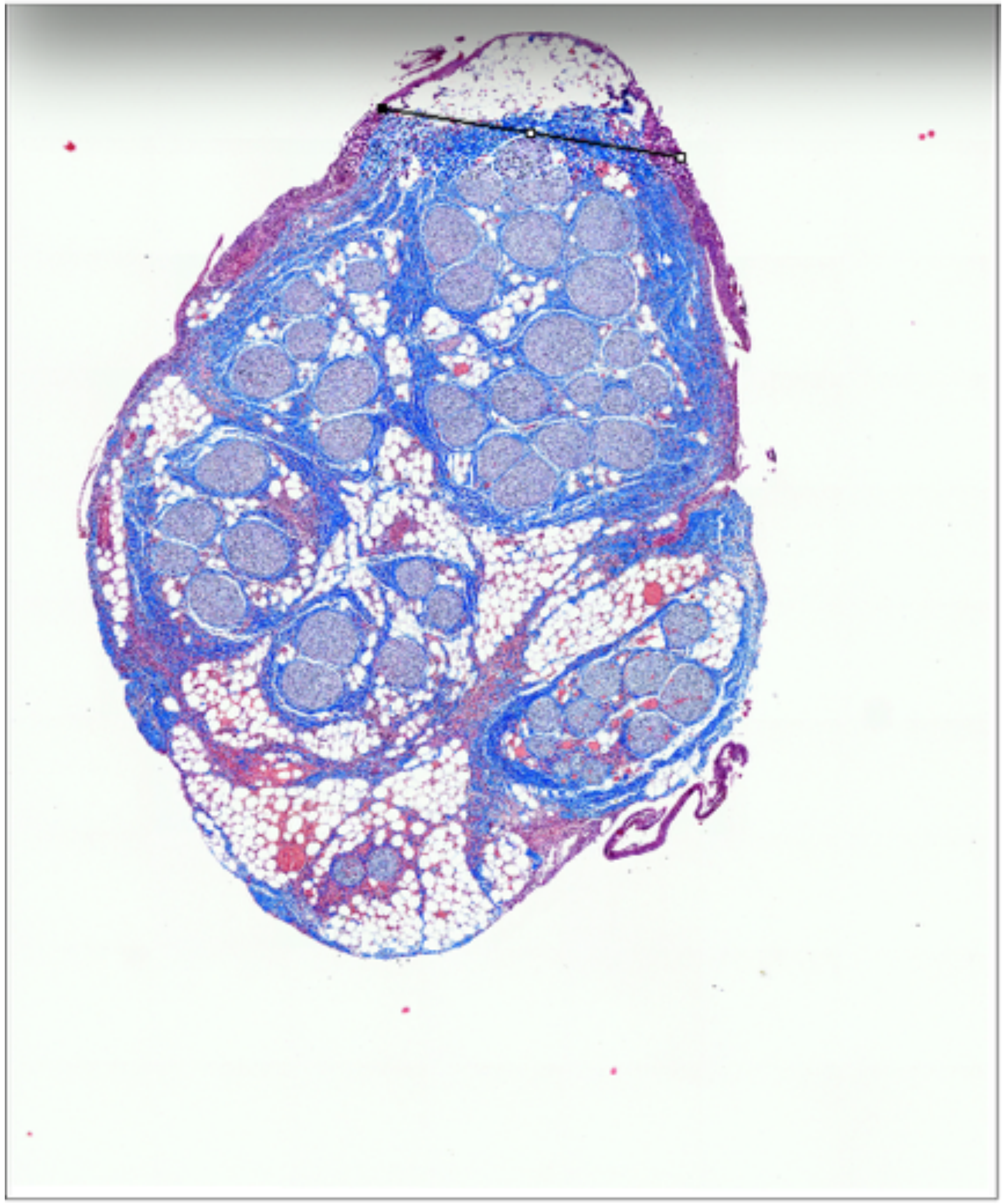

P786_LV1b_LivaNova

Widest and narrowest diameter

Mid-VN

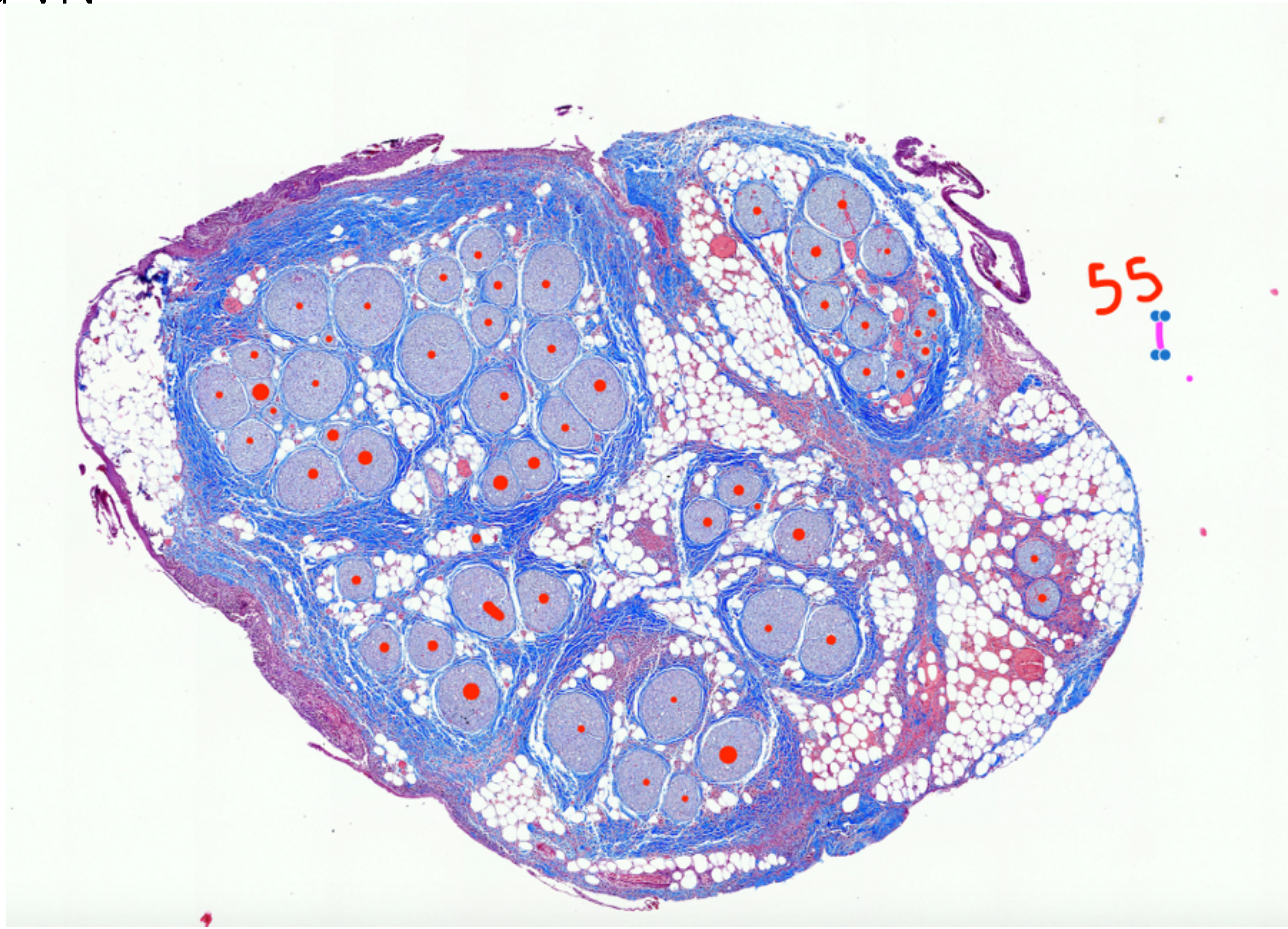


P786_LV1b_LivaNova

Fascicle count, 55

### Mid-VN

19P786_LV1b_LivaNova caudal half.tif (G) (16.7%)

4.12x5.39 mm (7801x10217); RGB; 304MB

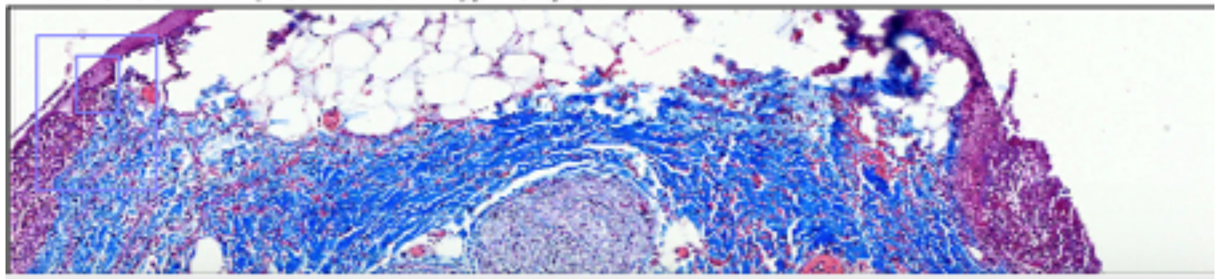

Results

| Area | Mean | Min | Max | Angle | Length |
|---|---|---|---|---|---|
| 0.010 | 184.250 | 12 | 255 | -9.828 | 0.272 |
| 0.009 | 189.742 | 15 | 255 | -16.361 | 0.240 |
| 0.010 | 184.770 | 9 | 255 | 5.156 | 0.282 |

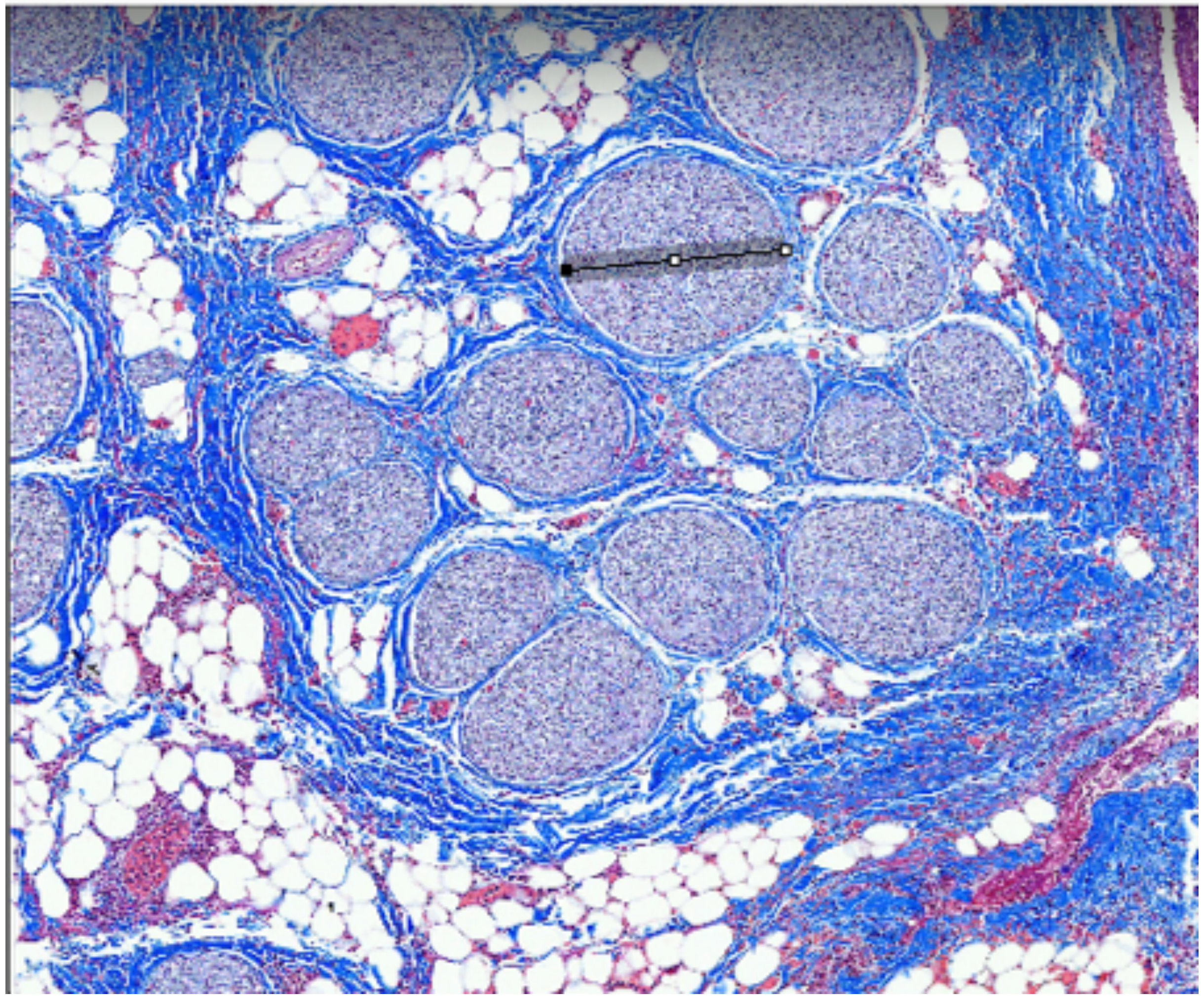

19P786_LV1b_LivaNova caudal half.tif (G) (33.3%)

4.12x5.39 mm (7801x10217); RGB; 304MB

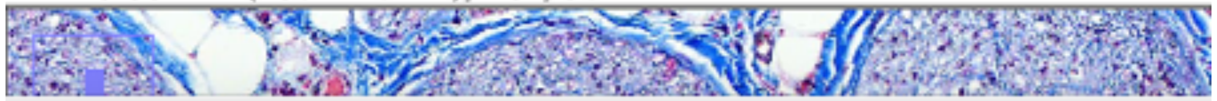

Results

| Area | Mean | Min | Max | Angle | Length |
|---|---|---|---|---|---|
| 0.010 | 184.770 | 9 | 255 | 5.156 | 0.282 |
| 0.009 | 184.742 | 8 | 255 | -86.260 | 0.243 |
| 0.009 | 185.522 | 8 | 255 | -90.000 | 0.244 |

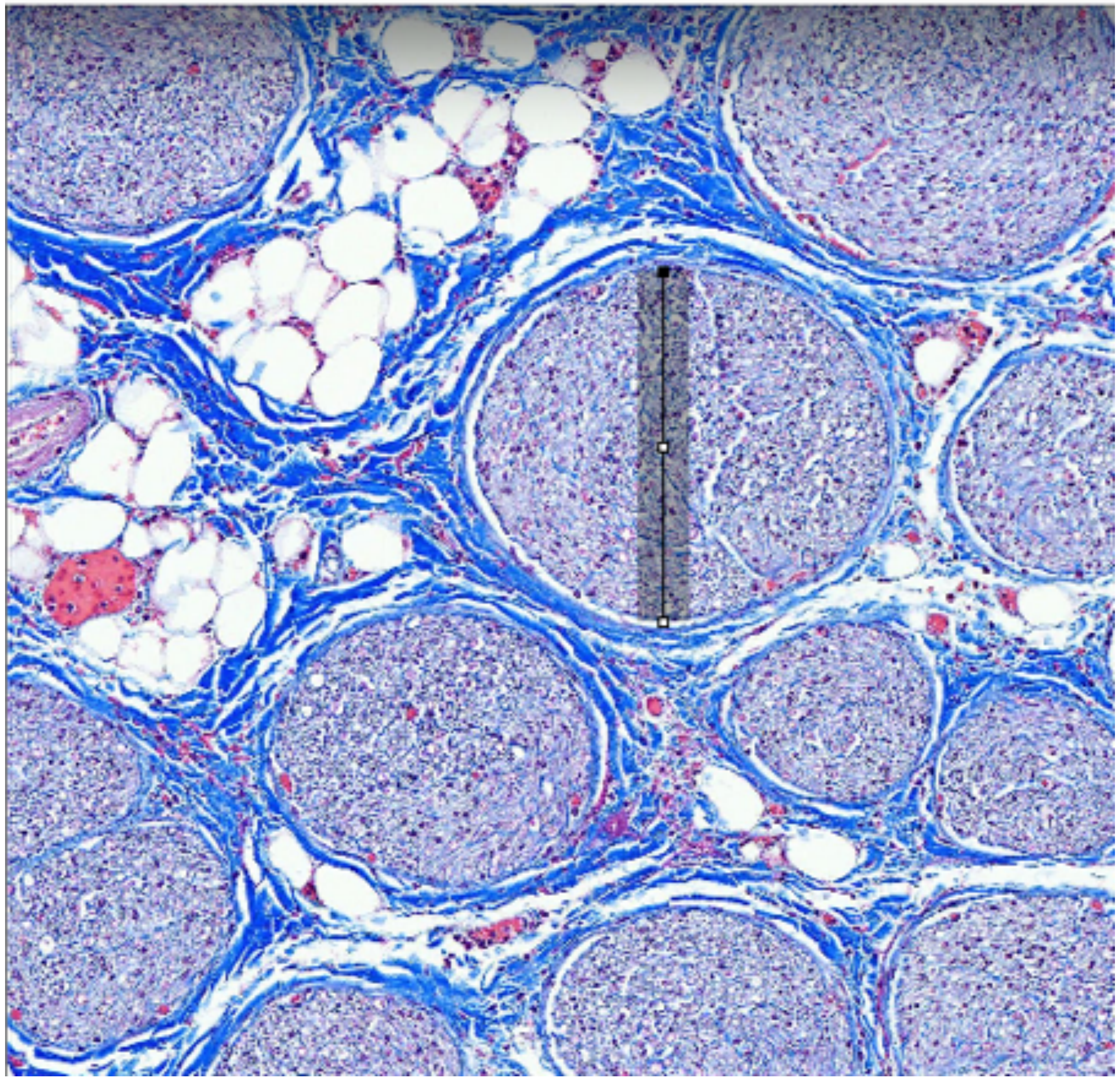

P786_LV1b_LivaNova

Largest fascicle_widest and narrowest diameter

Mid-VN

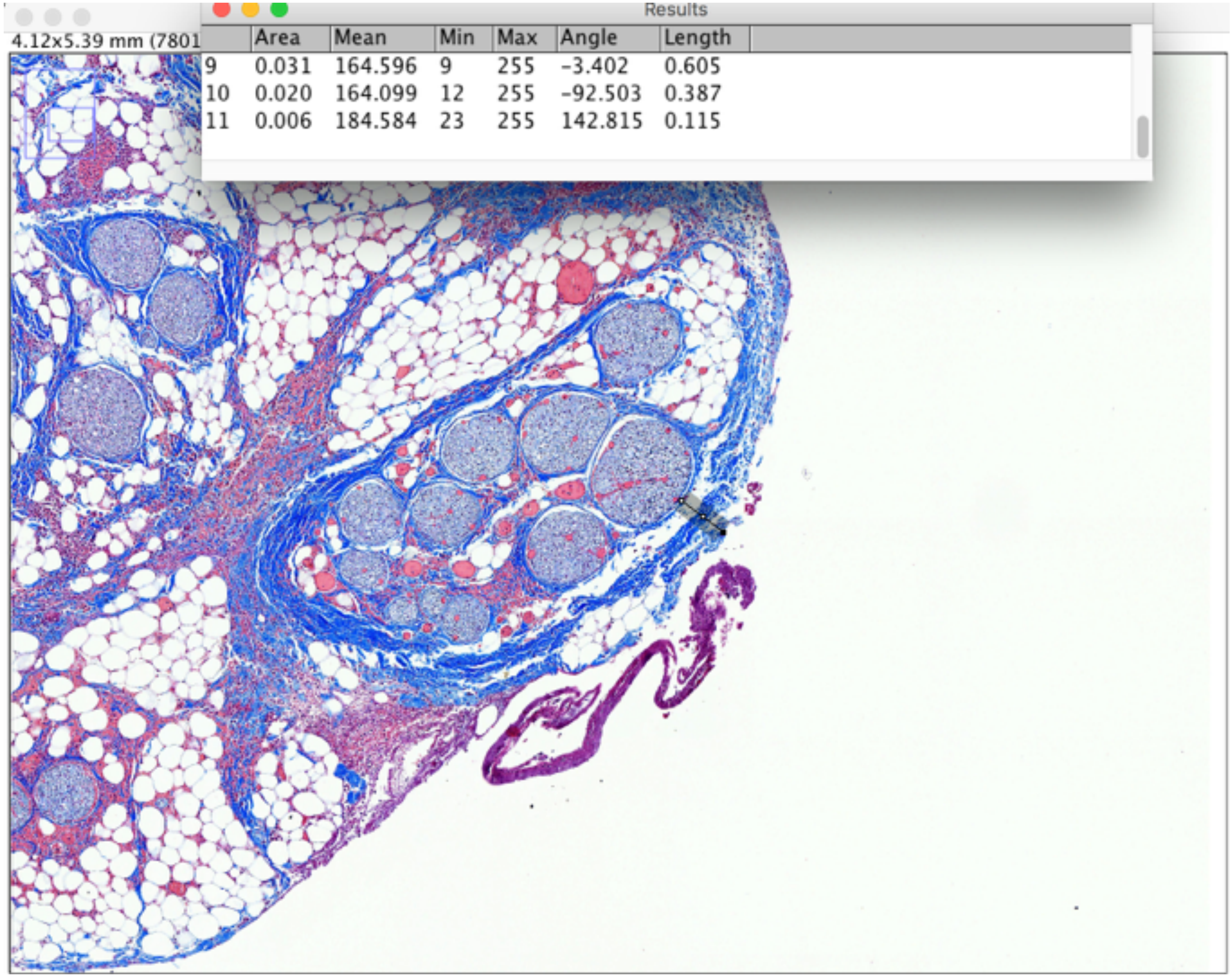


P786_LV1b_LivaNova

Fascicle depth

Subject 3_P824

### Nodose

Results

| Area | Mean | Min | Max | Angle | Length |
|---|---|---|---|---|---|
| 0.009 | 184.742 | 8 | 255 | -86.260 | 0.243 |
| 0.009 | 185.522 | 8 | 255 | -90.000 | 0.244 |
| 0.210 | 173.693 | 3 | 255 | -66.517 | 5.692 |

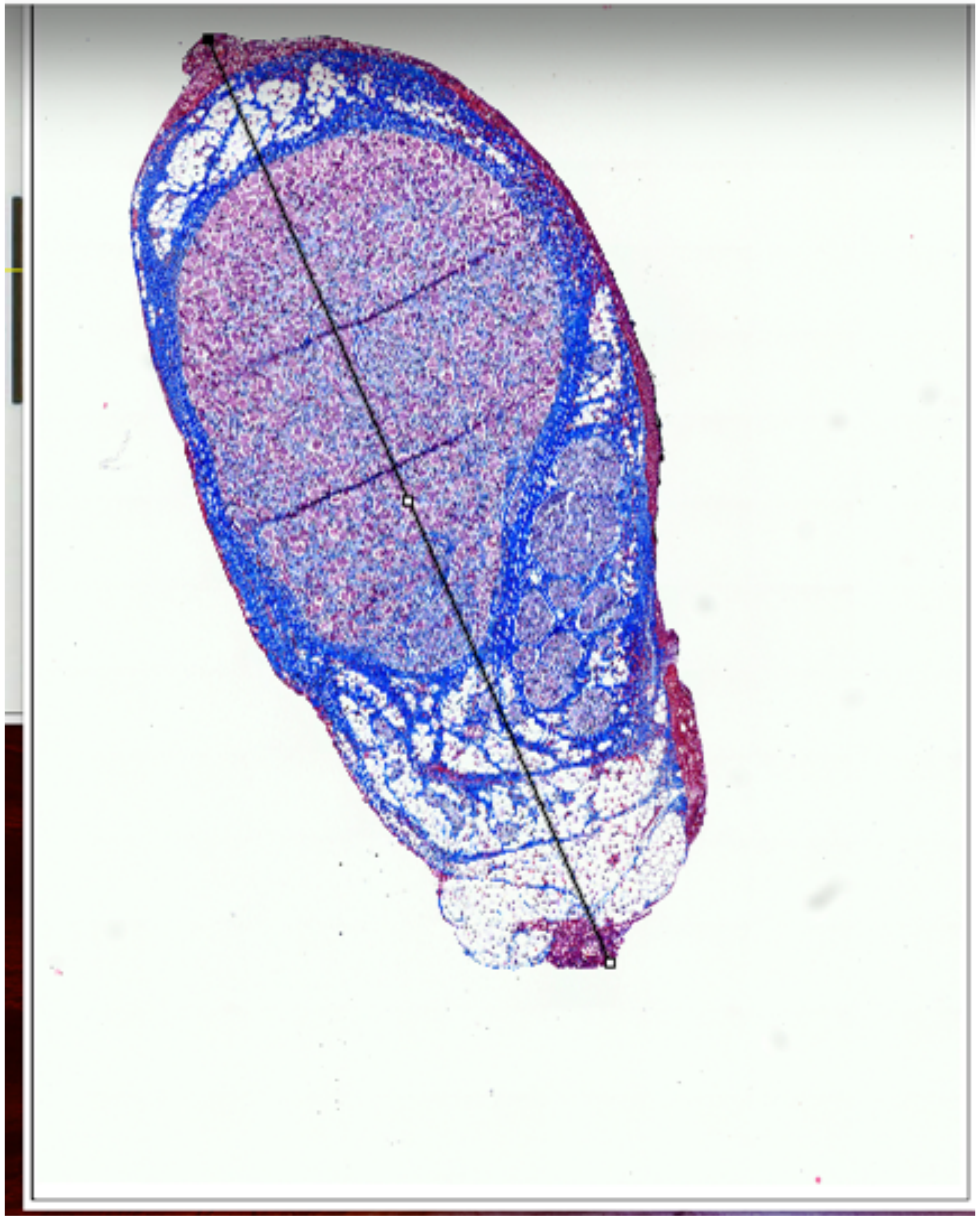

P824_nodose_a_2, nodose

Results

| | Area | Mean | Min | Max | Angle | Length |
|---|---|---|---|---|---|---|
| 1 | 0.052 | 139.610 | 5 | 255 | 21.444 | 2.010 |

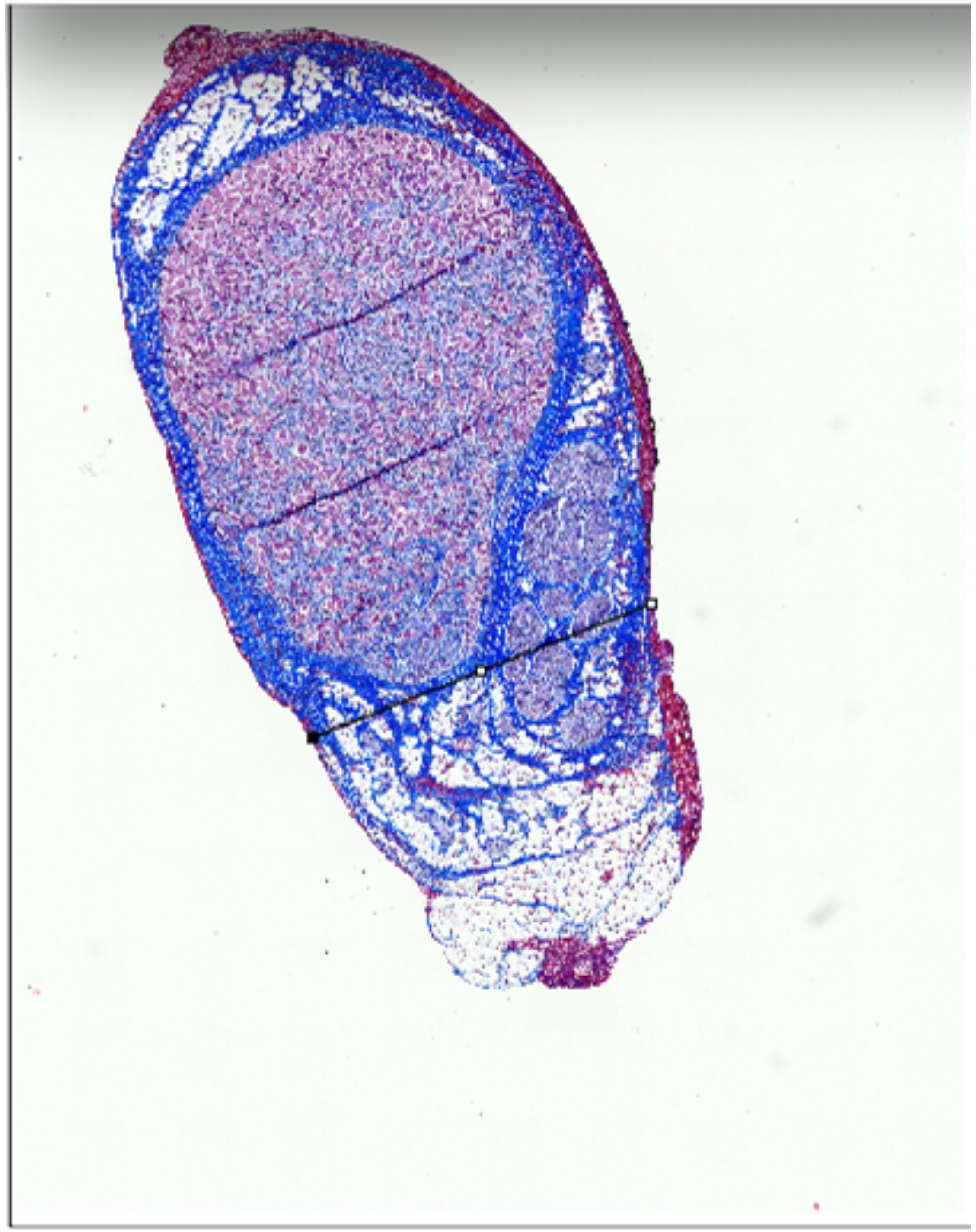

Widest and narrowest diameter

### Nodose

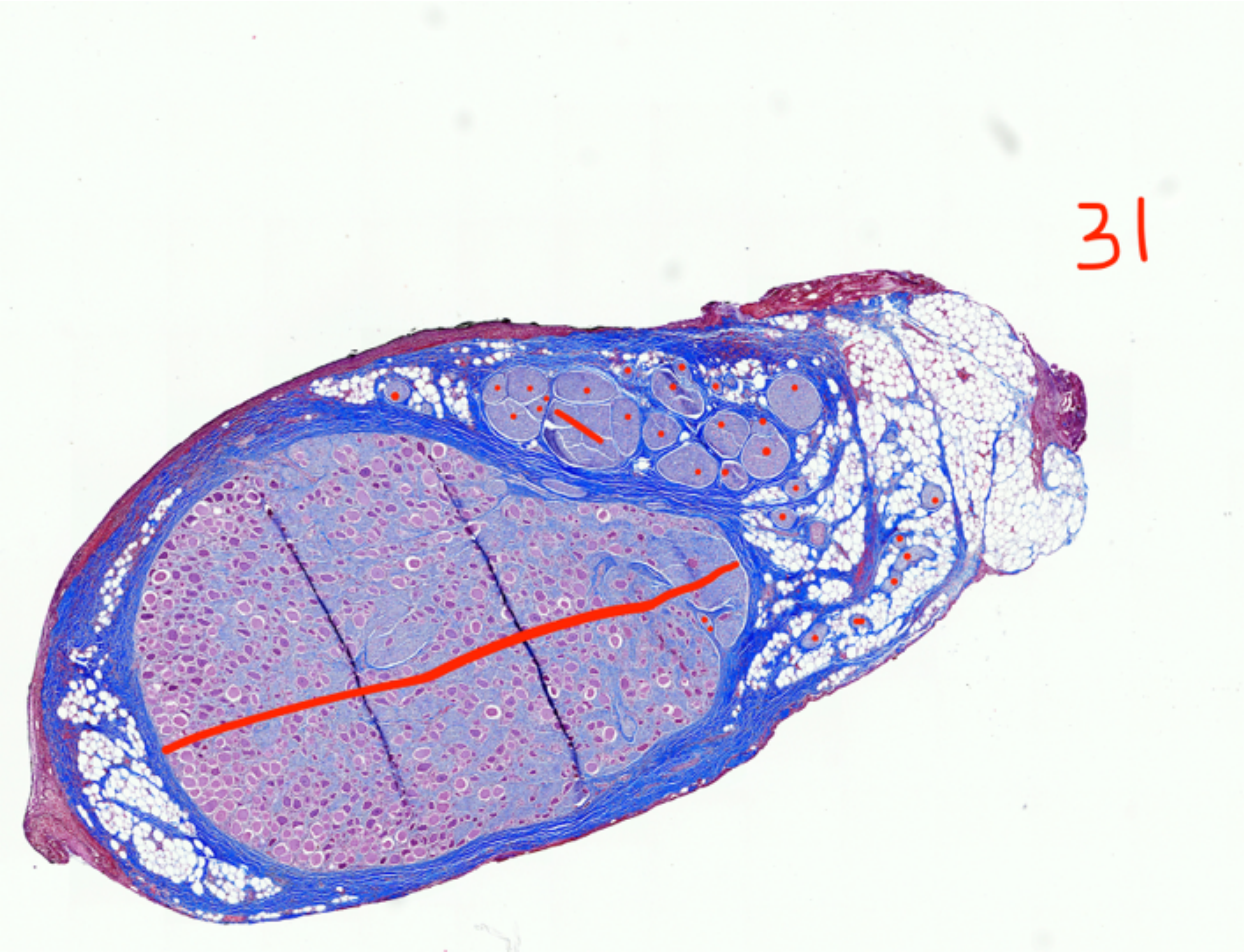


P824_nodose_a_2, nodose

Fascicle Count, 31

### Nodose

Results

| rea | Mean | Min | Max | Angle | Length |
|---|---|---|---|---|---|
| .210 | 173.693 | 3 | 255 | -66.517 | 5.692 |
| .097 | 163.632 | 5 | 255 | 25.320 | 2.607 |
| .112 | 167.214 | 6 | 255 | -74.766 | 3.014 |

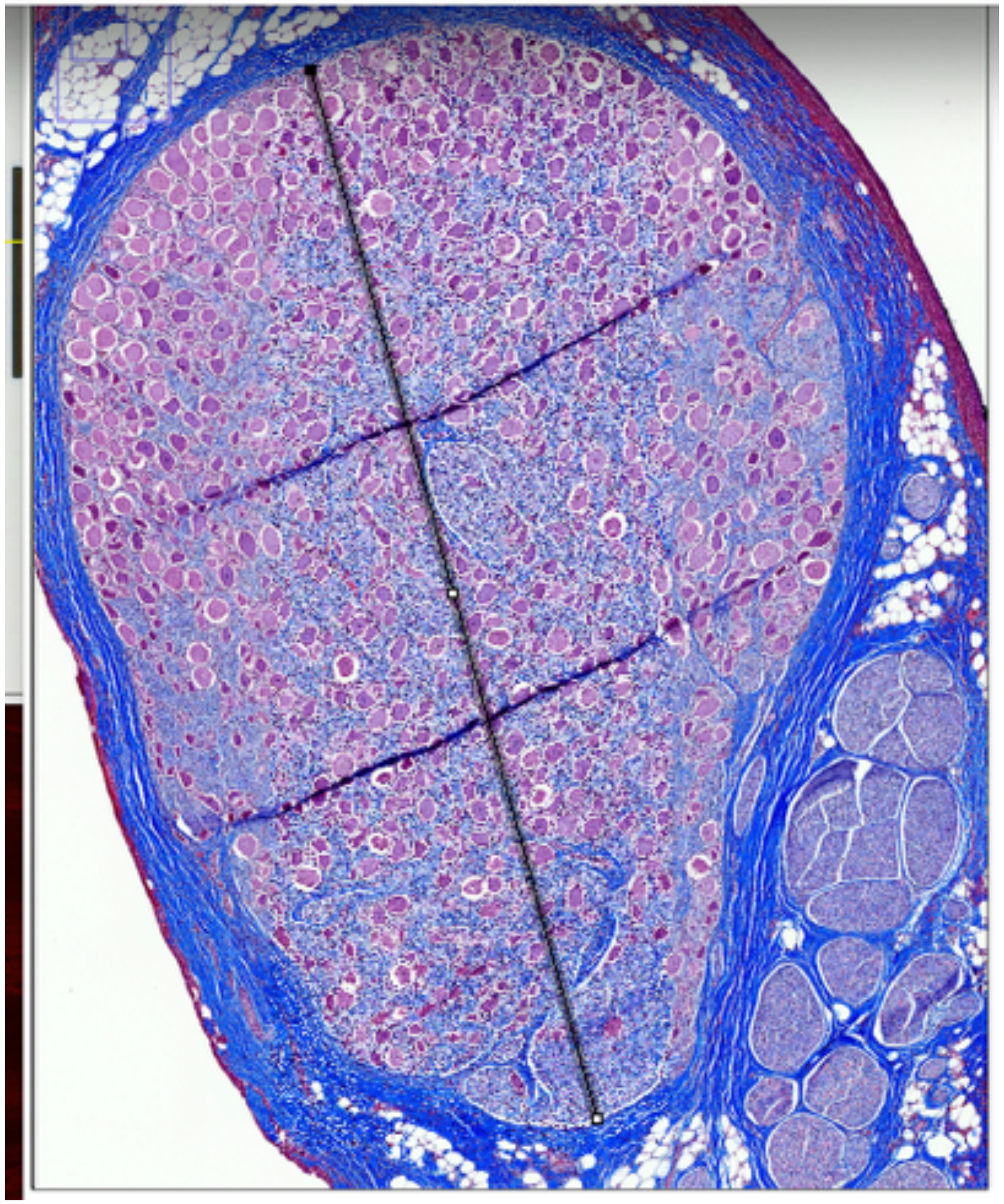

Results

| | Area | Mean | Min | Max | Angle | Length |
|---|---|---|---|---|---|---|
| 1 | 0.052 | 139.610 | 5 | 255 | 21.444 | 2.010 |
| 2 | 0.033 | 168.442 | 11 | 255 | 24.969 | 1.221 |
| 3 | 0.017 | 164.908 | 12 | 255 | 32.530 | 0.636 |

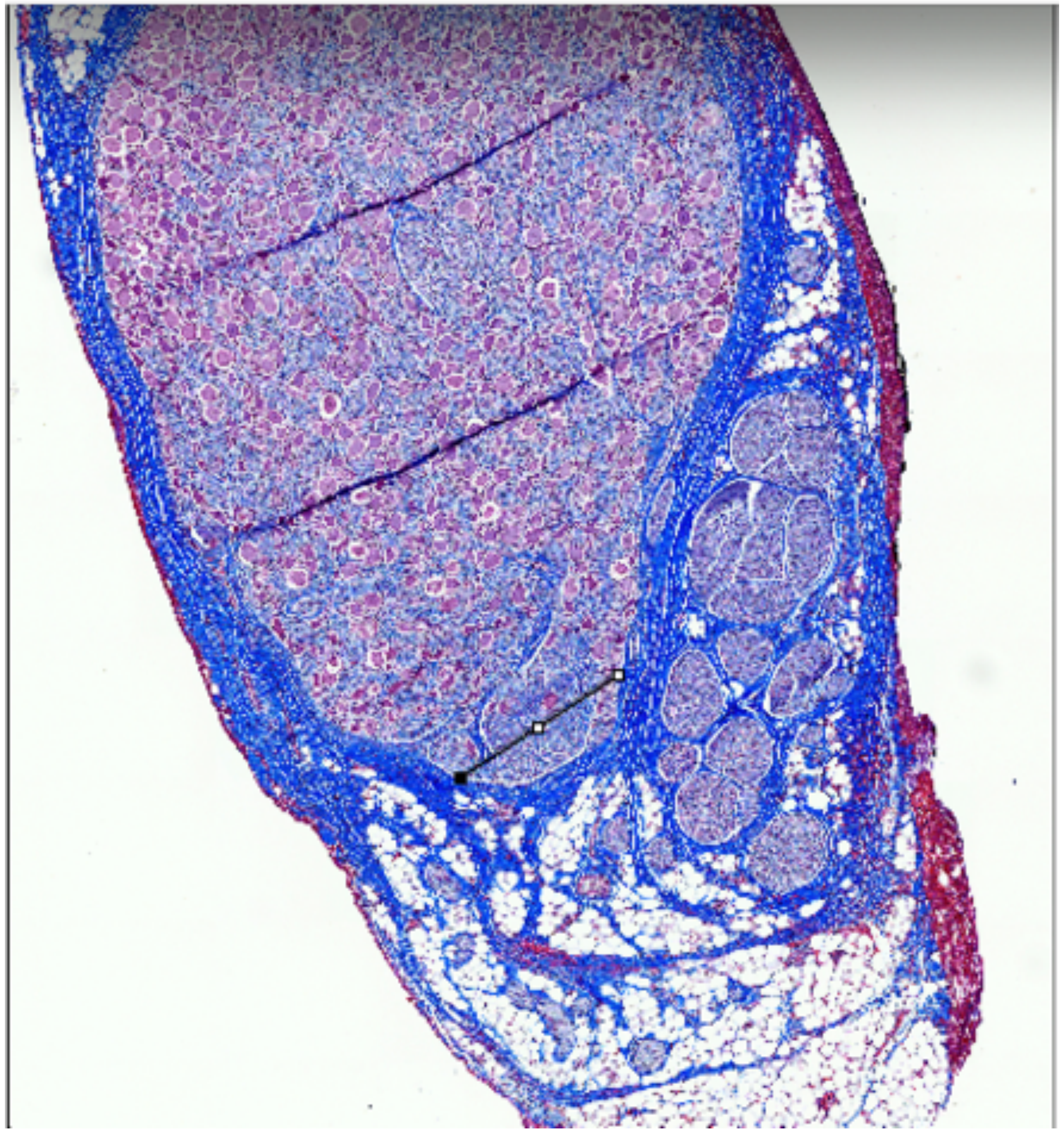

### P824_nodose_a_2, nodose

Largest fascicle_widest and narrowest diameters

Subject 4_P833

### Nodose

Results

| | Area | Mean | Min | Max | Angle | Length |
|---|---|---|---|---|---|---|
| 20 | 0.105 | 194.465 | 4 | 255 | -99.752 | 2.880 |
| 21 | 0.087 | 192.916 | 5 | 255 | -4.661 | 2.339 |
| 22 | 0.176 | 194.268 | 9 | 255 | 87.143 | 4.745 |

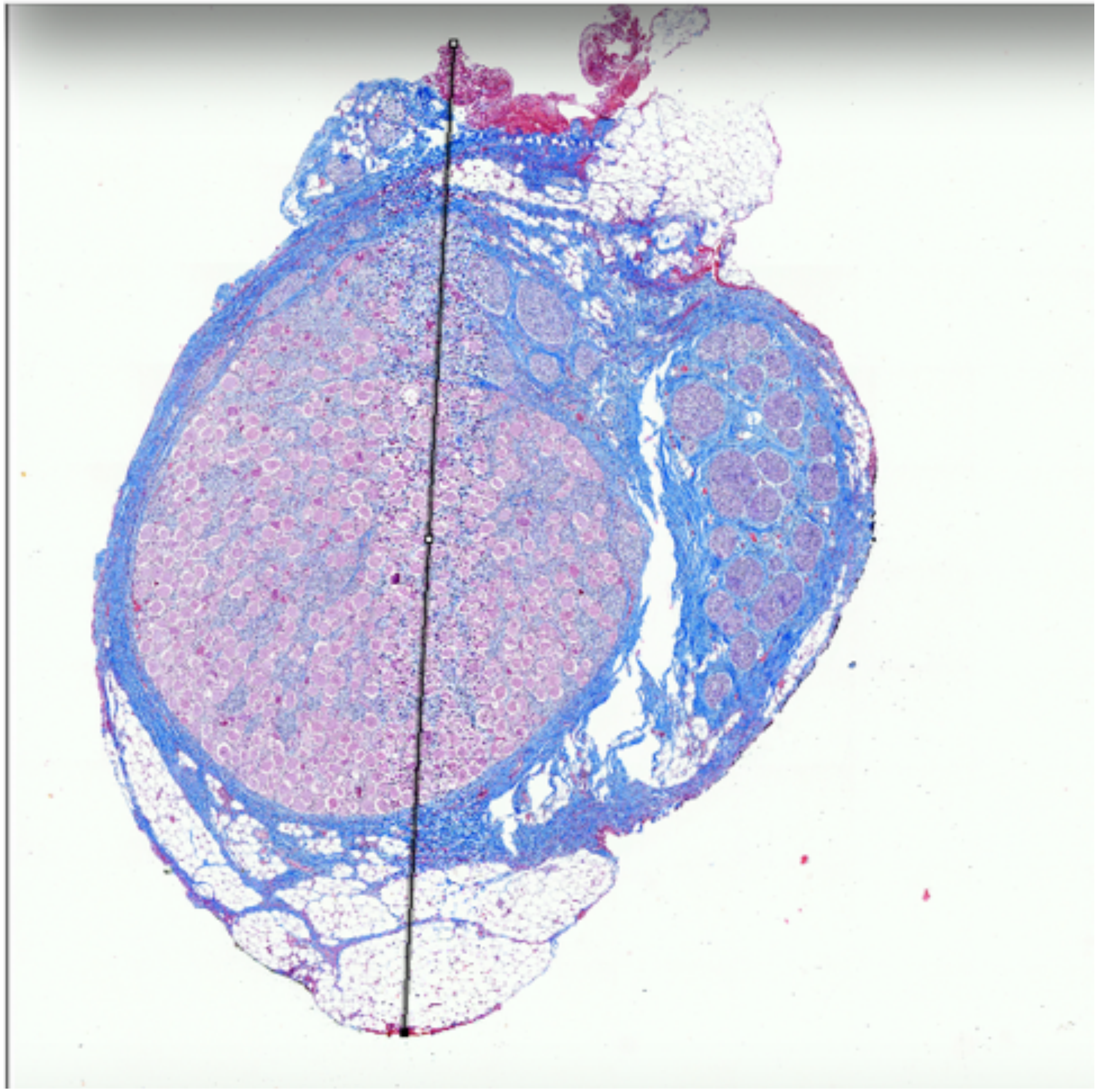

Results

| | Area | Mean | Min | Max | Angle | Length |
|---|---|---|---|---|---|---|
| 3 | 0.017 | 164.908 | 12 | 255 | 32.530 | 0.636 |
| 4 | 0.060 | 179.555 | 12 | 255 | -4.154 | 2.274 |
| 5 | 0.061 | 176.374 | 11 | 255 | 175.236 | 2.289 |

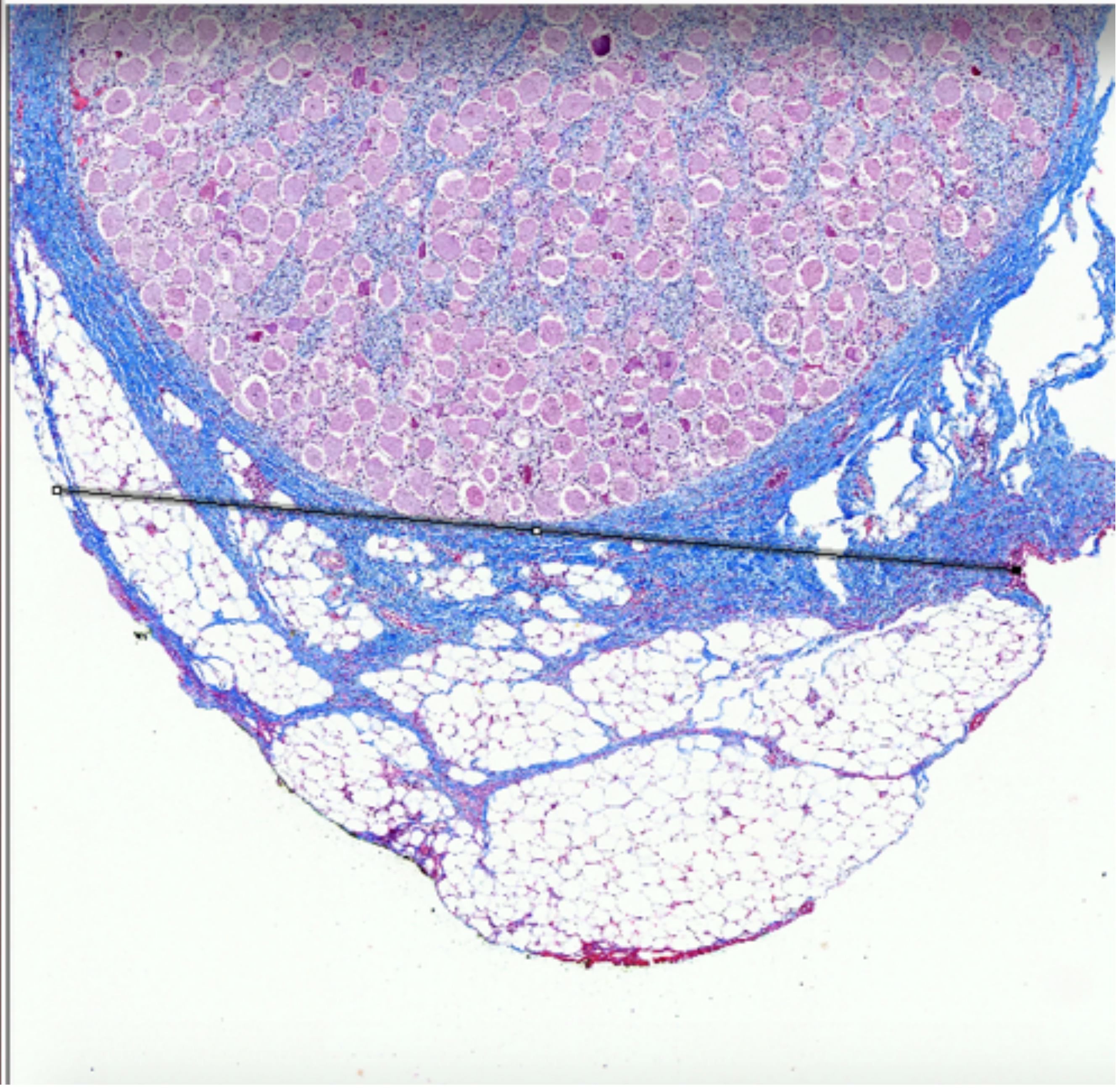

P833_RV1_18, nodose

Widest and narrowest diameter

Nodose

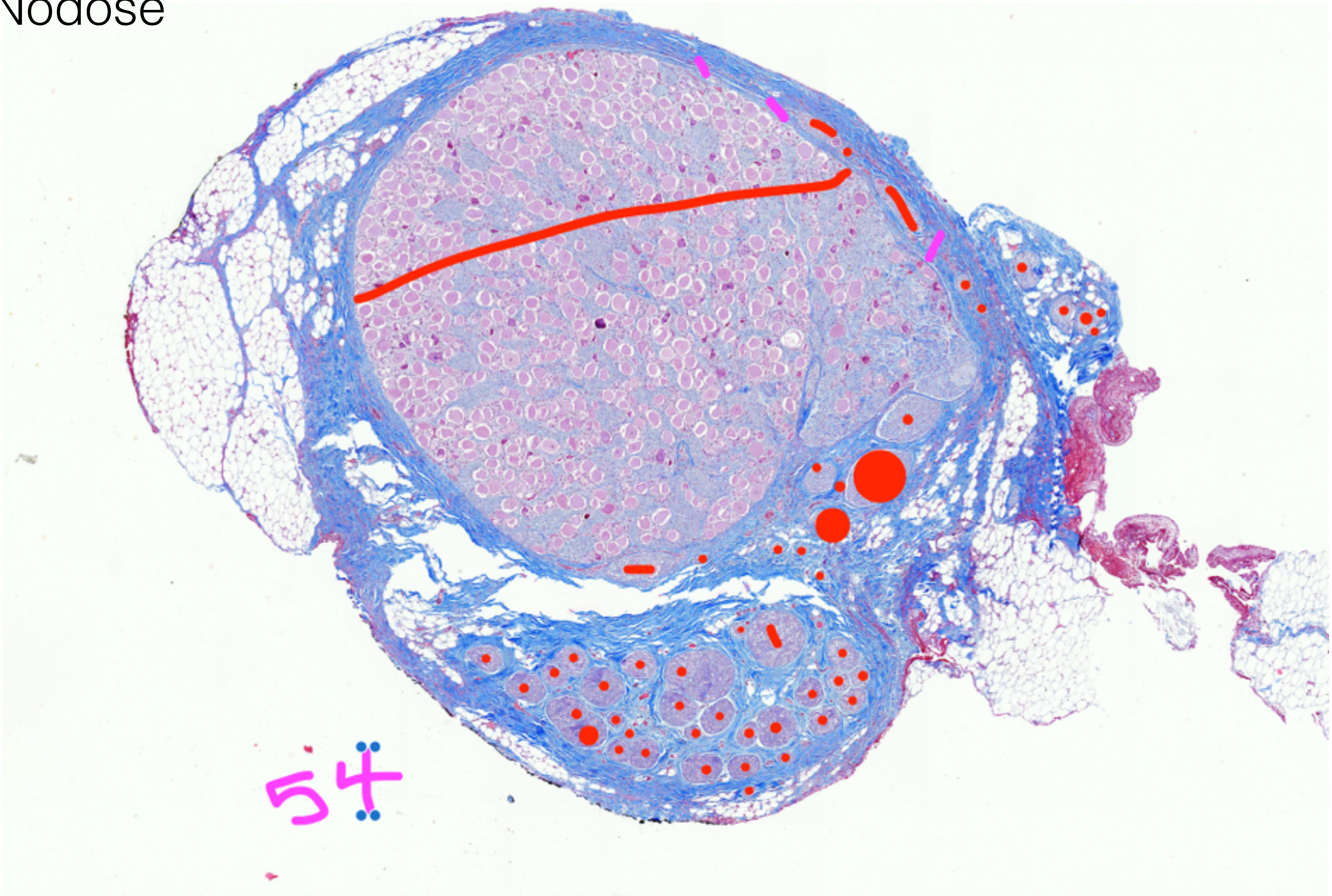


P833_RV1_18, nodose, fascicle count, 54

Fascicle Count, 54

### Nodose

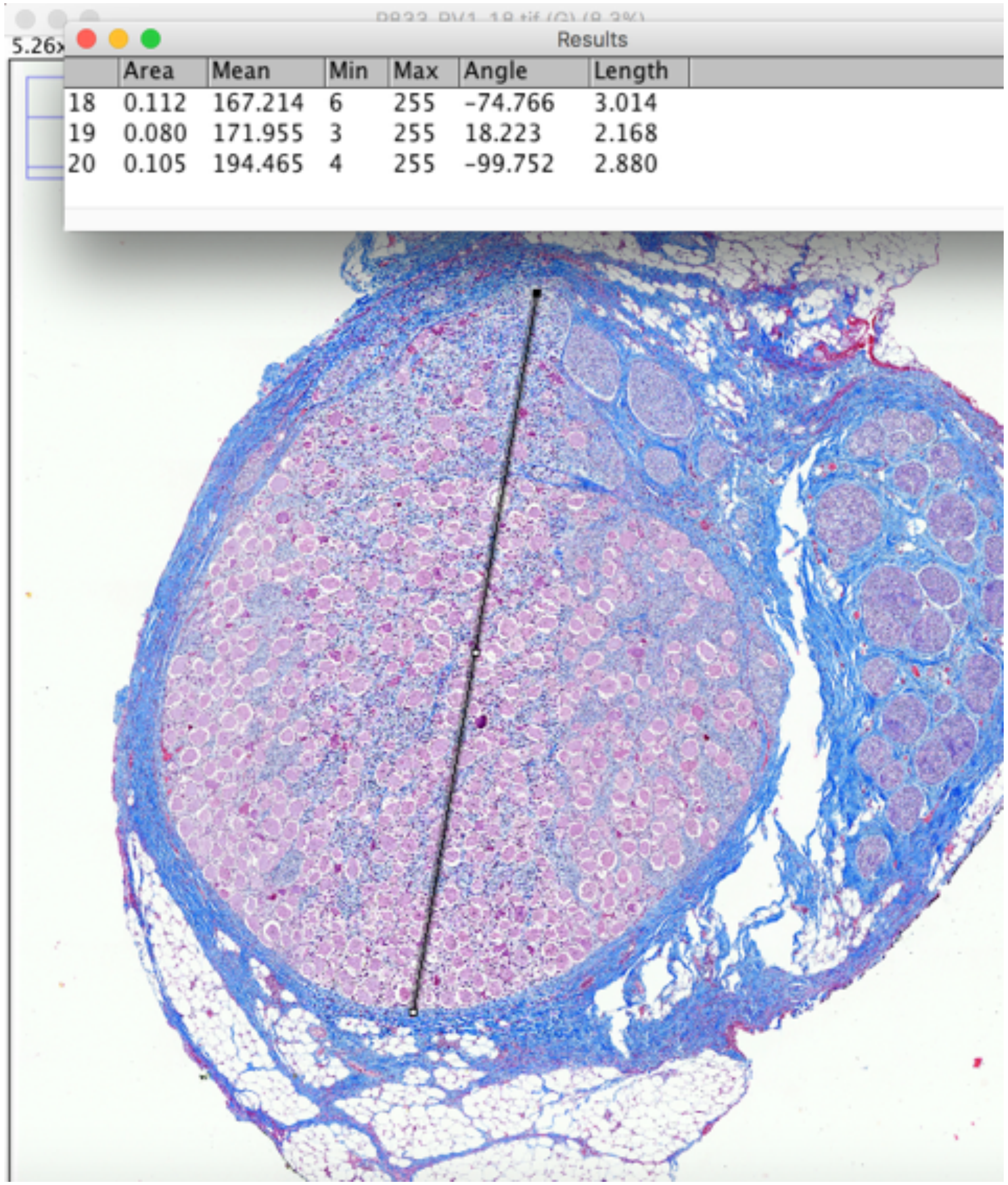


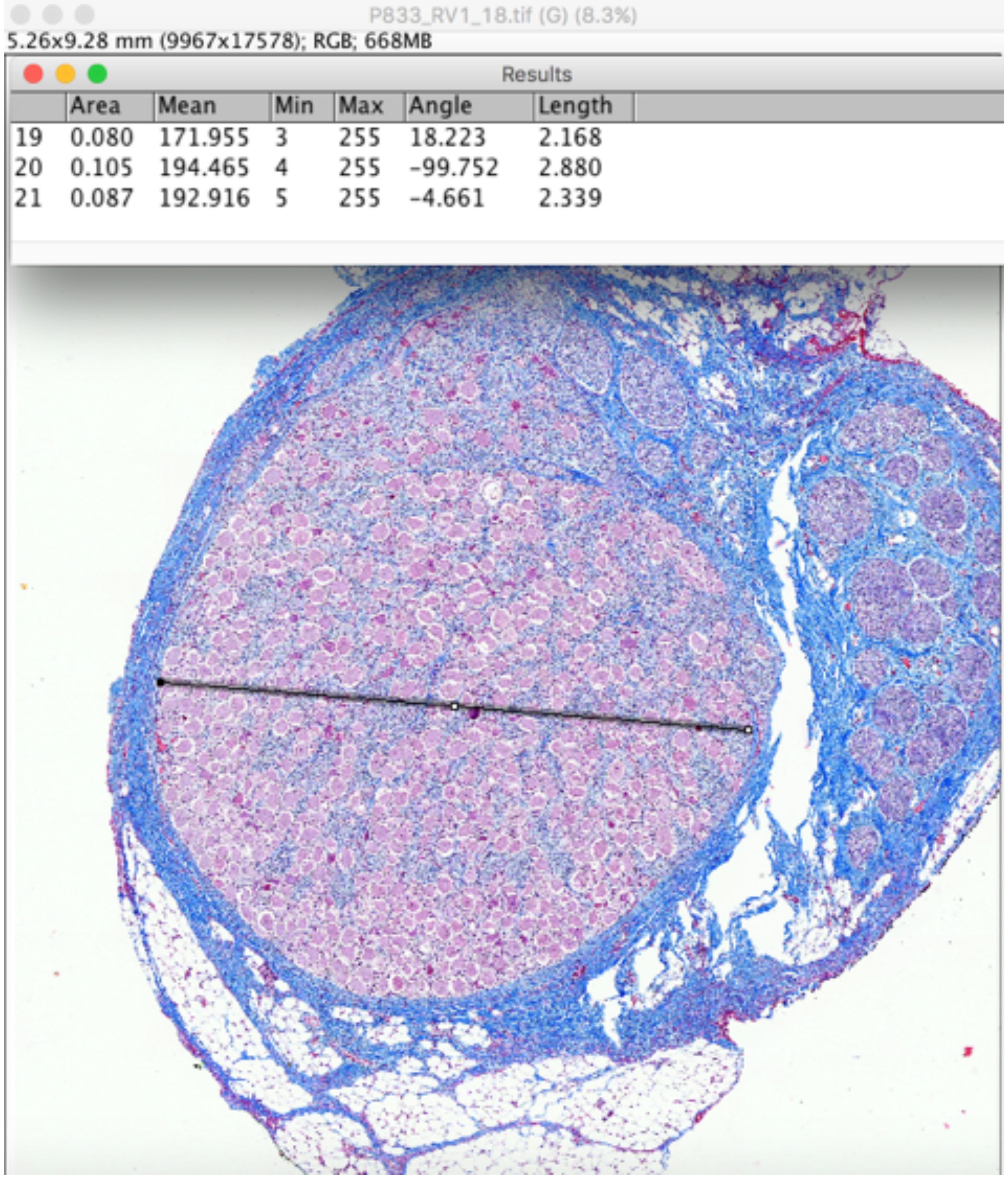


P833_RV1_18, nodose

Largest fascicle_widest and narrowest diameters

### Mid-VN

Results

| | Area | Mean | Min | Max | Angle | Length |
|---|---|---|---|---|---|---|
| 22 | 0.176 | 194.268 | 9 | 255 | 87.143 | 4.745 |
| 23 | 0.142 | 188.352 | 7 | 255 | 9.609 | 3.796 |
| 24 | 0.110 | 181.322 | 5 | 255 | -113.790 | 2.984 |

Results

| | Area | Mean | Min | Max | Angle | Length |
|---|---|---|---|---|---|---|
| 1 | 0.050 | 179.520 | 13 | 255 | -57.894 | 1.907 |

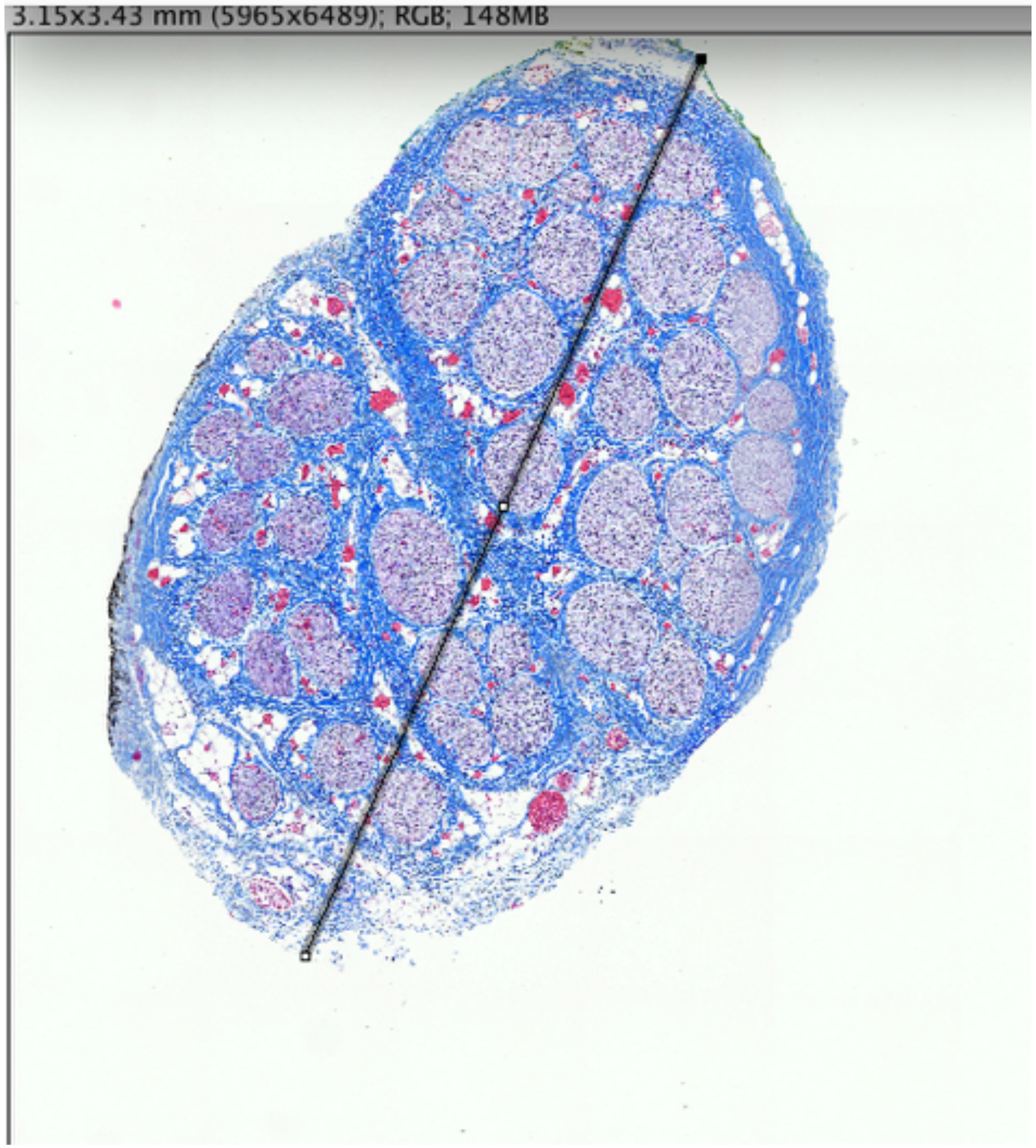


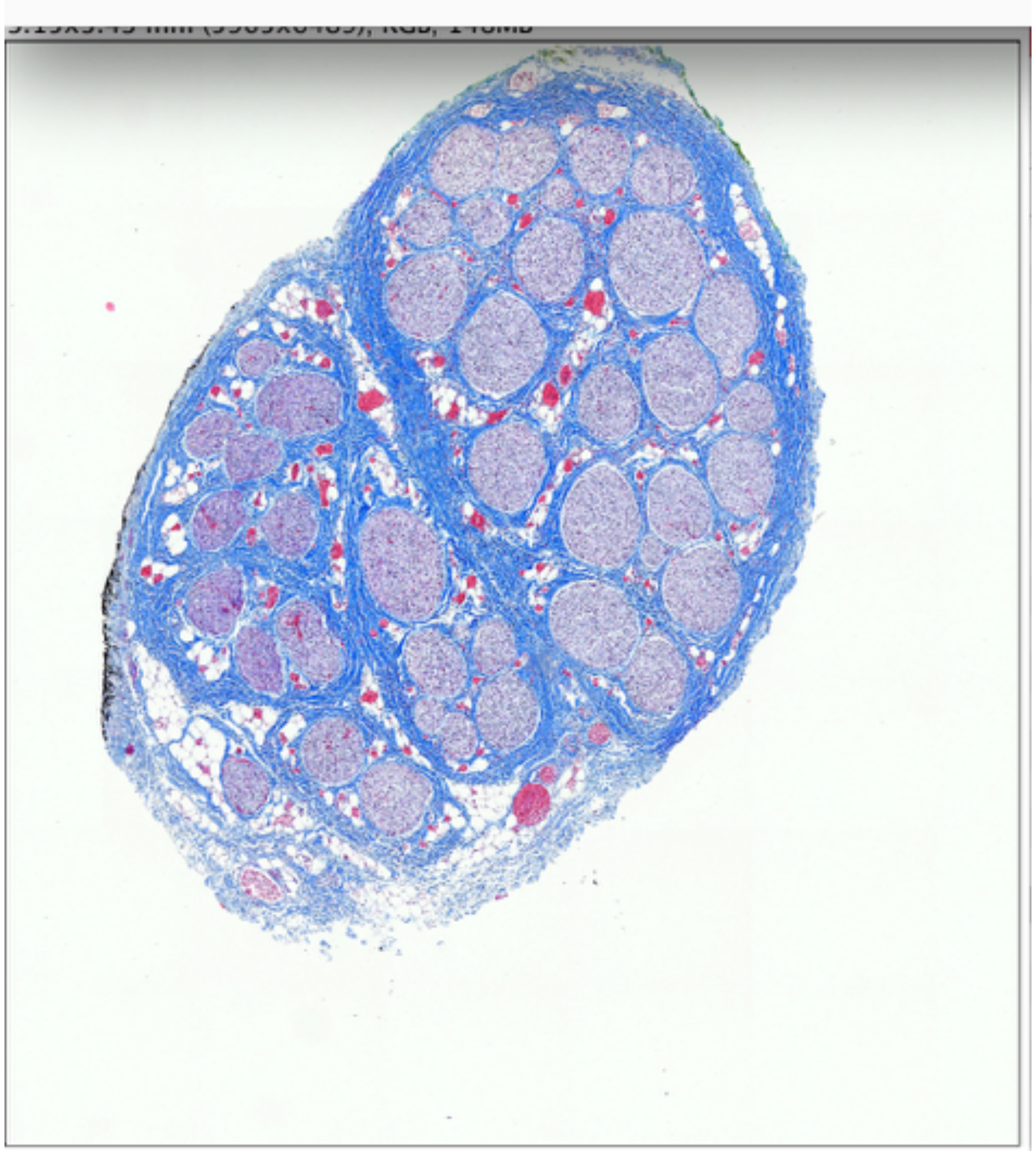

P833_RV2b_2

Widest and narrowest diameters

### Mid-VN

P833_RV2b_2.tif (G) (33.3%)

3.15x3.43 mm (5965x6489); RGB; 148MB

Results

| | Area | Mean | Min | Max | Angle | Length |
|---|---|---|---|---|---|---|
| 11 | 0.006 | 184.584 | 23 | 255 | 142.815 | 0.115 |
| 12 | 0.004 | 174.451 | 1 | 255 | -24.349 | 0.079 |
| 13 | 0.004 | 170.578 | 1 | 255 | -29.168 | 0.078 |

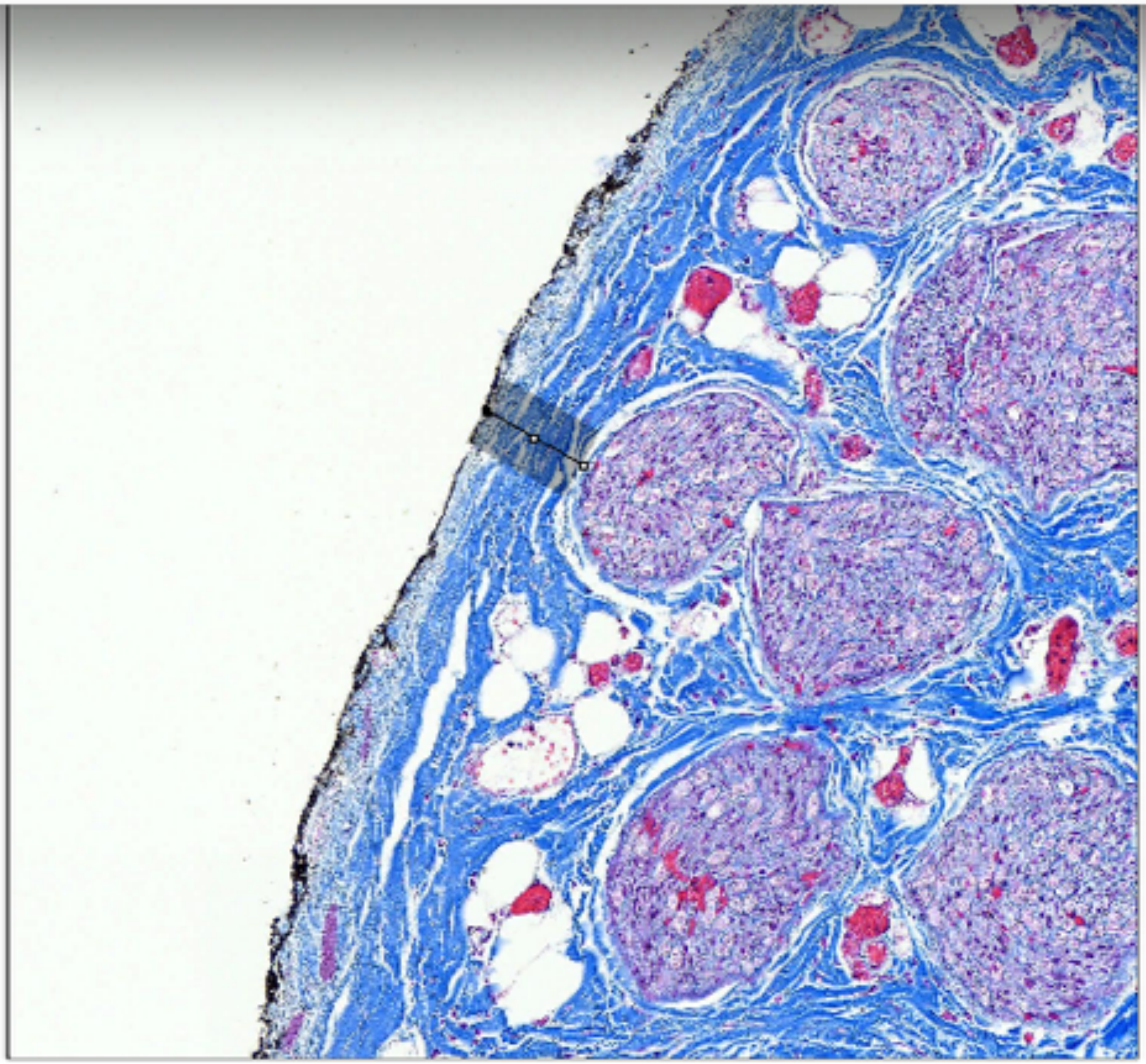

P833_RV2b_2

Fascicle Depth

Mid-VN

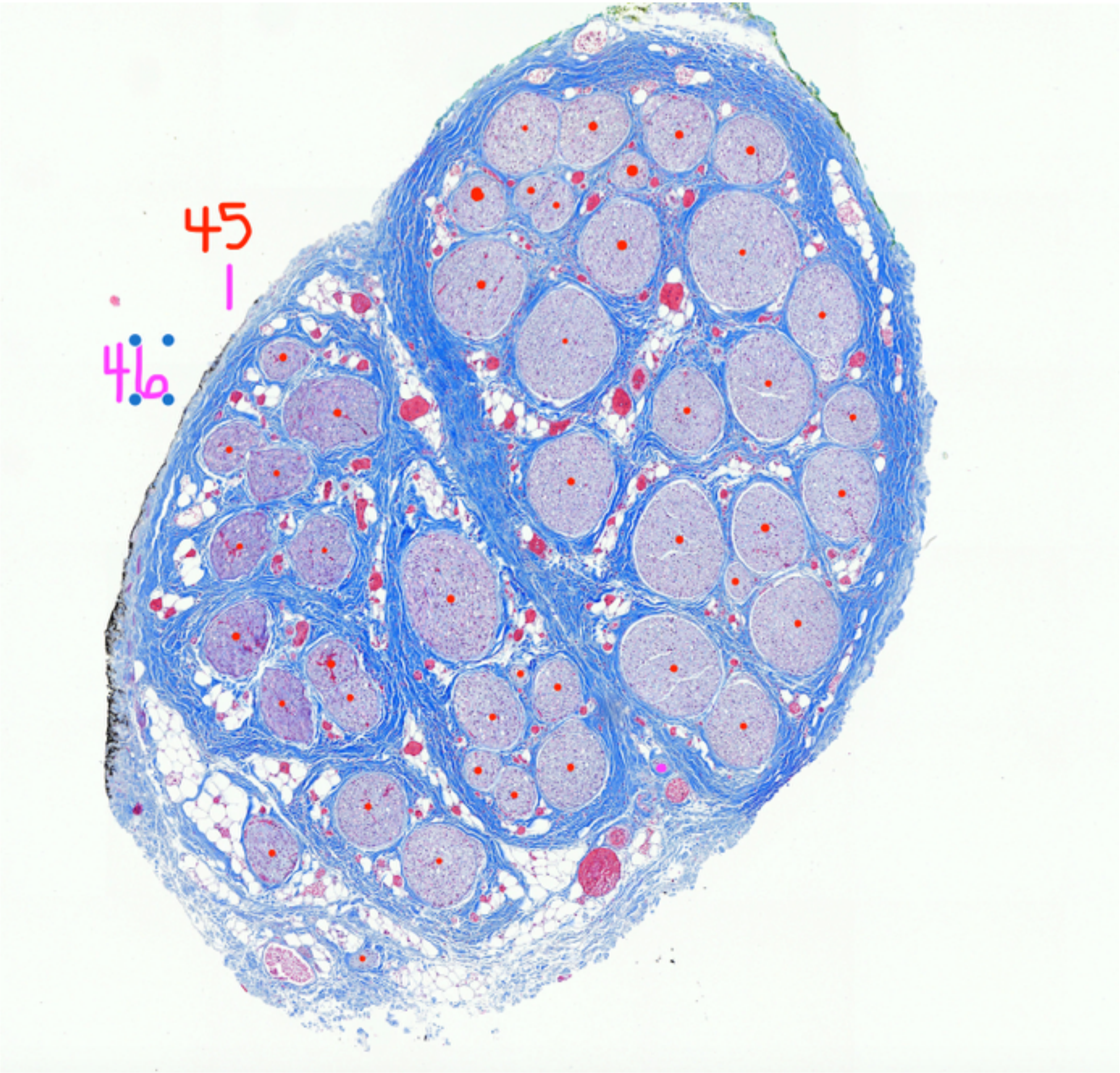


P833_RV2b_2

Fascicle count, 46

### Mid-VN

P833_RV2b_2.tif (G) (12.5%)

3.15x3.43 mm (5965x6489); RGB; 148MB

Results

| Area | Mean | Min | Max | Angle | Length |
|---|---|---|---|---|---|
| 0.110 | 181.322 | 5 | 255 | -113.790 | 2.984 |
| 0.072 | 179.130 | 4 | 255 | -49.179 | 1.968 |
| 0.013 | 190.754 | 9 | 255 | -69.085 | 0.355 |

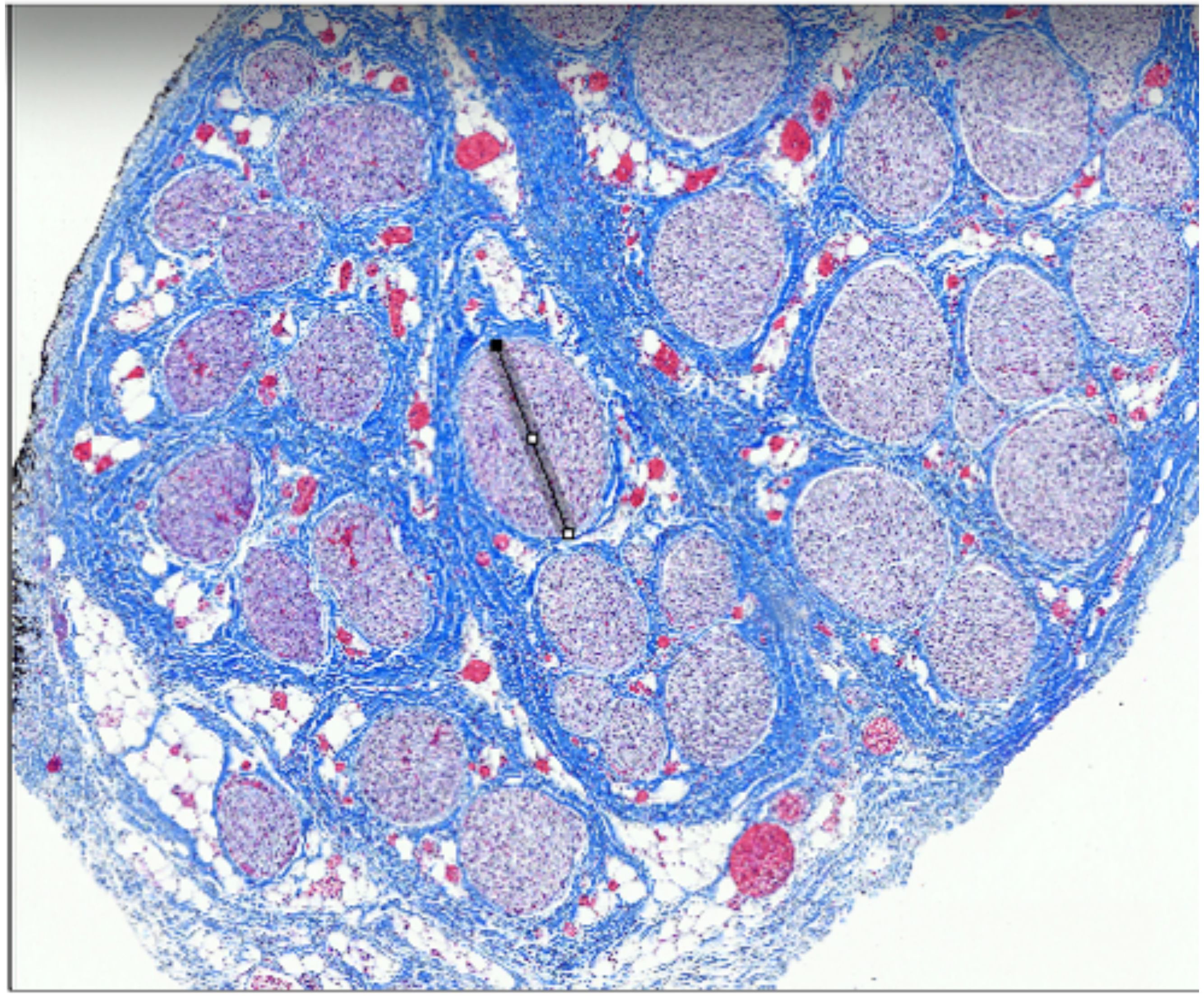

Results

| Area | Mean | Min | Max | Angle | Length |
|---|---|---|---|---|---|
| 0.072 | 179.130 | 4 | 255 | -49.179 | 1.968 |
| 0.013 | 190.754 | 9 | 255 | -69.085 | 0.355 |
| 0.009 | 190.395 | 7 | 255 | 27.229 | 0.245 |

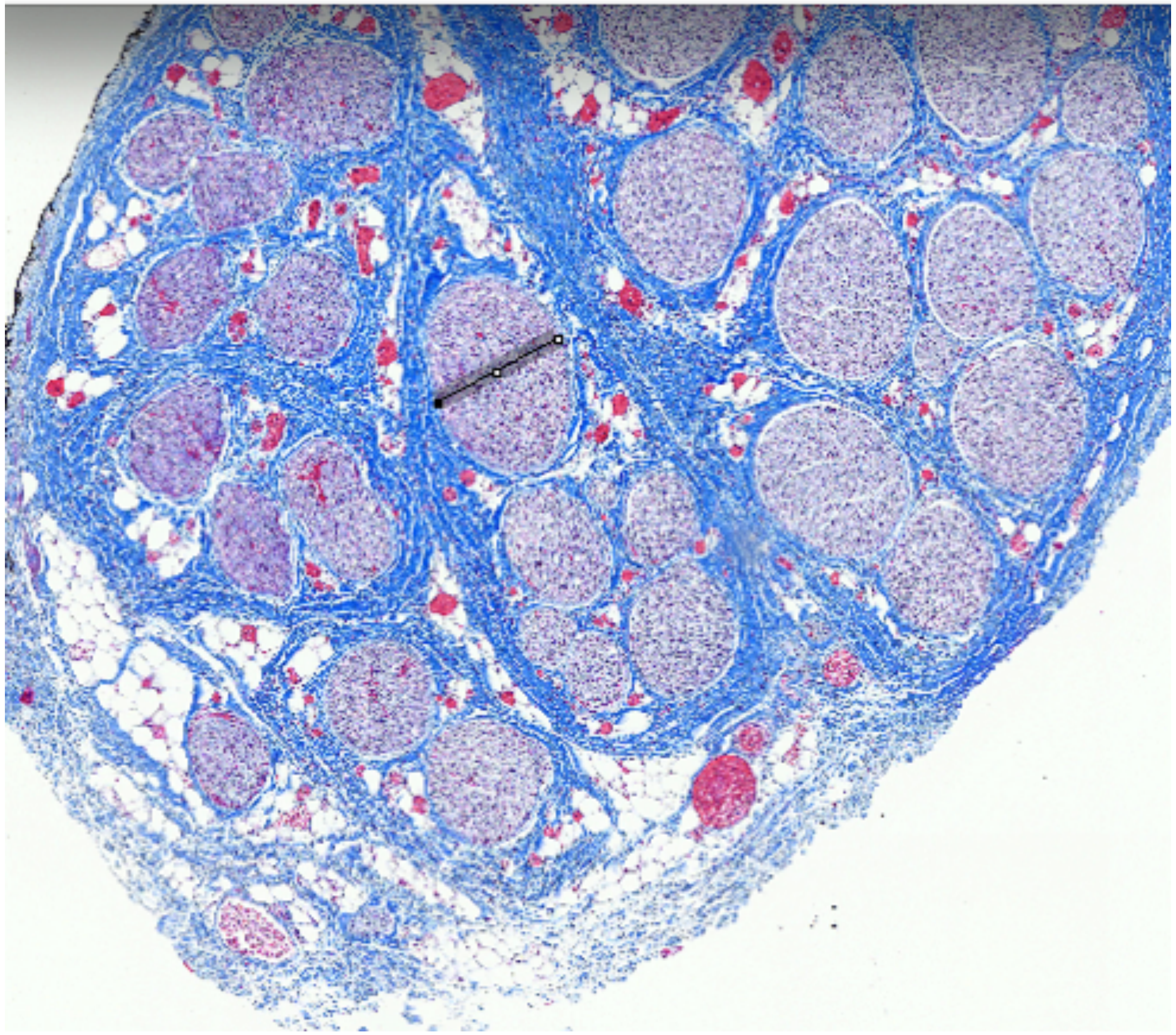

P833_RV2b_2

Largest Fascicle_Widest and narrowest diameters

RL

Results

| | Area | Mean | Min | Max | Angle | Length |
|---|---|---|---|---|---|---|
| 26 | 0.013 | 190.754 | 9 | 255 | -69.085 | 0.355 |
| 27 | 0.009 | 190.395 | 7 | 255 | 27.229 | 0.245 |
| 28 | 0.080 | 165.985 | 7 | 255 | -7.842 | 2.136 |

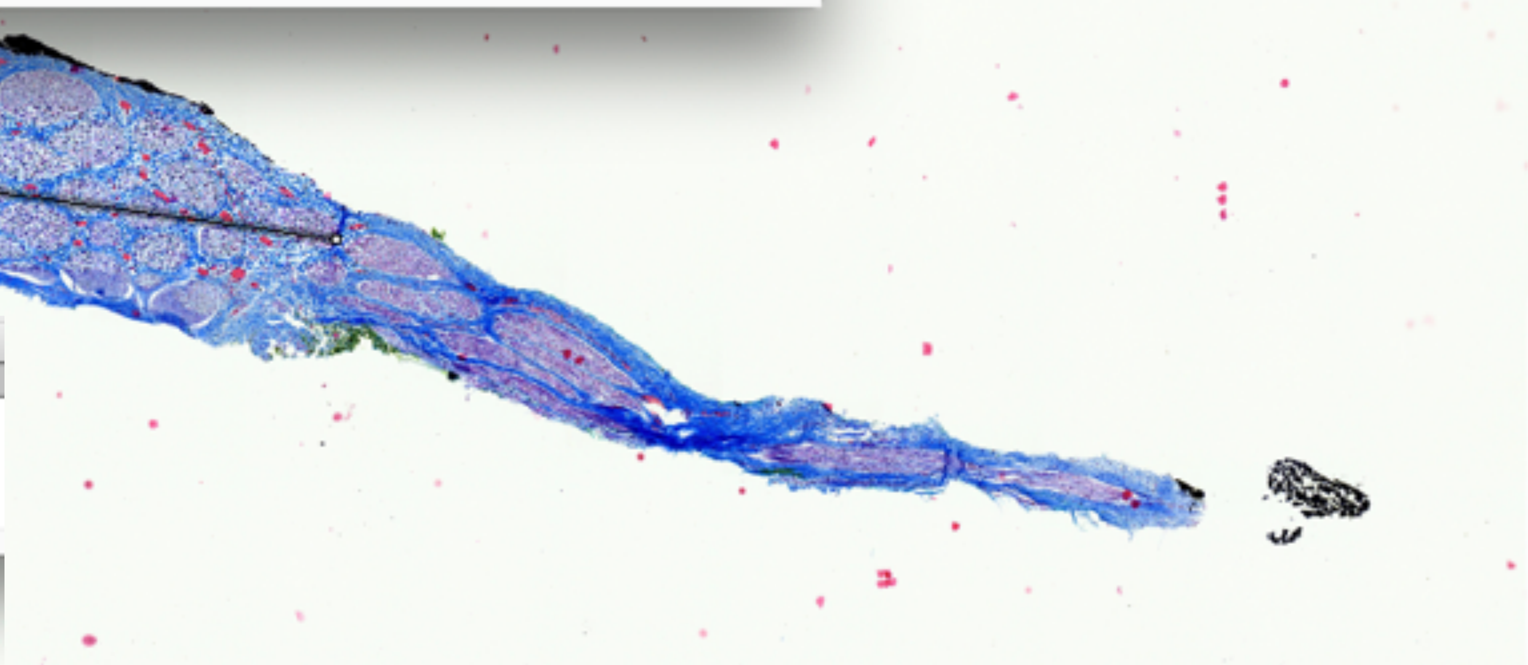

Results

| | Area | Mean | Min | Max | Angle | Length |
|---|---|---|---|---|---|---|
| 1 | 0.050 | 179.520 | 13 | 255 | -57.894 | 1.907 |
| 2 | 0.010 | 170.234 | 18 | 255 | -2.770 | 0.393 |

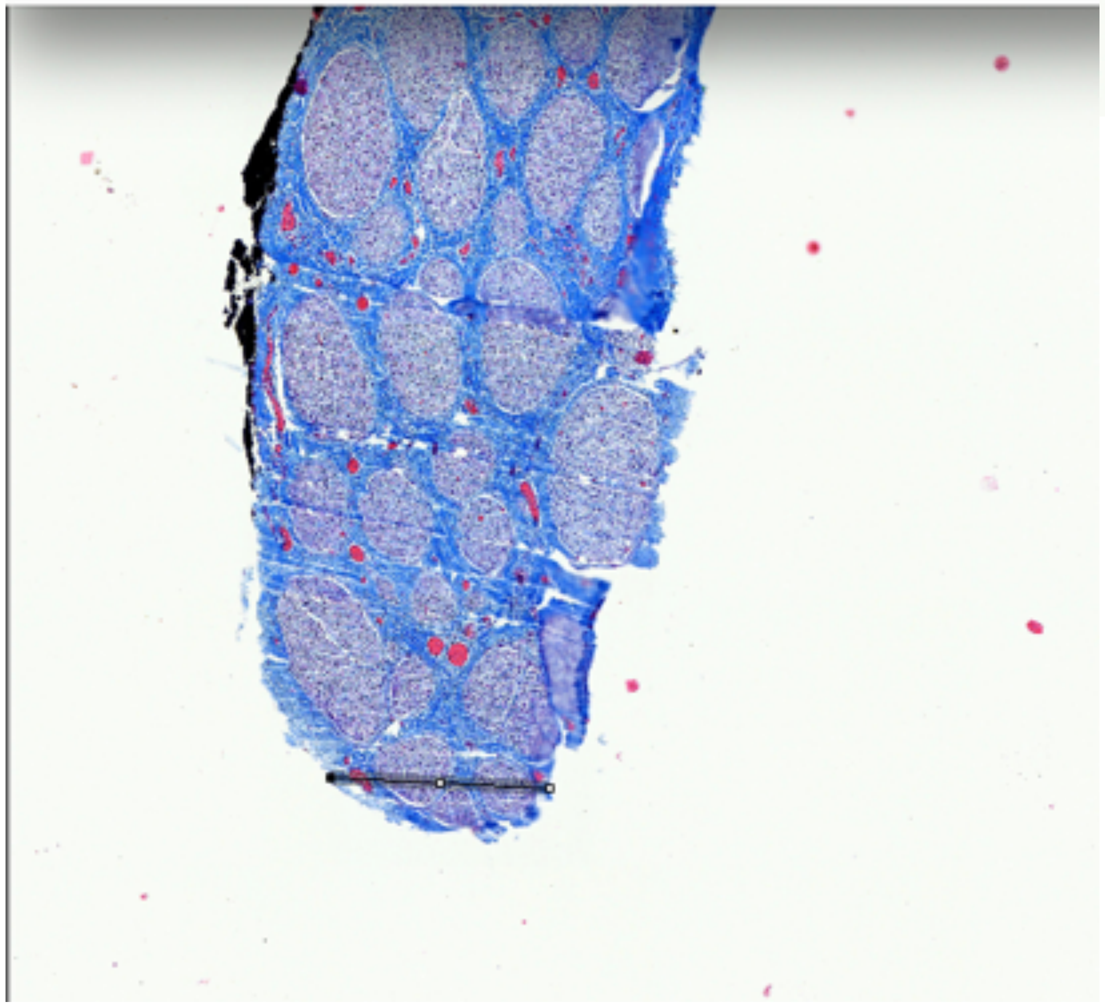

P833_RV3_5

Widest and narrowest diameters

RL

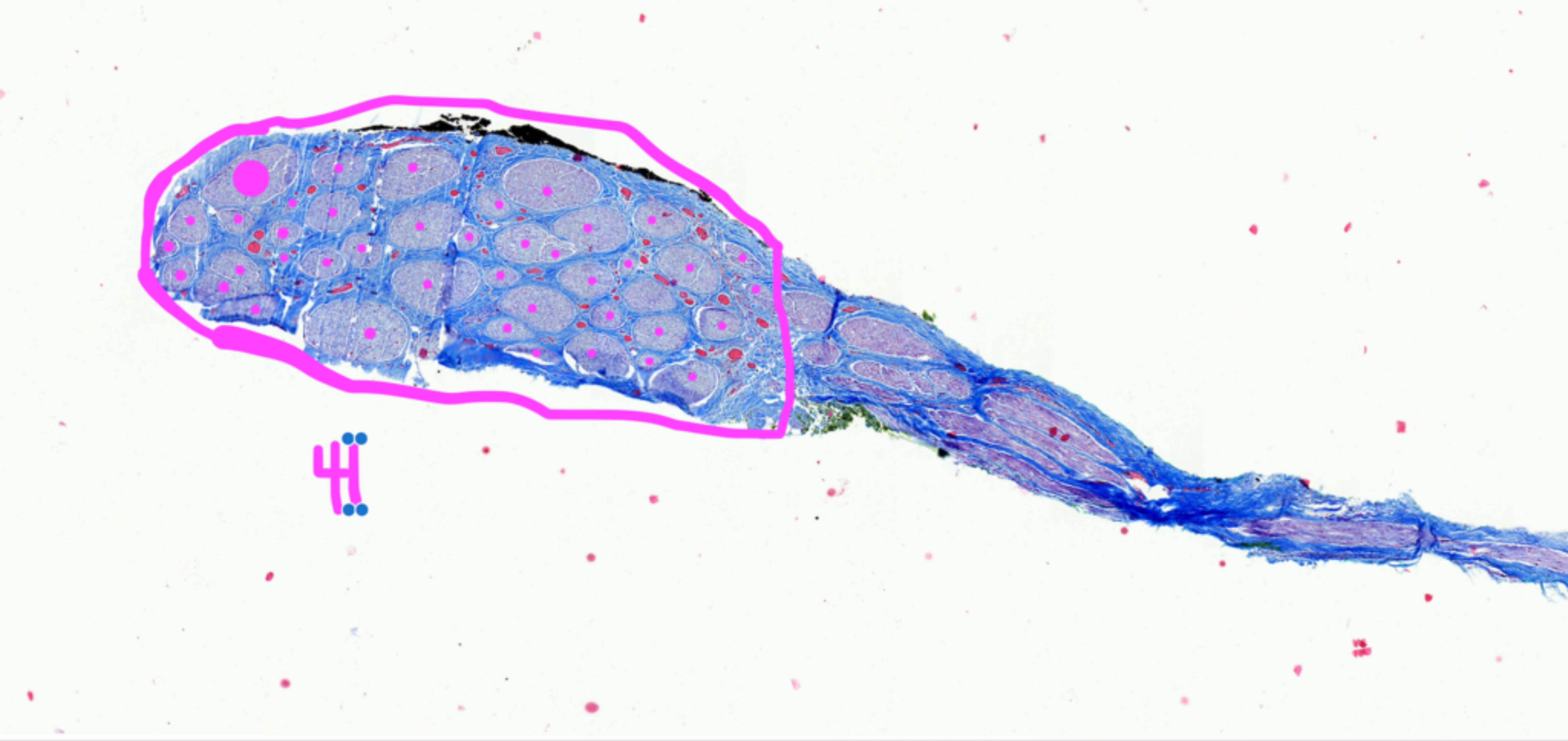


P833_RV3_5

Fascicle count,41

RL

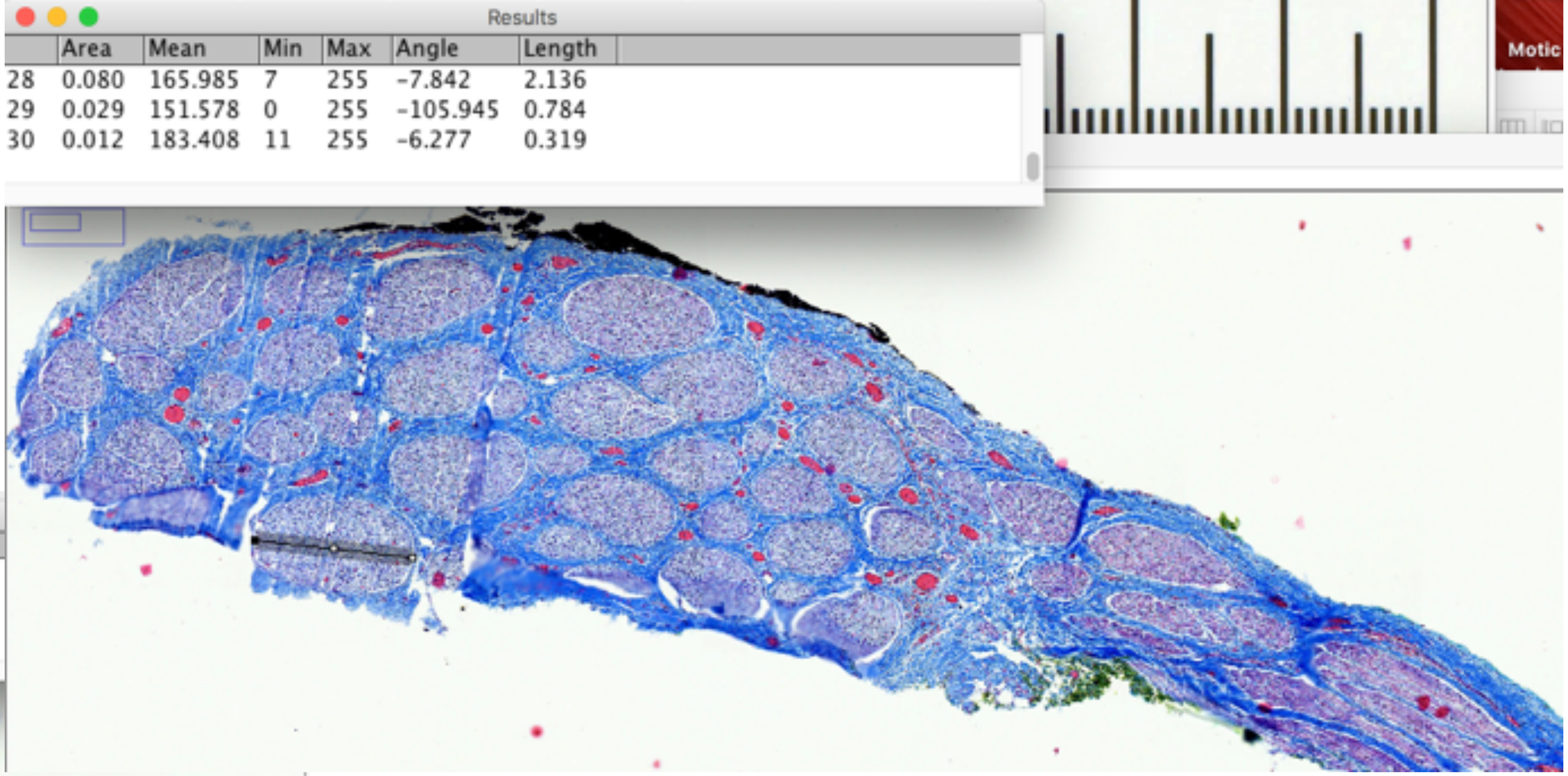


| | Area | Mean | Min | Max | Angle | Length |
|---|---|---|---|---|---|---|
| 28 | 0.080 | 165.985 | 7 | 255 | -7.842 | 2.136 |
| 29 | 0.029 | 151.578 | 0 | 255 | -105.945 | 0.784 |
| 30 | 0.012 | 183.408 | 11 | 255 | -6.277 | 0.319 |

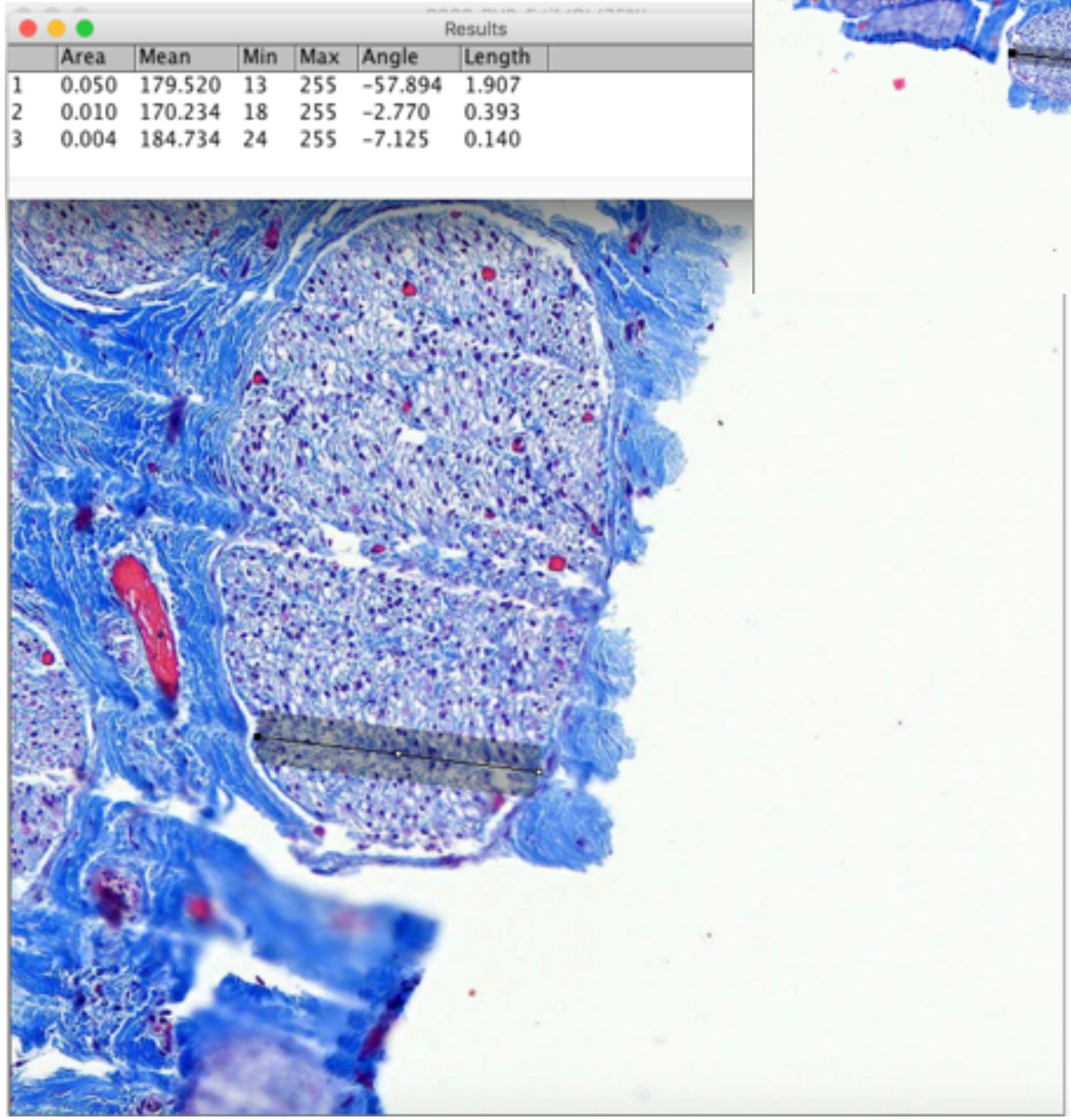


| | Area | Mean | Min | Max | Angle | Length |
|---|---|---|---|---|---|---|
| 1 | 0.050 | 179.520 | 13 | 255 | -57.894 | 1.907 |
| 2 | 0.010 | 170.234 | 18 | 255 | -2.770 | 0.393 |
| 3 | 0.004 | 184.734 | 24 | 255 | -7.125 | 0.140 |

P833_RV3_5

Largest fascicle_widest and narrowest diameter

Subject 5_P836

### Nodose

| | Area | Mean | Min | Max | Angle | Length |
|---|---|---|---|---|---|---|
| 30 | 0.012 | 183.408 | 11 | 255 | -6.277 | 0.319 |
| 31 | 0.007 | 180.125 | 16 | 255 | 72.665 | 0.191 |
| 32 | 0.205 | 200.621 | 2 | 255 | 0.087 | 5.550 |

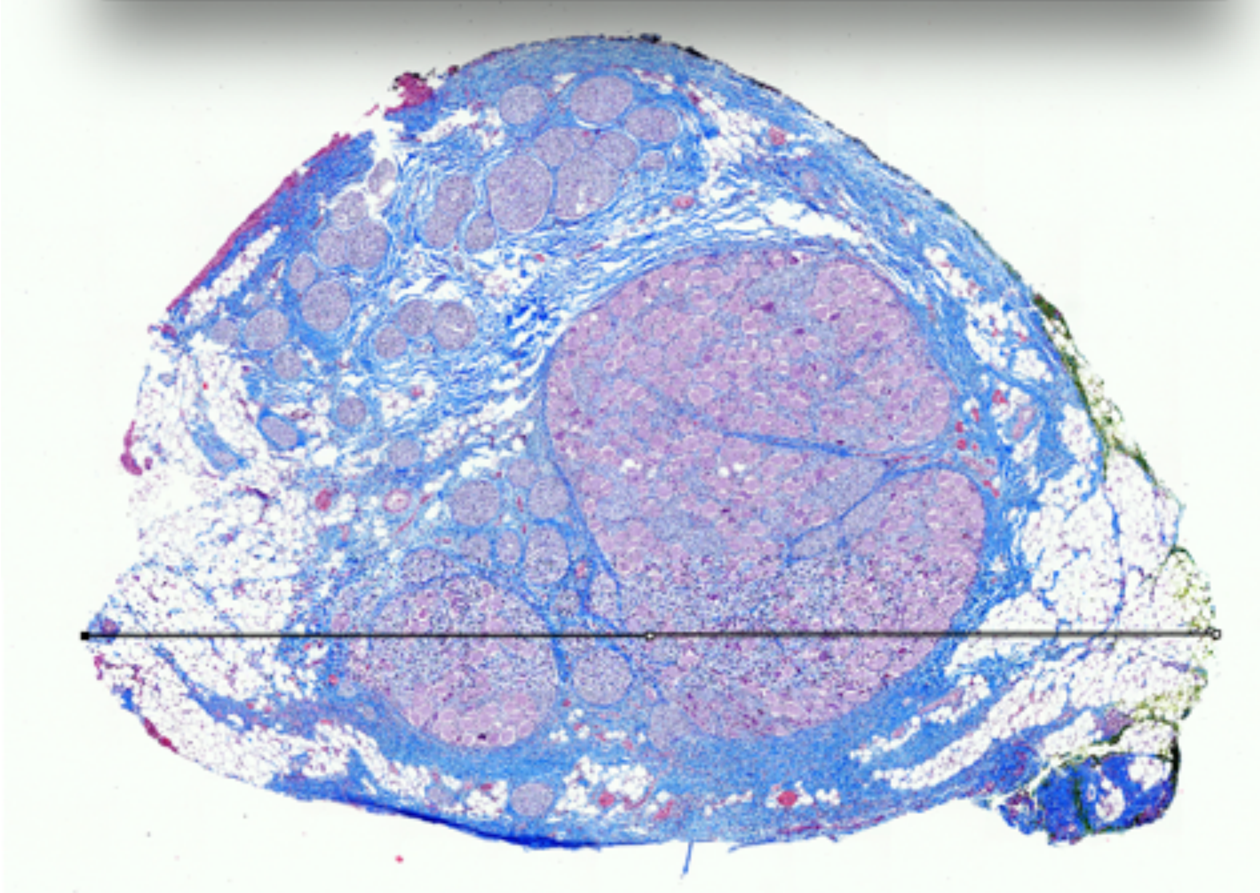

| | Area | Mean | Min | Max | Angle | Length |
|---|---|---|---|---|---|---|
| 1 | 0.070 | 187.943 | 2 | 255 | -93.594 | 2.628 |
| 2 | 0.069 | 186.855 | 4 | 255 | -95.683 | 2.623 |

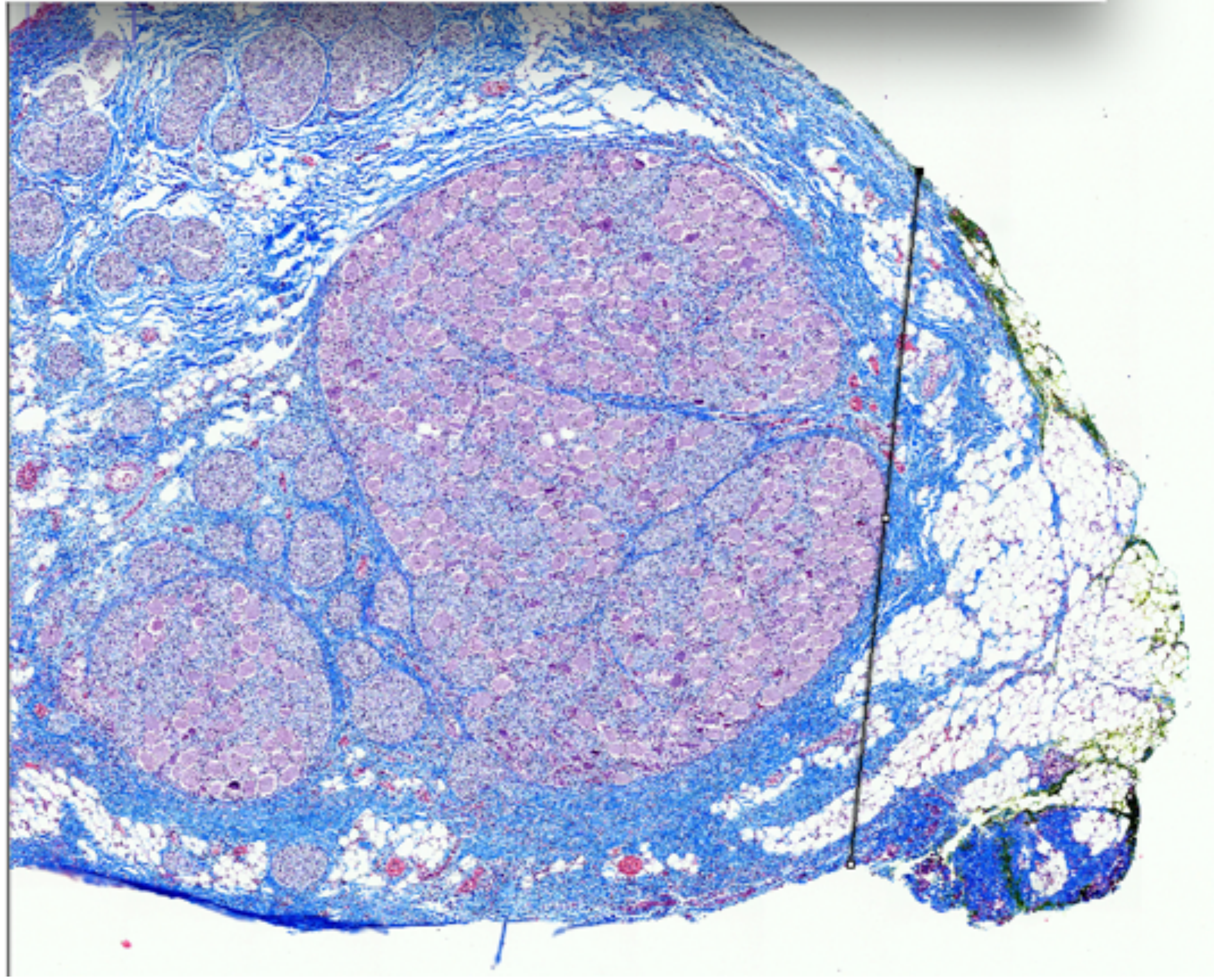

P836_RV3_41

Largest fascicle_widest and narrowest diameter

Nodose

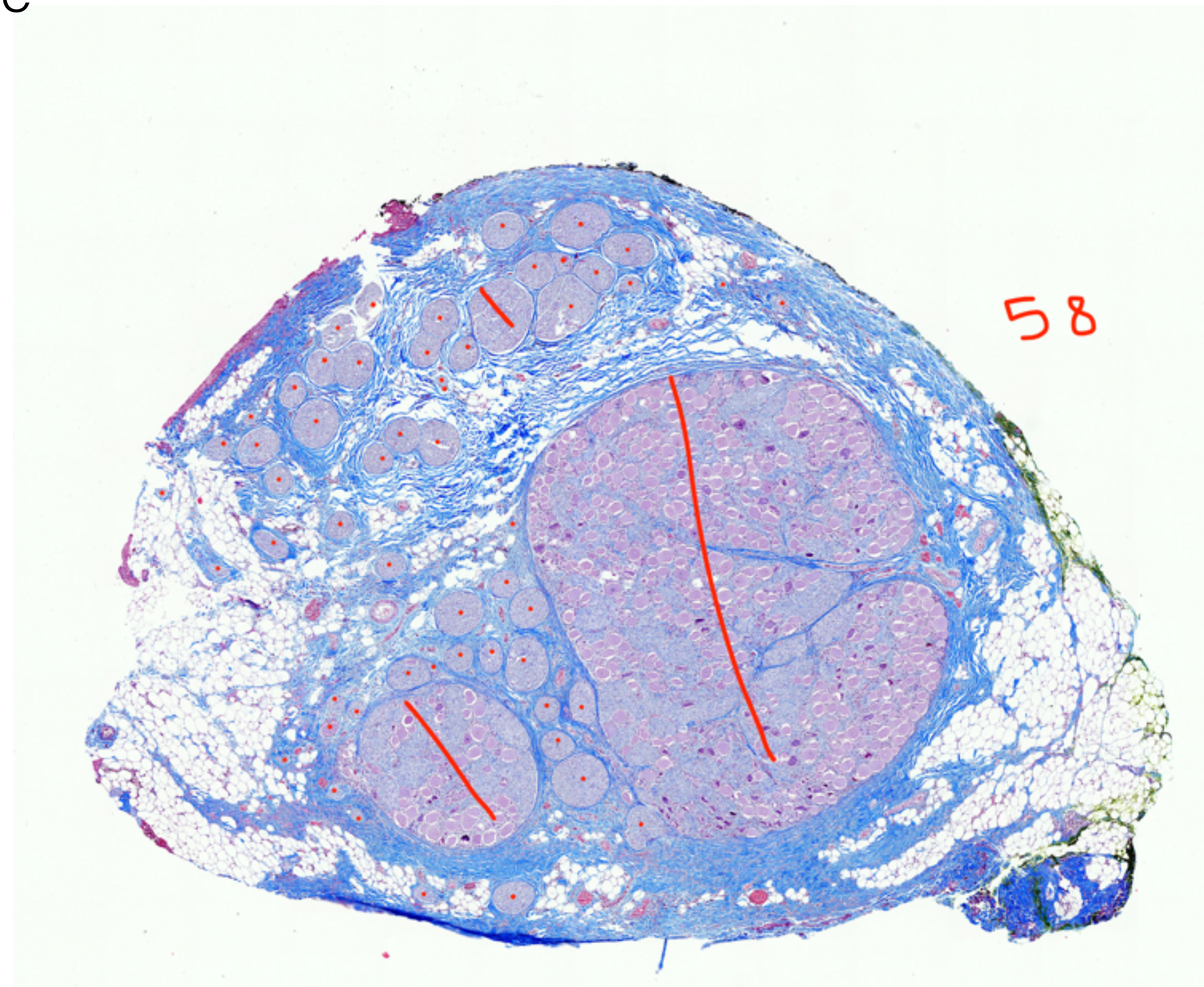


P836_RV3_41

Fascicle Count, 58

### Nodose

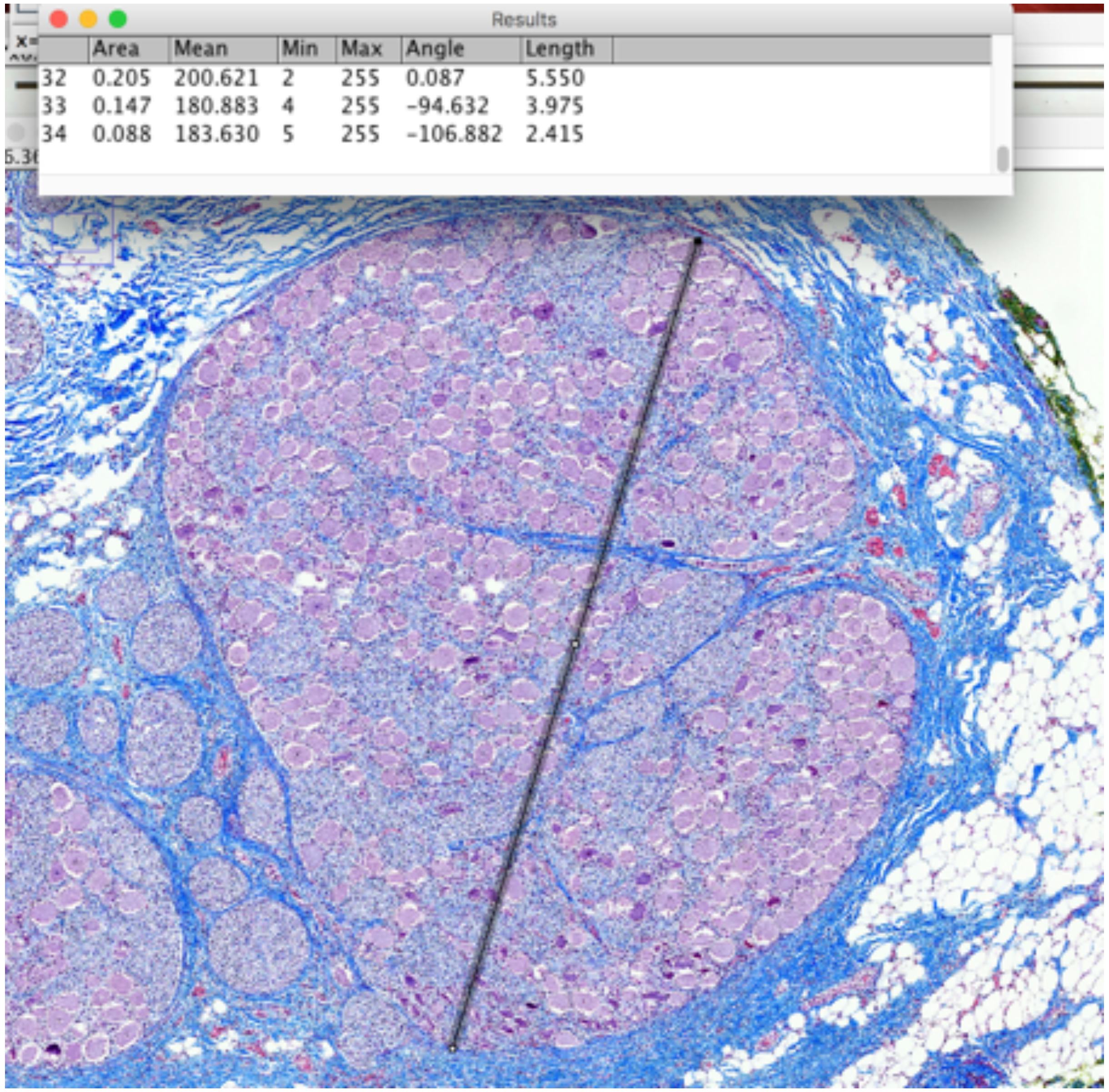

| | Area | Mean | Min | Max | Angle | Length |
|---|---|---|---|---|---|---|
| 32 | 0.205 | 200.621 | 2 | 255 | 0.087 | 5.550 |
| 33 | 0.147 | 180.883 | 4 | 255 | -94.632 | 3.975 |
| 34 | 0.088 | 183.630 | 5 | 255 | -106.882 | 2.415 |

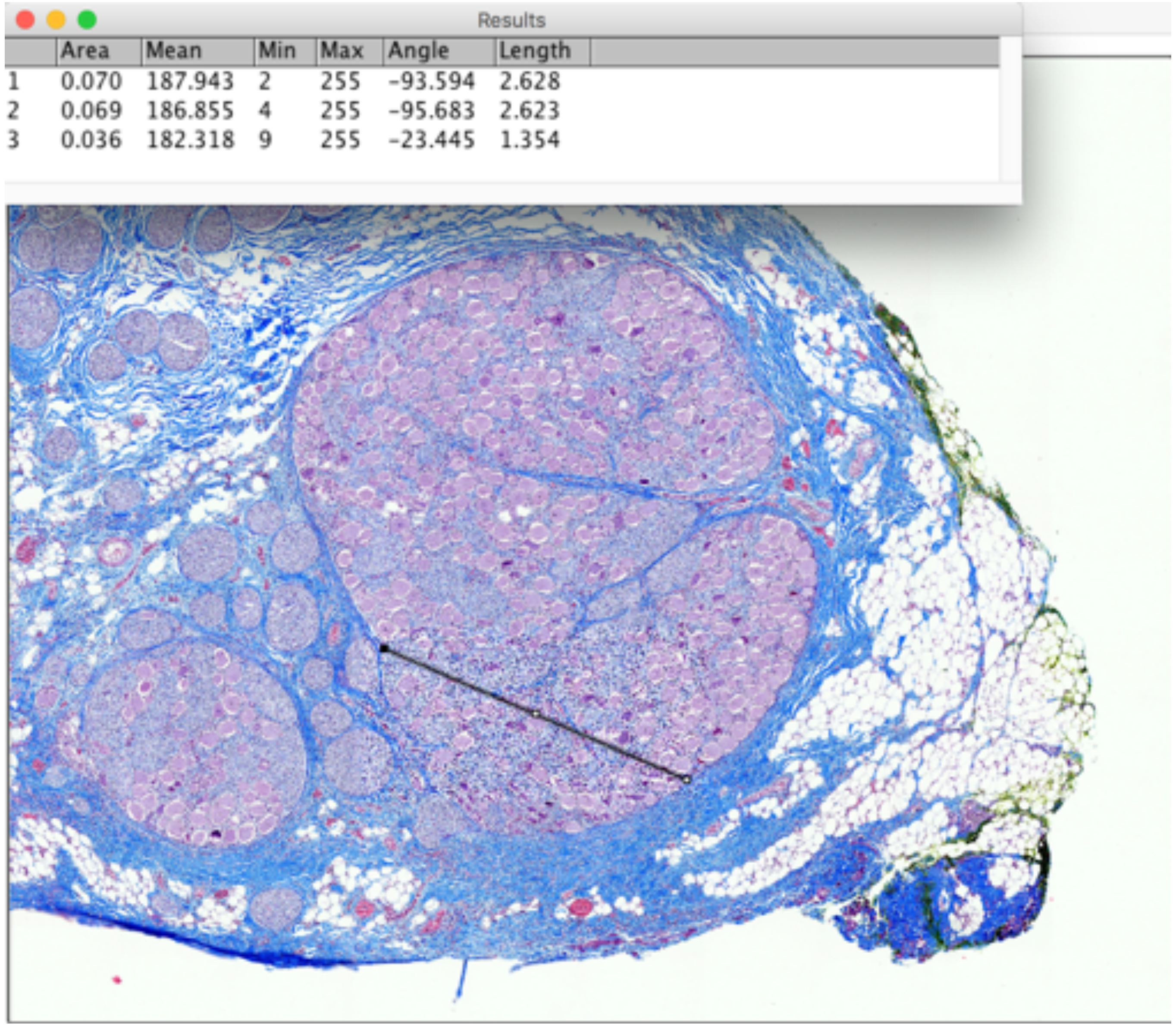

| | Area | Mean | Min | Max | Angle | Length |
|---|---|---|---|---|---|---|
| 1 | 0.070 | 187.943 | 2 | 255 | -93.594 | 2.628 |
| 2 | 0.069 | 186.855 | 4 | 255 | -95.683 | 2.623 |
| 3 | 0.036 | 182.318 | 9 | 255 | -23.445 | 1.354 |

P836_RV3_41

Largest Fascicle_Widest and narrowest diameter

### Mid-VN

Results

| Mean | Min | Max | Angle | Length |
|---|---|---|---|---|
| 183.630 | 5 | 255 | -106.882 | 2.415 |
| 188.232 | 11 | 255 | -31.581 | 2.355 |
| 168.908 | 2 | 255 | -90.201 | 3.618 |

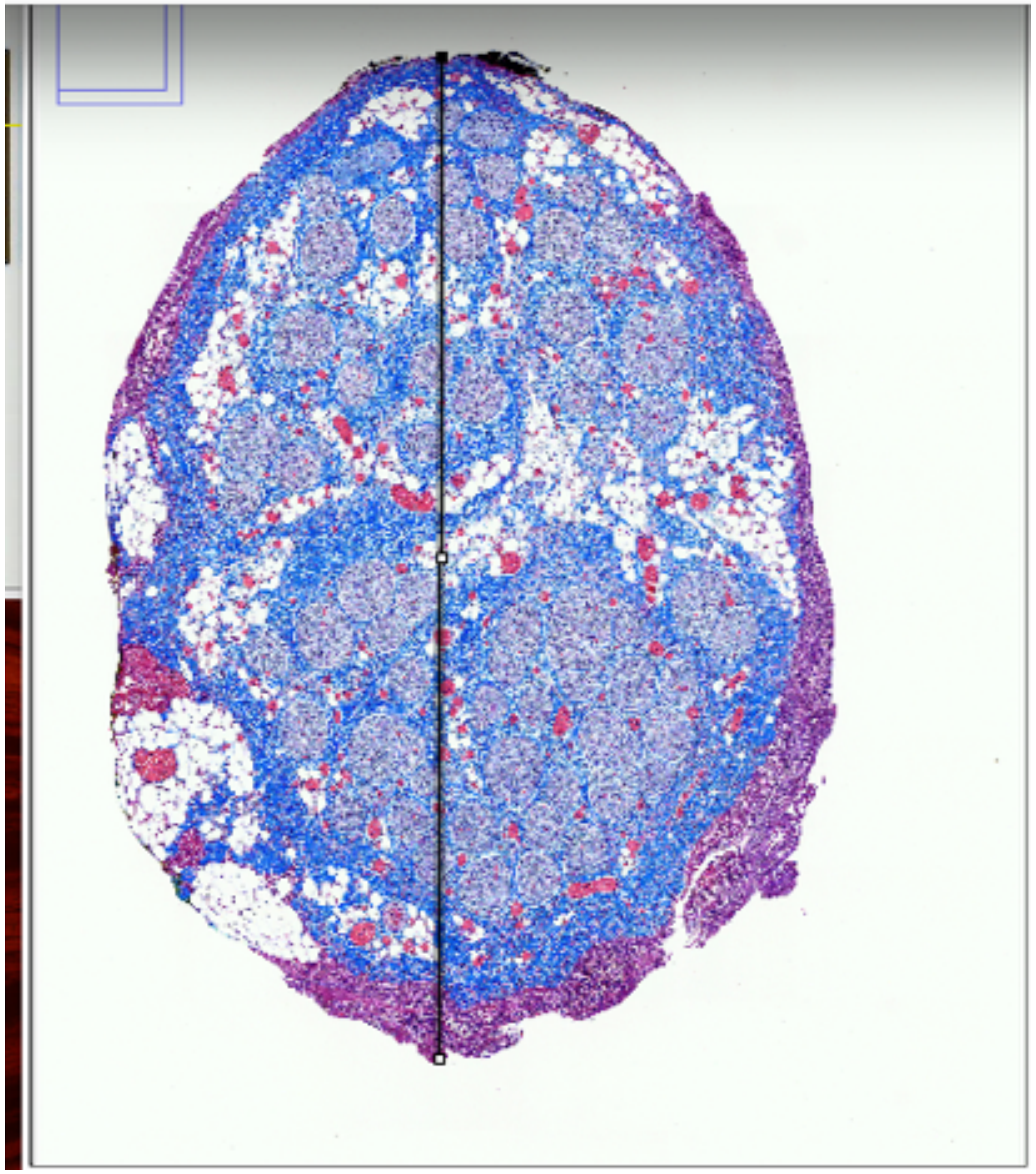

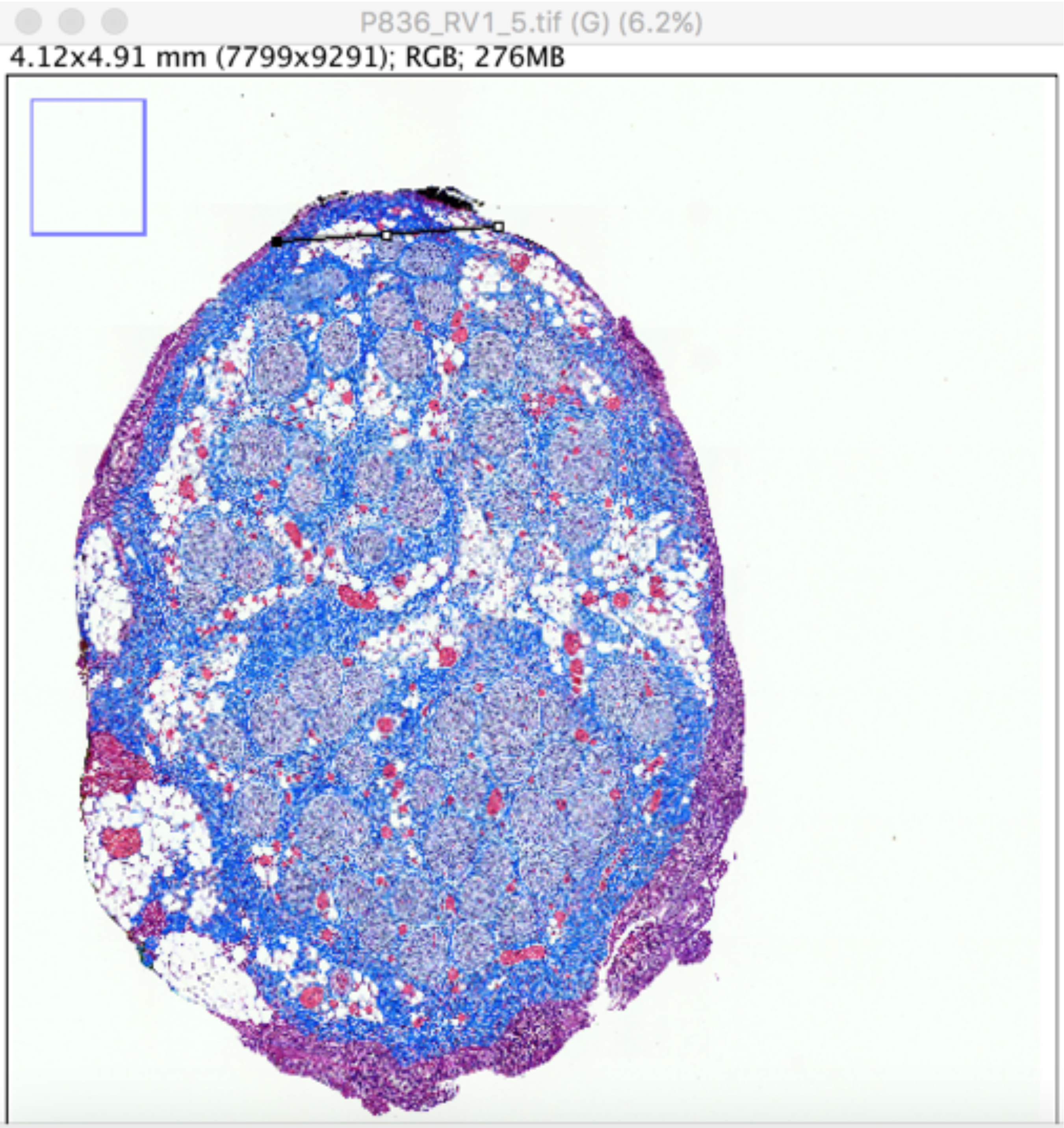


Results

| | Area | Mean | Min | Max | Angle | Length |
|---|---|---|---|---|---|---|
| 1 | 0.023 | 170.161 | 5 | 255 | 3.678 | 0.872 |

# P836_RV1_5

Widest and narrowest diameter

### Mid-VN

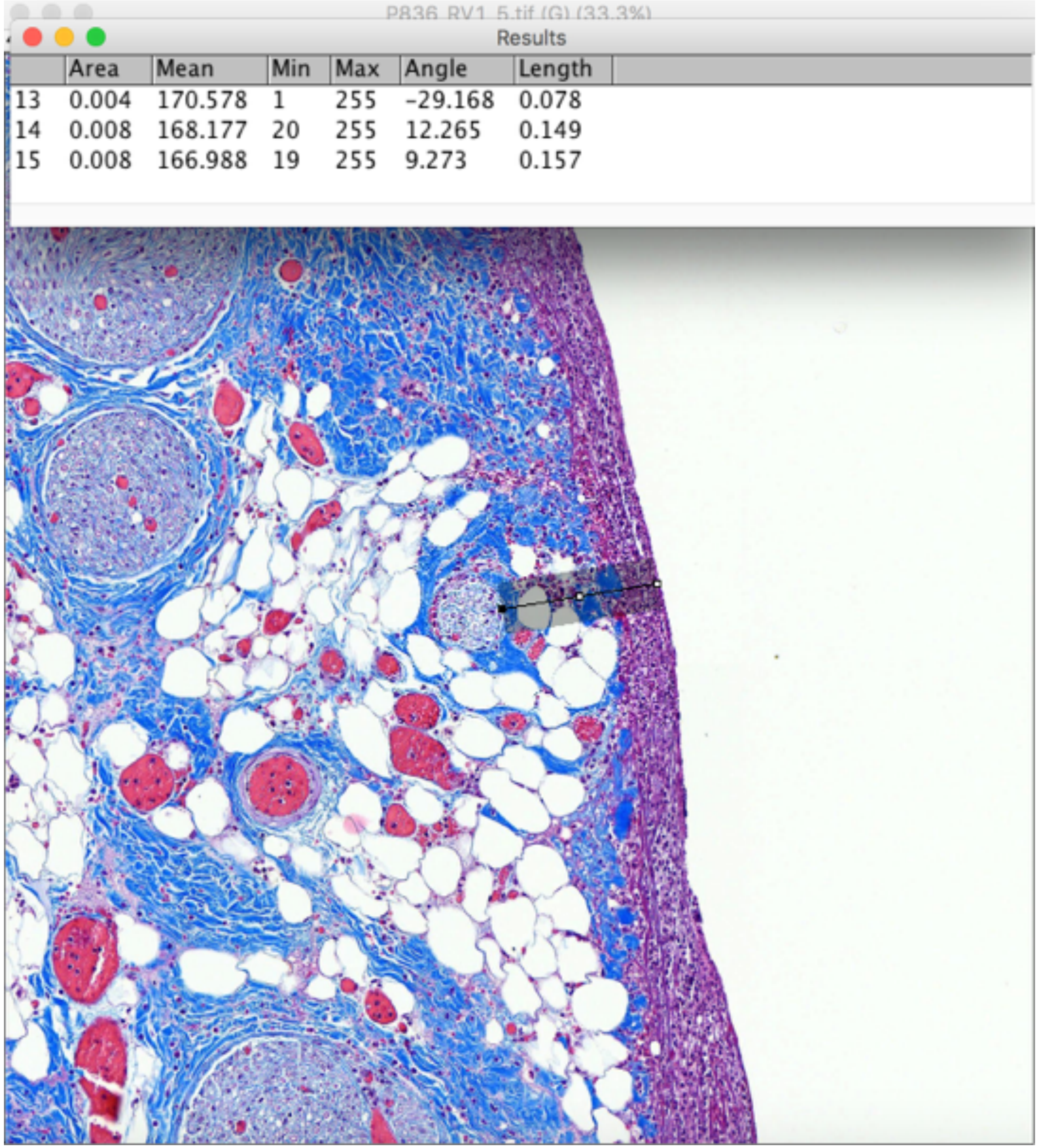


| | Area | Mean | Min | Max | Angle | Length |
|---|---|---|---|---|---|---|
| 13 | 0.004 | 170.578 | 1 | 255 | -29.168 | 0.078 |
| 14 | 0.008 | 168.177 | 20 | 255 | 12.265 | 0.149 |
| 15 | 0.008 | 166.988 | 19 | 255 | 9.273 | 0.157 |

P836_RV1_5

Fascicle depth

Mid-VN

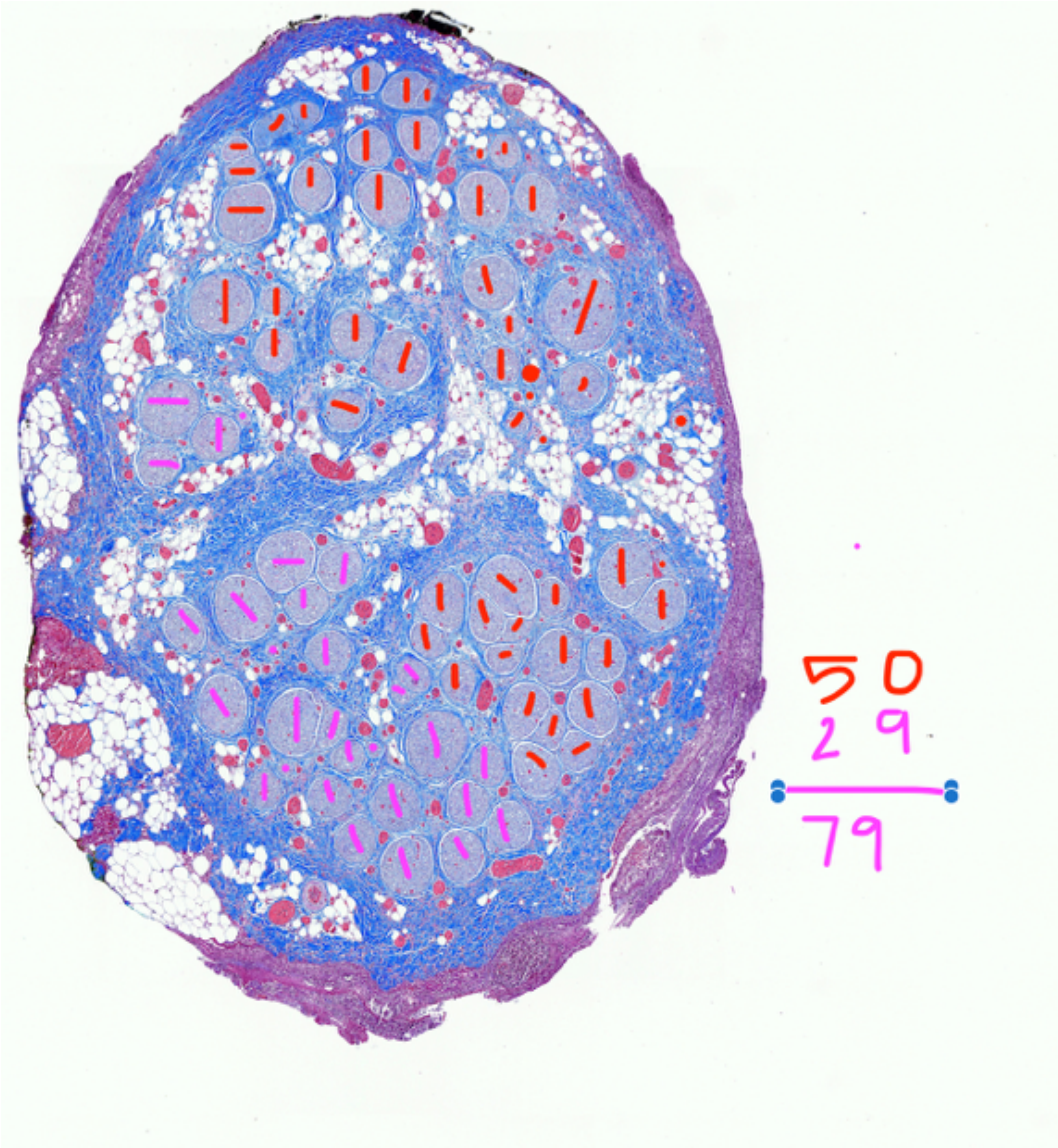


P836_RV1_5

Fascicle count

### Mid-VN

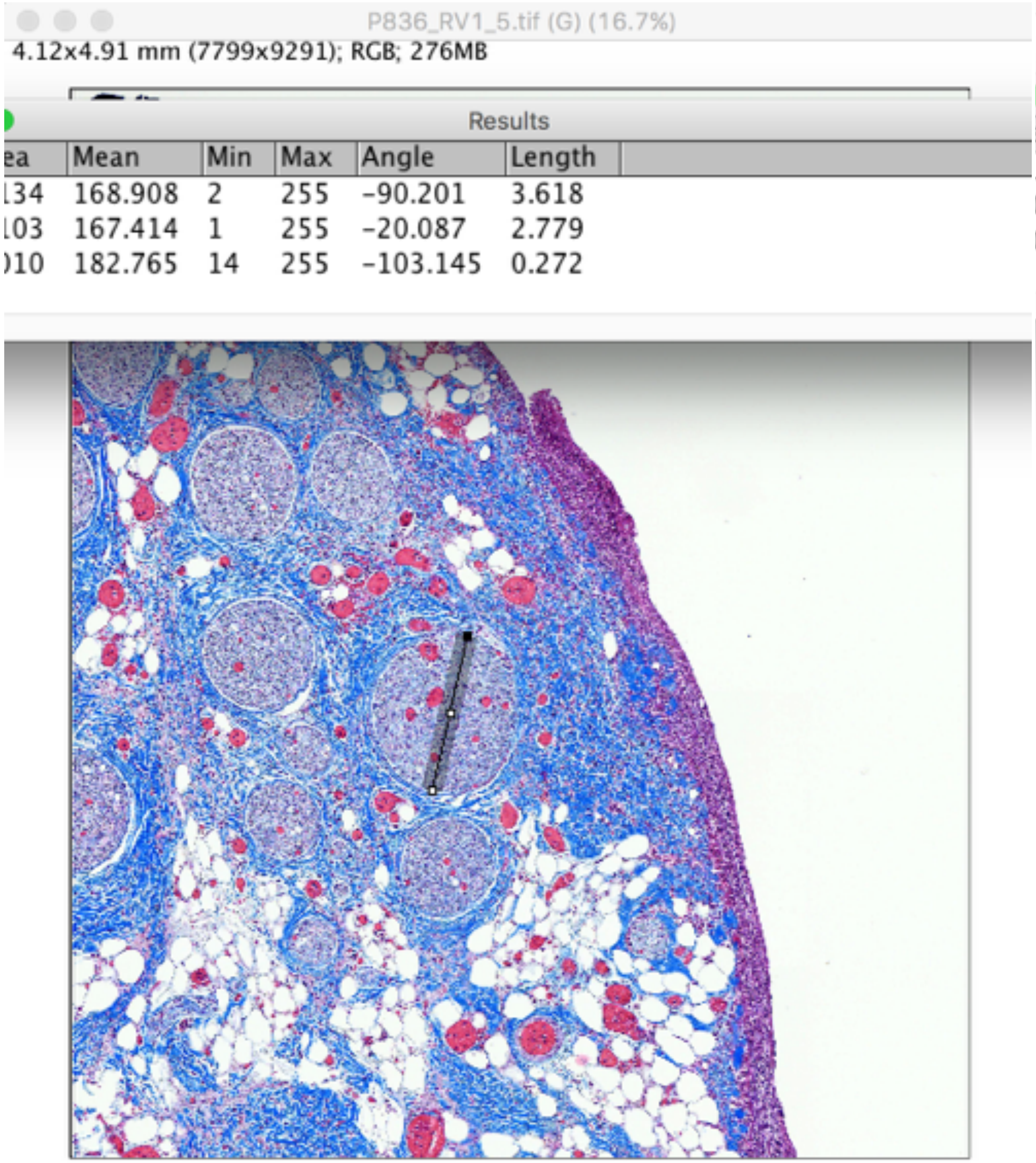


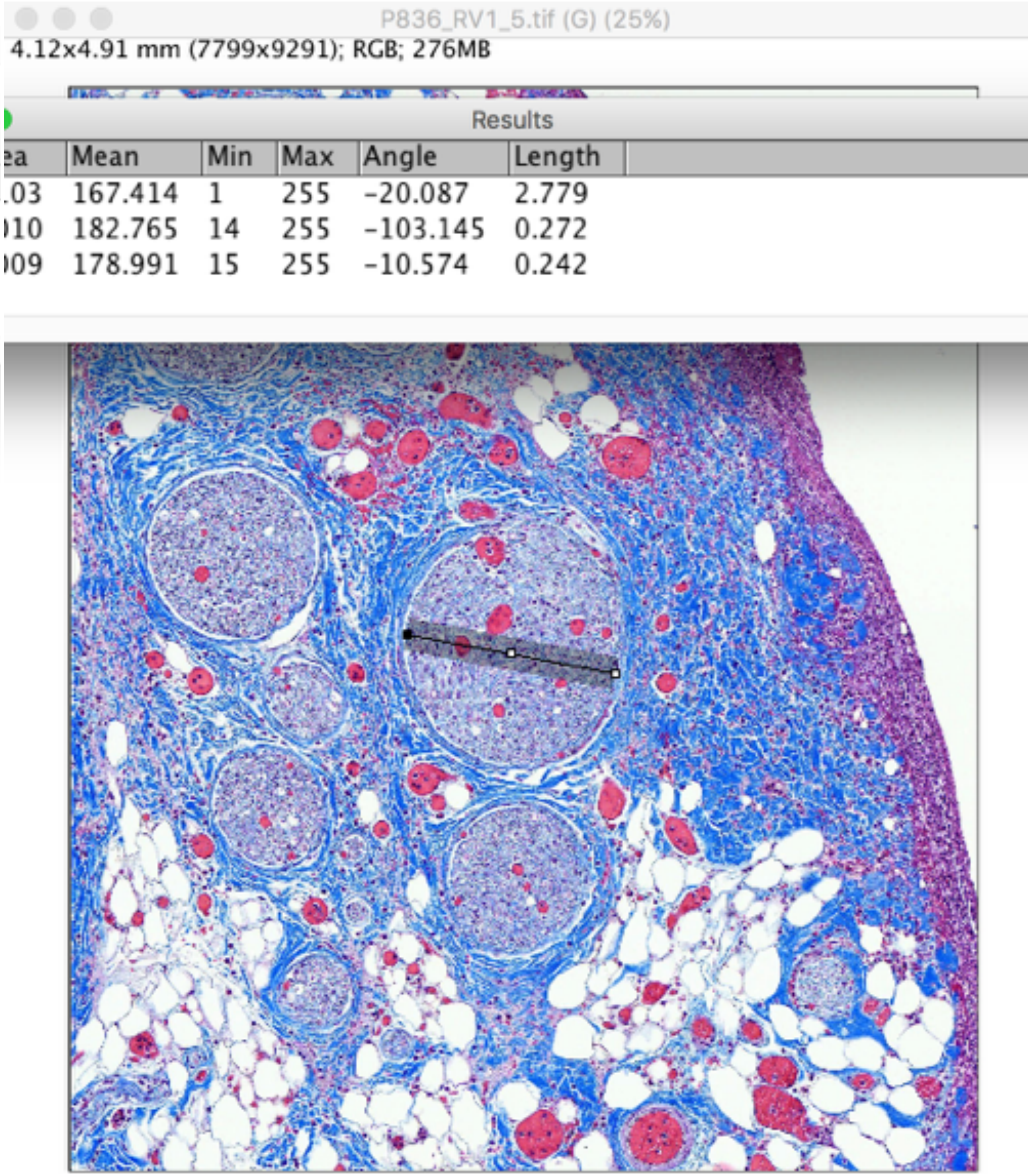


P836_RV1_5

Largest Fascicle_Widest and narrowest diameter

Subject 8_P858

### Pre-nodose

Results

| | Area | Mean | Min | Max | Angle | Length |
|---|---|---|---|---|---|---|
| 38 | 0.010 | 182.765 | 14 | 255 | -103.145 | 0.272 |
| 39 | 0.009 | 178.991 | 15 | 255 | -10.574 | 0.242 |
| 40 | 0.189 | 181.404 | 4 | 255 | -92.564 | 5.099 |

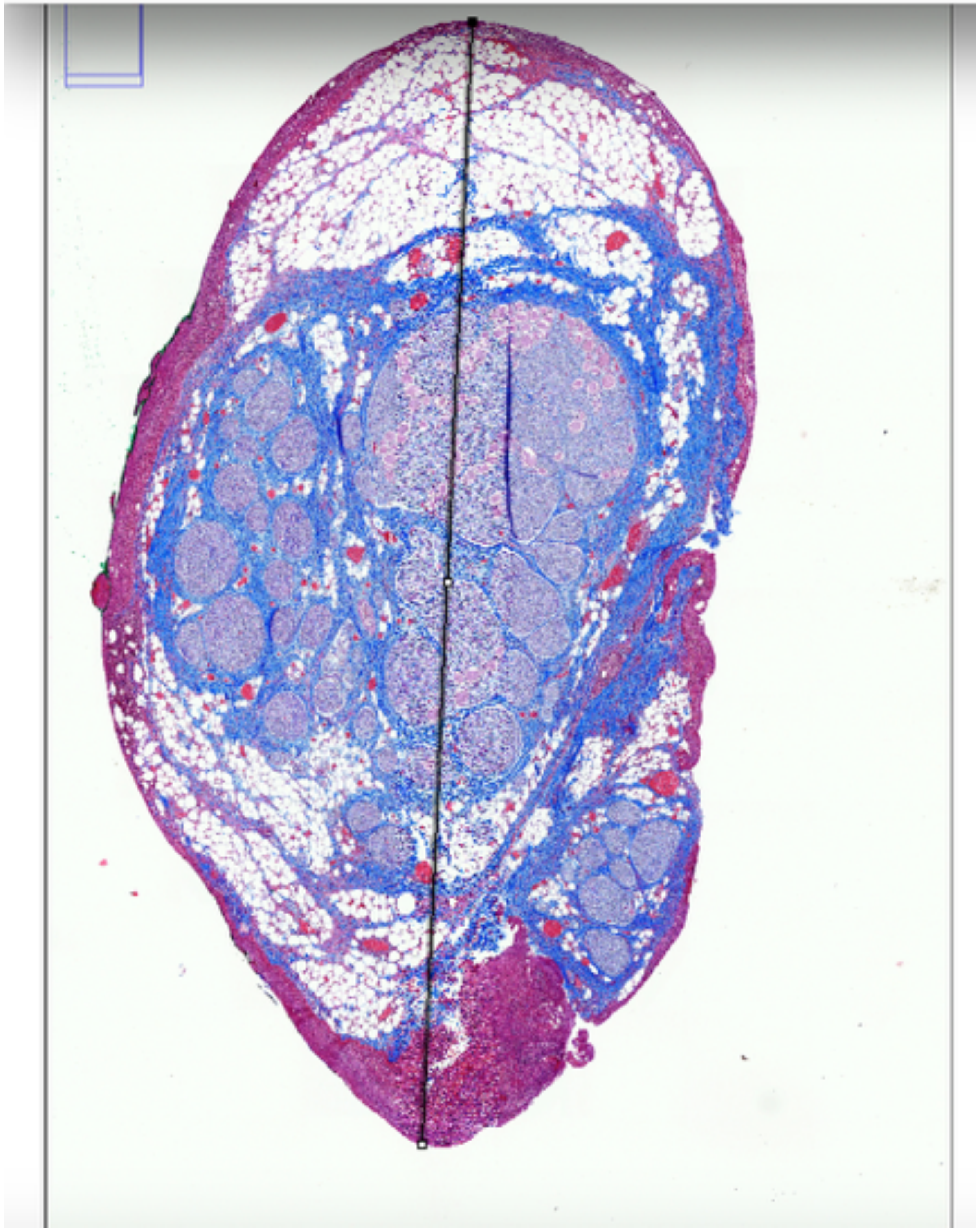

19P858_Nodose_35.tif (G) (4.2%)

4.11x6.36 mm (7785x12055); RGB; 358MB

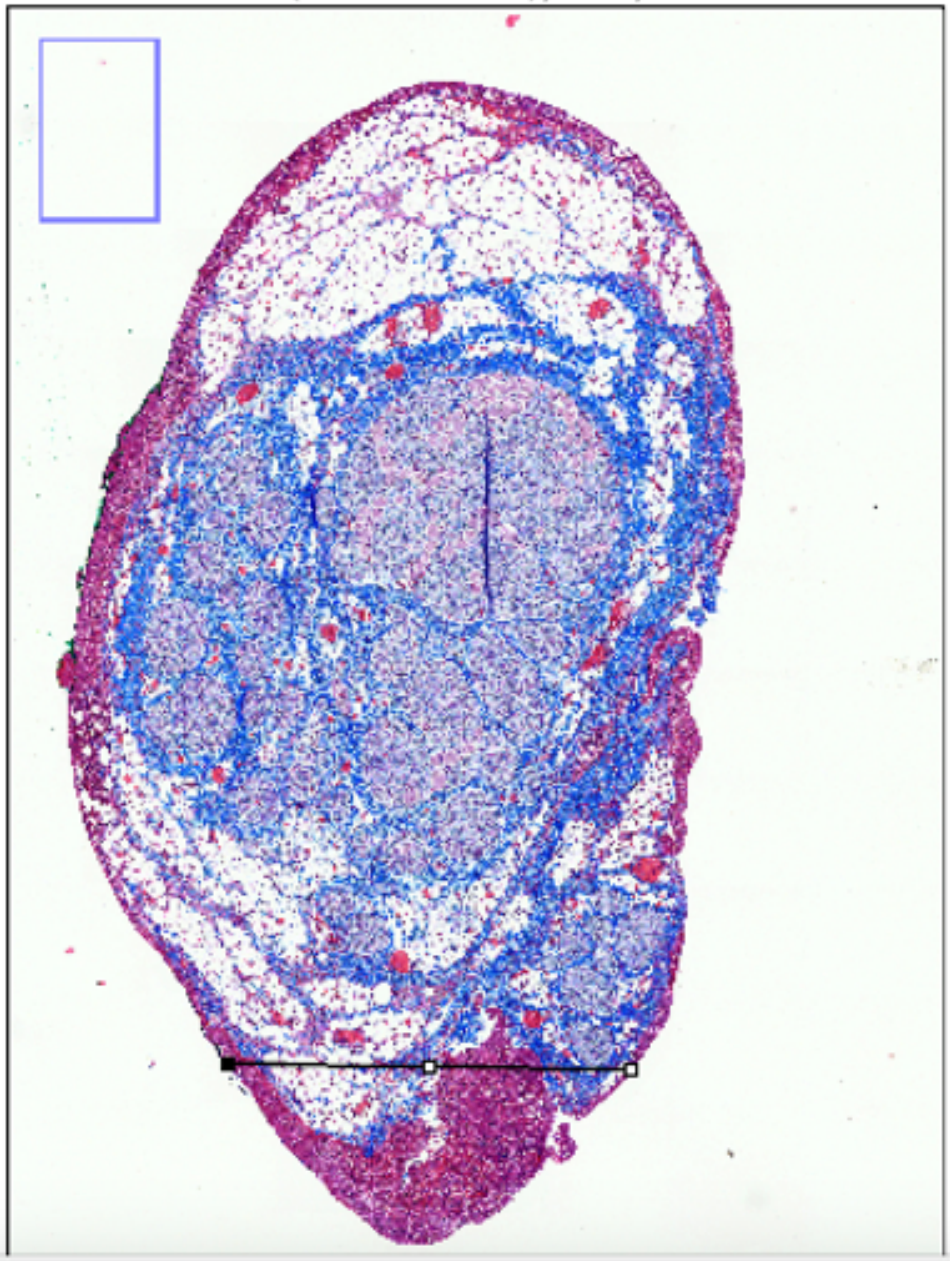

Results

| Area | Mean | Min | Max | Angle | Length |
|---|---|---|---|---|---|
| 0.023 | 170.161 | 5 | 255 | 3.678 | 0.872 |
| 0.047 | 158.753 | 4 | 255 | -1.144 | 1.772 |
| 0.047 | 158.753 | 4 | 255 | -1.144 | 1.772 |

P858_nodose_35

Widest and narrowest diameter

### Pre-nodose

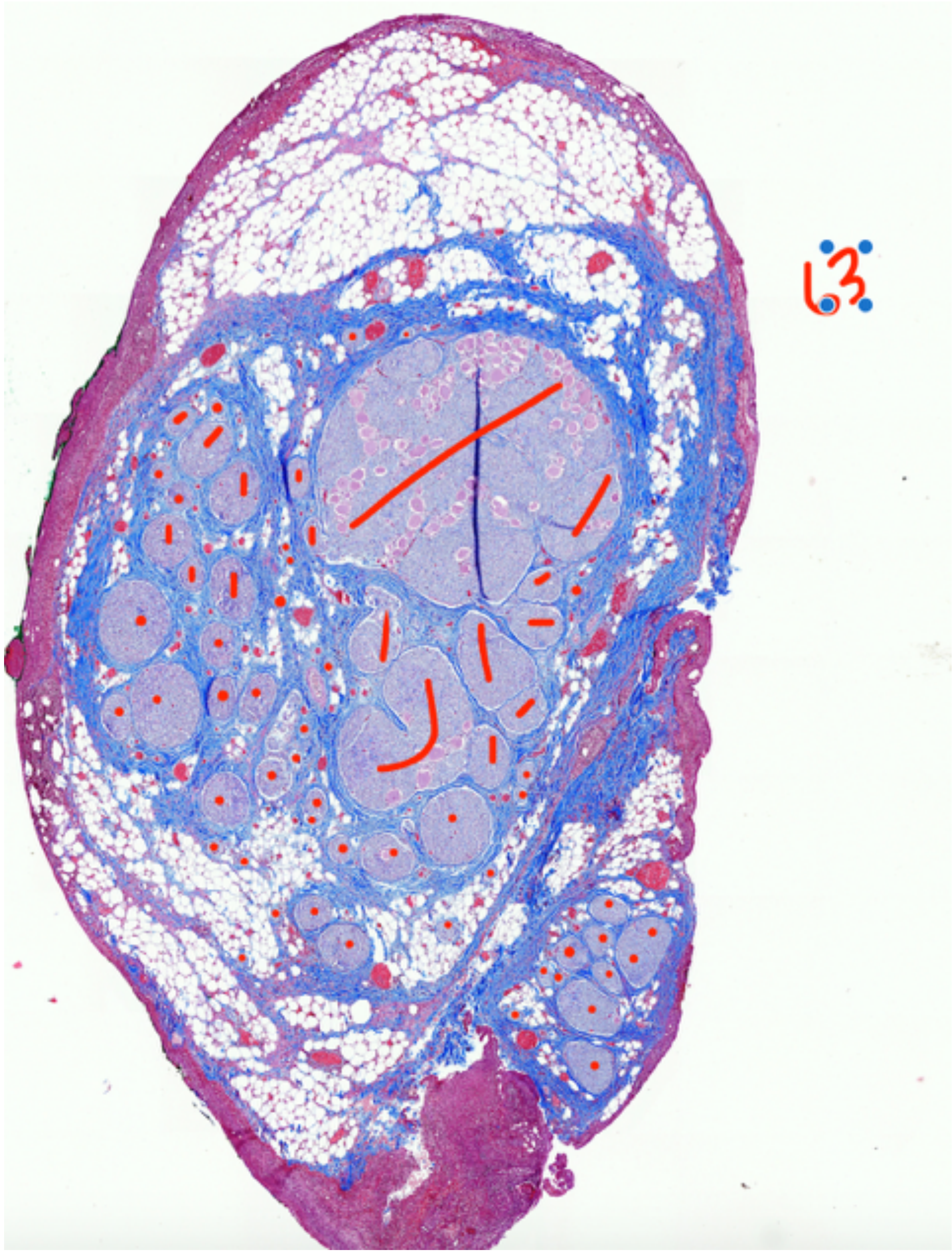


P858_nodose_35

Fascicle count, 63

### Pre-nodose

Results

| | Area | Mean | Min | Max | Angle | Length |
|---|---|---|---|---|---|---|
| 40 | 0.189 | 181.404 | 4 | 255 | -92.564 | 5.099 |
| 41 | 0.106 | 161.978 | 1 | 255 | -6.660 | 2.841 |
| 42 | 0.046 | 183.856 | 11 | 255 | 33.410 | 1.260 |

Results

| | Area | Mean | Min | Max | Angle | Length |
|---|---|---|---|---|---|---|
| 41 | 0.106 | 161.978 | 1 | 255 | -6.660 | 2.841 |
| 42 | 0.046 | 183.856 | 11 | 255 | 33.410 | 1.260 |
| 43 | 0.040 | 175.754 | 10 | 255 | -73.269 | 1.078 |

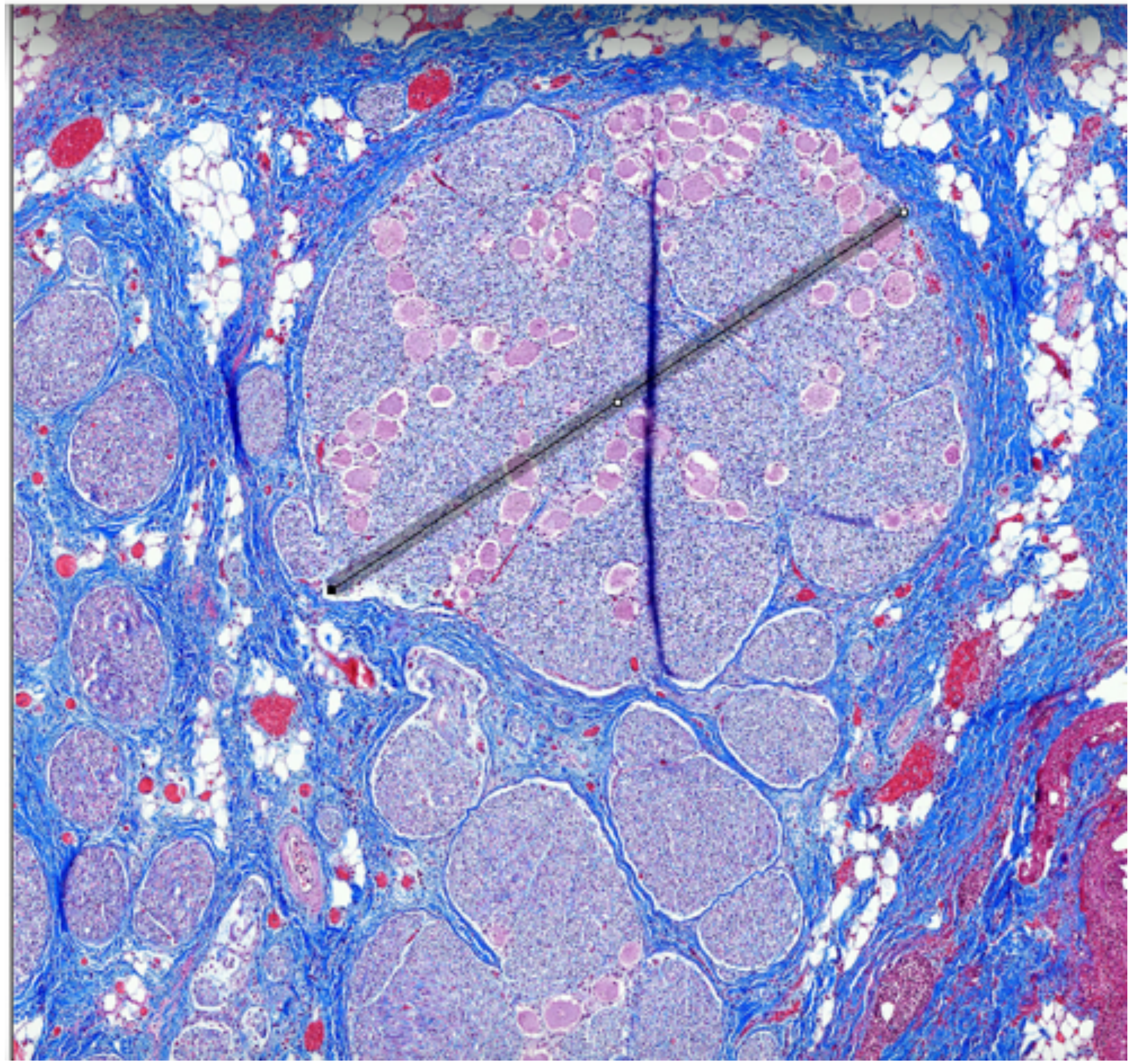

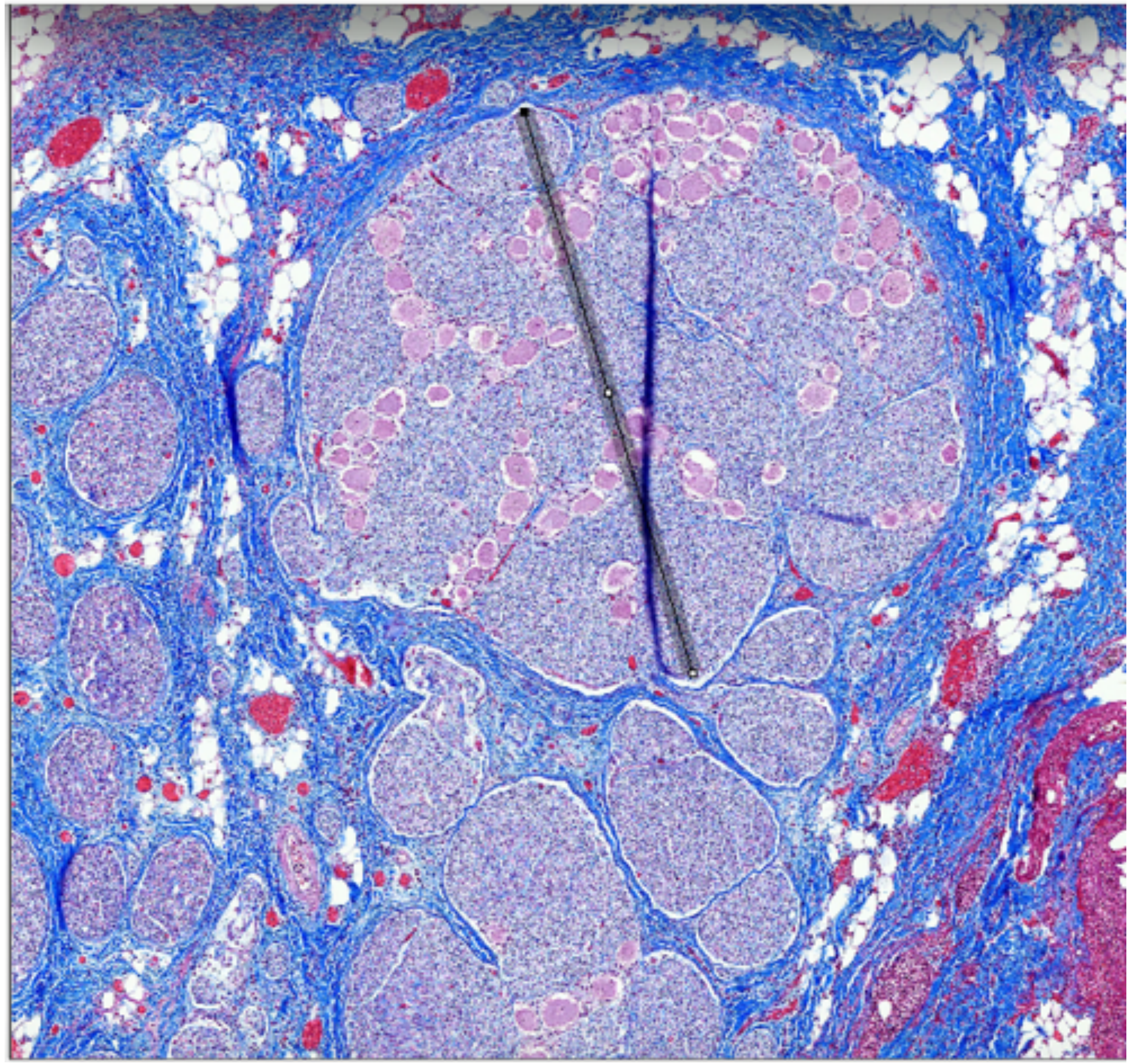

P858_nodose_35

Largest fascicle_Widest and narrowest diameter

### Nodose

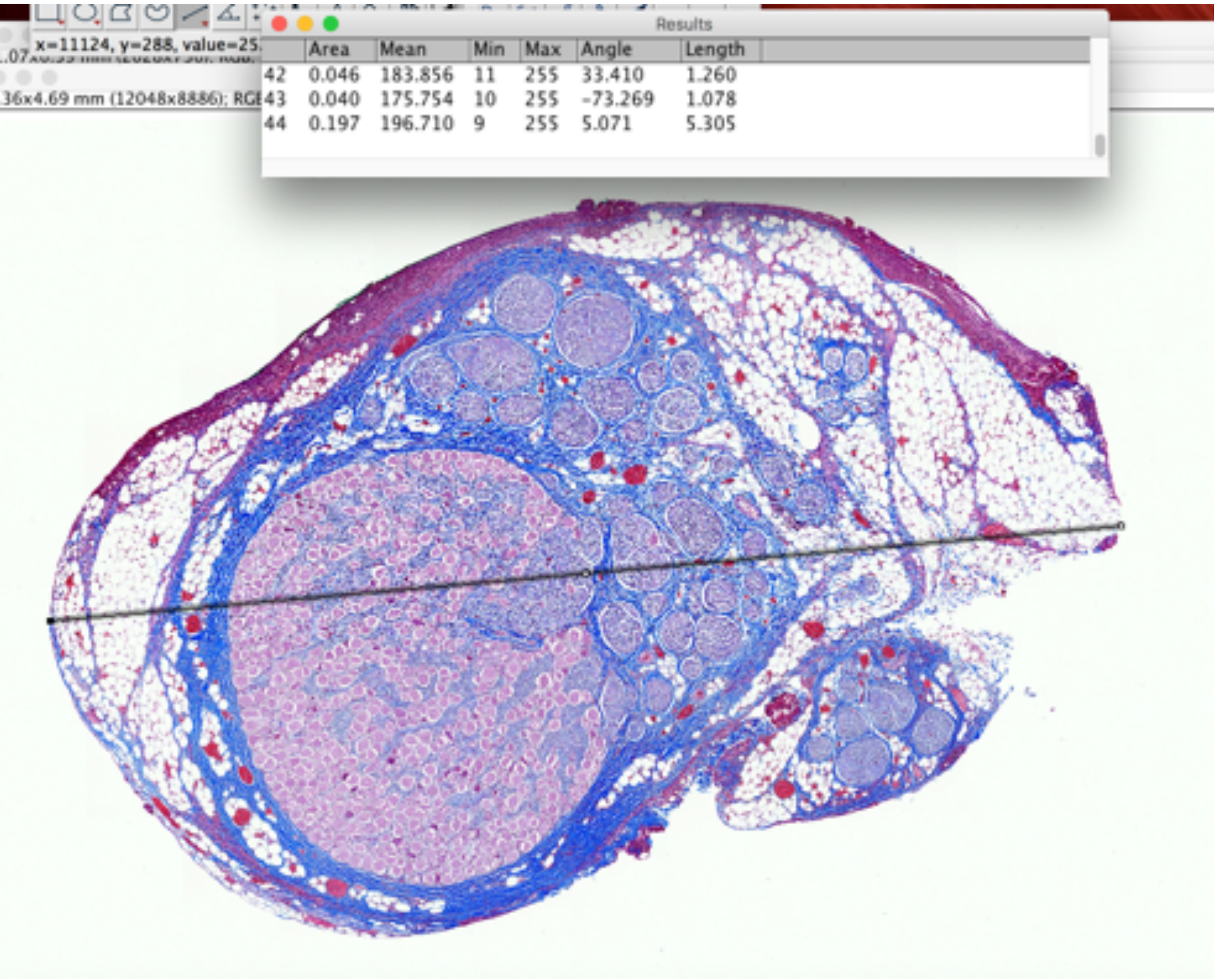


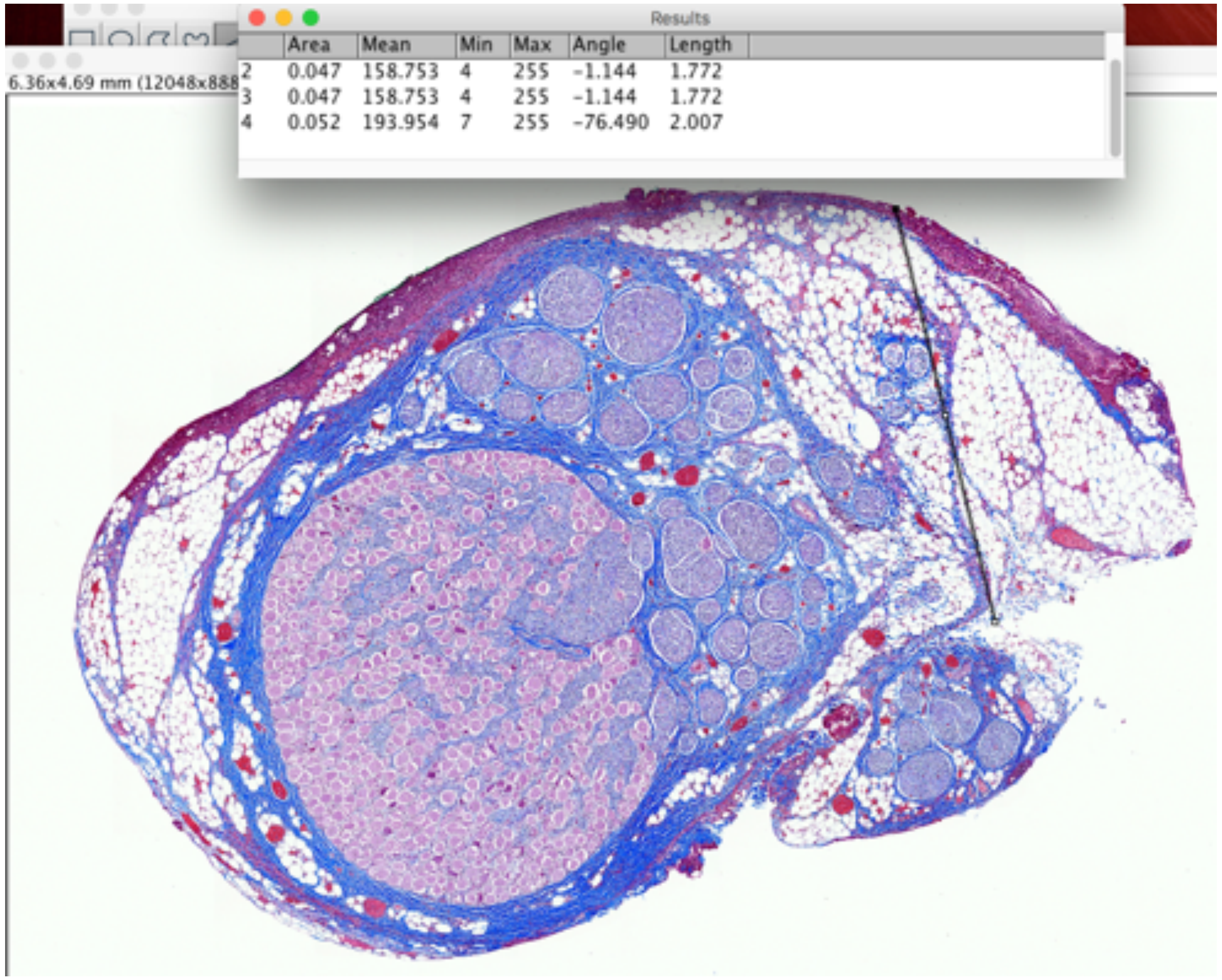


P858_nodose_8

Widest and narrowest diameter

### Nodose

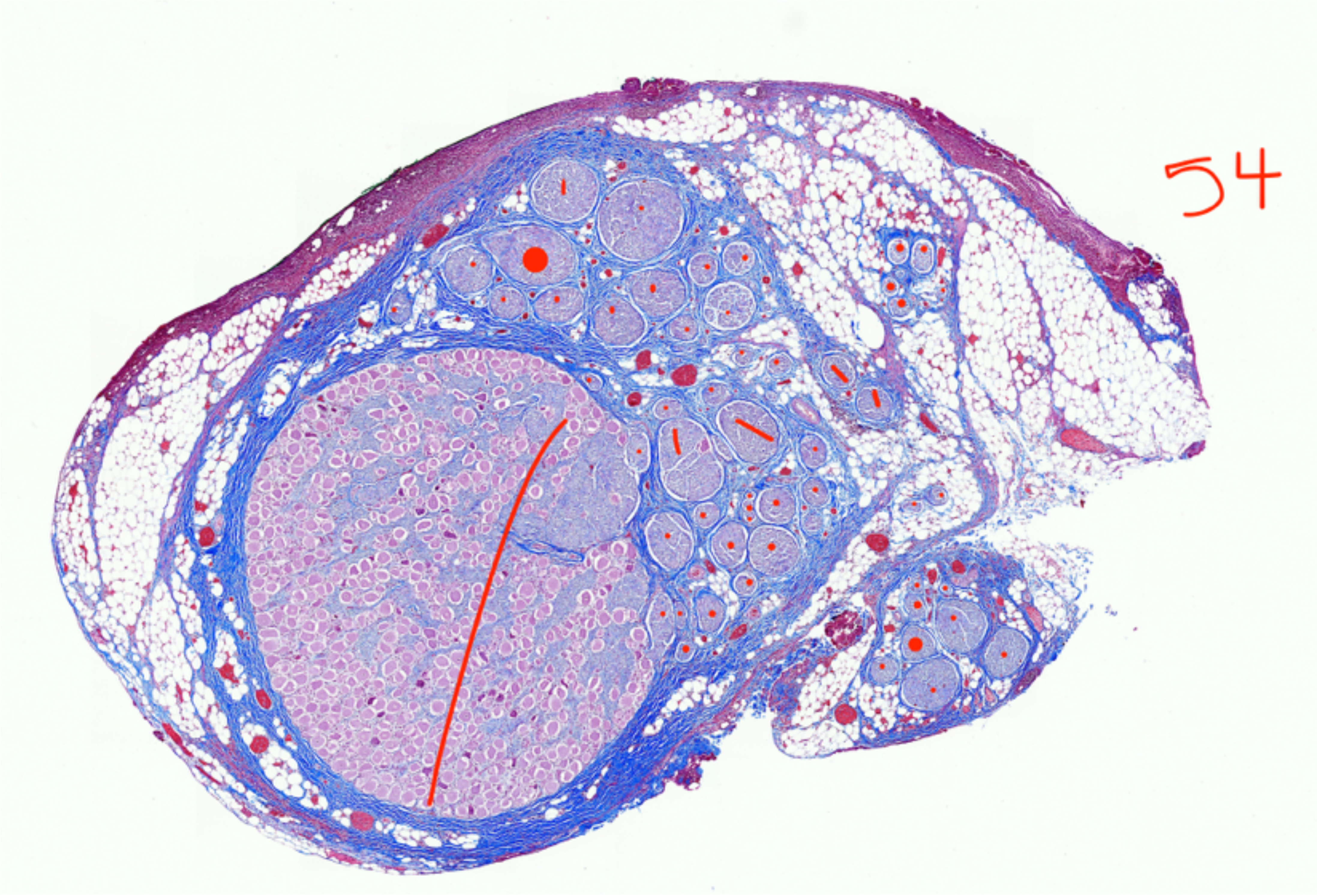


P858_nodose_8

Fascicle count, 54

### Nodose

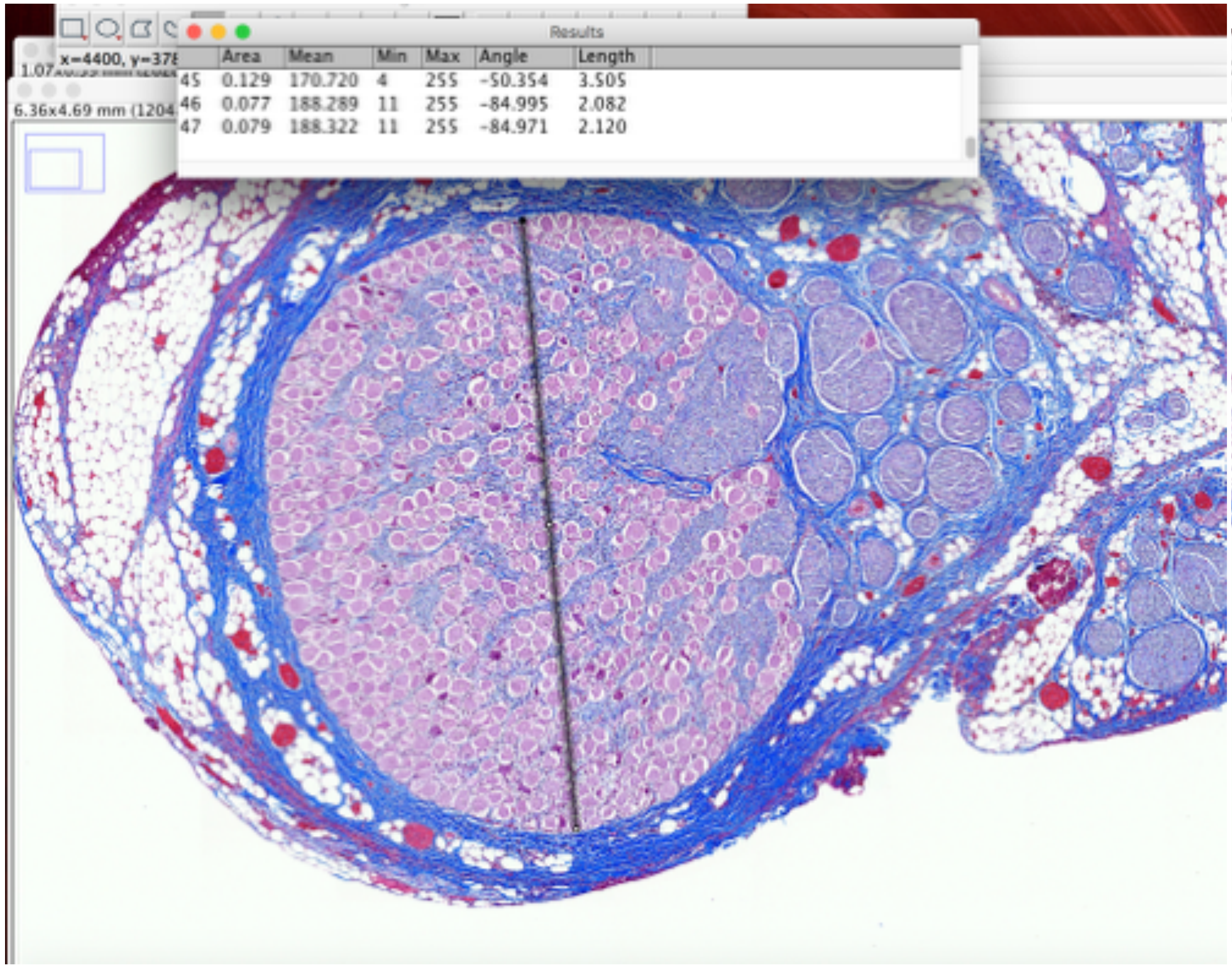


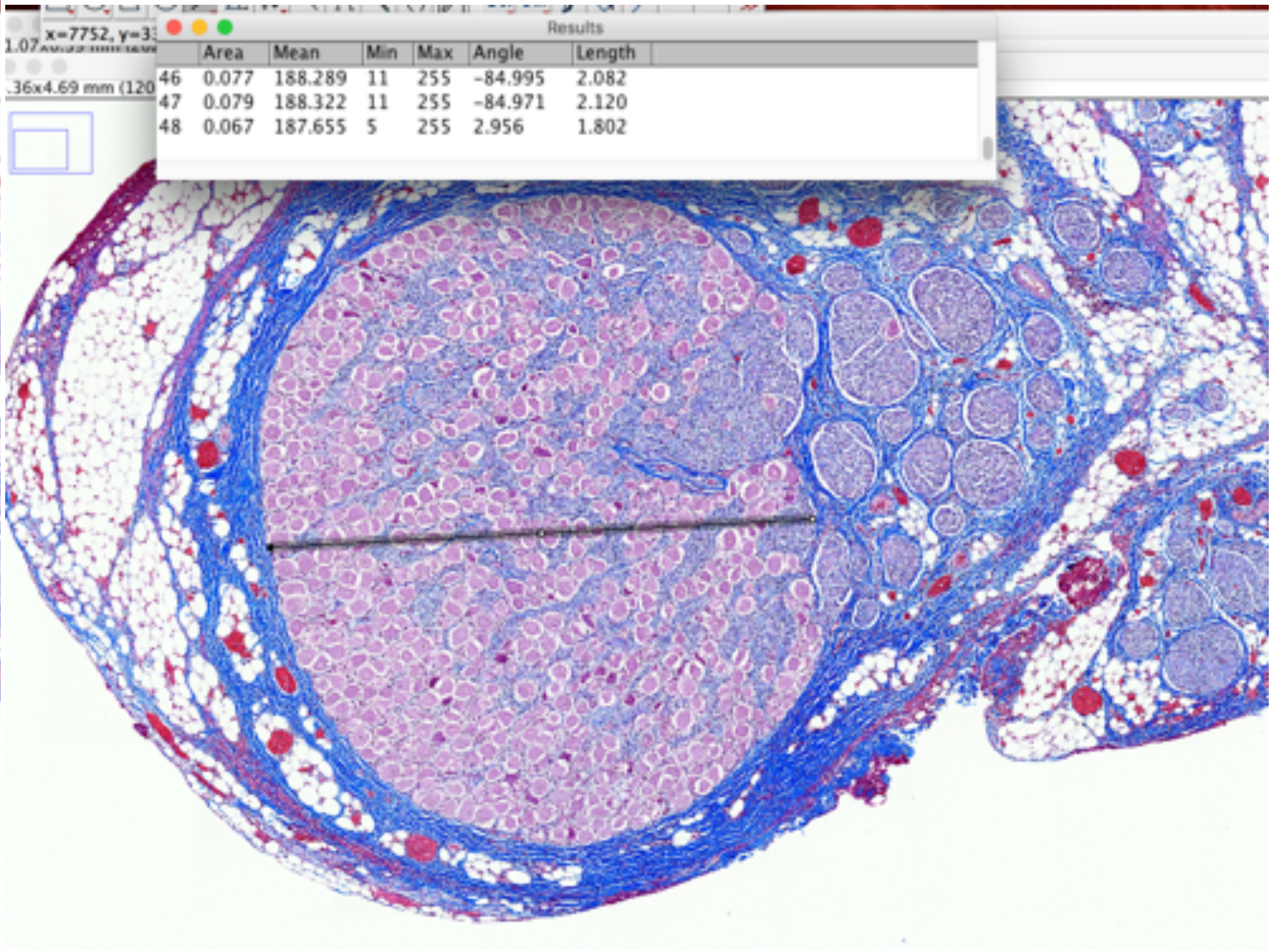


P858_nodose_8

Largest Fascicle_Widest and narrowest diameter

### Mid-VN

Results

| | Area | Mean | Min | Max | Angle | Length |
|---|---|---|---|---|---|---|
| 47 | 0.079 | 188.322 | 11 | 255 | -84.971 | 2.120 |
| 48 | 0.067 | 187.655 | 5 | 255 | 2.956 | 1.802 |
| 49 | 0.149 | 180.749 | 9 | 255 | 76.246 | 4.051 |

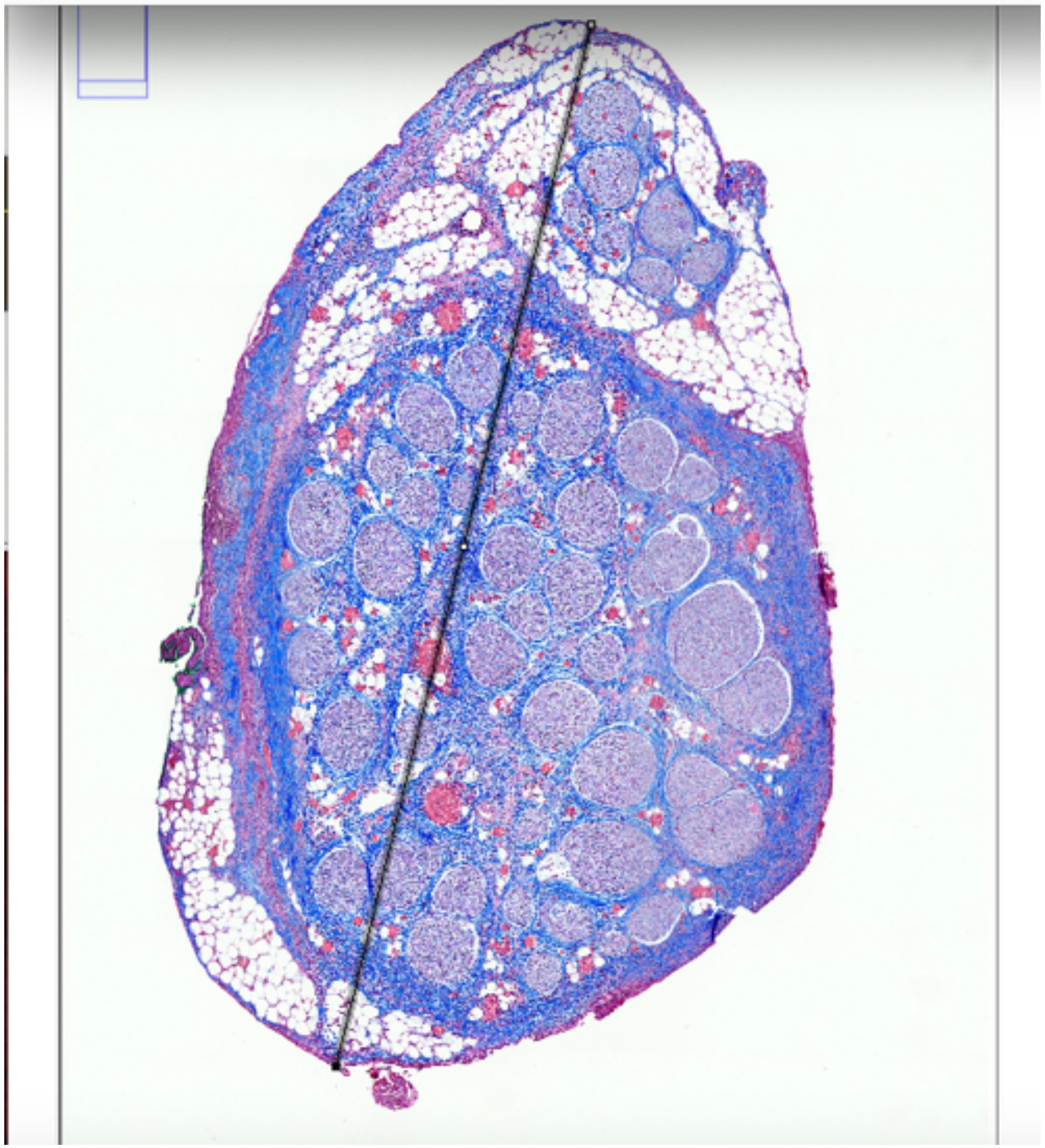

Results

| | Area | Mean | Min | Max | Angle | Length |
|---|---|---|---|---|---|---|
| 1 | 0.021 | 209.342 | 17 | 255 | -8.125 | 0.777 |

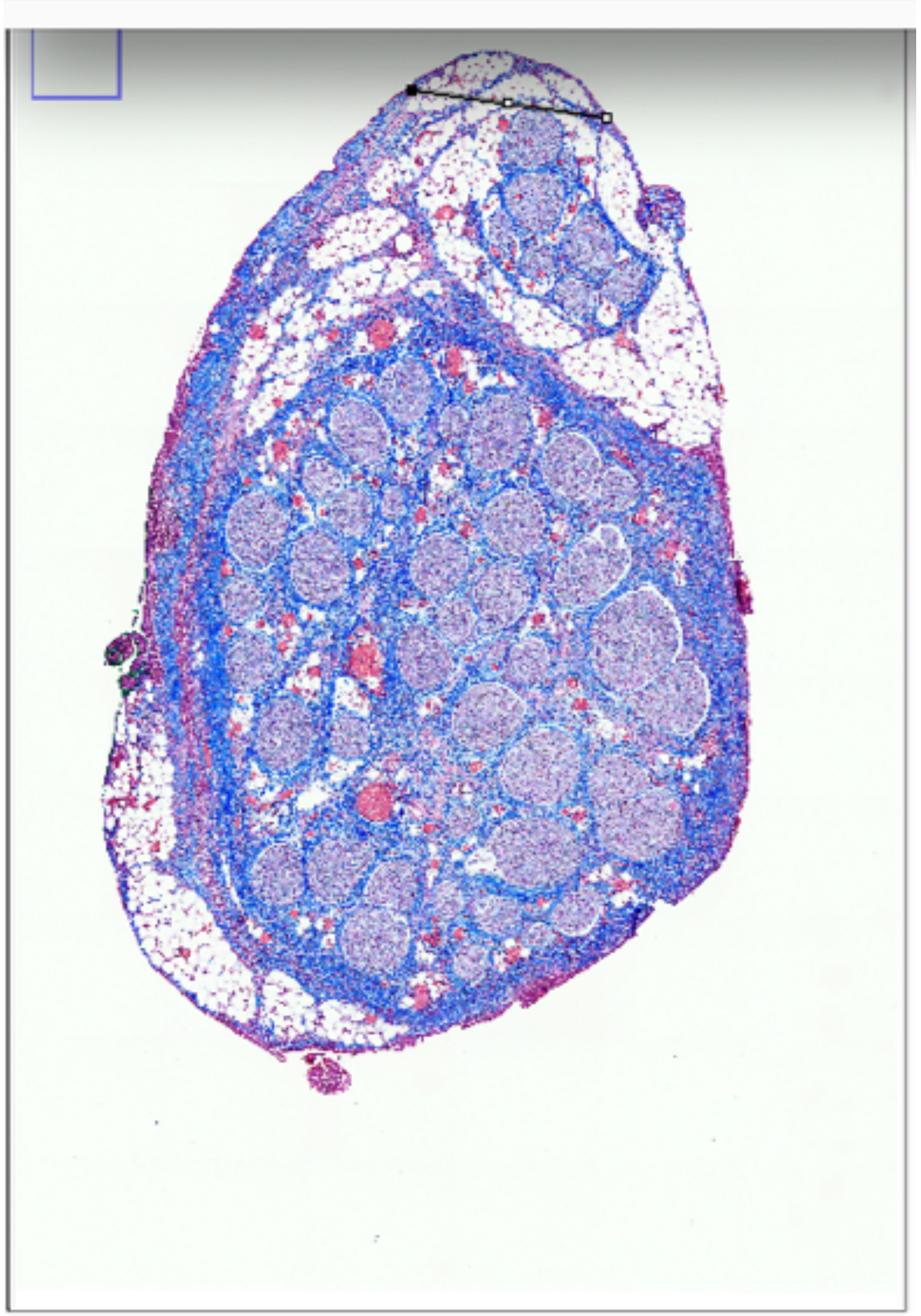

P858_LivaNova_43

Widest and narrowest diameter

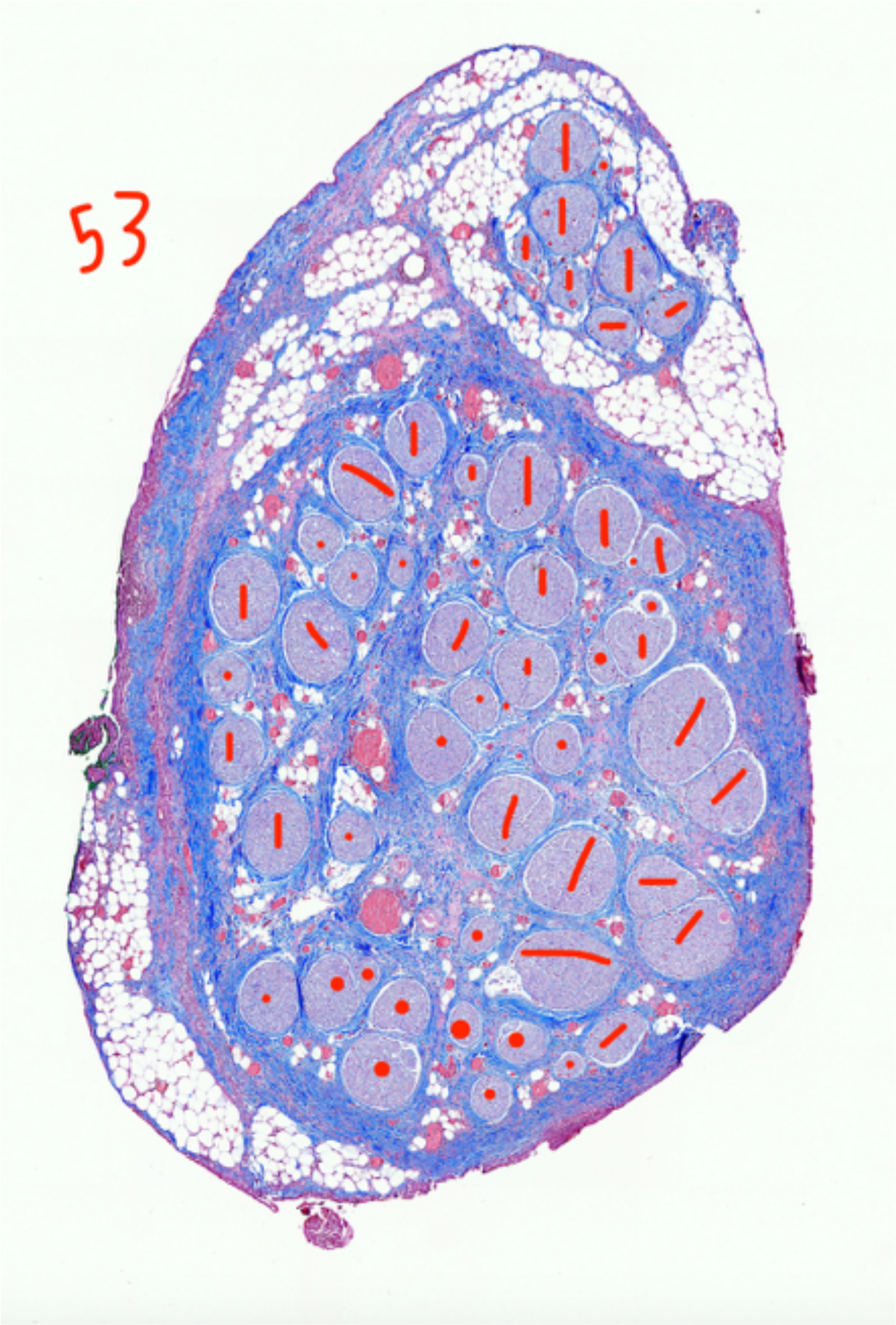


P858_LivaNova_43, mid shaft, fascicle numbers

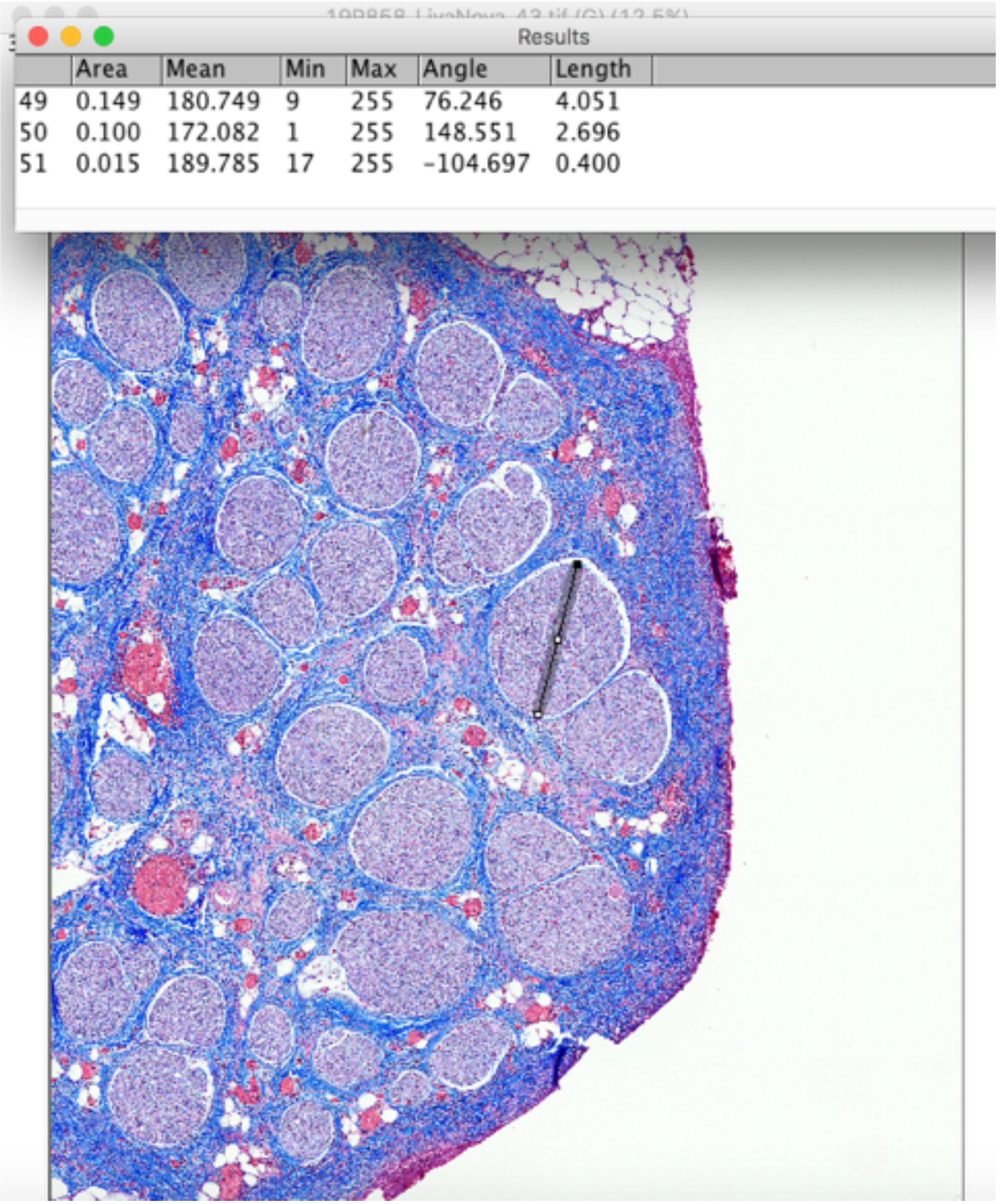


| | Area | Mean | Min | Max | Angle | Length |
|---|---|---|---|---|---|---|
| 49 | 0.149 | 180.749 | 9 | 255 | 76.246 | 4.051 |
| 50 | 0.100 | 172.082 | 1 | 255 | 148.551 | 2.696 |
| 51 | 0.015 | 189.785 | 17 | 255 | -104.697 | 0.400 |

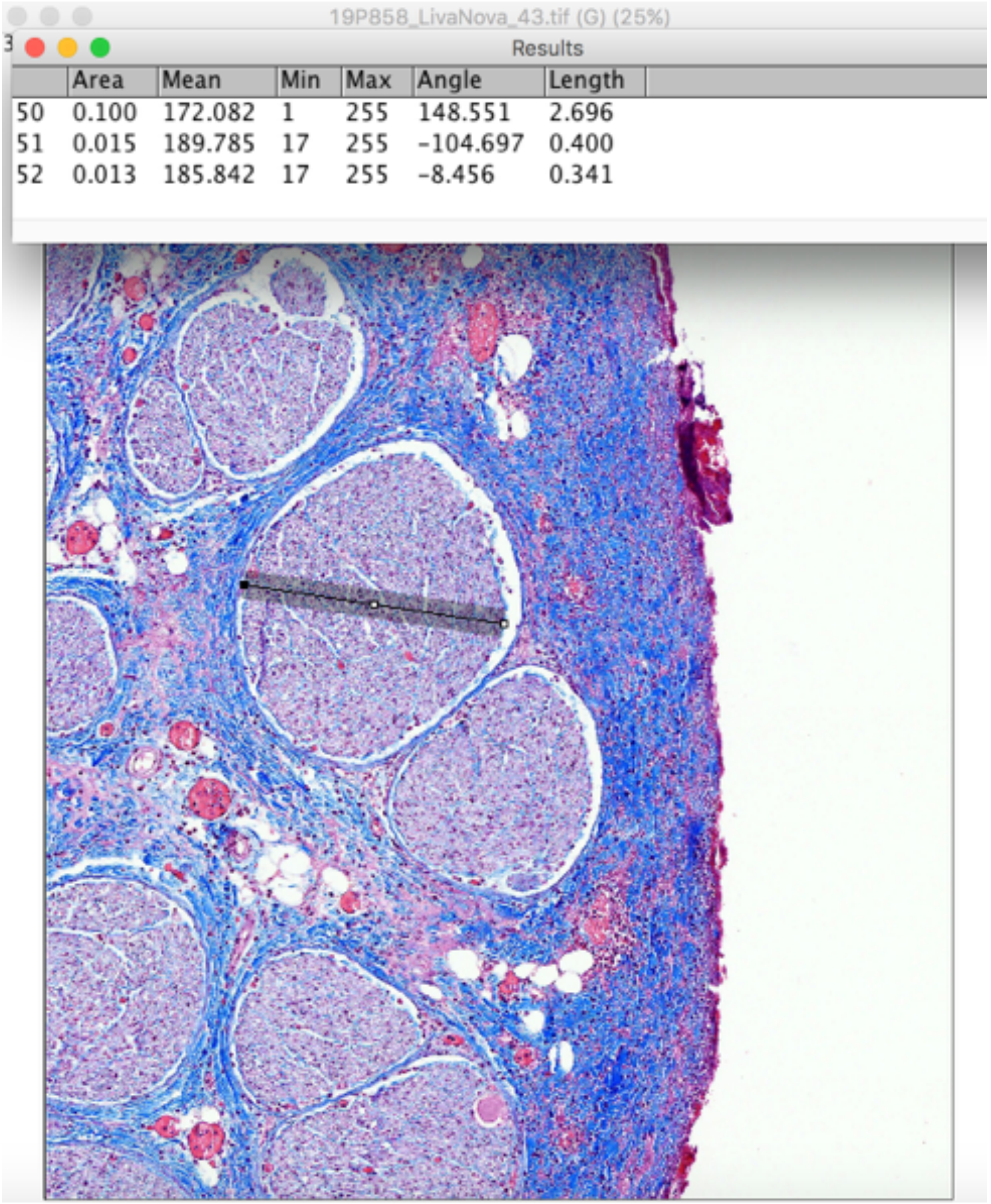


| | Area | Mean | Min | Max | Angle | Length |
|---|---|---|---|---|---|---|
| 50 | 0.100 | 172.082 | 1 | 255 | 148.551 | 2.696 |
| 51 | 0.015 | 189.785 | 17 | 255 | -104.697 | 0.400 |
| 52 | 0.013 | 185.842 | 17 | 255 | -8.456 | 0.341 |

P858_LivaNova_43, mid shaft, fascicle diameters

### Mid-VN

|  | Area | Mean | Min | Max | Angle | Length |
|---|---|---|---|---|---|---|
| 3 | 0.004 | 149.880 | 16 | 255 | -1.548 | 0.156 |
| 4 | 0.004 | 149.858 | 16 | 255 | -1.507 | 0.161 |
| 5 | 0.004 | 149.858 | 16 | 255 | -1.507 | 0.161 |

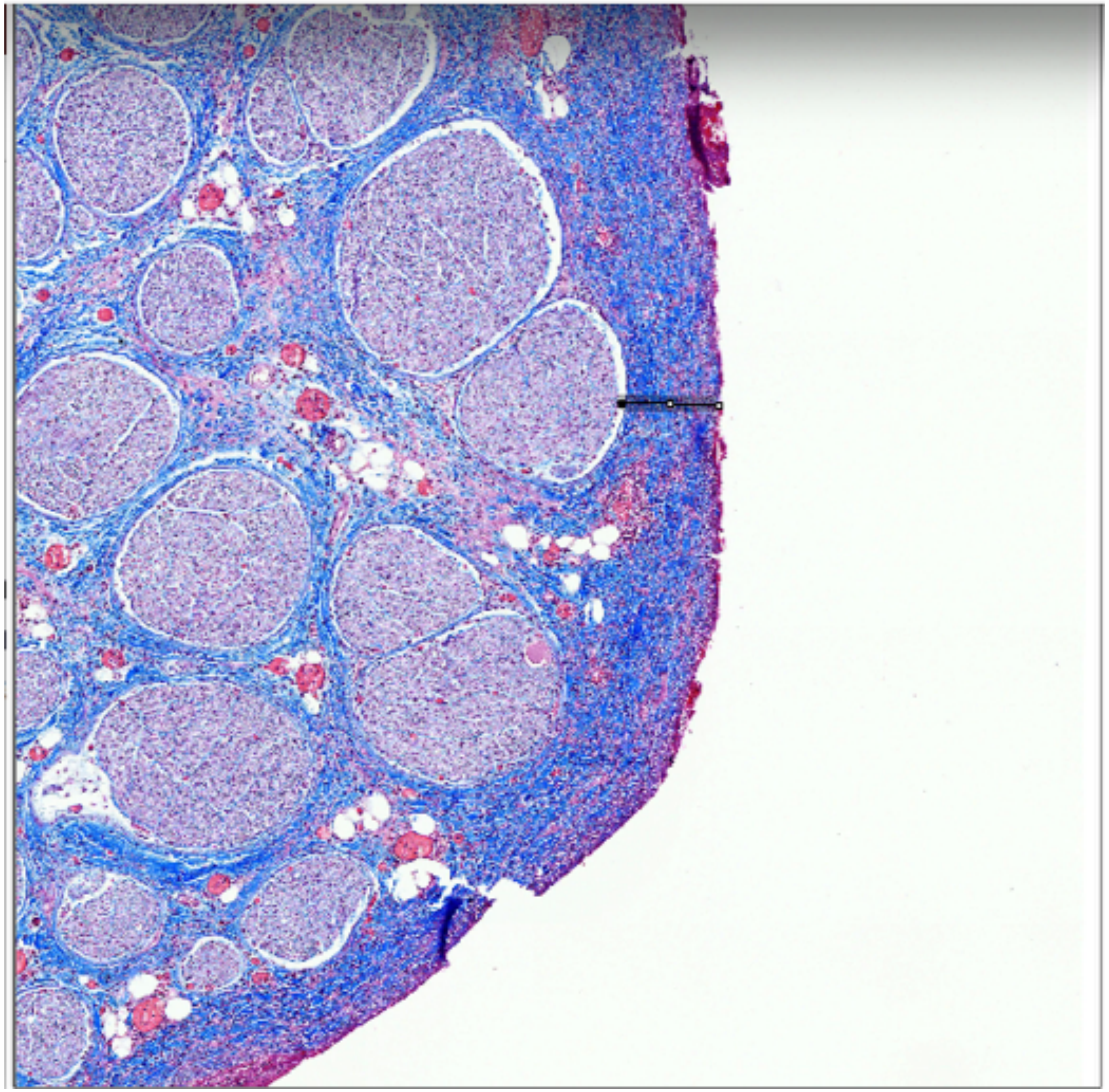

P858_LivaNova_43

Fascicle depth

Subject 10_P866

### Pre-nodose

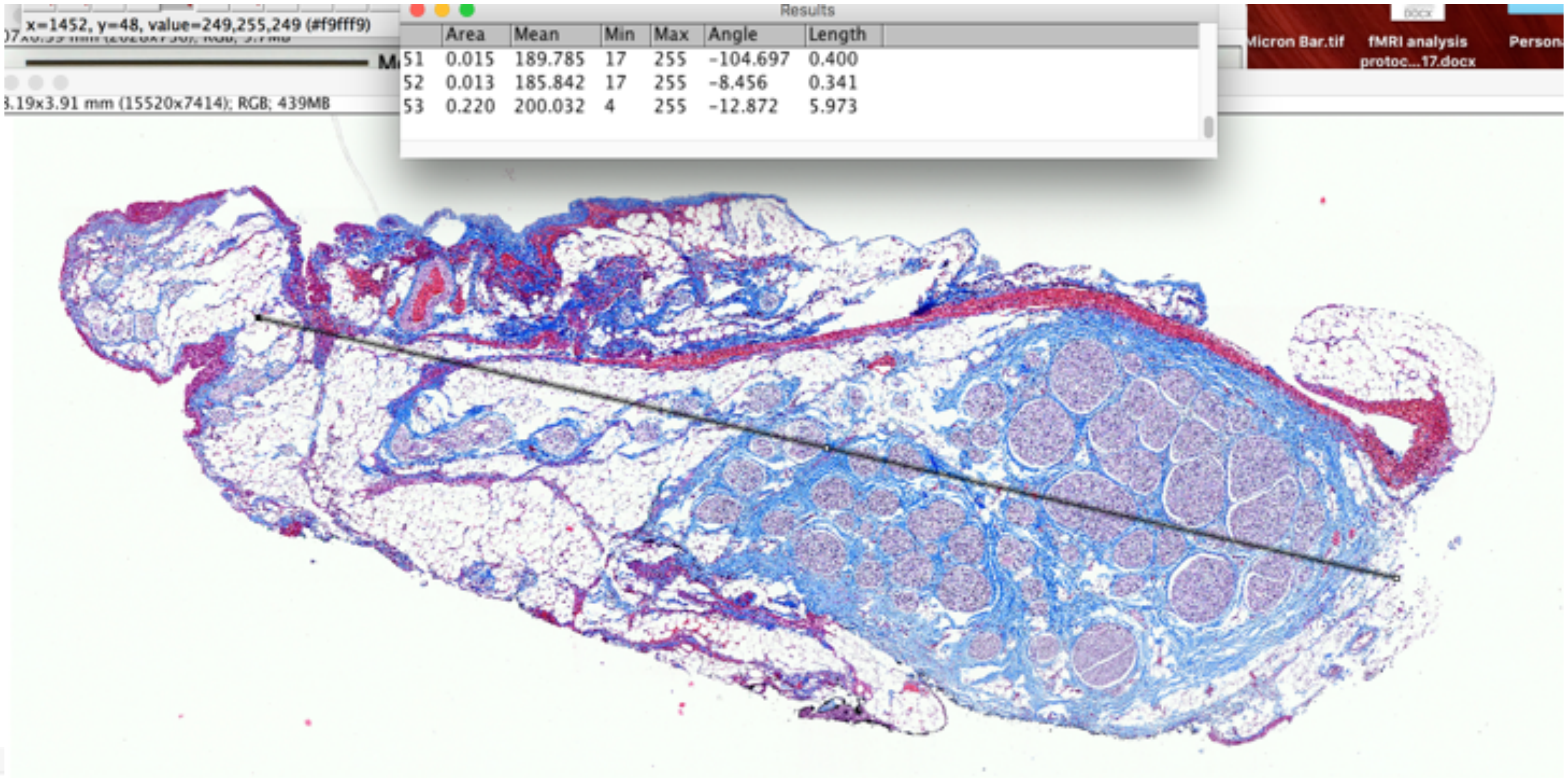


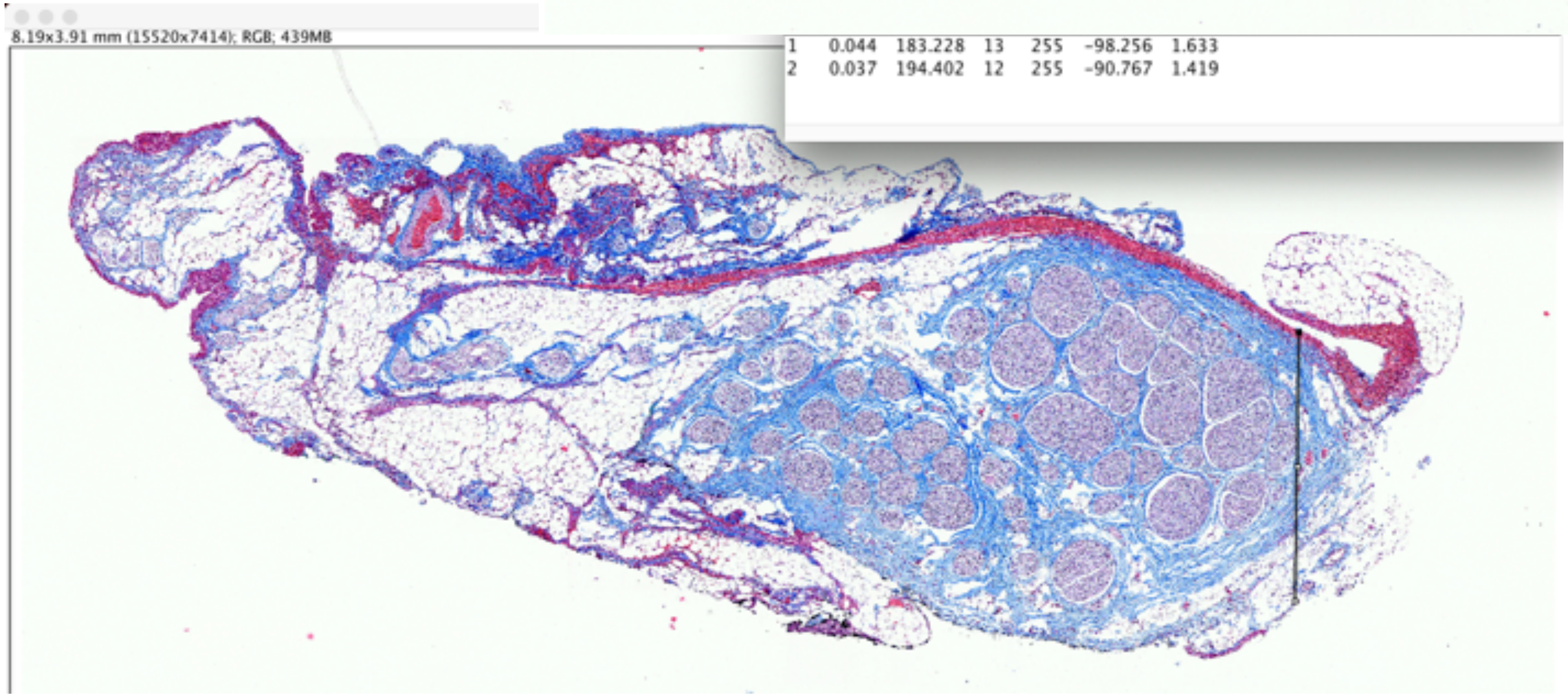


### P866_Nodose_152

Widest and narrowest diameter

Pre-nodose

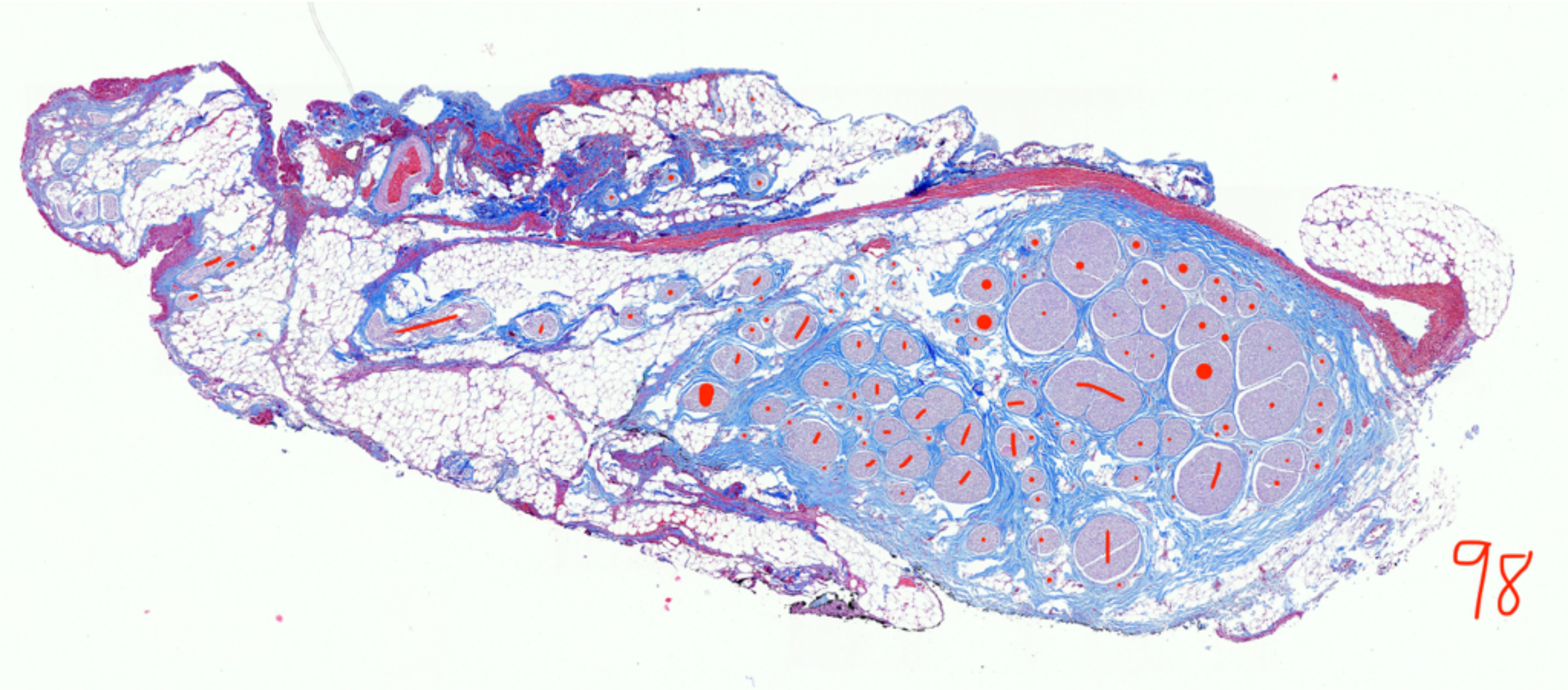


P866_Nodose_152

Fascicle count, 98

### Pre-nodose

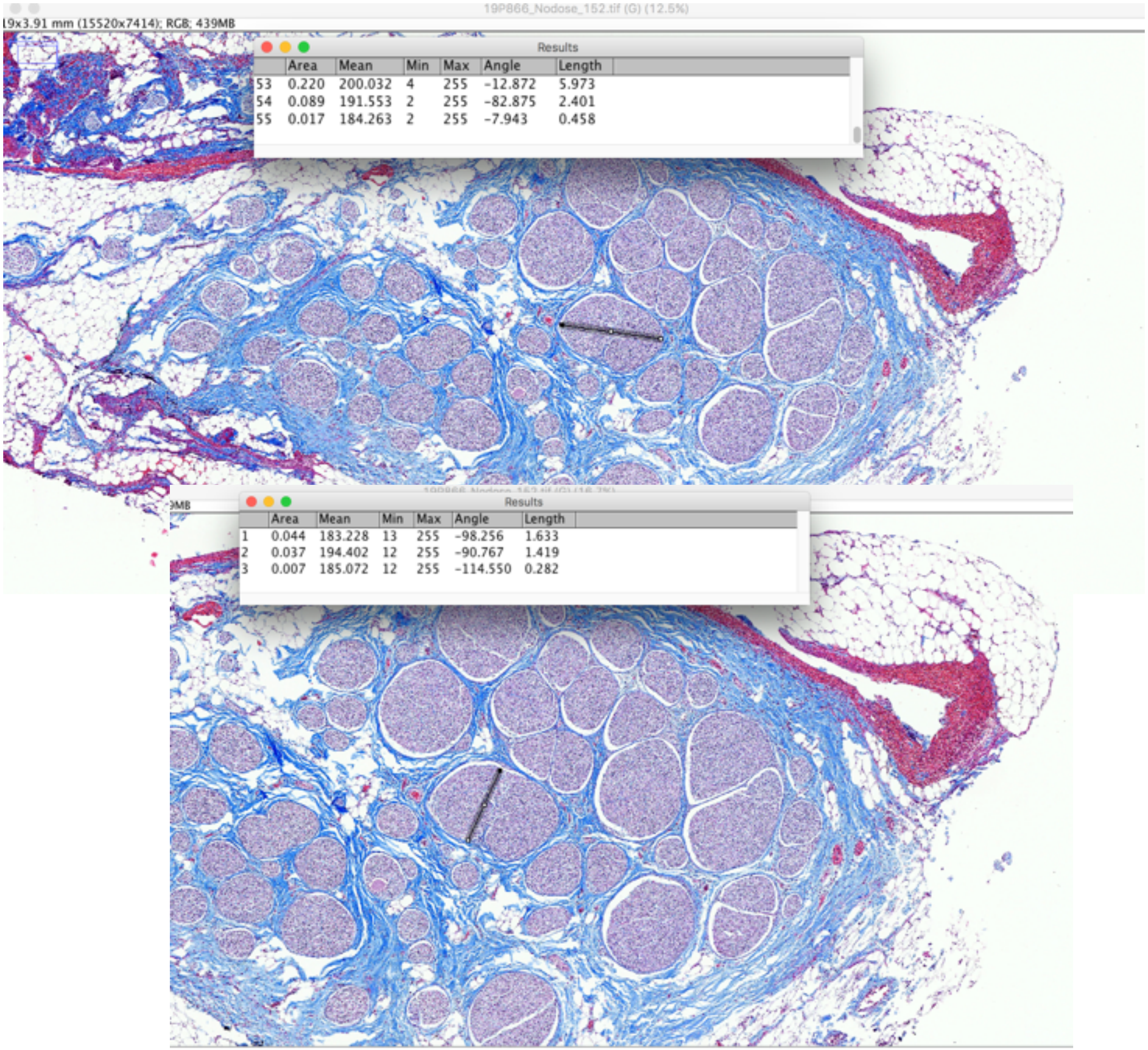


### P866_Nodose_152

Largest Fascicle_Widest and narrowest diameter

### Mid-VN

| | Area | Mean | Min | Max | Angle | Length |
|---|---|---|---|---|---|---|
| 56 | 0.013 | 185.229 | 7 | 255 | -78.417 | 0.347 |
| 57 | 0.119 | 175.308 | 5 | 255 | -44.263 | 3.250 |
| 58 | 0.121 | 181.479 | 0 | 255 | -39.005 | 3.288 |

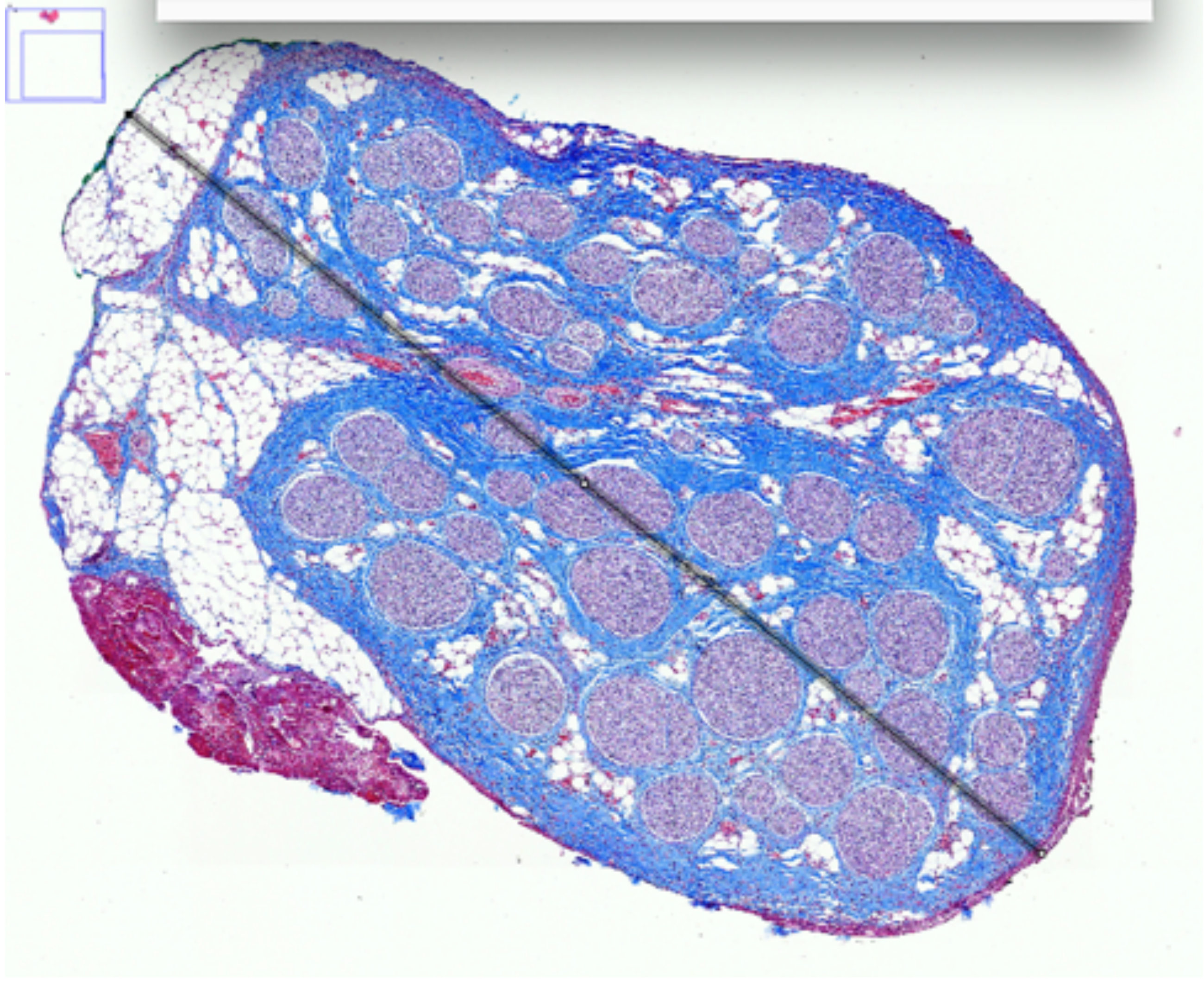

| | Area | Mean | Min | Max | Angle | Length |
|---|---|---|---|---|---|---|
| 1 | 8.946E-4 | 173.911 | 26.363 | 254.047 | -100.343 | 1.694 |
| 2 | 0.090 | 172.800 | 8.000 | 255.000 | -101.254 | 1.710 |

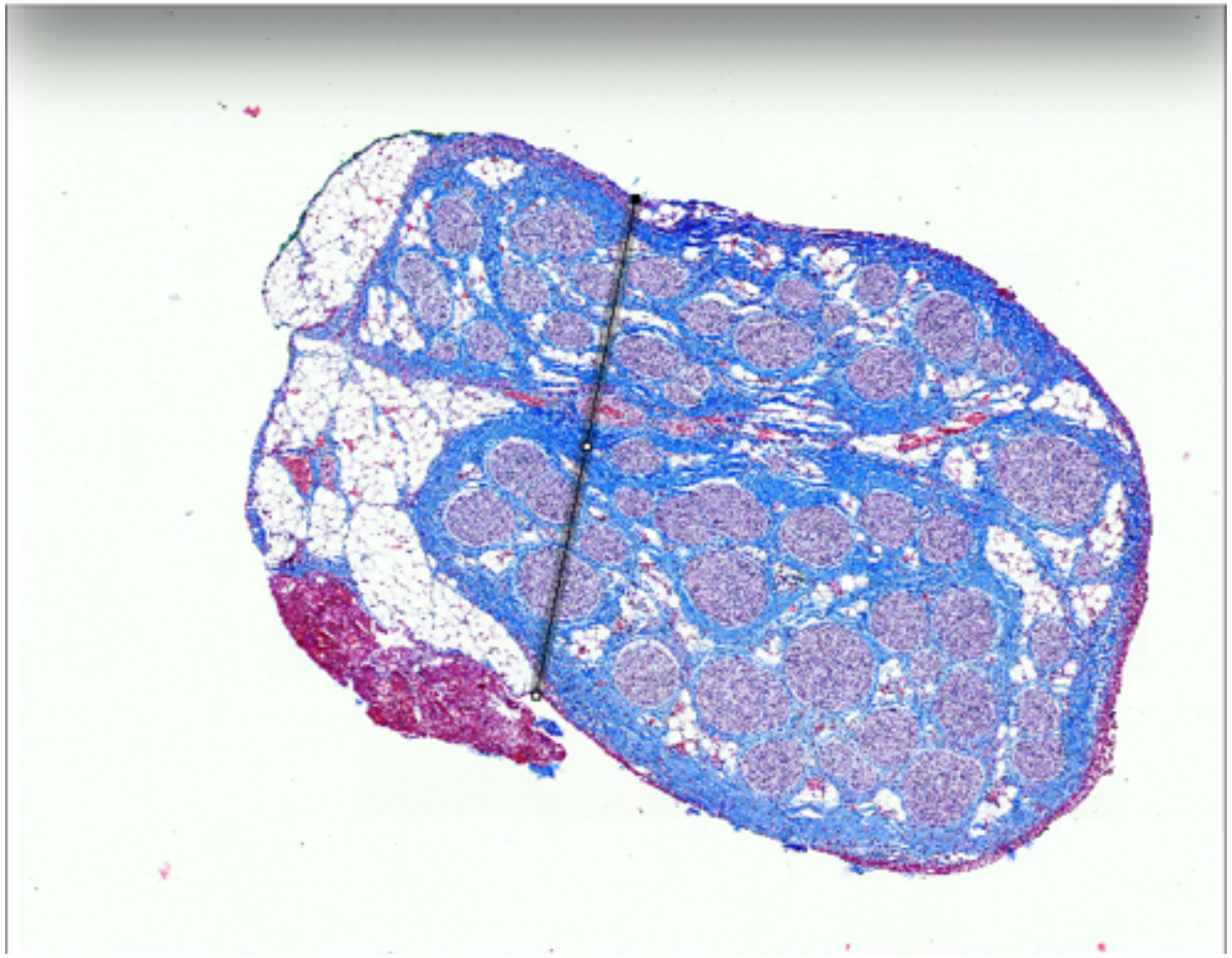

P866_VN1_A_154_blue

Widest and narrowest diameter

### Mid-VN

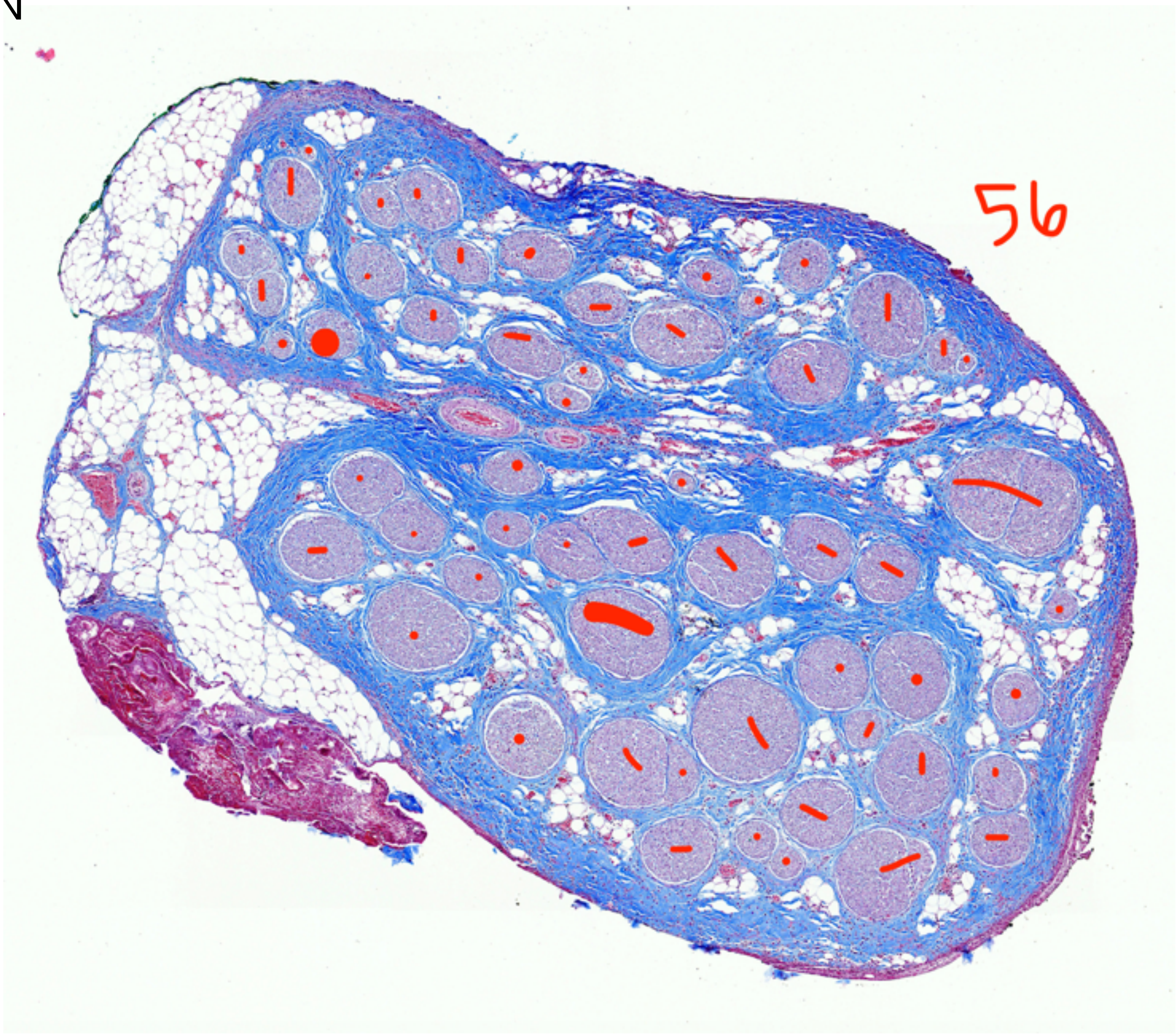


P866_VN1_A_154_blue

Fascicle count, 56

### Mid-VN

Results

| | Area | Mean | Min | Max | Angle | Length |
|---|---|---|---|---|---|---|
| 15 | 0.008 | 166.988 | 19 | 255 | 9.273 | 0.157 |
| 16 | 0.009 | 208.914 | 12 | 255 | 62.193 | 0.174 |
| 17 | 0.005 | 133.808 | 20 | 255 | 76.278 | 0.093 |

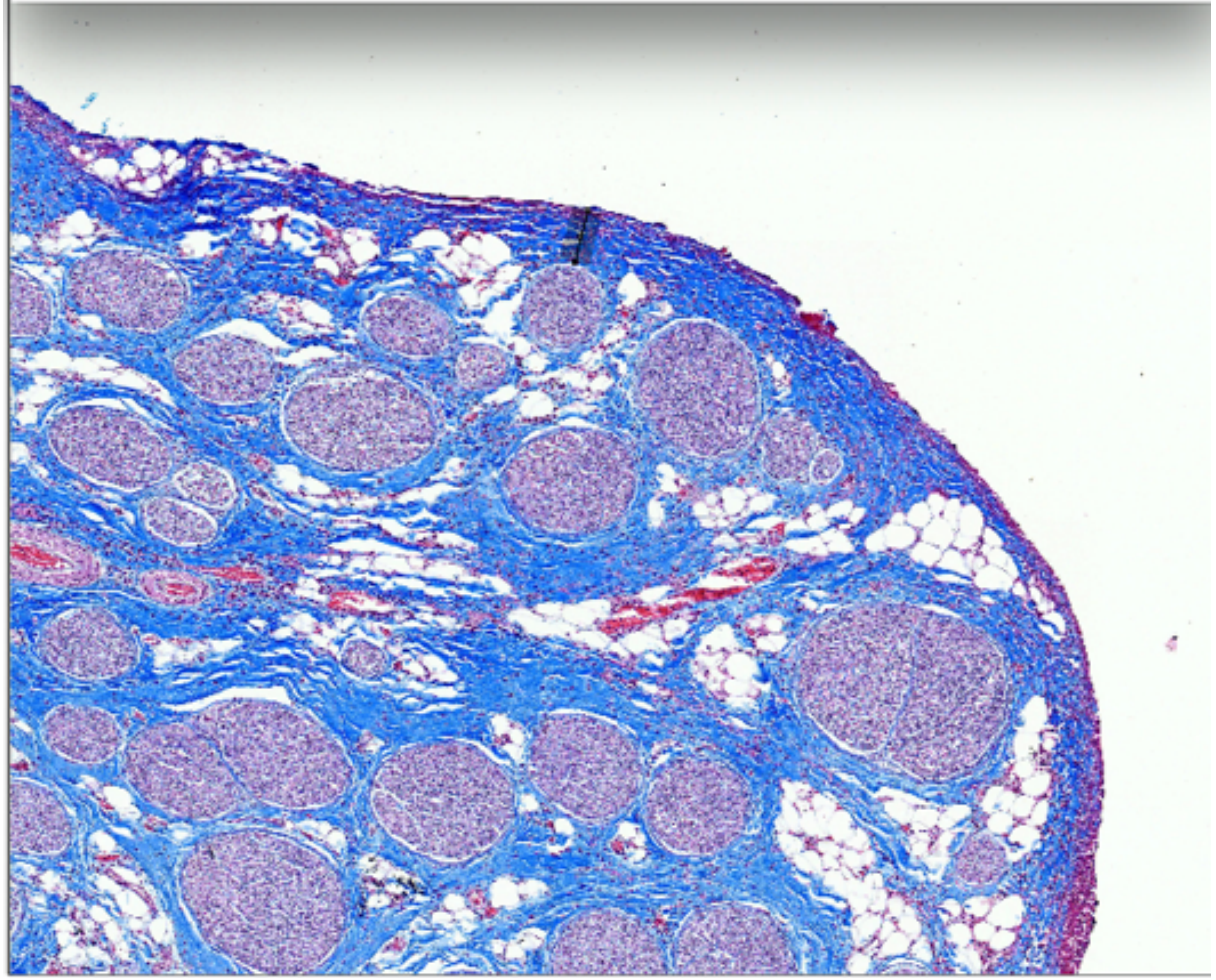

P866_VN1_A_154_blue

Fascicle depth

### Mid-VN

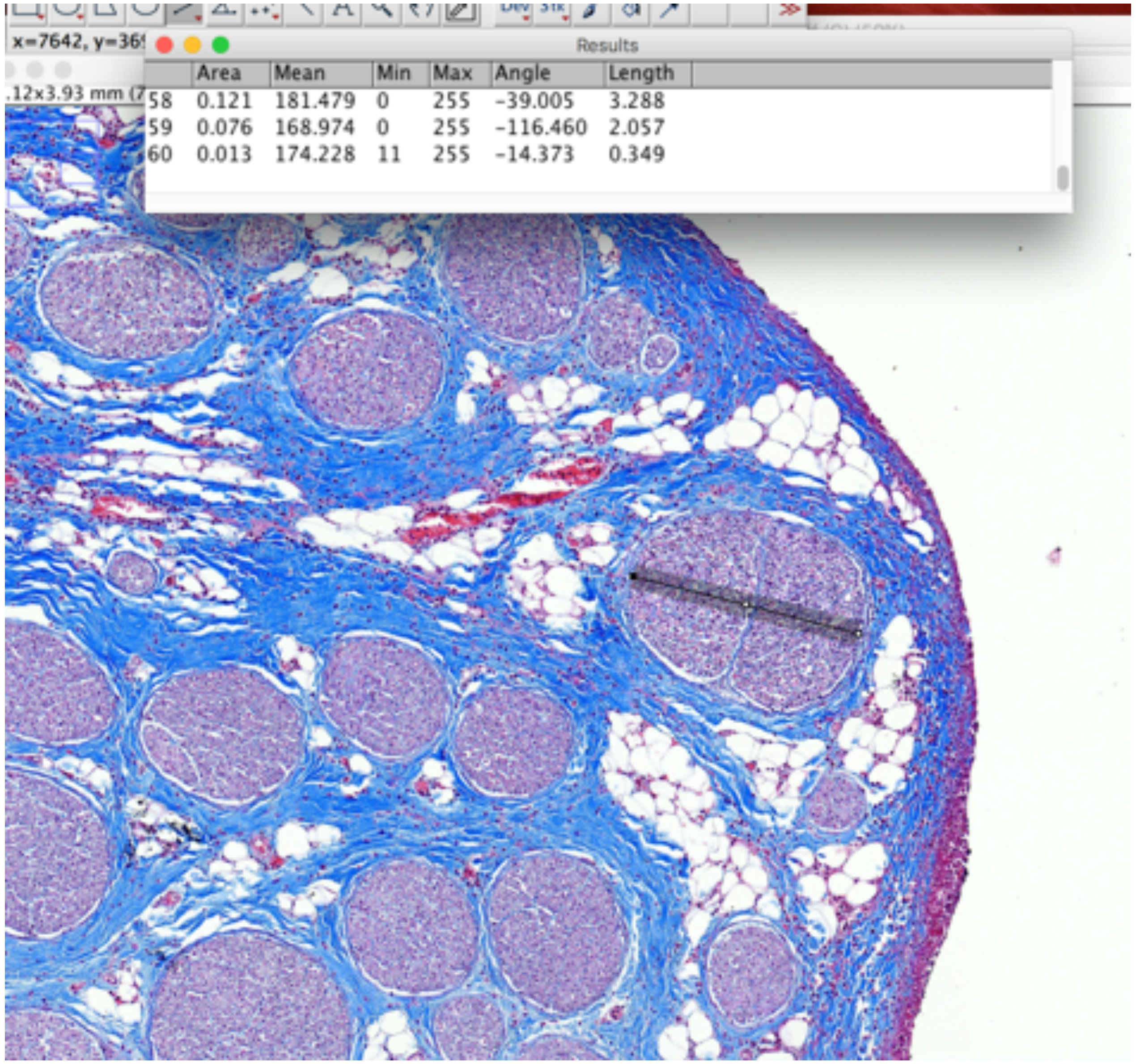


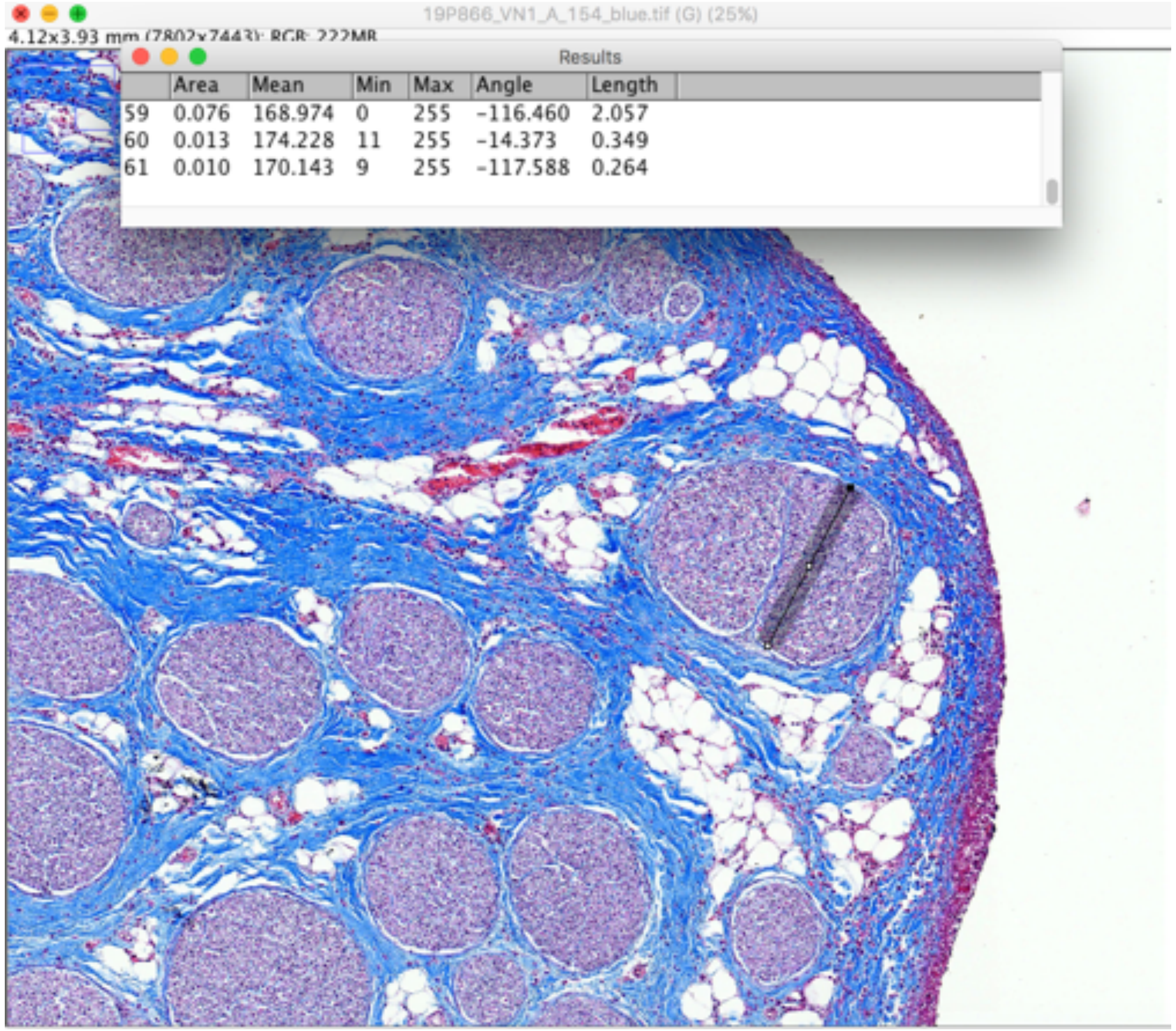


P866_VN1_A_154_blue

Largest fascicle_widest and narrowest diameter
